# Colchicine promotes atherosclerotic plaque stability independently of inflammation

**DOI:** 10.1101/2023.10.03.560632

**Authors:** Weizhen Li, Alexander Lin, Michael Hutton, Harkirat Dhaliwal, James Nadel, Julie Rodor, Sergey Tumanov, Tiit Örd, Matthew Hadden, Michal Mokry, Barend M Mol, Gerard Pasterkamp, Matthew P Padula, Carolyn L Geczy, Yogambha Ramaswamy, Judith C Sluimer, Minna U Kaikkonen, Roland Stocker, Andrew H Baker, Edward A Fisher, Sanjay Patel, Ashish Misra

## Abstract

Atherosclerosis is a chronic inflammatory disease which is driven in part by the aberrant *trans*-differentiation of vascular smooth muscle cells (SMCs). No therapeutic drug has been shown to reverse detrimental SMC-derived cell phenotypes into protective phenotypes, a hypothesized enabler of plaque regression and improved patient outcome. Herein, we describe a novel function of colchicine in the beneficial modulation of SMC-derived cell phenotype, independent of its conventional anti-inflammatory effects. Using SMC fate mapping in an advanced atherosclerotic lesion model, colchicine induced plaque regression by converting pathogenic SMC-derived macrophage-like and osteoblast-like cells into protective myofibroblast-like cells which thickened, and thereby stabilized, the fibrous cap. This was dependent on Notch3 signaling in SMC-derived plaque cells. These findings may help explain the success of colchicine in clinical trials relative to other anti-inflammatory drugs. Thus, we demonstrate the potential of regulating SMC phenotype in advanced plaque regression through Notch3 signaling, in addition to the canonical anti-inflammatory actions of drugs to treat atherosclerosis.

## Introduction

Atherosclerosis is the major cause of cardiovascular events, including myocardial infarctions and ischemic strokes, which are the leading cause of death worldwide. These acute complications are predominantly due to the rupture of unstable advanced atherosclerotic plaques (*1*). Vascular smooth muscle cells (SMCs) and their derivatives (SMC-derived cells) compose the majority of cells in advanced plaques, including the protective fibrous cap which provides mechanical strength and stability. The factors leading to plaque instability include thinning of the fibrous cap and the accumulation of inflammatory immune cells (*2*). No current therapies directly target the fibrous cap or SMC-derived cells to stabilize the plaque.

Recent clinical trials have focused on targeting inflammatory pathways. The anti-inflammatory drugs canakinumab and colchicine have shown reductions in cardiovascular events (*3-5*), whereas methotrexate has not shown the same efficacy (*6*). The precise underlying mechanisms accounting for these variations in anti-inflammatory action is poorly understood. Furthermore, it is unclear whether and if so, how anti-inflammatory interventions affect the fibrous cap. Contrary to expectations, anti-inflammatory (anti-IL-1β) therapy has a deleterious effect on SMC-derived plaque cells, causing a thinning of the fibrous cap (*7*). Thus, while anti-inflammatory agents have beneficial effects on immune cells, it is necessary to evaluate how these therapies affect non-immune cell populations.

Colchicine is the first anti-inflammatory agent to be FDA approved for cardiovascular disease patients. However, its success relative to other anti-inflammatory medication and its mechanism of action remains poorly understood. Colchicine’s anti-inflammatory effects on immune cells (*8-11*) are thought to facilitate the plaque stabilization seen in patients (*12*). Despite its success, it is unknown how colchicine influences SMC-derived plaque cells and their contribution to the fibrous cap. Herein, we describe a distinct athero-protective function for colchicine, independent of its anti-inflammatory effects. This role involves TGFβ and Notch3 signaling for the reversal of SMC-derived plaque cell phenotype. This previously unexplored athero-protective pathway for plaque stability reveals promising avenues for therapeutic intervention.

### Colchicine promotes the stability of advanced atherosclerotic plaques

To mimic colchicine intervention in advanced atherosclerotic cardiovascular disease patients in clinical trials (*4, 5*), apolipoprotein E null (*Apoe*^*-/-*^) mice were fed a Western diet (WD) for 24 weeks, with low dose colchicine (30μg/kg/day) intervention for the last 8 weeks (**Fig. 1A**). Treatment with colchicine had no influence on body mass (**Fig. S1A**). The bioavailability of colchicine was confirmed by the presence of the drug in arterial tissue and other organs as determined by liquid chromatography tandem mass spectrometry (LC-MS/MS) (Fig. 1B, S1B, C). To assess the potential anti-inflammatory effects of colchicine in atherogenic mice, our intervention with colchicine caused the differential expression of 59 proteins in the plasma (Fig. 1C, S1D). We observed the downregulation of multiple proteins associated with atherosclerotic disease severity, including A1AG1 (orosomucoid) (13) and SAMP (serum amyloid P) (*14*). Likewise, several proteins correlated with decreased atherosclerotic burden were upregulated by colchicine, including PROP (properdin) (*15*), APOH (apolipoprotein H) (*16*), and ALBU (albumin) (*17, 18*).

**Fig. 1.**
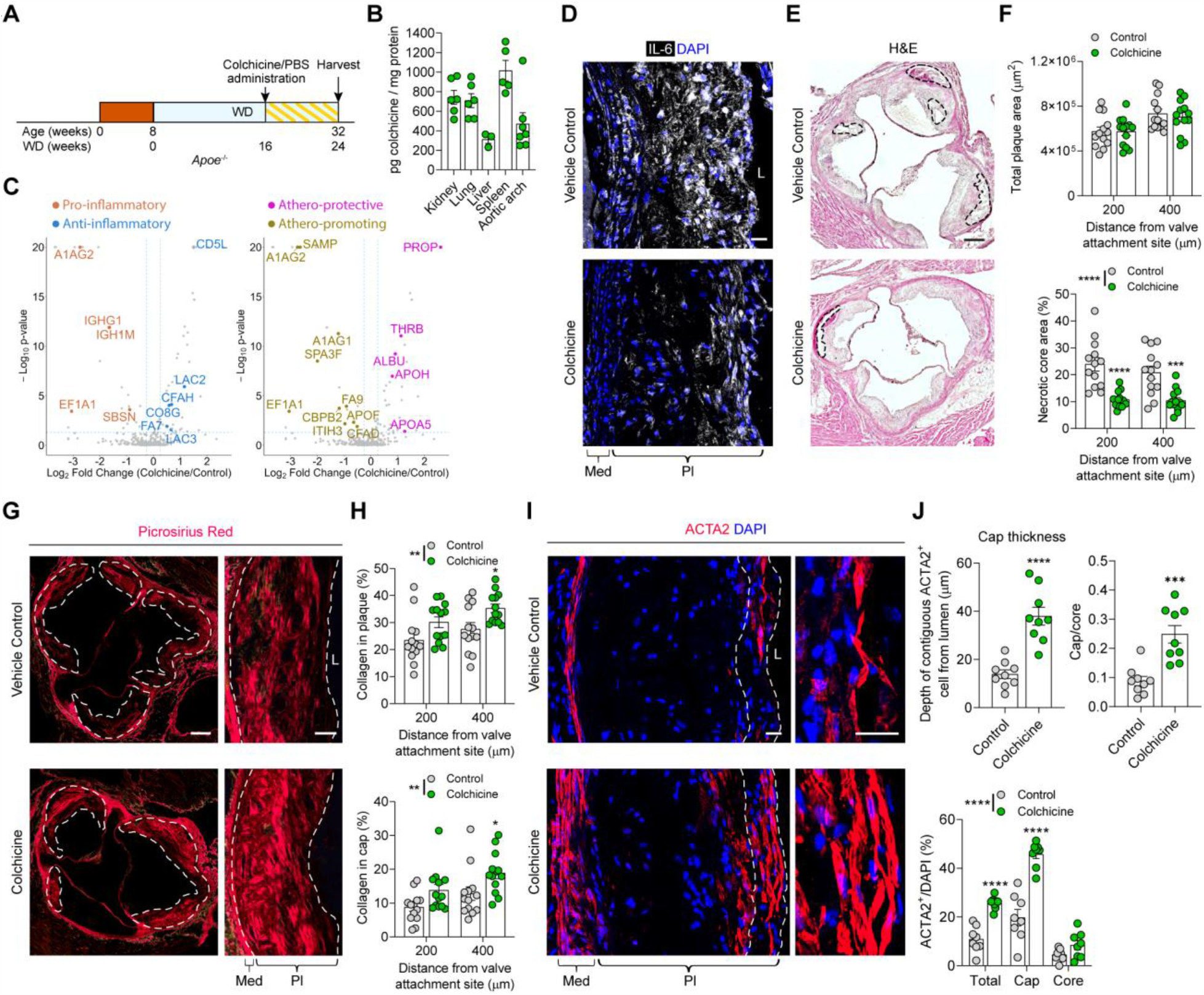
Colchicine treatment in advanced plaques promotes plaque stability. **A**, Atherosclerotic vessels were isolated from sex matched *Apoe*^*-/-*^ mice fed a WD for a total of 24 weeks, with colchicine (30μg/kg/day) or vehicle control intervention for the final 8 weeks. **B**, Colchicine was detected in the atherosclerotic aortic arch and other internal organs by LC-MS/MS. **C**, Volcano plots of proteins that are differentially expressed in plasma. Upregulated ‘anti-inflammatory’ proteins are shown in blue, downregulated ‘pro-inflammatory’ proteins are shown in orange. Upregulated ‘athero-protective’ proteins are shown in magenta, downregulated ‘athero-promoting’ proteins are shown in brown. **D**, Assessment of plaque inflammation by IL-6 staining showed reduced inflammation with colchicine treatment (representative of n = 8 for control and n = 8 for colchicine). **E**, Representative images of hematoxylin and eosin staining with necrotic core highlighted. **F**, No significant differences in total plaque area were observed (*p* = 0.7076), but colchicine intervention reduced necrotic core area (*p* < 0.0001). **G, H**, Representative Picrosirius Red staining (**g**) and quantification (**h**) demonstrates increased collagen deposition within the plaque (*p* = 0.0016) and fibrous cap (*p* = 0.0019) with colchicine intervention. **I, J**, Colchicine treatment increased the proportion of ACTA2^+^ cells within the lesion (*p* < 0.0001), fibrous cap to core ratio (*p* = 0.0003), and fibrous cap thickness (*p* < 0.0001), assessed by protein staining with cap area highlighted (**I**) and subsequent quantification (**J**). L, lumen; Med, tunica media; Pl, plaque. Scale bars: 200μm (**E, G**), 50μm (**G**, zoom), or 20μm (**D, I**). Data was analyzed using a two-way ANOVA with Sidak correction and multiple comparisons (**B, F, H, J** bottom) or a two-tailed unpaired t-test (**J** top). Individual dots represent biologically independent animals. Graphs show mean ± SEM. *P* values refer to two-way ANOVA between treatment conditions.

Atherosclerotic lesions were analyzed at the aortic root and brachiocephalic artery (BCA) (**Fig. S2A**). Colchicine’s anti-inflammatory properties were evident by reductions in intraplaque IL-6 and MCP-1 expression (**Fig. 1D, S2C**). Colchicine treatment did not significantly alter total lesion area (**Fig. 1E, F**) or intraplaque lipid levels (**Fig. S2D, E**). However, we observed a notable increase in indices of plaque stability, including a 50%-55% decrease in necrotic core size (**Fig. 1E, F**) and ∼28% and 48%-57% increase in collagen content within the lesion and fibrous cap region respectively (**Fig. 1G, H**). It is well established that ACTA2^+^ myofibroblast-like cells are the predominant source of collagen within the atherosclerotic plaque, and continuous ACTA2^+^ cell depth has been used to define the fibrous cap region (*19*). Colchicine intervention resulted in a 137% increase in ACTA2^+^ cells within the lesion, which were predominantly localized towards the luminal side (**Fig. 1I, J**). Analysis of the continuous ACTA2^+^ cells from the lumen demonstrated an average fibrous cap thickness of 38.0μm in the colchicine treated group, compared to 13.9μm in the vehicle control group (**Fig. 1I, J**). Thus, hereafter, 38.0μm was used to define the fibrous cap region in aortic root plaques. Taken together, these results indicate that colchicine improves indices of plaque stability, including increasing fibrous cap thickness, which may or may not be coupled to its anti-inflammatory effects.

We also investigated colchicine’s effects in the BCA, considering that different vascular regions have different mechanical stresses (*20*), and SMCs in different segments of the aorta originate from diverse developmental origins and exhibit varying responses to stimuli (*21*). BCA lesions also displayed indices of increased plaque stability (**Fig. S3**), as we observed a 146-187% increase in collagen content within the fibrous cap (**Fig. S3F, G**), 8.1μm increase in the thickness of continuous ACTA2^+^ cells from 9.3μm to 17.4μm (**Fig. S3H, I**), a 31-82% increase in ACTA2^+^ cells within the fibrous cap (**Fig. S3H, I**), and a 29-33% decrease in necrotic core size (**Fig. S3A, B**). Hereafter, 17.5μm (mean ACTA2^+^ depth in colchicine treated plaques) was used to define the fibrous cap region in BCA plaques. In comparison to the aortic root, there was an 18-32% decrease in Oil Red O^+^ staining area (**Fig. S3D, E**). These results suggest that the increase in athero-protective ACTA2^+^ cells following colchicine treatment is not dependent on the origin of SMCs.

### Colchicine intervention beneficially alters the phenotype of SMC-derived plaque cells

To investigate the origin of ACTA2^+^ cells within the thicker colchicine-induced fibrous cap, we generated hyperlipidemic SMC lineage tracing mice (*Apoe*^*-/-*^; *Myh11-*CreER^T2^; *ROSA26*^*mTmG/+*^ or *Apoe*^*-/-*^; *Myh11-*CreER^T2^; *ROSA26*^*tdTomato/+*^), given that medial SMCs are the dominant contributor to the athero-protective fibrous cap (*19*). These animals were induced with tamoxifen (1mg/day) for 5 days before WD initiation (**Fig. 2A**) to genetically mark mature Myh11^+^ cells and their progeny, with a high labelling efficiency as previously reported (*22*). We found there was a 101% increase in lineage^+^ ACTA2^+^ cells within the fibrous cap, and a 106% increase in the total lesion (**Fig. 2B, C**), indicating that colchicine increased the number of SMC-derived ACTA2^+^ cells for fibrous cap stability. Similar findings were demonstrated in the BCA (**Fig. S4**).

**Fig. 2.**
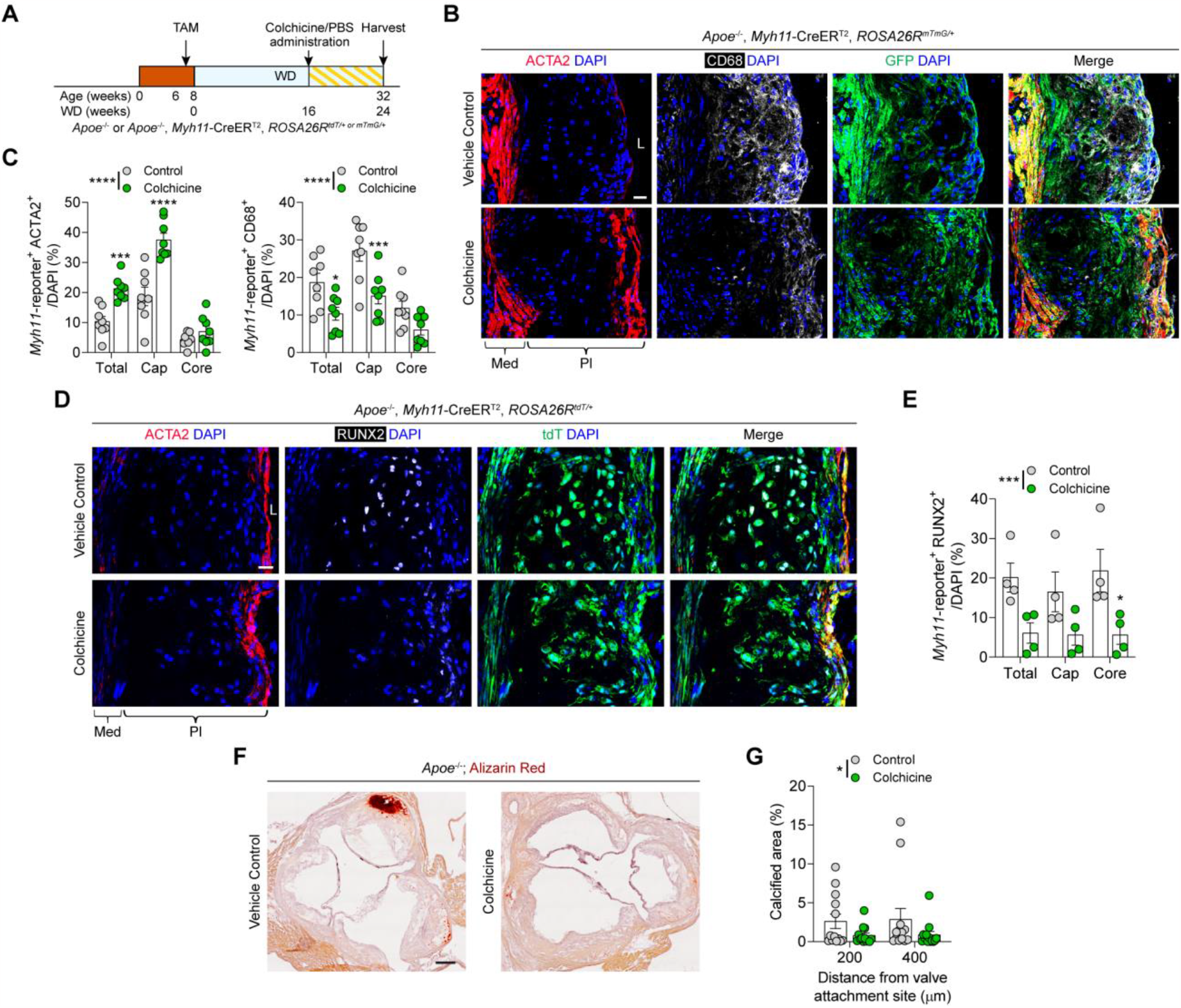
Colchicine increases the number of ACTA2^+^ SMC-derived cells within the lesion and ameliorates SMC *trans*-differentiation to macrophage-like and osteoblast-like phenotypes. **A**, SMC fate mapping mice (*Apoe*^*-/-*^, *Myh11-*CreER^T2^, *Rosa26*^*mTmG/+*^ *or Rosa26*^*tdTomato/+*^) were used for colchicine intervention studies. **B, C**, Colchicine intervention increased the proportion of SMC-derived ACTA2^+^ myofibroblast-like (*p* < 0.0001) cells and reduced the proportion of SMC-derived CD68^+^ macrophage-like cells (*p* < 0.0001) within the lesion, as assessed by marker protein staining (**B**) and subsequent quantification (**C**). **D, E**, Colchicine treatment also reduced the proportion of SMC-derived RUNX2^+^ osteoblast-like cells (*p* = 0.0003) within the lesion, as assessed by marker protein staining (**D**) and subsequent quantification (**E**). **F, G**, Representative Alizarin Red staining (**F**) and quantification (**G**) shows reduced intraplaque calcification (*p* = 0.0361) in the colchicine treatment group. TAM, tamoxifen; L, lumen; Med, tunica media; Pl, plaque. Scale bars: 20μm (**B, D**) or 200μm (**F**). Data was analyzed using a two-way ANOVA with Sidak correction and multiple comparisons. Individual dots represent biologically independent animals. Graphs show mean ± SEM. *P* values refer to two-way ANOVA between treatment conditions.

SMCs are highly plastic within the atherosclerotic plaque, and it has been shown that over 80% of SMCs lose their characteristic gene expression in the advanced plaque (*22*). These phenotypically switched SMCs have been shown to adopt a variety of plaque-burdening phenotypes, including a CD68^+^ macrophage-like phenotype which contributes to foam cell formation and plaque inflammation (*22-25*), and a RUNX2^+^ osteoblast-like phenotype which promotes intraplaque calcification (*26, 27*). Given that colchicine induces ACTA2 expression in SMC-derived cells, we investigated the impact of colchicine treatment on the phenotypic modulation of SMC-derived cells within the lesion. We observed numerous CD68^+^ SMC-derived macrophage-like cells in the 16 weeks WD advanced plaque prior to colchicine treatment (**Fig. S5**). Remarkably at 24 weeks WD, we found a 45% decrease in SMC-lineage^+^ CD68^+^ cells (**Fig. 2B, C**) and a 70% decline in SMC-lineage^+^ RUNX2^+^ cells within the total plaque after colchicine intervention (**Fig. 2D, E**). In accordance with the decline in calcification-driving cells, we observed a 70% decrease in calcified regions as indicated by Alizarin Red staining (**Fig. 2F, G**). These results suggest that colchicine mediated the regression of advanced atherosclerotic plaques by promoting an ACTA2^+^ phenotype in SMC-derived cells and reducing detrimental *trans*-differentiated SMC-derived cells.

### Colchicine regresses SMC-derived plaque burdening phenotypes via TGFβ signaling

Colchicine reduced the proportion of SMC-derived cells that had switched phenotypes. To identify whether colchicine was regressing these plaque-burdening phenotypes, we utilized a cholesterol loading assay that induces the *trans*-differentiation of SMCs to a macrophage-like phenotype (*28*). Human aortic SMCs (HASMCs) were loaded with cholesterol for 24 hours, followed by 24 hours of colchicine treatment (**Fig. 3A, S6A**). Cholesterol loading reduced the expression of SMC gene transcripts, whereas colchicine upregulated these SMC genes (**Fig. S6B**). For an unbiased approach, we performed RNA sequencing (RNA-seq), with principal component analysis revealing distinct clustering between treatment groups (**Fig. S7A**). We identified 701 genes regulated by cholesterol loading in SMCs (compared to the negative control) (**table S1**), and 964 genes were differentially expressed after colchicine intervention (compared to cholesterol loading) (**table S2**). Gene Ontology analysis showed that these genes are involved in cell adhesion, anatomical structure development and locomotion (**Fig. 3B, table S3**). Interestingly, the top SMC-related enriched Go Term was ‘Smooth Muscle Contraction’, suggesting that colchicine treatment may affect SMC contractility and reverse some of the changes induced by cholesterol loading. KEGG pathway analysis confirmed the changes of genes involved in cell adhesion and showed the enrichment of the TGFβ signaling pathway (**Fig. 3B, table S4**). 96 of the cholesterol regulated genes (14%) were regulated in the opposite direction after colchicine intervention, including known SMC contractile markers *ACTA2, TGLN, CNN1*, as well as several collagen genes *COL1A1, COL4A1, COL4A2*, and *COL12A1* (**Fig. 3C**). Gene Ontology analysis also revealed that the genes upregulated by cholesterol and repressed by colchicine (**Fig. S7B**) are involved in ‘inflammatory response’ and ‘signaling’, while the genes downregulated by cholesterol and re-activated by colchicine are linked to ‘cell adhesion’ (**Fig. S7C**).

**Fig. 3.**
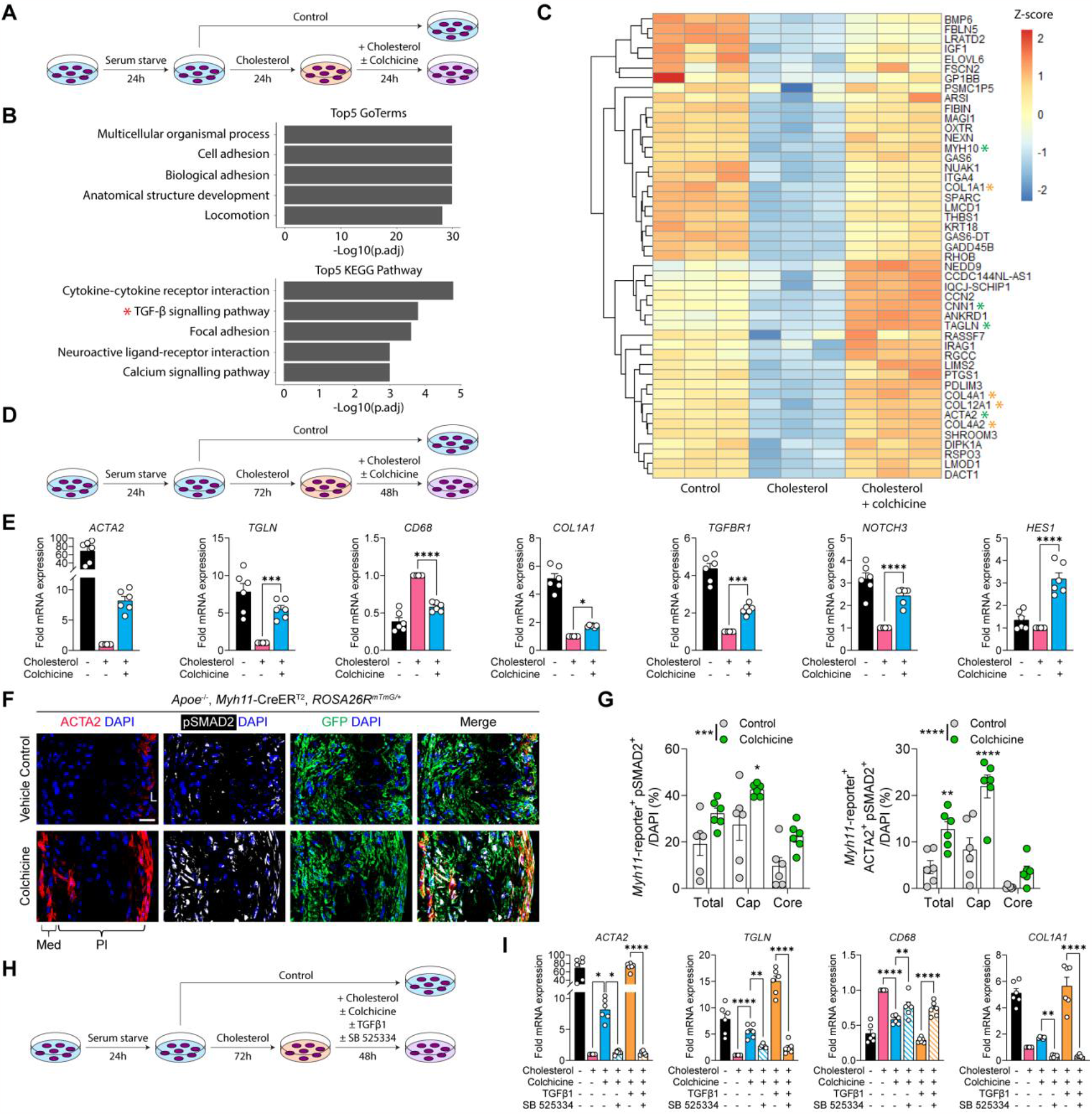
Colchicine reverts plaque burdening SMC phenotypes through the TGFβ pathway. **A**, Schematic demonstrating colchicine intervention after short-term macrophage-like *trans*-differentiation. Briefly, HASMCs were cholesterol loaded (10μg/mL) for 24 hours followed by treatment with colchicine (50nM) for 24 hours. **B**, Top 5 enriched Go-Terms (top) and KEGG pathway (bottom) for genes regulated by colchicine intervention compared to cholesterol loading. **C**, Heatmap of genes significantly downregulated by cholesterol and re-activated by colchicine intervention. Collagen related genes are marked with an orange asterisk, SMC related genes are marked with a green asterisk. **D**, Schematic demonstrating colchicine intervention after longer-term macrophage-like *trans*-differentiation. Briefly, HASMCs were cholesterol loaded (10μg/mL) for 72 hours followed by treatment with colchicine (50nM) for 48 hours. **E**, mRNA transcript levels of myofibroblast markers *ACTA2, TGLN, COL1A1*, macrophage marker *CD68*, TGFβ signaling mediator *TGFBR1*, and Notch signaling mediators *NOTCH3, HES1* were analyzed with qPCR. **F, G**, Colchicine intervention increases the proportion of cells expressing the downstream TGFβ mediator pSMAD2 as evident by protein staining (**F**) and quantification (**G**) (*p* = 0.0002 and *p* < 0.0001). **H**, Schematic demonstrating TGFβ-dependent colchicine intervention after macrophage-like *trans*-differentiation. Briefly, HASMCs were cholesterol loaded (10μg/mL) for 72 hours followed by treatment with colchicine (50nM) and/or TGFβ1 (2.5ng/mL) and/or the TGFβR1 inhibitor SB525334 (5μg/mL) for 48 hours. **I**, mRNA transcript levels of myofibroblast markers *ACTA2, TGLN, COL1A1*, and macrophage marker *CD68* were analyzed with qPCR. L, lumen; Med, tunica media; Pl, plaque. Scale bars: 20μm. Data was analyzed using a one-way ANOVA with Sidak correction and multiple comparisons (**E, I**) or a two-way ANOVA with Sidak correction and multiple comparisons (**G**). Individual dots represent independent experiments (**E, I**) or biologically independent animals (**G**). Graphs show mean ± SEM.

To examine colchicine’s effects after prolonged periods of cholesterol loading, HASMCs were loaded with cholesterol for 72 hours, followed by 48 hours of colchicine treatment (**Fig. 3D**), hereafter referred to as the cholesterol regression experiment. Relative to the cholesterol loaded cells, colchicine treatment increased transcript levels of the SMC markers *ACTA2, TGLN*, as well as the collagen isoform *COL1A1*, while decreasing the macrophage marker *CD68* (**Fig. 3E**). During atherogenesis, cholesterol is shuttled into the plaque by apolipoprotein B-containing lipoproteins (eg. low-density lipoproteins (LDLs), very low-density lipoproteins, and chylomicron remnants) (*29, 30*). Notably, colchicine was found to reduce the uptake of Dil-labeled oxidized LDL, which may contribute to the reductions in SMC-derived macrophage-like cells (**Fig. S9**).

As SMCs can adopt an osteoblast-like plaque-burdening phenotype (*31*), HASMCs were subjected to osteogenic medium treatment for 16 days, with colchicine intervention initiated from day 7 (**Fig. S6C**). Alizarin red staining confirmed osteoblast-like *trans*-differentiation over 16 days (**Fig. S6D**). Remarkably, we observed a trend of reduced calcification after colchicine treatment (**Fig. S6E**). These findings provide further support to our *in vivo* results, indicating that colchicine has the potential to reverse the macrophage-like and osteoblast-like phenotypes of plaque-burdening SMCs and restore them to an ACTA2^+^ phenotype.

In order to investigate the potential underlying mechanisms of colchicine-induced regression, we delved deeper into the involvement of TGFβ and Notch signaling pathways. These pathways have an established role in the maturation of SMCs (*32-35*). We found transcript level increases in the TGFβ signaling mediator *TGFBR1*, as well as the Notch pathway mediators *NOTCH3* and *HES1* (**Fig. 3E**). Within the plaque, we observed that colchicine increased the expression of TGFβR1 and the downstream protein pSMAD2 (**Fig. 3F, S9A**). Quantification of these SMC-derived cells demonstrated a 55% increase and 69% increase in SMC-lineage^+^ pSMAD2^+^ cells in the cap and plaque respectively, along with a 164% increase and 178% increase in SMC-lineage^+^ ACTA2^+^ pSMAD2^+^ cells (**Fig. 3G**).

To determine the potential association between TGFβ signaling and plaque cap thickness in humans, we collected human coronary plaque samples, which were classified histologically as stable or unstable plaques. We then analyzed TGFβ signaling in both the thick and thin cap regions of these plaques. Consistent with the findings from colchicine-treated mice, our observation revealed increases in the expression of TGFβR1 and phosphorylated SMAD2 (pSMAD2) in thick fibrous cap plaques compared to thin capped plaques (**Fig. S10A, B**). Additionally, we observed a concurrent increase in ACTA2 expression. TGFβ signaling thus appears to contribute to fibrous cap thickness in mature human coronary atherosclerotic plaques, thereby presenting a potential mechanism through which colchicine therapy may stabilize vulnerable atherosclerotic lesions in humans.

To confirm whether colchicine’s mechanism of action to induce ACTA2 expression is TGFβ dependent, we performed the cholesterol loading regression experiment in the presence of the TGFβR1 inhibitor SB525334 (**Fig. 3H**). Co-treatment with SB525334 prevented colchicine-mediated SMC gene induction and CD68 reduction (**Fig. 3I**). Immunostaining corroborated these findings (**Fig. S9B**), indicating that colchicine may regulate SMC phenotype through TGFβ mediated pathways.

### Colchicine regulates SMC-derived cell phenotype independent of anti-inflammatory pathways

To investigate whether the colchicine-mediated increase in ACTA2 expression in SMCs is independent of its anti-inflammatory actions, C57BL/6 mice were fed a normal chow diet and treated with colchicine for a period of 3 weeks. Colchicine treatment increased the expression of ACTA2, SMMHC, NOTCH3, as well as the number of pSMAD2^+^ cells within the healthy medial wall (**Fig. 4A, B, C, D**). These findings indicate that colchicine has the potential to induce the expression of favorable SMC-related proteins, even in the absence of atherosclerosis and its associated chronic inflammation.

**Fig. 4.**
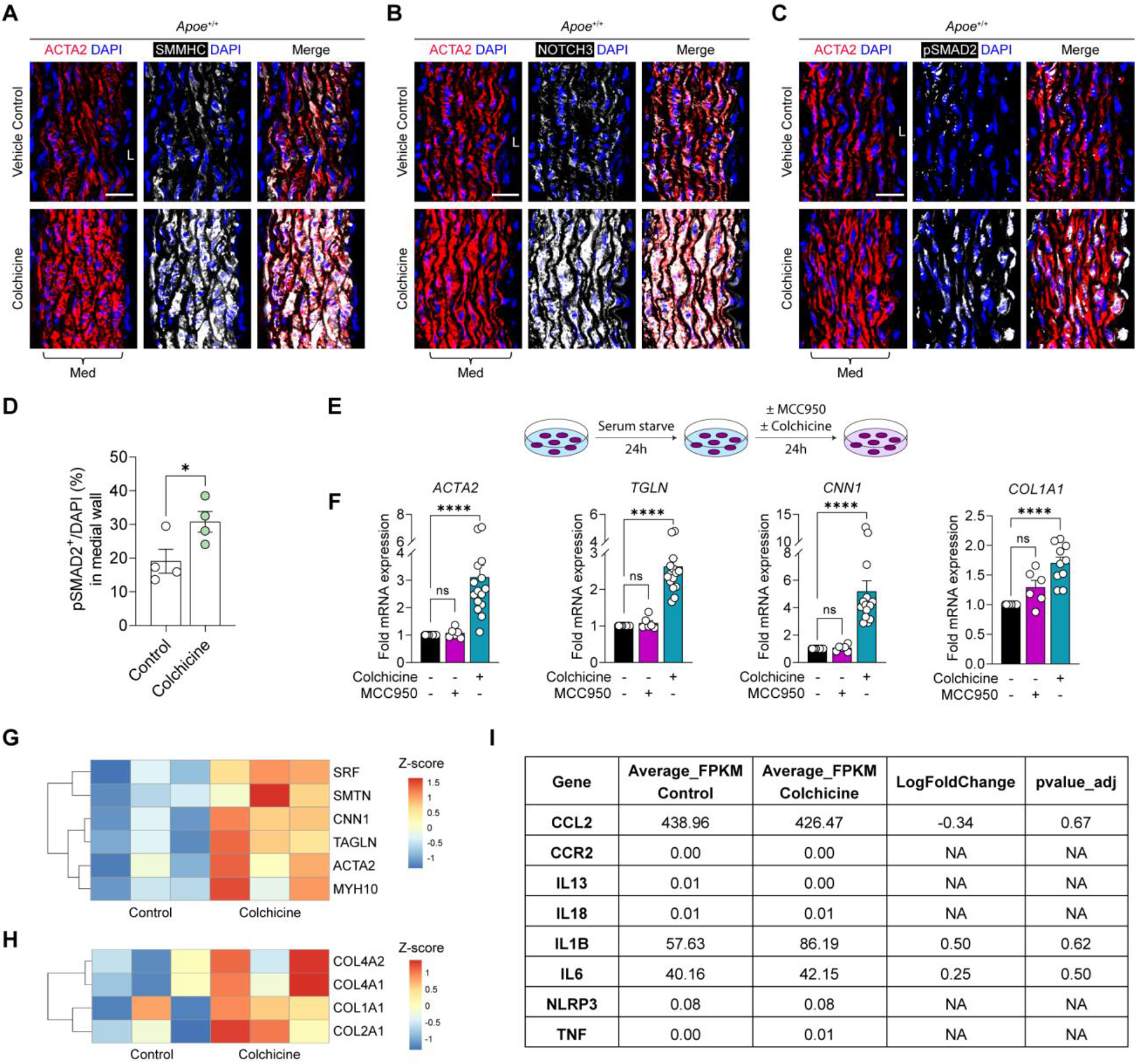
Colchicine influences the phenotype of SMC-derived cells independently of inflammation. **A, B**, Protein staining demonstrates that colchicine increases the expression of ACTA2 (**A, B**), SMMHC (**A**), and NOTCH3 (**B**) within the medial wall of healthy mice (representative of n = 4 for each group). **C, D**, Colchicine also induced pSMAD2 expression, as evident by protein expression (**C**) and subsequent quantification (**D**) (*p* = 0.0474). **E**, Schematic demonstrating colchicine and MCC950 loading of non-inflamed SMCs. Briefly, HASMCs were loaded with colchicine (50nM) or the NLRP3 inhibitor MCC950 (10μM) for 24 hours. **F**, mRNA transcript levels of the SMC markers *ACTA2, TGLN, CNN1* and the collagen isoform *COL1A1* were analyzed with qPCR. **G, H**, Heatmap of SMC related genes (**G**) and collagen genes (**H**) that were upregulated with colchicine treatment. **I**, No changes in inflammatory gene expression were observed after colchicine treatment of healthy HASMCs. L, lumen; Med, tunica media; NA, not applicable. Scale bars: 20μm. Data was analyzed using a two-tailed unpaired t-test (**D**) or a one-way ANOVA with Sidak correction and multiple comparisons (**F**). Individual dots represent biologically independent animals (**D**) or independent experiments (**F**). Graphs show mean ± SEM.

Colchicine is thought to inhibit the NLRP3 inflammasome and thereby suppress pro-inflammatory cytokine secretion (*9, 36*). To further demonstrate that colchicine has a direct effect on SMCs in the absence of inflammation, HASMCs were cultured with colchicine or the NLRP3 inhibitor MCC950 for 24 hours (**Fig. 4E**). Colchicine treatment upregulated SMC gene expression, while MCC950 did not have the same effect (**Fig. 4F**). This indicates that the observed phenotype with colchicine cannot be solely attributed to the suppression of the NLRP3 inflammasome.

To better understand the effects of colchicine alone on SMCs, we performed RNA-seq on the colchicine treated HASMCs (**Fig. S11A, B**). This identified 1249 differentially expressed genes (656 upregulated and 593 downregulated) (**table S5**). Gene Ontology analysis showed the enrichment of genes involved in anatomical structure morphogenesis, cell adhesion, and locomotion (**Fig. S11C, table S6**), similarly to the changes observed in colchicine intervention after cholesterol loading. In this condition, we showed the upregulation of several SMC contractility markers *CNN1, TAGLN, SMTN* and *SRF* (**Fig. 4G**), as well as the collagen genes *COL4A1, COL4A2* and *COL12A1* (**Fig. 4H**) which were also upregulated by colchicine in cholesterol-loaded SMCs. In contrast, we did not observe any change in expression of inflammatory markers (**Fig. 4I**). Taken together, these findings suggest that colchicine has a direct effect in promoting a favorable ACTA2^+^ phenotype in SMC-derived plaque cells, which is independent of its anti-inflammatory effects on the NLRP3 inflammasome.

### Notch signaling mediates the myofibroblast-like phenotype induced by colchicine

The data so far shows that colchicine induces TGFβ signaling and Notch3 signaling independently of anti-inflammatory pathways. This implies a potential synergistic interaction between TGFβ and Notch signaling, enhancing the impact of colchicine. To further explore the connections between TGFβ signaling and the Notch pathway, we investigated the binding of SMAD3 to Notch downstream mediators in human coronary artery smooth muscle cells (HCASMCs) using ChIP-Seq. The results demonstrate that SMAD3 binds to *RBPJ* and *HEY1* promoters and to an active enhancer element near *HES1* (**Fig. S12**). These findings further strengthen the notion of a regulatory crosstalk between the Notch pathway and the SMAD3-mediated TGFβ signaling cascade.

Indeed, in the advanced plaque regression model, colchicine treatment enhanced Notch3 expression in the fibrous cap, which showed substantial colocalization with ACTA2 (**Fig. 5A**). Moreover, there was a concurrent increase in downstream HES1 expression (**Fig. 5B**). A similar pattern was observed in human coronary plaques, where an increased Notch3 expression was observed in thick fibrous caps (**Fig. S10C**). To determine whether colchicine’s beneficial effects were Notch dependent, we cotreated HASMCs after cholesterol loading with colchicine and the γ-secretase inhibitor DAPT (*N*-[(3,5-difluorophenyl)acetyl]-L-alanyl-2-phenyl]glycine-1,1-dimethylethyl ester) to inhibit Notch signaling after cholesterol loading (**Fig. S13A**). Similar to SB525334, we observed that DAPT prevented the colchicine-mediated increase in SMC gene expression and reverted the beneficial change in gene expression of CD68 (**Fig. S13B**). This was further supported by immunofluorescent staining for ACTA2 (**Fig. S13C**), reinforcing the hypothesis that diminished Notch signaling promotes SMC phenotypic switching.

**Fig. 5.**
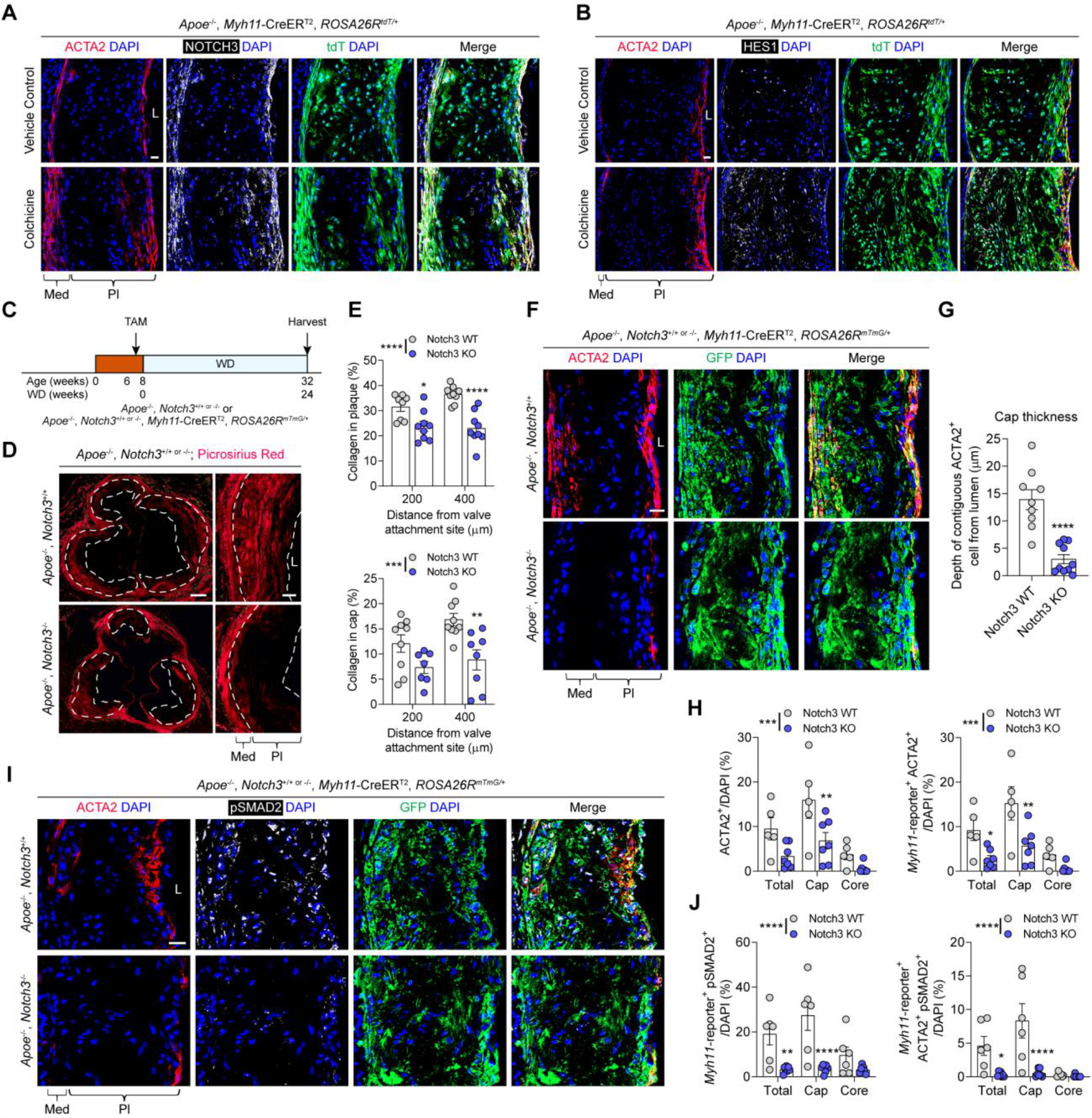
Notch3 signaling regulates indices of plaque stability and is essential for SMC-derived cells to acquire an ACTA2^+^ phenotype. **A**, Representative image showing Notch3 upregulation during colchicine mediated plaque regression. **B**, Representative image demonstrating downstream Notch mediator HES1 induction after colchicine treatment. **C**, To investigate the role of Notch3 in advanced plaques, *Apoe*^*-/-*^, *Notch3*^*-/-*^ or *Apoe*^*-/-*^, *Notch3*^*-/-*^, *Myh11-*CreER^T2^, *Rosa26*^*mTmG/+*^ mice were fed a WD for 24 weeks. **D, E**, Representative Picrosirius Red staining (**D**) and quantification (**E**) demonstrates reduced collagen deposition in the atherosclerotic plaque (*p* < 0.0001) and its fibrous cap (*p* = 0.0005) of *Notch3*^*-/-*^ mice. **F, G, H**, *Notch3* deletion also reduces the proportion of ACTA2^+^ cells (*p* = 0.0006), SMC-derived ACTA2^+^ myofibroblasts (*p* = 0.0001) and fibrous cap thickness (*p* < 0.0001), as assessed by protein staining (**F**) and associated quantification (**G, H**). TGFβ signaling was also impaired, indicated by a decline in SM-derived pSMAD2^+^ and ACTA2^+^ pSMAD2^+^ cells (**I, J**) (*p* < 0.0001 and *p* < 0.0001). L, lumen; Med, tunica media; Pl, plaque. Scale bars: 200μm (**D**), 50μm (**D**, zoom) or 20μm (**A, B, F, I**). Results are representative of n = 9 mice for each group (**A, B**). Data was analyzed using a two-way ANOVA with Sidak correction and multiple comparisons (**E, H, J**) or a two-tailed unpaired t-test (**G**). Individual dots represent biologically independent animals. Graphs show mean ± SEM. *P* values refer to two-way ANOVA between genotypes.

### Plaque stability is exacerbated in the absence of *Notch3*

To further define the specific role of Notch3 in SMC phenotypic switching, we generated quadruple transgenic mice (*Apoe*^*-/-*^; *Notch3*^*-/-*^; *Myh11-*CreER^T2^; *ROSA26*^*mTmG/+*^) that allow atherosclerotic induction and SMC lineage tracing in a *Notch3*^*-/-*^ background (**Fig. 5C**). *Apoe*^*-/-*^, *Notch3*^*-/-*^ mice were used where fate mapping was not required. Consistent with our hypothesis, Picrosirius red staining for collagen showed a 24-37% decrease in collagen content within the lesion and a 39-47% decrease in collagen content within the fibrous cap following *Notch3* deletion (**Fig. 5D, E**). Similar results were observed in BCA lesions (**Fig. S14A, B**). This was associated with a 65% decrease and a 57% decrease in ACTA2^+^ cells within the total lesion and fibrous cap respectively (**Fig. 5F, H**). Likewise, there was a 70% decrease and a 63% decrease in SMC-lineage^+^ ACTA2^+^ cells within the total lesion and fibrous cap respectively. Analysis of these ACTA2^+^ cells indicated a reduction in fibrous cap thickness from 13.9μm to 3.0μm (**Fig. 5G**). Consistent with this observation, we detected a decrease in TGFβ signaling in the plaques with *Notch3* deletion, as evident by reduced expression of TGFβR1 (**Fig. S14C**) and a decline in SMC-derived pSMAD2^+^ cells (**Fig. 5I, J**). Collectively, these findings indicate that the global knockout of *Notch3* causes a thinning of the athero-protective fibrous cap, a key feature suggesting reduced plaque stability.

### Colchicine-mediated regression is dependent on *Notch3*

Next, to demonstrate *in vivo* that colchicine’s effect on fibrous cap thickness was mediated Notch3, we fed *Apoe*^*-/-*^; *Notch3*^*-/-*^ or *Apoe*^*-/-*^; *Notch3*^*-/-*^; *Myh11-*CreER^T2^; *ROSA26*^*mTmG/+*^ mice a WD for 24 weeks, with colchicine (30μg/kg/day) or vehicle intervention from 16 weeks (**Fig. 6A**). Unlike *Apoe*^*-/-*^; *Notch3*^*+/+*^ animals, colchicine treatment had no influence on fibrous cap thickness as indicated by the unchanged number of ACTA2^+^ cells, contiguous ACTA2^+^ cell depth and collagen content (**Fig. 6D, E, F, G**). Furthermore, there were no changes in other indices of plaque stability including necrotic core area, Oil Red O^+^ area and calcified regions (**Fig. 6B, C, S15E, F, K, L**). This was similarly demonstrated in the BCA (**Fig. S15A, B, C, D, G, H, I, J**). These results indicate that colchicine’s plaque-stabilizing effects are dependent on Notch3 signaling.

**Fig. 6.**
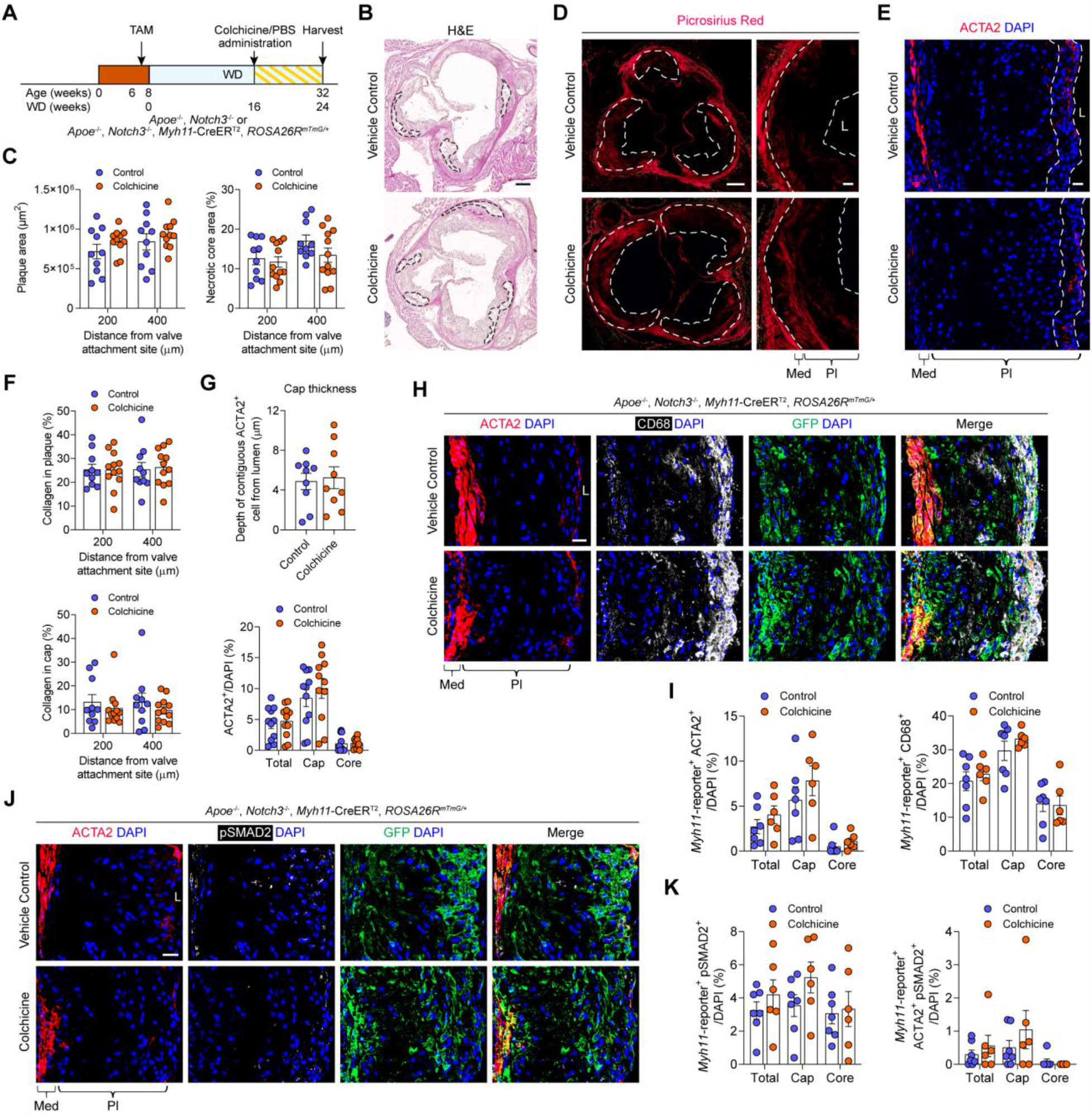
Colchicine intervention in *Apoe*^*-/-*^, *Notch3*^*-/-*^ mice does not improve indices of plaque stability. **A**, *Apoe*^*-/-*^; *Notch3*^*-/-*^ or *Apoe*^*-/-*^; *Notch3*^*-/-*^; *Myh11-*CreER^T2^; *ROSA26*^*mTmG/+*^ mice were used for colchicine intervention studies. **B**, Representative images of hematoxylin and eosin staining with necrotic core highlighted. **C**, No difference in plaque area (*p* = 0.1341) or necrotic core proportion (*p* = 0.1628) were observed after colchicine intervention. **D, F**, Colchicine intervention had no influence on collagen content within the plaque as assessed by Picrosirius Red staining (**D**) and associated quantification (**F**) within the fibrous cap (*p* = 0.2583) and lesion (*p* = 0.8656). **E, G**, Representative images of ACTA2 staining with cap area highlighted (**E**) and corresponding quantification (**G**) shows no change in ACTA2^+^ cell proportion (*p* = 0.4116) or fibrous cap thickness (*p* = 0.7829) with colchicine treatment. **H, I**, No difference in the proportion of SMC-derived ACTA2^+^ or CD68^+^ cells were observed after colchicine treatment, as demonstrated by protein staining (**H**) and subsequent quantification (**I**) (*p* = 0.1303 and *p* = 0.4031). **J, K**, Colchicine failed to induce pSMAD2 expression in SMC-derived cells as assessed by protein staining (**J**) and subsequent quantification (**K**) (*p* = 0.1361 and *p* = 0.2894). L, lumen; Med, tunica media; Pl, plaque. Scale bars: 200μm (**B, D**), 50μm (**D**, zoom) or 20μm (**E, H, J**). Data was analyzed using a two-way ANOVA with Sidak correction and multiple comparisons (**C, F, G** bottom, **I, K**) or a two-tailed unpaired t-test (**G** top). Individual dots represent biologically independent animals. Graphs show mean ± SEM. *P* values refer to two-way ANOVA between treatment conditions.

We observed that colchicine treatment induced TGFβ signaling and reduced the number of SMC-lineage^+^ CD68^+^ cells when Notch3 was present. Furthermore, in *Notch3*^-/-^ mice treated with colchicine, we did not observe any change in fibrous cap thickness. Therefore, we investigated whether colchicine-induced TGFβ signaling and its beneficial effects on SMC-derived cell fate were impaired in the absence of Notch3. Colchicine intervention had no influence on the proportion of SMC-lineage^+^ ACTA2^+^ cells, and there was no change in the percentage of SMC-lineage^+^ CD68^+^ cells (**Fig. 6H, I**). Consistent with this notion, we observed no changed in TGFβR1 expression (**Fig. S15M**), and no increase in the proportions of SMC-lineage^+^ pSMAD2^+^ cells or SMC-lineage^+^ ACTA2^+^ pSMAD2^+^ cells (**Fig. 6J, K**). In conclusion, these findings suggest that colchicine has the potential to enhance indices of plaque stability and thus reduce cardiovascular disease by positively modulating the phenotype of SMC-derived plaque cells through Notch3 dependent mechanisms.

## Discussion

The CANTOS trial provided proof of principle evidence that targeting inflammation lowers atherosclerotic cardiovascular events independently of lipid levels (*3*). Subsequent clinical trials with colchicine further supported this notion (*4, 5*), whereas methotrexate has not yielded the same results (*6*). However, canakinumab was rejected by the FDA for cardiovascular patients. Furthermore, the success of IL-1β suppression has been called into question with preclinical studies suggesting that suppressing the IL-1 axis may prevent fibrous cap stabilization (*7, 19*). In a similar plaque regression model with anti-IL-1β antibodies, Gomez *et al*. (*7*) showed a decline in ACTA2^+^ SMC-derived cells and collagen content in the fibrous cap, accompanied by an increased accumulation of macrophages. Our findings with colchicine directly contrast with these results as we show the reversal of detrimental SMC-derived cell phenotypes, which thickened the fibrous cap. Given that both therapies suppress IL-1β mediated pathways, this suggests that colchicine possesses additional independent protective effects that canakinumab lacks.

The LoDoCo2 trial showed that colchicine intervention led to a 31% relative risk reduction in primary composite cardiovascular events (*4*), leading to its recent FDA approval for atherosclerotic cardiovascular disease. While colchicine’s anti-inflammatory effects in immune cells have been widely recognized (*8-11*), the present study reveals an additional remarkable effect. Colchicine also has the exceptional capacity to revert deleterious SMC-derived plaque cell phenotypes back into a protective ACTA2^+^ phenotype independently of inflammatory pathways, increasing fibrous cap thickness and overall plaque stability. The beneficial modulation of SMC-derived plaque cells has been hypothesized but has remained elusive to date. These dual-action benefits likely explain the success of colchicine relative to other anti-inflammatory therapies, and strongly suggests that targeting plaque cell *trans-*differentiation holds promise for improving patient outcomes.

In order to achieve these remarkable effects on SMC-derived cell phenotype, our study revealed that colchicine activates both TGFβ and Notch3 signaling. There is cumulative evidence suggesting that the impairment of TGFβ signaling promotes detrimental SMC phenotypic switching and disease burden (*37, 38*), but no pharmacological agent has yet been shown to augment this pathway. Similarly, deficiencies in SMC Notch signaling have been shown to reduce fibrous cap thickness (*39*). We identify that the Notch3 receptor is responsible for mediating these effects. Crucially, colchicine-mediated regression does not occur in the absence of *Notch3*, emphasizing the importance of Notch3 mediated pathways in atherosclerosis and the tractability of targeting such mechanisms.

In summary, our findings indicate that colchicine is protective in SMC-derived plaque cells in advanced atherosclerosis, which likely contributed to its success in clinical trials (*4, 5*). This effect is independent of its other beneficial anti-inflammatory properties. Colchicine promoted fibrous cap thickening and collagen deposition by converting detrimental SMC-derived plaque cell phenotypes into protective ACTA2^+^ myofibroblast-like cells, in a Notch3 dependent manner. In light of recent clinical trials (*3-6*) and emerging fate mapping studies of SMCs (*7*), our results emphasize the importance of non-immune cells in anti-inflammatory therapy and the need for additional measures of atherosclerotic cardiovascular disease risk beyond the current gold standard indicators of high sensitivity C-reactive protein and low density lipoprotein cholesterol levels. Ultimately, our data suggests that SMC-derived plaque cell phenotype could be targeted through Notch3 by therapeutics in the future for improving plaque stability and patient outcome, a paradigm which has yet to be established in clinical practice.

## Acknowledgments

The authors acknowledge the Australian Genome Research Facility in performing RNA sequencing. We thank Shaun Jackson for critical review of the manuscript.

## Author contributions

Conceptualization: AM

Methodology: AM

Investigation: WL, AL, MHutton, HD, JN, JR, ST, TÖ, MHadden, MM, BMM, MPP, AM

Visualization: WL, AL, MHutton, AM

Funding acquisition: JCS, AHB, SP, AM

Project administration: AM

Supervision: GP, CLG, YR, JCS, MUK, RS, AHB, EAF, SP, AM

Writing – original draft: WL, AL, AM

Writing – review & editing: WL, AL, CG, JCS, MUK, RS, AHB, EAF, AM

## Data and materials availability

Data is available in the main text or the supplementary materials. The RNA-seq datasets are deposited in the Gene Expression Omnibus (GEO) repository (accession numbers will be provided upon manuscript acceptance for publication).

## Supplementary Materials

Materials and Methods

Figs. S1 to S15

Tables S1 to S4

## References and Notes

1. M. J. Davies, A. Thomas, Thrombosis and Acute Coronary-Artery Lesions in Sudden Cardiac Ischemic Death. New England Journal of Medicine 310, 1137–1140 (1984).

2. R. Virmani, F. D. Kolodgie, A. P. Burke, A. Farb, S. M. Schwartz, Lessons From Sudden Coronary Death. Arteriosclerosis, Thrombosis, and Vascular Biology 20, 1262–1275 (2000).

3. P. M. Ridker et al., Antiinflammatory Therapy with Canakinumab for Atherosclerotic Disease. New England Journal of Medicine 377, 1119–1131 (2017).

4. S. M. Nidorf et al., Colchicine in Patients with Chronic Coronary Disease. New England Journal of Medicine 383, 1838–1847 (2020).

5. J.-C. Tardif et al., Efficacy and Safety of Low-Dose Colchicine after Myocardial Infarction. New England Journal of Medicine 381, 2497–2505 (2019).

6. P. M. Ridker et al., Low-Dose Methotrexate for the Prevention of Atherosclerotic Events. New England Journal of Medicine 380, 752–762 (2018).

7. D. Gomez et al., Interleukin-1β has atheroprotective effects in advanced atherosclerotic lesions of mice. Nature Medicine 24, 1418–1429 (2018).

8. G. J. Martínez et al., Colchicine Acutely Suppresses Local Cardiac Production of Inflammatory Cytokines in Patients With an Acute Coronary Syndrome. J Am Heart Assoc 4, e002128 (2015).

9. S. Robertson et al., Colchicine therapy in acute coronary syndrome patients acts on caspase-1 to suppress NLRP3 inflammasome monocyte activation. Clinical Science 130, 1237–1246 (2016).

10. N. Schwarz et al., Colchicine exerts anti-atherosclerotic and -plaque-stabilizing effects targeting foam cell formation. The FASEB Journal 37, e22846 (2023).

11. K. Vaidya et al., Colchicine Inhibits Neutrophil Extracellular Trap Formation in Patients With Acute Coronary Syndrome After Percutaneous Coronary Intervention. Journal of the American Heart Association 10, e018993 (2021).

12. K. Vaidya et al., Colchicine Therapy and Plaque Stabilization in Patients With Acute Coronary Syndrome: A CT Coronary Angiography Study. JACC Cardiovasc Imaging 11, 305–316 (2018).

13. J. Berntsson et al., Orosomucoid, Carotid Plaque, and Incidence of Stroke. Stroke 47, 1858–1863 (2016).

14. N. S. Jenny, A. M. Arnold, L. H. Kuller, R. P. Tracy, B. M. Psaty, Serum Amyloid P and Cardiovascular Disease in Older Men and Women. Arteriosclerosis, Thrombosis, and Vascular Biology 27, 352–358 (2007).

15. T. Steiner et al., Protective role for properdin in progression of experimental murine atherosclerosis. PLoS One 9, e92404 (2014).

16. X. Wang et al., Anti-β2GPI antibodies enhance atherosclerosis in ApoE-deficient mice. Biochemical and Biophysical Research Communications 512, 72–78 (2019).

17. S. Beddhu et al., Association of serum albumin and atherosclerosis in chronic hemodialysis patients. American Journal of Kidney Diseases 40, 721–727 (2002).

18. A. Ronit et al., Plasma Albumin and Incident Cardiovascular Disease. Arteriosclerosis, Thrombosis, and Vascular Biology 40, 473–482 (2020).

19. A. A. C. Newman et al., Multiple cell types contribute to the atherosclerotic lesion fibrous cap by PDGFRβ and bioenergetic mechanisms. Nature Metabolism 3, 166–181 (2021).

20. C. G. Caro, J. M. Fitz-Gerald, R. C. Schroter, Arterial Wall Shear and Distribution of Early Atheroma in Man. Nature 223, 1159–1161 (1969).

21. M. W. Majesky, Developmental Basis of Vascular Smooth Muscle Diversity. Arteriosclerosis, Thrombosis, and Vascular Biology 27, 1248–1258 (2007).

22. L. S. Shankman et al., KLF4-dependent phenotypic modulation of smooth muscle cells has a key role in atherosclerotic plaque pathogenesis. Nature Medicine 21, 628–637 (2015).

23. S. Allahverdian, A. C. Chehroudi, B. M. McManus, T. Abraham, G. A. Francis, Contribution of Intimal Smooth Muscle Cells to Cholesterol Accumulation and Macrophage-Like Cells in Human Atherosclerosis. Circulation 129, 1551–1559 (2014).

24. Y. Wang et al., Smooth Muscle Cells Contribute the Majority of Foam Cells in ApoE (Apolipoprotein E)-Deficient Mouse Atherosclerosis. Arteriosclerosis, Thrombosis, and Vascular Biology 39, 876–887 (2019).

25. L. Dobnikar et al., Disease-relevant transcriptional signatures identified in individual smooth muscle cells from healthy mouse vessels. Nature Communications 9, 4567 (2018).

26. M. E. Lin et al., Runx2 deletion in smooth muscle cells inhibits vascular osteochondrogenesis and calcification but not atherosclerotic lesion formation. Cardiovasc Res 112, 606–616 (2016).

27. M. Y. Speer et al., Smooth Muscle Cells Give Rise to Osteochondrogenic Precursors and Chondrocytes in Calcifying Arteries. Circulation Research 104, 733–741 (2009).

28. J. X. Rong, M. Shapiro, E. Trogan, E. A. Fisher, Transdifferentiation of mouse aortic smooth muscle cells to a macrophage-like state after cholesterol loading. Proc Natl Acad Sci U S A 100, 13531–13536 (2003).

29. A. D. Sniderman et al., Apolipoprotein B Particles and Cardiovascular Disease: A Narrative Review. JAMA Cardiol 4, 1287–1295 (2019).

30. N. A. Marston et al., Association of Apolipoprotein B–Containing Lipoproteins and Risk of Myocardial Infarction in Individuals With and Without Atherosclerosis: Distinguishing Between Particle Concentration, Type, and Content. JAMA Cardiology 7, 250–256 (2022).

31. G. F. Alencar et al., Stem Cell Pluripotency Genes Klf4 and Oct4 Regulate Complex SMC Phenotypic Changes Critical in Late-Stage Atherosclerotic Lesion Pathogenesis. Circulation 142, 2045–2059 (2020).

32. T. Grieskamp, C. Rudat, T. H. W. Lüdtke, J. Norden, A. Kispert, Notch Signaling Regulates Smooth Muscle Differentiation of Epicardium-Derived Cells. Circulation Research 108, 813–823 (2011).

33. V. Domenga et al., Notch3 is required for arterial identity and maturation of vascular smooth muscle cells. Genes & Development 18, 2730–2735 (2004).

34. M. B. Hautmann, C. S. Madsen, G. K. Owens, A transforming growth factor beta (TGFbeta) control element drives TGFbeta-induced stimulation of smooth muscle alpha-actin gene expression in concert with two CArG elements. J Biol Chem 272, 10948–10956 (1997).

35. Y. Liu, S. Sinha, G. Owens, A transforming growth factor-beta control element required for SM alpha-actin expression in vivo also partially mediates GKLF-dependent transcriptional repression. J Biol Chem 278, 48004–48011 (2003).

36. G. J. Martínez, D. S. Celermajer, S. Patel, The NLRP3 inflammasome and the emerging role of colchicine to inhibit atherosclerosis-associated inflammation. Atherosclerosis 269, 262–271 (2018).

37. P. Y. Chen et al., Smooth Muscle Cell Reprogramming in Aortic Aneurysms. Cell Stem Cell 26, 542–557.e511 (2020).

38. P. Cheng et al., Smad3 regulates smooth muscle cell fate and mediates adverse remodeling and calcification of the atherosclerotic plaque. Nature Cardiovascular Research 1, 322–333 (2022).

39. C. J. Martos-Rodríguez et al., Fibrous Caps in Atherosclerosis Form by Notch-Dependent Mechanisms Common to Arterial Media Development. Arteriosclerosis, Thrombosis, and Vascular Biology 41, e427–e439 (2021).

40. A. Wirth et al., G12-G13–LARG–mediated signaling in vascular smooth muscle is required for salt-induced hypertension. Nature Medicine 14, 64–68 (2008).

41. M. D. Muzumdar, B. Tasic, K. Miyamichi, L. Li, L. Luo, A global double-fluorescent Cre reporter mouse. genesis 45, 593–605 (2007).

42. L. Madisen et al., A robust and high-throughput Cre reporting and characterization system for the whole mouse brain. Nature Neuroscience 13, 133–140 (2010).

43. L. T. Krebs et al., Characterization of Notch3-deficient mice: Normal embryonic development and absence of genetic interactions with a Notch1 mutation. genesis 37, 139–143 (2003).

44. A. Misra et al., Integrin beta3 regulates clonality and fate of smooth muscle-derived atherosclerotic plaque cells. Nature Communications 9, 2073 (2018).

45. A. B. Nair, S. Jacob, A simple practice guide for dose conversion between animals and human. J Basic Clin Pharm 7, 27–31 (2016).

46. A. Al Shoyaib, S. R. Archie, V. T. Karamyan, Intraperitoneal Route of Drug Administration: Should it Be Used in Experimental Animal Studies? Pharm Res 37, 12 (2019).

47. J. Nadel et al., Intraplaque Myeloperoxidase Activity as Biomarker of Unstable Atheroma and Adverse Clinical Outcomes in Human Atherosclerosis. JACC: Advances 0.

48. H. C. Stary et al., A Definition of Advanced Types of Atherosclerotic Lesions and a Histological Classification of Atherosclerosis. Circulation 92, 1355–1374 (1995).

49. H. F. Dovey et al., Functional gamma-secretase inhibitors reduce beta-amyloid peptide levels in brain. J Neurochem 76, 173–181 (2001).

50. E. T. Grygielko et al., Inhibition of gene markers of fibrosis with a novel inhibitor of transforming growth factor-beta type I receptor kinase in puromycin-induced nephritis. J Pharmacol Exp Ther 313, 943–951 (2005).

51. R. C. Coll et al., A small-molecule inhibitor of the NLRP3 inflammasome for the treatment of inflammatory diseases. Nature Medicine 21, 248–255 (2015).

