## Supplementary Information for "Colchicine promotes atherosclerotic plaque stability independently of inflammation"

**Content:**

Materials and Methods
Figs. S1 to S15
Tables S1 to S4

### Materials and Methods

Animals

All experimental protocols involving mice were approved by the Sydney Local Health District. All atherosclerotic animals used were on an *Apoe*$^{-/-}$ background for plaque development. SMC lineage tracing was performed using the *Myh11*-CreER$^{T2}$ transgene and the *ROSA26*$^{mTmG/+}$ or *ROSA26*$^{tdTomato/+}$ reporter transgenes (*40-42*). Notch3 studies were performed using the *Notch3*$^{-/-}$ mutation (*43*), with and without lineage tracing transgenes. *Apoe*$^{-/-}$,*Notch*$^{+/-}$ were bred with each other to generate littermate controls with the genotype *Apoe*$^{-/-}$,*Notch*$^{+/+}$ for *Apoe*$^{-/-}$,*Notch*$^{-/-}$. Experiments on SMC lineage tracing mice were restricted to males due to limitations with the *Myh11*-CreER$^{T2}$ transgene location on the Y chromosome. All other experiments were performed with both sexes. Cre-*lox* recombination was induced in 6-week-old mice via intraperitoneal injections of tamoxifen (1mg/day; Sigma-Aldrich, T5648) for 5 days (*44*). At 8 weeks old, mice were fed a Western diet containing 22% fat and 0.15% cholesterol (SF00-219, Specialty Feeds) for up to 24 weeks. Colchicine (30µg/kg/day; Sigma-Aldrich, C9754) or vehicle was administered intraperitoneally after 16 weeks WD for the remaining 8 weeks. For healthy non-atherosclerotic experiments, 7-11 week old C57BL/6 (*Apoe*$^{+/+}$) animals were administered intraperitoneally with colchicine (30µg/kg/day), or vehicle for 3 weeks. The dose of colchicine used approximates the FDA approved 0.5 mg/day human dose, based on species difference in body surface area (*45*) and differences in drug bioavailability between oral and intraperitoneal route of administration (*46*).

Human tissue

The Human Research Ethics Committee of St Vincent's Hospital, Darlinghurst, Australia approved this study that conformed to the ethical guidelines of the 1975 Declaration of Helsinki. Anonymized datasets, analyzing methods and study materials can be made available to other researchers for the purpose of reproducing the results or replicating procedures by contacting the corresponding author.

Colchicine extraction from murine tissues

20-130 mg of tissue samples were weighed and added to a lysing matrix tube containing 300µL ice-cold PBS and HALT™ Protease Inhibitor Cocktail. Cell lysis was performed using a Precellys homogenizer with the following settings: 6,000 rpm, 3 x 30s cycles, 10s rest. The resulting homogenate was transferred to fresh Eppendorf tubes then centrifuged to pellet cell debris. Protein concentration was measured via NanoDrop. 20µL aliquots of homogenate were mixed with 180µL of ice-cold methanol and the internal standard D6-colchicine, vortexed, and centrifuged at 15,000 g for 10 min at 4°C. Extracts were then transferred to fresh Eppendorf tubes and evaporated in a vacuum centrifuge. Dried extracts were resuspended in 50µL of methanol and transferred to LC/MS vials for LC/MS analysis.

LC-MS/MS quantification of colchicine

Colchicine concentration was determined by LC-MS/MS using a Shimadzu triple quadrupole LCMS-8050. 10µL of each sample was injected to a Hypersil™ 1.9 µm Gold VANQUISH 100 x 2.1mm column held at 40°C and eluted with a gradient of mobile phase A (water with 0.1% formic acid) and mobile phase B (acetonitrile with 0.1% formic acid). The LC gradient was as follows: 25% mobile phase B at 0.4 mL/min for 0-0.5 min, mobile phase B was increased to 100% from 0.5-5 min then held at 100% from 5-7 min, mobile phase B was decreased to 25% from 7-7.5 min. Colchicine and D6-colchicine were analyzed in positive ion mode with electrospray ionization.

The ESI settings were as follows: interface temperature 300°C; nebulizing gas flow 3 L/min; heating gas flow 6 L/min, DL temperature 210°C, heat block temperature 400°C, drying gas flow 6 L/min. Quantification of colchicine and D6-colchicine was achieved by multiple reaction monitoring with the following energies, respectively: m/z 399.9→310 with collision energy of -27.8 eV and m/z 405.9→313.1 with collision energy of 29 eV. Analysis of LC-MS data was performed using Labsolutions software (Shimadzu).

Immunohistochemistry

*Murine samples*

After anesthesia, blood was drawn, before perfusion through the left ventricle with 20ml of PBS. Mouse organs, aortic root and BCA were collected and fixed with 4% paraformaldehyde (PFA) for 2 hours on ice and then incubated in 30% sucrose at 4°C. Tissue samples were embedded in Optimal Cutting Temperature compound (Tissue Tek, IA018) and cryosectioned at 10μm per section.

For immunofluorescent staining, slides were incubated for an hour in blocking buffer (5% bovine serum albumin (BSA), 2% Tween-20 in PBS), and then incubated in primary antibody solution (in 5% BSA, 1% Tween-20 in PBS) overnight at 4°C. Slides were then incubated in secondary antibody solution for 1 hour. The primary antibodies were; FITC conjugated anti-SMA (1:500, Sigma-Aldrich, F3777), Cy3 conjugated anti-SMA (1:500, Sigma-Aldrich, C6198), eFluor™ 660 conjugated anti-SMA (1:200, Invitrogen, 50-9760-82), anti-SMMHC (1:100, Abcam, ab53219), anti-IL-6 (1:100, Abcam, ab208113), anti-CD68 (1:100, BIO-RAD, MCA1957), anti-Mac2 (1:100, Cedarlane, CL8942AP), anti-GFP (1:100, Abcam, ab13970), anti-RUNX2 (1:100, Abcam, ab192256), anti-NOTCH3 (1:100, Abcam, ab23426), anti-TGFβR1 (1:100, Abcam, ab31013), anti-pSMAD2 (1:100, Invitrogen, 44-244G), anti-HES1 (1:100, Cell Signaling Technology, 11988S). The secondary antibodies were; Alexa Fluor-488, -594, -647 (1:400, Invitrogen). DAPI (1:250, Sigma-Aldrich, D9542) was used to stain the nucleus. Slides were washed with PBST (0.1% Tween-20 in PBS) between antibodies, and mounted using Dako mounting medium (Agilent, S302380-2). It should be noted that the tdTomato fluorescence in *ROSA26*$^{mTmG/+}$ mice is virtually undetected without amplification and thus the red channel was used for other antibody staining.

For immunohistochemistry staining, slides were fixed for an additional 10 min in formalin. Hematoxylin and eosin staining was used for overall vessel morphology and necrotic core identification – slides were incubated with Mayer’s hematoxylin (Sigma, MHS80) for 10 min, Scott’s bluing agent (Leica, 3802901E) for 1 min, and 0.2% eosin in 95% ethanol for 1 min. Oil Red O staining was used for neutral lipids – slides were incubated with 0.18%(w/v) Oil Red O (Sigma, O0625) in 60% isopropanol for 20 min before counterstaining with Mayer’s hematoxylin for 10 min and Scotch bluing buffer for 1 min. Picrosirius red staining was used for collagen content identification – slides were incubated with Picrosirius red (0.3% (w/v) direct red 80 in picric acid solution) for 1 hour. Alizarin red staining was used to identify calcium deposition – slides were stained with 2% Alizarin Red S (pH 4.1-4.3, Sigma-Aldrich, A5533) for 5 min. Slides were mounted with DPX (Sigma, O6522). Hematoxylin and eosin, oil red o, and alizarin red staining was visualized using a ZEISS Axio Scan Z1 Slide Scanner. Picrosirius red staining was imaged using a ZEISS Axio Imager Z2 light microscope under polarized light.

All staining was performed in two locations in the aortic root approximately 200µm apart, and 3 locations in the BCA approximately 300µm apart (**Fig. S2A**). The sample size is indicated in all graphs by individual dots.

*Human samples*

Fixed human coronary artery samples were embedded in paraffin and sectioned at 5µm per section. Sections were deparaffinized with xylene and rehydrated with ethanol. Samples were classified as previously described (*47*). Briefly, plaques were categorized based on AHA histological classification (*48*). Stable or unstable lesions from plaques AHA grade IV or higher were used. Unstable lesions were classified as AHA grade VI, or if ≥1 of the following criteria were met: fibrous cap < 65um, lipid core >25% of section area, >100 inflammatory cells or >14 within the fibrous cap, previous plaque rupture, intraplaque hemorrhage or neovascularization. Thick fibrous cap regions from stable plaques or thin fibrous cap regions from unstable plaques were used for analysis.

For immunohistochemistry staining, antigen retrieval was performed using pepsin (0.5% in 5mM HCl, pH 2) for 10 min at 37°C and 10 min at room temperature. Slides were blocked for endogenous peroxidases using 0.3% hydrogen peroxide in methanol for 20 min, before incubation for 30 min in blocking buffer (5% BSA, 0.1% Triton-X 100 in TBS). Primary antibody incubation was performed overnight at 4°C. Slides were then washed, and incubated at room temperature in secondary antibody solution for 1 hour. Washing was performed between blocking and antibody steps using TBST (0.1% Triton-X 100 in TBS). Chromogen detection was performed using 3,3'-diaminobenzidine (Agilent, K346811-2) for 10 min. Slides were stained for hematoxylin and dehydrated before mounting with DPX. The primary antibodies were anti-pSMAD2 (1:100, Invitrogen, 44-244G), anti-SMA (1:100, Sigma, A2547). The secondary antibodies were; anti-mouse (1:250, Agilent, P044701-2), anti-rabbit (1:250, Agilent, P044801-2).

For immunofluorescent staining, slides were rehydrated with TNT buffer (0.1M Tris-HCl, 0.2% Tween-20 in saline, pH 7.5) for 10 min before antigen retrieval with sodium citrate buffer (10mM, pH 6) for 20 min at 90°C or pepsin (0.5% in 5mM HCl, pH 2) for 10 min at 37°C and 10 min at room temperature. Slides were washed again with TNT buffer for 10 min before performing staining as per murine samples (described above). The primary antibodies were; FITC conjugated anti-SMA (1:500, Sigma-Aldrich, F3777), Cy3 conjugated anti-SMA (1:500, Sigma-Aldrich, C6198), anti-TGFβR1 (1:100, Abcam, ab31013), anti-NOTCH3 (1:100, Abcam, ab23426). The secondary antibodies were; Alexa Fluor-488, -594 (1:400, Invitrogen).

<u>Imaging</u>

Immunofluorescent images were obtained using the Zeiss LSM800 confocal microscope. Image counting and processing was performed using Zen Blue/Black, Adobe Photoshop, and Image J.

<u>Defining the fibrous cap</u>

The fibrous cap thickness was determined by the average depth of continuous $ACTA2^+$ cells from the lumen, in a similar manner described by Newman *et al.*(*19*). In brief, evenly spaced lines were drawn perpendicular to the endothelium (**Fig. S2B**). The distance of uninterrupted $ACTA2^+$ cells (not separated by $ACTA2^-$ cells) along these perpendicular lines was used. Averages of $ACTA2^+$ depths along 30 perpendicular lines were used to determine the fibrous cap in the aortic root, or along 15 perpendicular lines in the BCA per animal. Unlike Newman *et al.*(*19*), the fibrous cap thickness was defined by averaging the mean thickness per animal, rather than taking the maximum thickness, as colchicine treatment led to some animals with disproportionately large fibrous cap to core ratios, which would have hindered analysis of the lesion core. The average $ACTA2^{+}$ cell depth (*Apoe*$^{-/-}$) after colchicine treatment was 38.0µm in the aortic root and 17.5µm in the BCA. Thus, throughout this study, the region we used for general fibrous cap analysis was 38.0µm in aortic root plaques, and 17.5µm in BCA plaques.

Image quantification and analysis of murine lesions

The lesion and necrotic core area as well as lumen and the internal elastic lamina area were measured using ImageJ. Oil Red O staining, Picrosirius red staining, Alizarin red staining were quantified by measuring positive area within threshold of selective color filtered with color deconvolution using ImageJ.

Cell counting was performed using Zeiss Zen software to quantify cell composition within cap, core, and lesion.

Proteomics

2µL of plasma was diluted in 1% sodium deoxycholate in 100mM HEPES pH 8.5 and protein disulphides reduced and alkylated with 5mM Tris (2-carboxyethyl) phosphine (TCEP; final concentration) and 10mM iodoacetamide (IAA; final concentration). Samples were heated to 95°C for 10 min and left to cool at room temperature for 50 min. 1µg of trypsin (Promega) was added to each sample and incubated at 37°C for 16 hours. Digested samples were diluted with 10 volumes of 90% acetonitrile, 1% trifluoroacetic acid (TFA), centrifuged at 16,500g and peptides captured and cleaned using SDB-RPS-based STAGE tips. After washing with 9% acetonitrile, 0.1% TFA, peptides were eluted in 50µL of 1% ammonia in 80% acetonitrile which was then removed by rotary evaporation and the peptides resuspended in 25µL of 2% acetonitrile, 0.1% TFA for analysis by LC-MS/MS.

Using an Acquity M-class nanoLC system (Waters, USA), 5µL of the sample was loaded at 15µL/min for 3 min onto a nanoEase Symmetry C18 trapping column (180µm x 20mm) before being washed onto a PicoFrit column (75 µmID x 350 mm; New Objective, Woburn, MA) packed with SP-120-1.7-ODS-BIO resin (1.7µm, Osaka Soda Co, Japan) heated to 45°C. Peptides were eluted from the column and into the source of a Q Exactive Plus mass spectrometer (Thermo Scientific) using the following program: 5-30% MS buffer B (98% acetonitrile + 0.2% formic acid) over 90 min, 30-80% MS buffer B over 3 min, 80% MS buffer B for 2 min, 80-5% for 3 min. The eluting peptides were ionized at 2400V. A Data Dependant MS/MS (dd-$MS^2$) experiment was performed, with a survey scan of 350-1500 Da performed at 70,000 resolution for peptides of charge state 2+ or higher with an AGC target of 3e6 and maximum Injection Time of 50ms. The Top 12 peptides were selected and fragmented in the HCD cell using an isolation window of 1.4 m/z, an AGC target of 1e5 and maximum injection time of 100ms. Fragments were scanned in the Orbitrap analyser at 17,500 resolution and the product ion fragment masses measured over a mass range of 120-2000 Da. The mass of the precursor peptide was then excluded for 30s.

The MS/MS data files were searched using Peaks Studio X Pro against the UniProt Mouse reference proteome database and a database of common contaminants with the following parameter settings. Fixed modifications: none. Variable modifications: carbamidomethyl, oxidised methionine, deamidated asparagine. Enzyme: semi-trypsin. Number of allowed missed cleavages: 3. Peptide mass tolerance: 10 ppm. MS/MS mass tolerance: 0.05 Da. The results of the search were then filtered to include peptides with a $-\log_{10}P$ score that was determined by the False Discovery Rate (FDR) of <1%, the score being that where decoy database search matches were <1% of the total matches. Label Free Quantification (LFQ) was performed using the PEAKS Q module.

Cell culture

Human aortic smooth muscle cells (HASMCs; Sigma-Aldrich, 354-05A) were cultured in Smooth Muscle Cell Growth Medium-2 Bullet Kit (Lonza, CC-3182) at 37°C, 5% $CO_2$. At 70% confluency, the cells were serum starved (0.2% BSA in high glucose DMEM) for 24 hours, and the treatments were given as indicated. For macrophage-like regression, cells were loaded with cholesterol-methyl-β-cyclodextrin (10µg/mL, Sigma-Aldrich, C4951) for 24-72 hours (*28*), and then treated with colchicine (50nM, Sigma-Aldrich, C9754) for an additional 24-48 hours. For oxidized LDL uptake experiments, cells were treated with oxidized LDL (10µg/ml, ThermoFisher, L34358) and colchicine (50nM). For Notch inhibition experiments, cells were treated with cholesterol, the γ-secretase inhibitor DAPT (*N*-[(3,5-difluorophenyl)acetyl]-L-alanyl-2-phenyl]glycine-1,1-dimethylethyl ester) (5µM, Sigma-Aldrich, D5942) and/or colchicine (50nM). For TGFβ experiments, cells were treated with cholesterol, TGFβ1 (2.5ng/mL, Sigma-Aldrich, T7039 or BioLegend, 580702) and/or the TGFβ inhibitor SB-525334 (5µg/mL, Sigma-Aldrich) and/or colchicine (50nM). For osteoblast-like regression, the cells were not serum starved, and instead grown in osteogenic medium (ThermoFisher, A1007201) for 16 days, with colchicine intervention (25nM) at day 7 (media changed every 3 days). For NLRP3 inflammasome experiments, cells were treated with MCC950 (10µM, Sigma-Aldrich, PZ0280) or colchicine (50nM). Inhibitor concentrations used were well above IC50 to ensure pathway inhibition and account for differences in cell type with published values (*49-51*).

Quantitative real-time PCR

At the appropriate endpoints, RNA was extracted according to QIAGEN guidelines using the QIAzol lysis reagent (QIAGEN, 79306). Complementary DNA was synthesized using the SensiFAST cDNA synthesis kit (Bioline, BIO-65054) in a Bio-Rad Thermal Cycler. SensiFAST SYBR No-ROX (Bioline, CSA-01133) and appropriate primers (see below) were used to amplify cDNA of interest in a Bio-Rad Real-Time PCR Detection System. Analysis was performed using the $2^{-\Delta\Delta ct}$ method, using glyceraldehyde 3-phosphate dehydrogenase (GAPDH) as the housekeeping gene for normalization.

qPCR Primers: *Homo Sapiens*

| Gene Name | Forward Primer | Reverse Primer |
|---|---|---|
| ***ABCA1*** | CAGGCTACTACCTGACCTTGGT | CTGCTCTGAGAAACACTGTCCTC |
| ***ACTA2*** | AGCCAAGCACTGTCAGGAATC | GAGCCCAGAGCCATTGTCAC |
| ***CD68*** | GCTACATGGCGGTGGAGTACAA | ATGATGAGAGGCAGCAAGATGG |
| ***CNN1*** | ATGTCCTCTGCTCACTTCAAC | GCTGGTGGTCATACTTCTGG |

| | | |
|---|---|---|
| ***COL1A1*** | GGACACAGAGGTTTCAGTGGT | CACCATCATTTCCACGAGCA |
| ***GAPDH*** | GAAGGCTGGGGCTCATTT | CAGGAGGCATTGCTGATGAT |
| ***HES1*** | AAGAAAGATAGCTCGCGGCA | CGGAGGTGCTTCACTGTCAT |
| ***NOTCH3*** | CGTGGCTACACTGGACCTC | AGATACAGGTGAACTGGCCTAT |
| ***TGFBR1*** | GACAACGTCAGGTTCTGGCTCA | CCGCCACTTTCCTCTCCAAACT |
| ***TGLN*** | AGTCTTCACTCCTTCCTG | CTCGTCATACTTCTTCTCG |

Bulk RNA-seq and analysis

RNA was extracted using the PureLink™ RNA Mini kit (Invitrogen, 12183018A) according to the manufacturer's instructions. Quality control and RNA library preparation was performed by the Australian Genome Research Facility, using the Illumina Stranded Total RNA Prep Ligation with Ribo-Zero Plus kit and sequencing using the Illumina NovaSeq platform.

Between 49 and 73 million paired reads (average 62 million) were obtained for the colchicine intervention experiment (**Fig. S7A**). Between 53 and 75 million paired reads (average 72 million) were obtained for the colchicine loading experiment (**Fig. S11A**). Trimming of adapters and low-quality nucleotides was performed using TrimGalore version 0.4.1. Gene quantification (read count and FPKM) was obtained using RSEM version 1.3.0 (mapping using bowtie 2.4.2) (options: -bowtie2 -pairedend), based on GENCODE annotation (Release 38). Mapping rate to the transcriptome ranged from 75 to 79% in the colchicine intervention experiment, and from 78 to 84% in the colchicine loading experiment.

Downstream analysis was performed on R version 4.2.2.

Principal component analysis and differential gene expression was performed using DESeq2 version 1.38.3. For the colchicine loading experiment, differential gene expression analysis was done using a paired analysis as the PCA plot showed a separation based on replicate on the first principal component. To identify relevant differential expressed genes, we used a 1.5 fold change and an adjusted (Benjamini-Hochberg corrected) pvalue below 0.05 thresholds and considered genes with an expression of 1 Fragments Per Kilobase per Million mapped fragments (FPKM) in at last 2 samples.

Heatmap was obtained using package pheatmap version 1.0.12.

Gene ontology analysis was done using TopGo version 2.50.0 using org.Hs.eg.db_3.16.0. Only significant ($p<0.01$) enriched GoTerms with more than 2 genes were considered. The top5 GoTerm (based on p-value) of colchicine regulated genes were extracted after removing redundant terms using Revigo (http://revigo.irb.hr/ with small list output).

Cell staining

Cells were washed with PBS and fixed in 4% PFA for 20 min. Immunofluorescent staining was performed in a similar manner to murine tissue samples. The primary antibodies used were; Cy3 conjugated anti-SMA (1:500, Sigma-Aldrich, C6198), anti-pSMAD2 (1:100, Invitrogen, 44-244G), anti-HES1 (1:100, Cell Signaling Technology, 11988S). The secondary antibodies used were; Alexa Fluor-488 (1:400, Invitrogen). For alizarin red staining, fixed cells were stained for 45 min with 2% Alizarin Red S Solution, before washing with distilled water.

Alizarin Red quantification

Well images were taken using the Hirox HRX-01 microscope. For quantification, cells were shaken for 30 min in 10% acetic acid, heated to 95°C, then centrifuged at 20,000g for 15 min at 4°C before isolation of the supernatant. pH was elevated to 4.1-4.3, and absorbance at 405nm was measured using a CLARIOstar microplate reader.

Statistics

All graphs were created in GraphPad Prism version 9.3, and subsequently analyzed statistically. Error bars represent mean ± SEM. Individual dots represent biologically independent animals or independent experiments. For data comparing multiple locations (i.e. animals examined at 2 locations across aortic root and 3 locations across BCA) or regions within lesions (i.e. animals quantified in cap, core and total lesion) between treatment groups, statistical significance was determined by a two-way ANOVA with multiple comparisons and Sidak correction. Data for individual locations between treatment groups were analyzed by a two-sided, unpaired Mann–Whitney U test. P-values $\leq 0.05$ were considered statistically significant.

Data Access

The RNA-seq datasets are deposited in the Gene Expression Omnibus (GEO) repository and accession numbers will be provided upon manuscript acceptance for publication.

The coverage file (bigwig) and peak file (narrowpeak) for SMAD3 ChIP-Seq and ATAC-Seq from human coronary artery smooth muscle cells was downloaded from GEO under accession numbers GSM3175516 and GSM1876021, respectively.

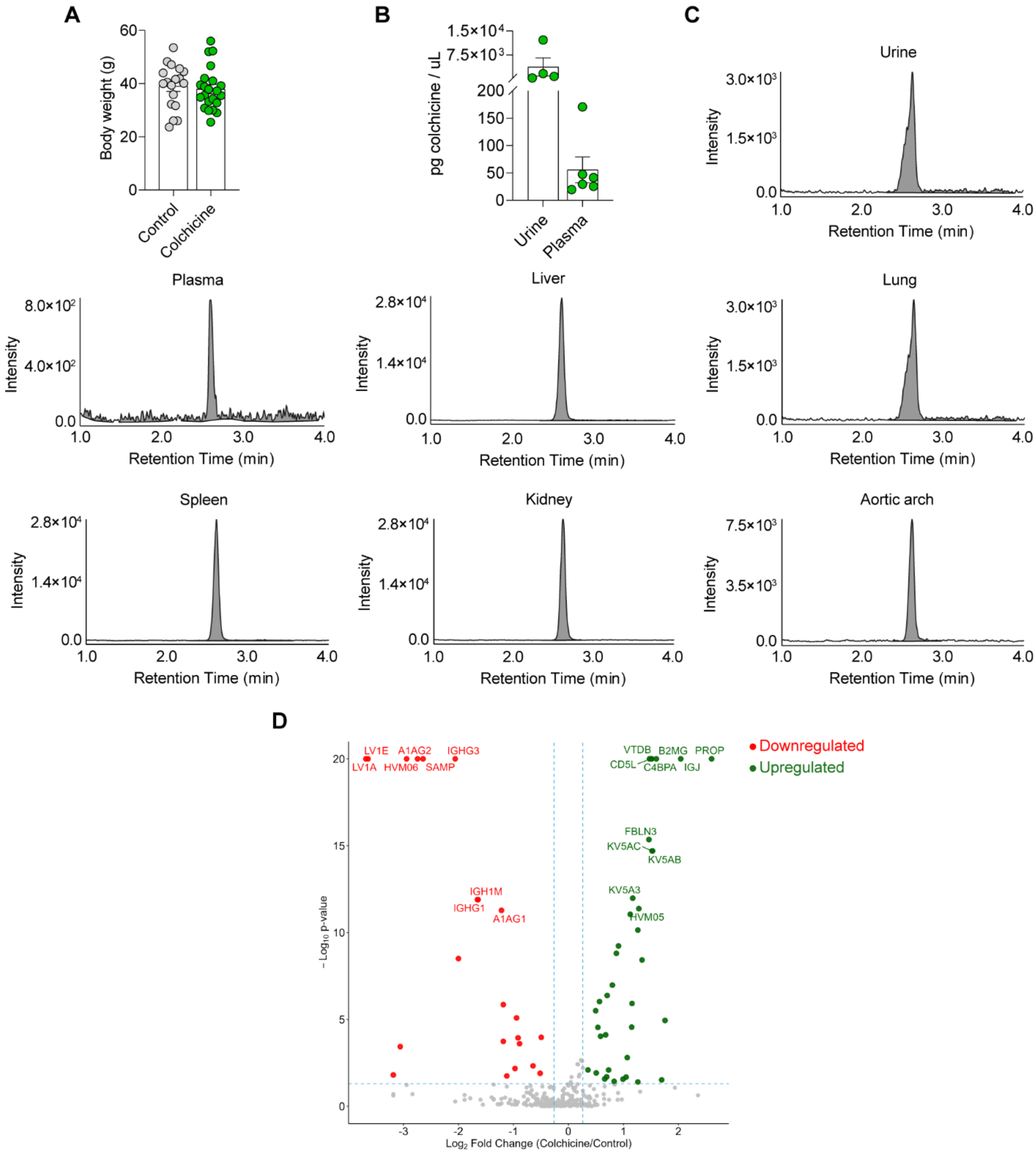

**Fig. S1. The administration of a low dose of colchicine leads to its systemic distribution and induces changes in the plasma proteome. A,** Body weight at harvest remained unchanged after colchicine intervention ($p$ = 0.6777). **B,** Colchicine is detected within the urine and plasma. **C,** Representative chromatograms of colchicine detection throughout the body. **D,** Volcano plot of identified proteins from proteomic analysis of plasma, with top 20 significantly changed proteins labelled. Data was analyzed with a two-tailed unpaired t-test. Individual dots represent biologically independent animals. Graphs show mean ± SEM.

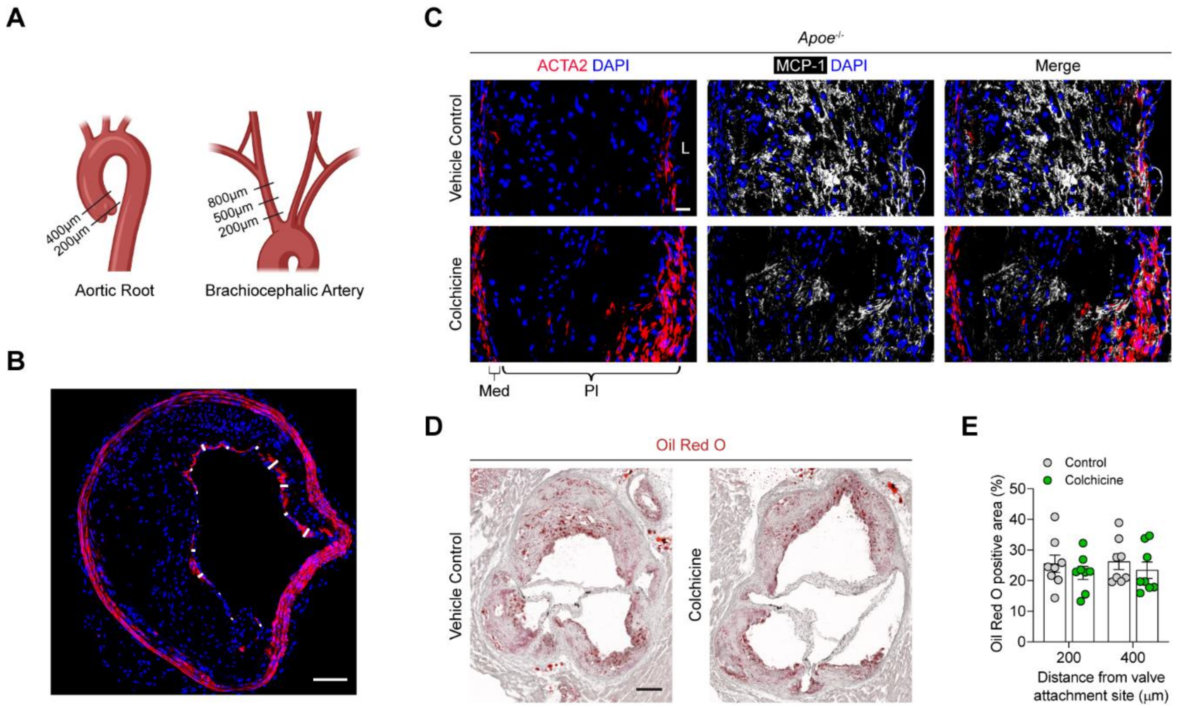

**Fig. S2. Colchicine intervention reduces MCP-1 expression but does not impact lipid uptake in atherosclerotic plaques of the aortic root. A,** Plaques from the aortic root and brachiocephalic artery were analyzed in multiple locations. **B,** Representative image of measurements made to determine continuous depth of ACTA2$^{+}$ cells. **C,** Colchicine reduced the expression of MCP-1. **D, E,** Representative image of Oil Red O staining after colchicine treatment (**D**), with associated quantification (**E**) ($p = 0.2657$). L, lumen; Med, tunica media; Pl, plaque. Scale bar: 20μm (**C**), 100μm (**B**), 200μm (**D**). Results are representative of n = 11 mice for control and n = 11 for colchicine (**C**). Data was analyzed using a two-way ANOVA with Sidak correction and multiple comparisons. Individual dots represent biologically independent animals. Graphs show mean ± SEM. *P* values refer to two-way ANOVA between treatment conditions.

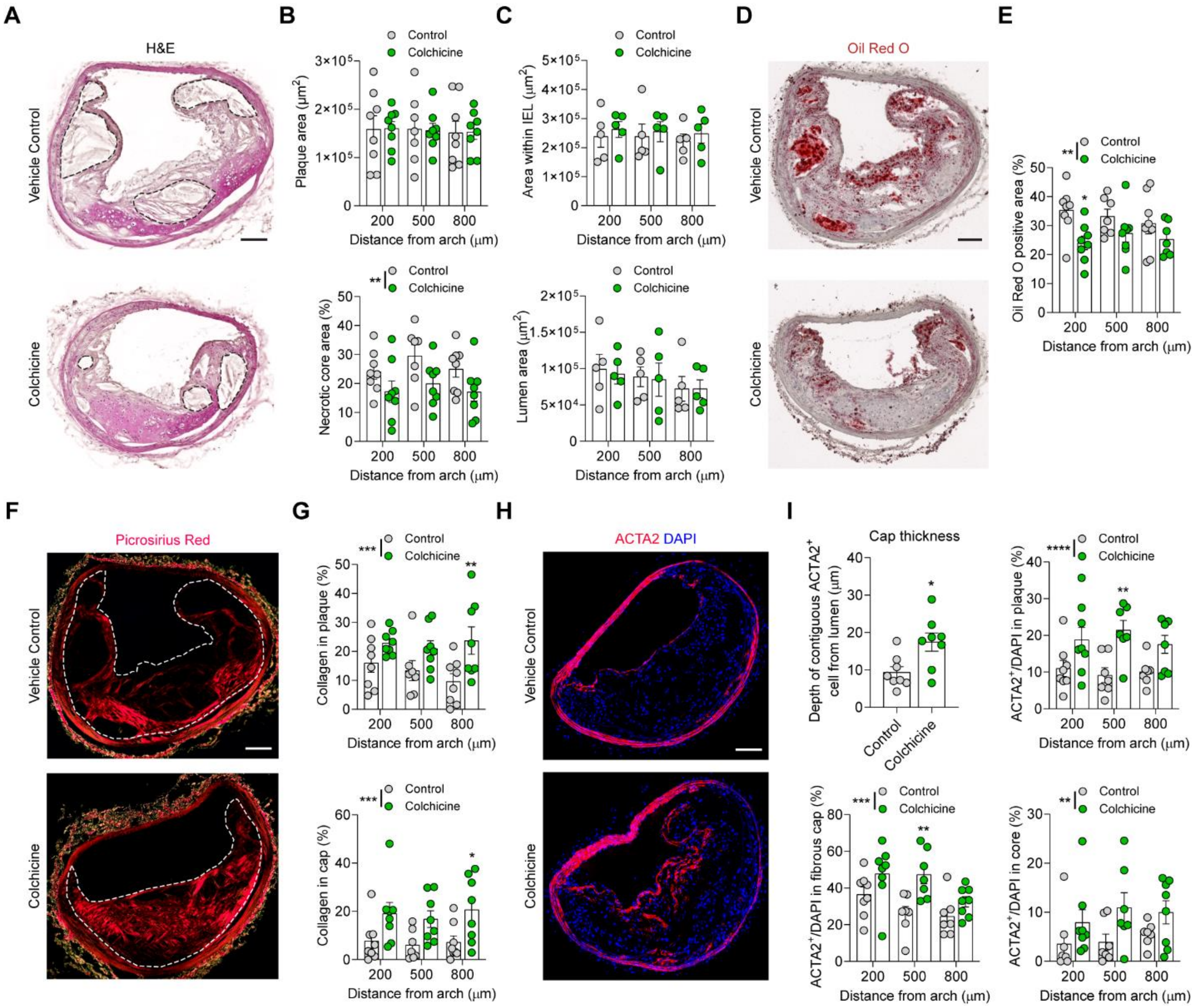

**Fig. S3. Colchicine promotes multiple indices of plaque stability in brachiocephalic lesions.** **A,** Representative images of hematoxylin and eosin staining with necrotic core highlighted. **B,** Quantification of these lesions revealed no change in total lesion area ($p$ = 0.9915), but a reduction in necrotic core area ($p$ = 0.0030) after colchicine intervention. **B, C,** Beneficial outward remodeling was unchanged with colchicine intervention, as assessed by the combination of total lesion area (**B**), area within the internal elastic lamina (**C**) ($p$ = 0.4242), and total lumen area (**C**) ($p$ = 0.8025). **D, E,** Colchicine treatment reduced intraplaque lipid and triglyceride levels ($p$ = 0.0022), assessed by Oil Red O staining (**D**) and associated quantification (**E**). **F, G,** Collagen deposition was increased in the lesion ($p$ = 0.0007) and its fibrous cap ($p$ = 0.0003), indicated by Picrosirius Red staining (**F**) and corresponding quantification (**G**). **H, I,** Representative image of ACTA2 staining (**H**) demonstrating increased fibrous cap thickness ($p$ = 0.0135), as well as increased proportions of $ACTA2^+$ cells in the total lesion ($p < 0.0001$), fibrous cap ($p$ = 0.0005), and plaque core ($p$ = 0.0063) (**I**). Scale bars: 100µm. Data was analyzed using a two-way ANOVA with Sidak correction and multiple comparisons (**B, C, E, G, I**) or a two-tailed unpaired t-test (**I** top left). Individual dots represent biologically independent animals. Graphs show mean ± SEM. $P$ values refer to two-way ANOVA between treatment conditions.

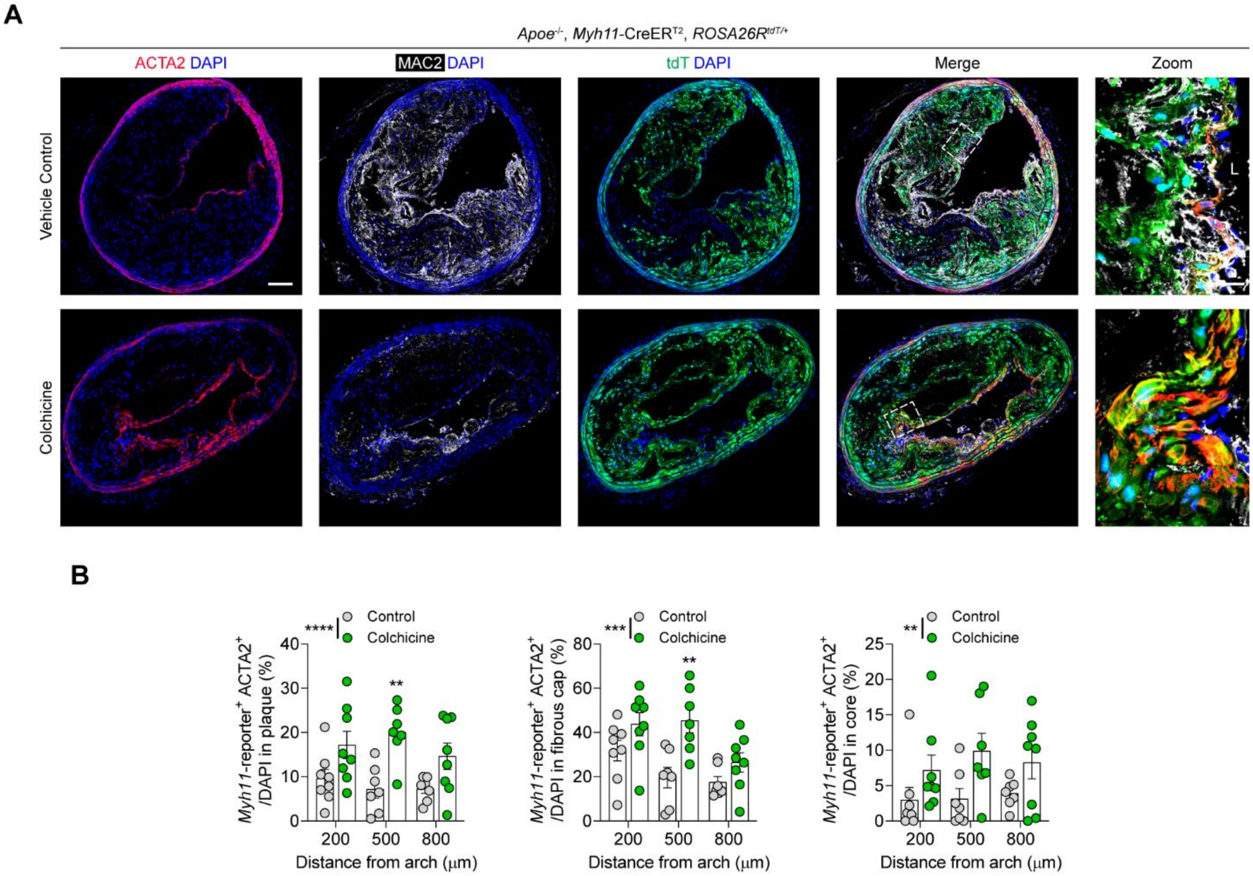

**Fig. S4. Colchicine intervention promotes an ACTA2$^+$ myofibroblast-like phenotype in SMC-derived cells of brachiocephalic lesions. A, B,** Brachiocephalic artery plaques demonstrated increased proportions of SMC-lineage$^+$ ACTA2$^+$ cells of total plaque cells within the total lesion ($p < 0.0001$), fibrous cap ($p = 0.0002$), and core ($p = 0.0027$), as evident by representative ACTA2 staining (**A**) and associated quantification (**B**). L, lumen .Scale bar: 100μm (**A**), 20μm (**A**; zoom). Data was analyzed using a two-way ANOVA with Sidak correction and multiple comparisons. Individual dots represent biologically independent animals. Graphs show mean ± SEM. *P* values refer to two-way ANOVA between treatment conditions.

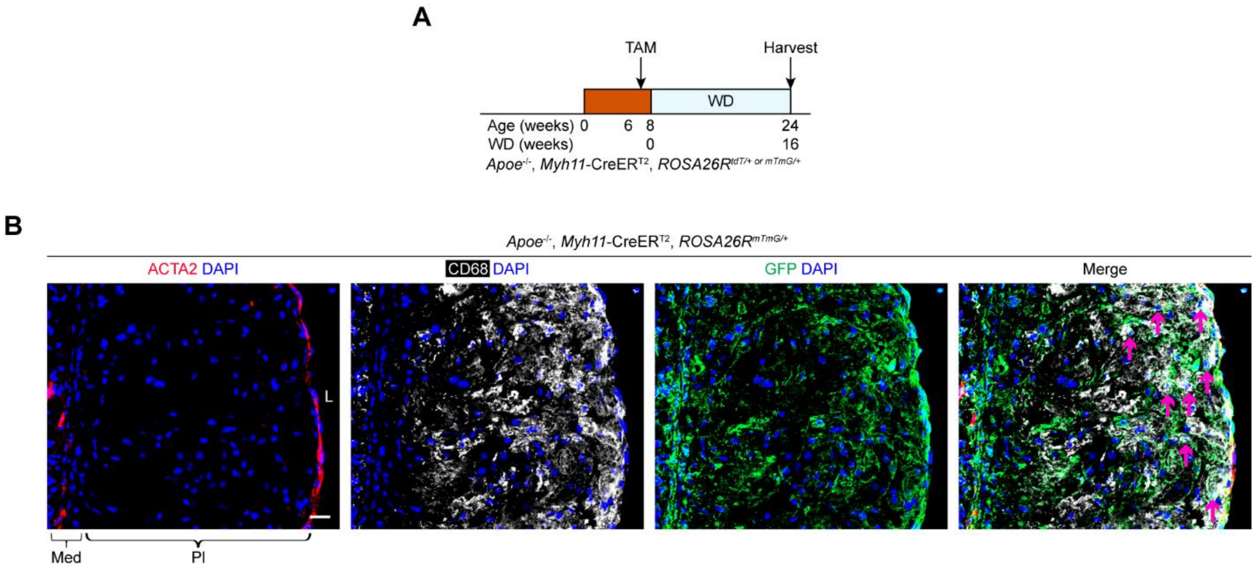

**Fig. S5. SMC-derived cells *trans*-differentiate into CD68$^+$ cells after 16 weeks WD. A,** SMC lineage tracing mice were labelled with tamoxifen and fed a WD for 16 weeks. **B,** SMC-derived CD68$^+$ cells are present in 16-week WD advanced plaques prior to colchicine intervention. Arrows point to SMC-lineage$^+$ CD68$^+$ cells. L, lumen; Med, tunica media; Pl, plaque. Scale bar: 20μm. Results are representative of n = 7 mice.

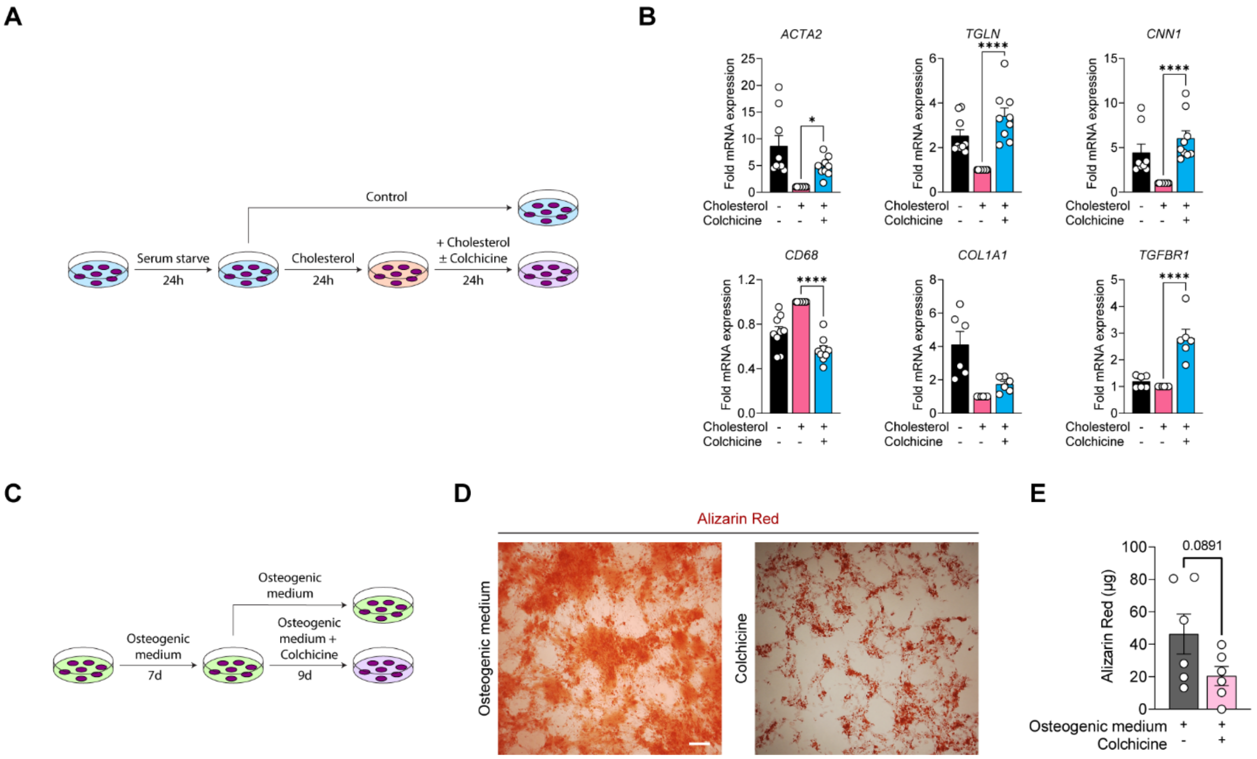

**Fig. S6. Colchicine reverses athero-promoting phenotypes in cultured SMCs. A,** Schematic demonstrating colchicine intervention after short-term macrophage-like *trans*-differentiation. Briefly, HASMCs were cholesterol loaded (10μg/mL) for 24 hours followed by treatment with colchicine (50nM) for 24 hours. **B,** mRNA transcript levels of myofibroblast markers *ACTA2*, *TGLN*, *CNN1*, *COL1A1*, macrophage marker *CD68*, TGFβ signaling mediator *TGFBR1* were analyzed with qPCR. **C,** Schematic demonstrating colchicine intervention during osteoblast-like *trans*-differentiation. Briefly, HASMCs were grown in osteogenic medium for 16 days, with colchicine intervention (25nM) from day 7. **D, E,** Representative Alizarin Red staining (**D**) and quantification (**E**) after colchicine treatment. Scale bars: 500μm. Data was analyzed using a one-way ANOVA with Sidak correction and multiple comparisons (**B**) or a two-tailed unpaired t-test (**E**). Individual dots represent independent experiments. Graphs show mean ± SEM.

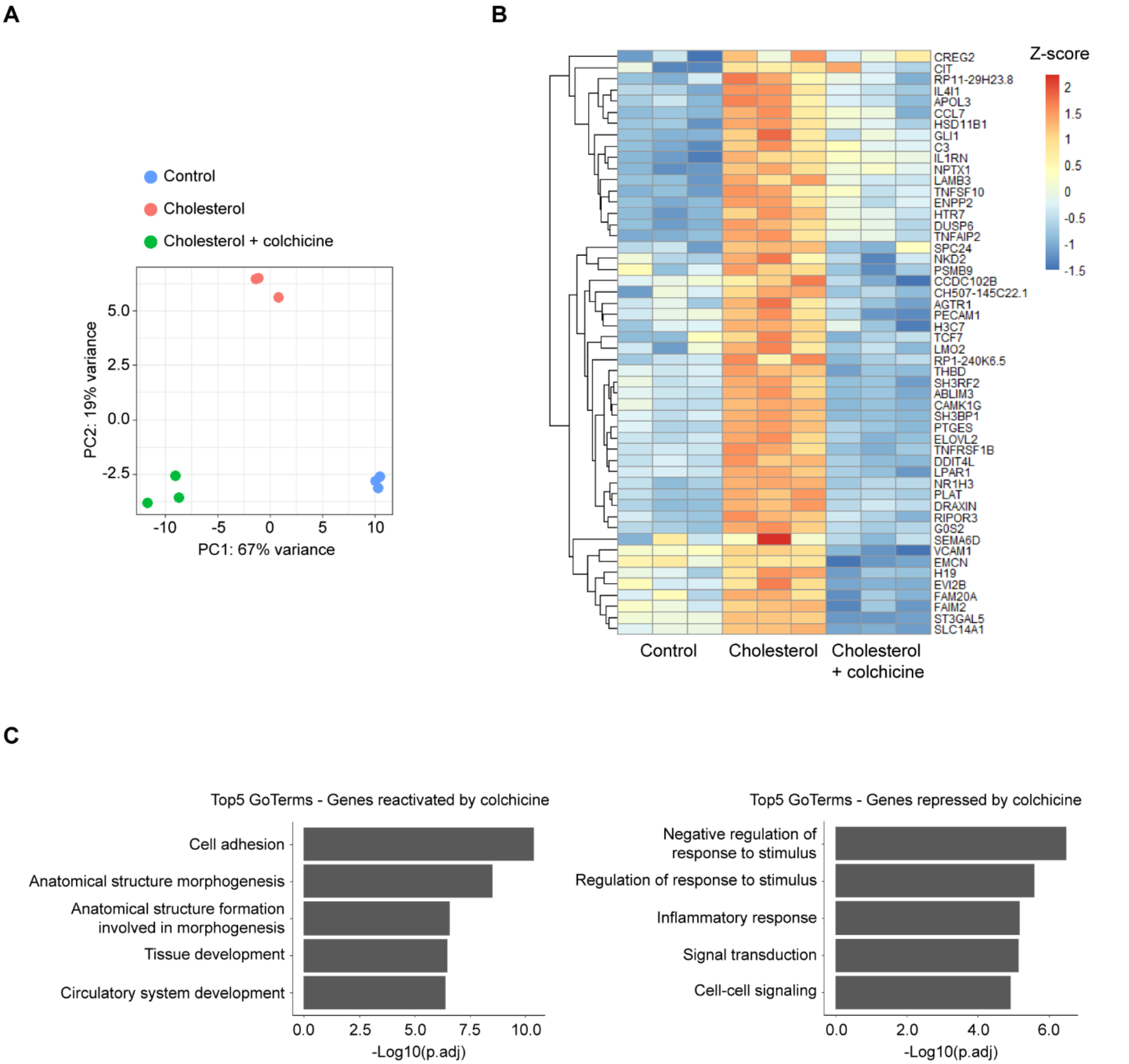

**Fig. S7. Transcriptional changes in SMCs in response to colchicine intervention after cholesterol loading.** **A,** Principal component analysis of the RNA-seq. **B,** Heatmap of genes significantly upregulated by cholesterol and repressed with colchicine intervention. **C,** Top 5 enriched Go-Terms that were significantly downregulated by cholesterol and reactivated by colchicine (left) or upregulated by cholesterol and repressed by colchicine (right).

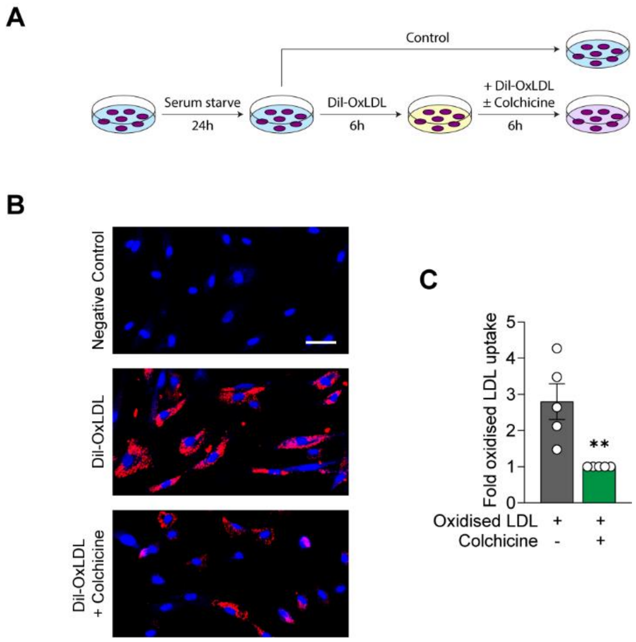

**Fig. S8. Colchicine reduces lipid uptake in cultured human aortic SMCs. A,** Schematic demonstrating colchicine intervention of oxLDL loading. Briefly, HASMCs were loaded with dil-oxLDL (10µg/mL) for 6 hours followed by treatment with colchicine (50nM) for 6 hours. **B, C,** Colchicine intervention reduced oxLDL accumulation within SMCs ($p$ = 0.0065) as assessed by representative image (**B**) and associated quantification (**C**). Scale bars: 50µm. Data was analyzed using a two-tailed unpaired t-test. Individual dots represent independent experiments. The graph shows mean ± SEM.

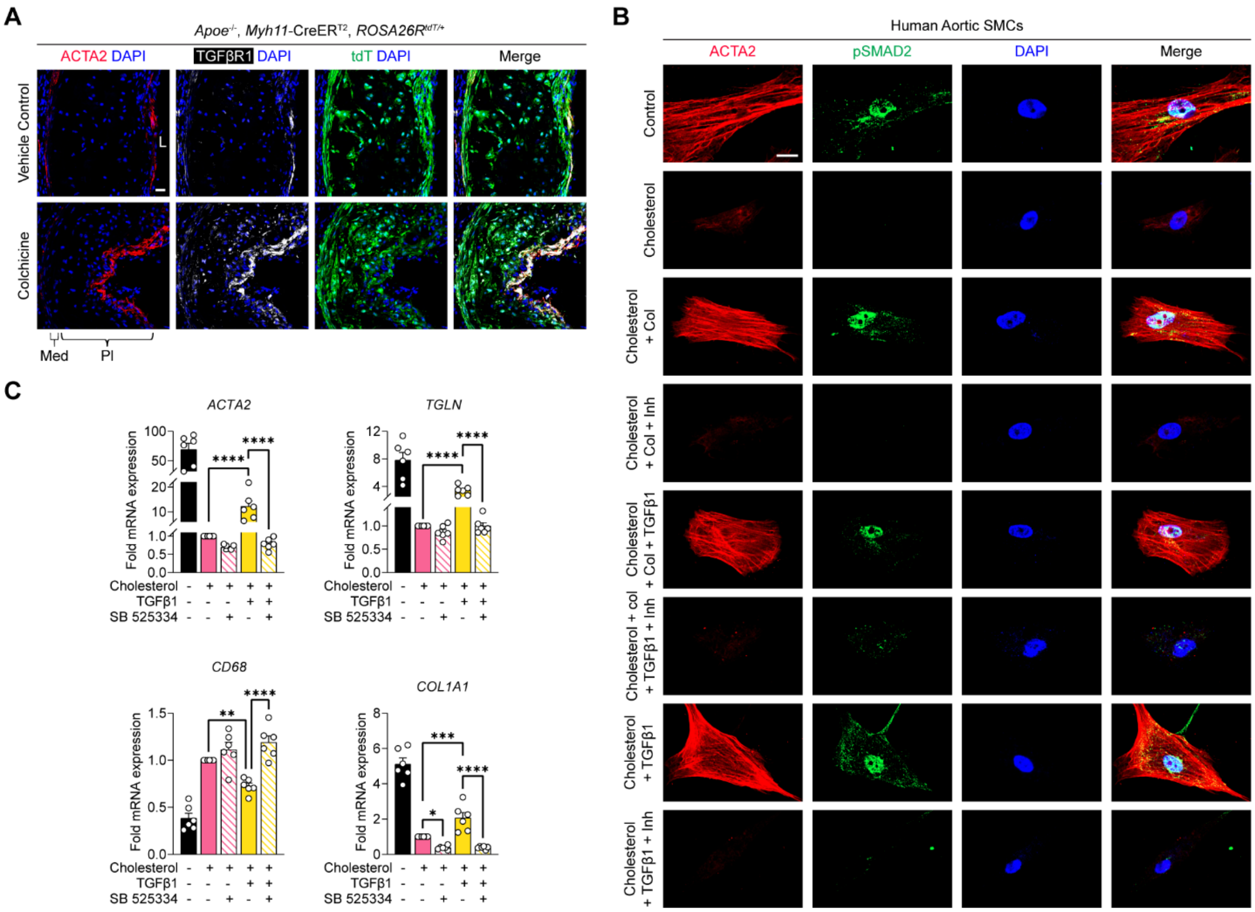

**Fig. S9. Colchicine reverses athero-promoting phenotypes in SMCs by inducing the TGFβ pathway. A,** Representative image demonstrating an increase in TGFβR1 expression after colchicine treatment (n = 9 mice for each group). **B, C,** HASMCs were loaded with cholesterol, and phenotypically rescued with colchicine (50nM) and/or TGFβ1 (2.5ng/mL) in the presence of the TGFβR1 inhibitor SB525334 (5µg/mL). Co-treatment with SB525334 ameliorates the effects of colchicine (**B**), and TGFβ1 (**B, C**). L, lumen; Med, tunica media; Pl, plaque; Col, colchicine; Inh, SB525334. Scale bar: 20µm. Data was analyzed using a one-way ANOVA with Sidak correction and multiple comparisons. Individual dots represent independent experiments. Graphs show mean ± SEM.

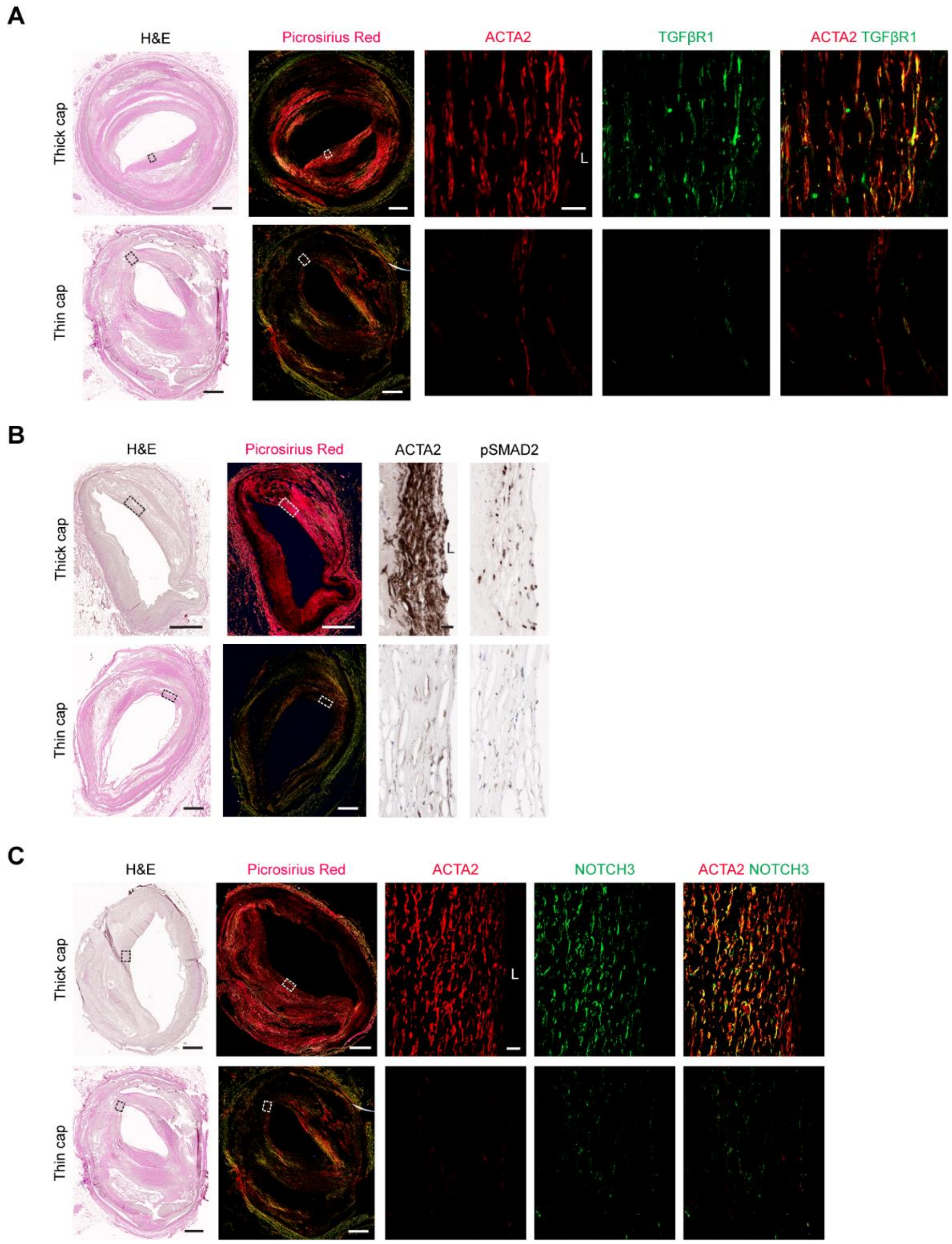

**Fig. S10. Human coronary plaques with thick fibrous caps exhibit increased TGFβ signaling and Notch3 expression. A, B, C,** Human coronary atherosclerotic plaques were classified as stable or unstable based on histological analysis. Thick fibrous cap regions from stable plaques and thin fibrous cap regions from unstable plaques were stained for TGFβ and Notch signaling mediators. TGFβ signaling (**A, B**) and Notch signaling (**C**) were elevated in thick cap stable plaques. Scale bar: 500μm (total lesion) or 20μm (zoom). Results are representative of n = 9 samples for each group.

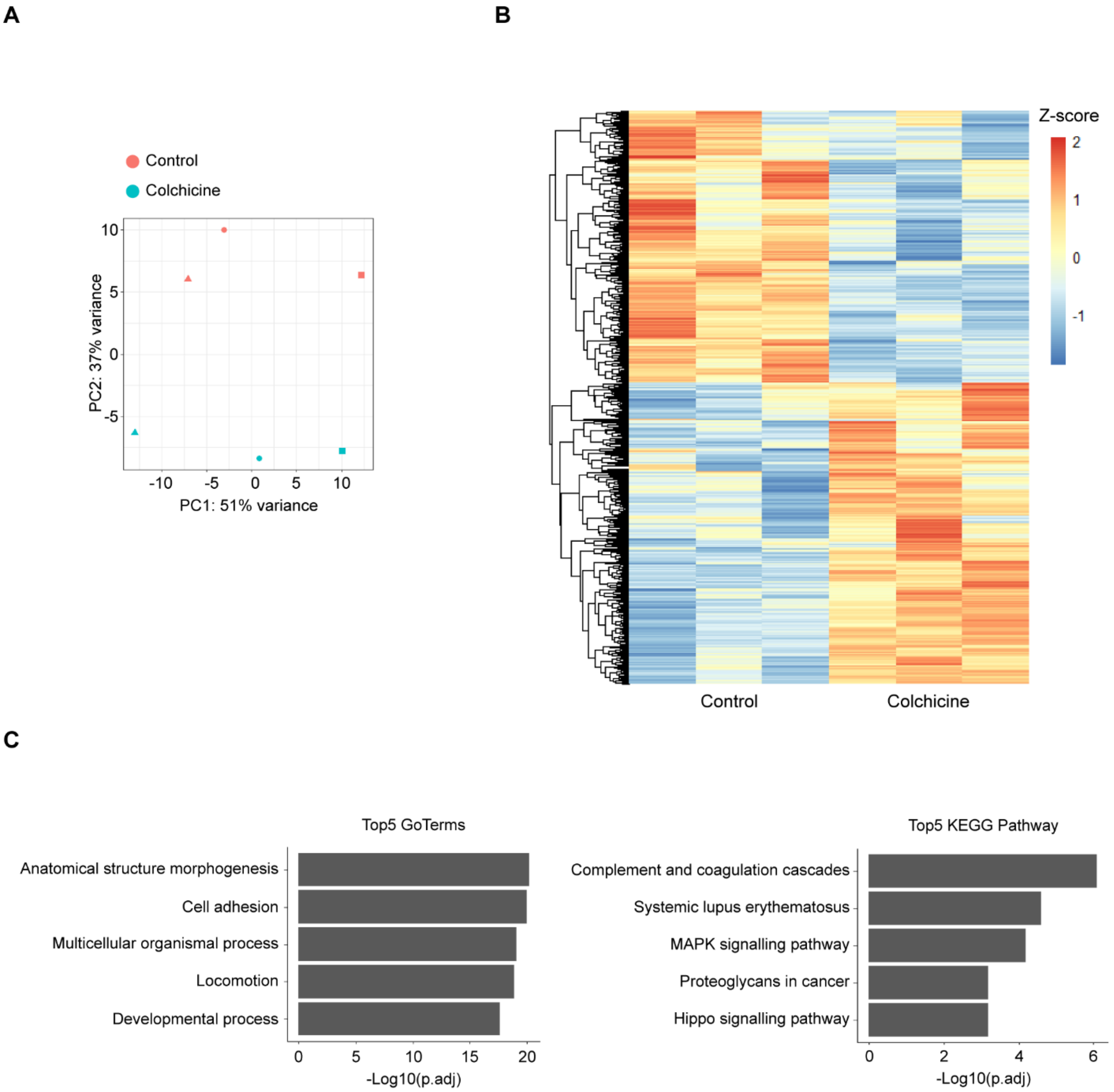

**Fig. S11. Colchicine treatment of human aortic SMCs induces widespread transcriptional changes.** **A,** Principal component analysis of the RNA-seq. **B,** Heatmap of genes differentially regulated by colchicine treatment. **C,** Top 5 enriched Go-Terms (left) and KEGG pathways (right) that were significantly upregulated by colchicine.

A

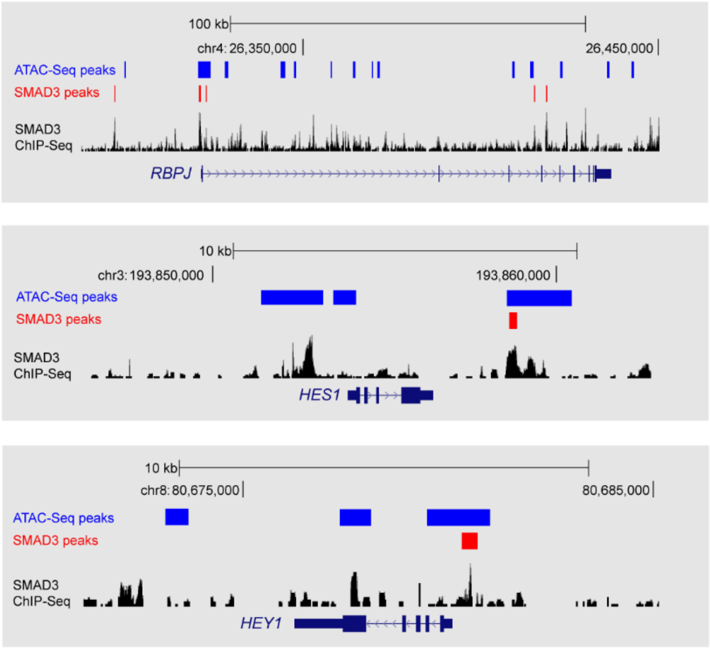

**Fig. S12. Binding of SMAD3 to Notch downstream mediators in human coronary artery SMCs. A,** UCSC browser images of ChIP-Seq data from human coronary artery SMCs, demonstrating the binding of SMAD3 to the promoters of *RBPJ* and *HEY1* and to nearby enhancer element(s) of *HES1*, defined by open chromatin (ATAC-Seq).

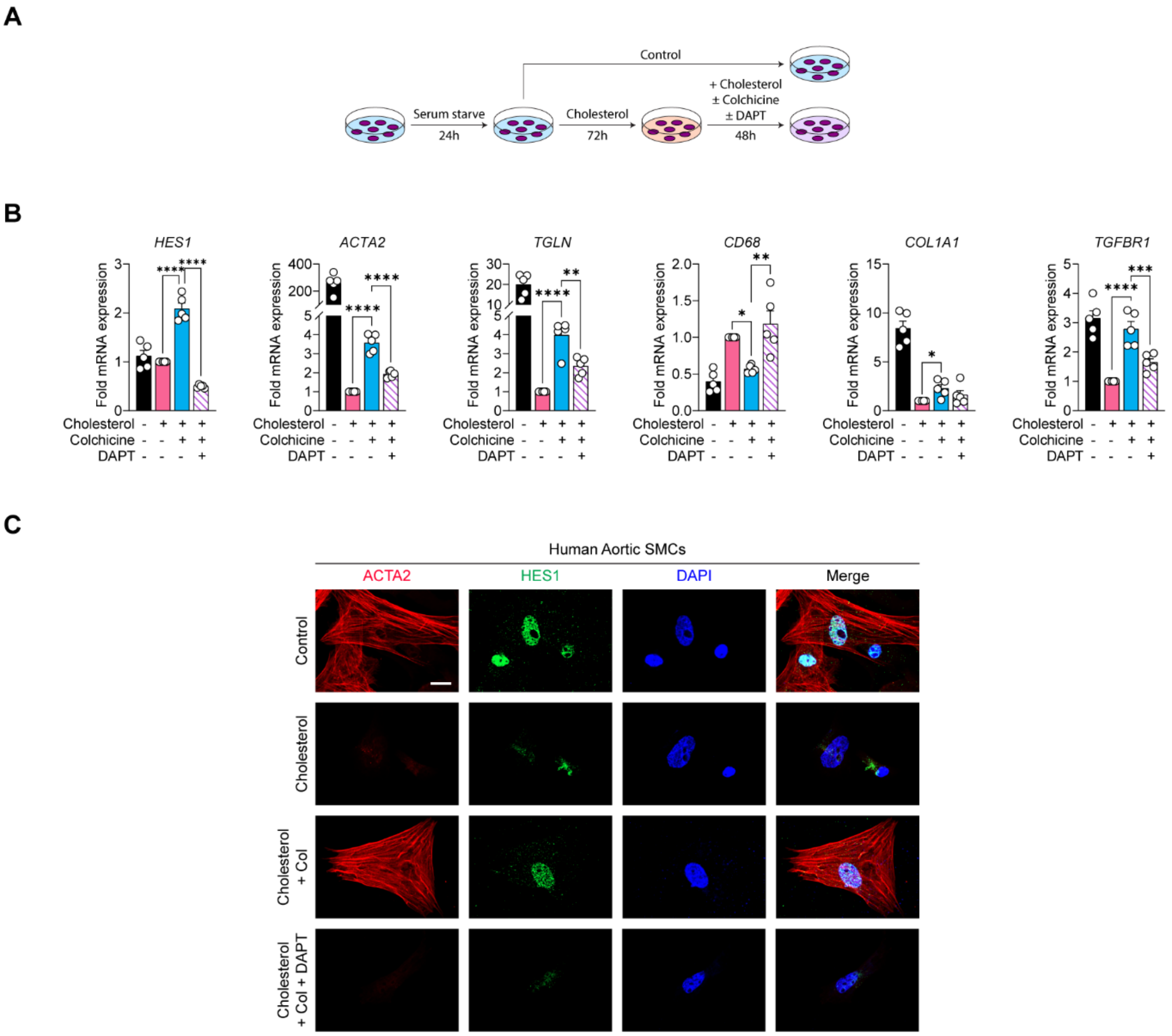

**Fig. S13. DAPT ameliorates the effects of colchicine after cholesterol loading. A,** Experimental schematic showing Notch dependent colchicine regression after long-term macrophage-like *trans*-differentiation. Briefly, HASMCs were cholesterol loaded (10µg/mL) for 72 hours followed by treatment with colchicine (50nM) and/or DAPT (5µM) for 48 hours. **B,** mRNA transcript levels of myofibroblast markers *ACTA2*, *TGLN*, *COL1A1*, macrophage marker *CD68*, the Notch signaling mediator *HES1*, and the TGFβ mediator *TGFBR1* were analyzed with qPCR. **C,** Representative images of cultured HASMCs showing protein expression of ACTA2 and HES1. Col, Colchicine. Scale bars: 20µm. Data was analyzed using a one-way ANOVA with Sidak correction and multiple comparisons. Individual dots represent independent experiments. Graphs show mean ± SEM.

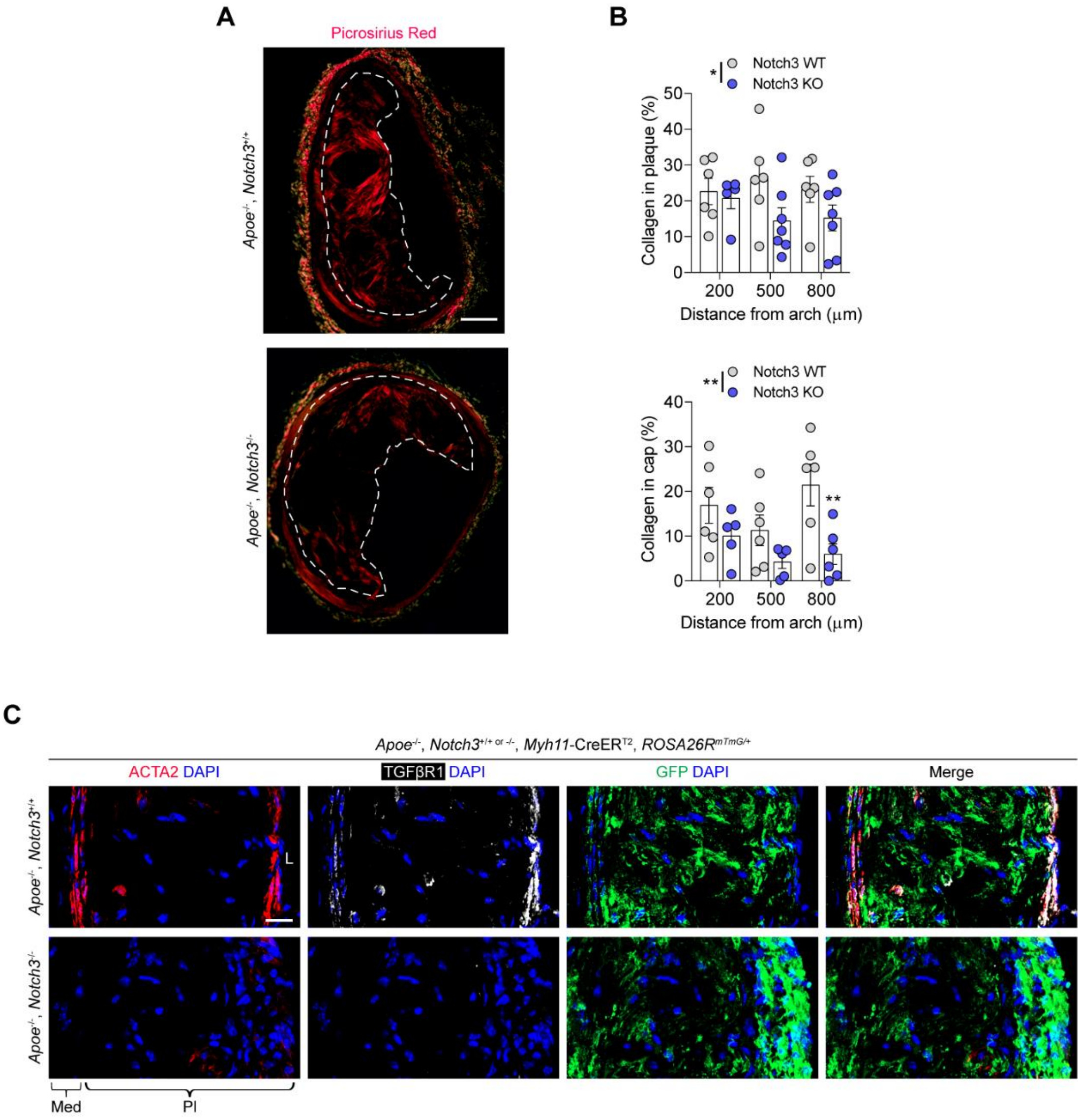

**Fig. S14. Deletion of *Notch3* markedly reduces both collagen content and TGFβR1 within atherosclerotic lesions. A, B,** Representative Picrosirius Red staining (**A**) and associated quantification (**B**) demonstrates a reduction in collagen content within brachiocephalic artery plaques ($p$ = 0.0311) and their fibrous caps ($p$ = 0.0014) in *Apoe*$^{-/-}$ *Notch3*$^{-/-}$ mice. **C,** Representative image demonstrating reduced TGFβR1 expression in aortic root plaques in the absence of *Notch3* (n = 9 mice for WT and n = 10 mice for KO). L, lumen; Med, tunica media; Pl, plaque. Scale bars: 100μm (**A**), 20μm (**C**). Data was analyzed using a two-way ANOVA with Sidak correction and multiple comparisons. Individual dots represent biologically independent animals. Graphs show mean ± SEM. *P* values refer to two-way ANOVA between genotypes.

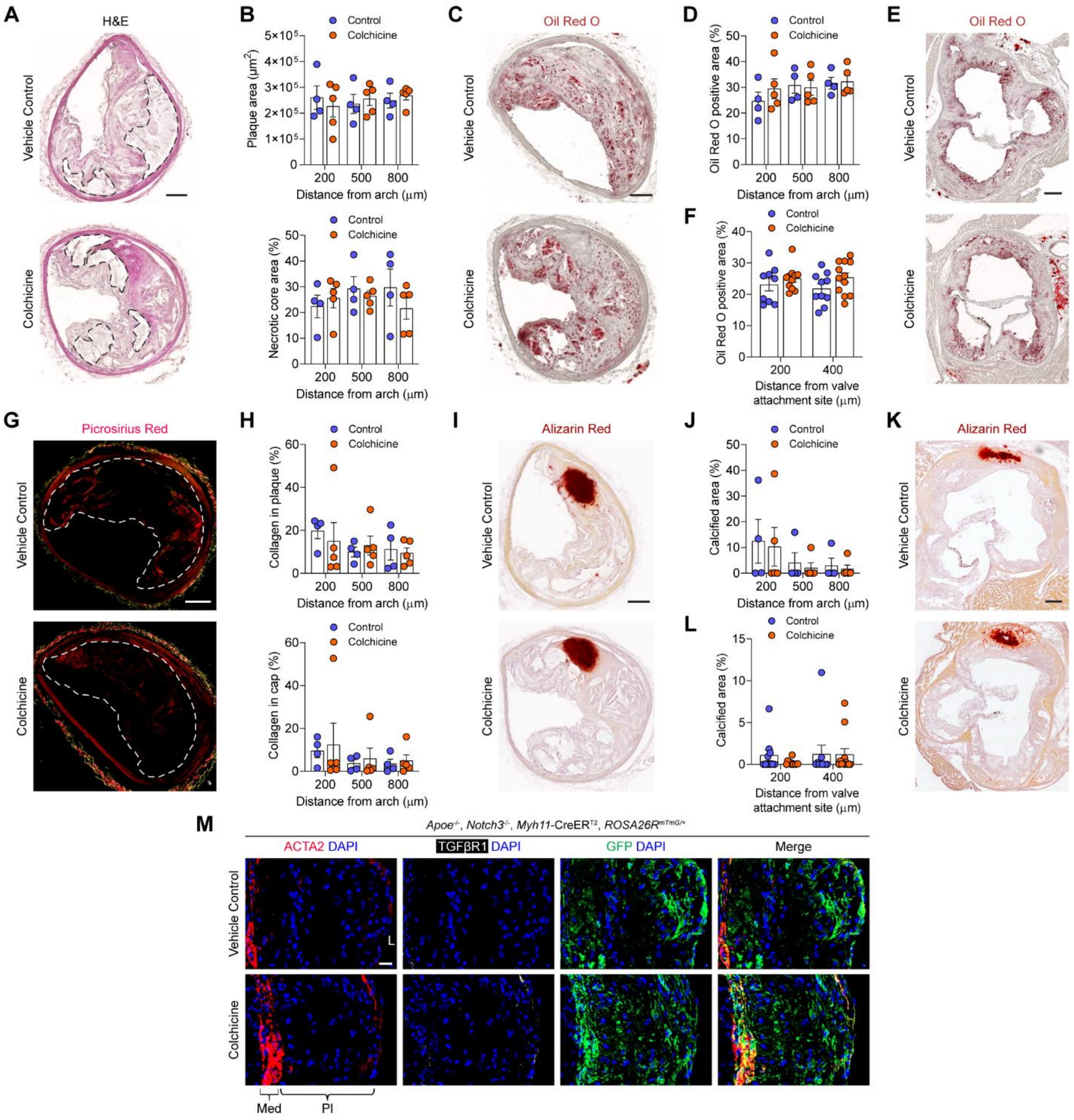

**Fig. S15. Colchicine intervention in *Apoe*$^{-/-}$, *Notch3*$^{-/-}$ mice does not lead to any enhancement in measures of plaque stability. A,** Representative images of hematoxylin and eosin staining of BCA plaques with necrotic core highlighted. **B,** Quantification of these lesions revealed no change in total lesion area ($p$ = 0.9533), nor necrotic core area ($p$ = 0.4775) after colchicine intervention. **C, D, E, F,** Colchicine treatment did not influence intraplaque lipid and triglyceride levels in the BCA ($p$ = 0.5601) or aortic root ($p$ = 0.0848), assessed by Oil Red O staining (**C, E**) and associated quantification (**D, F**). **G, H,** Collagen deposition was unchanged in BCA lesions ($p$ = 0.7844) and their fibrous caps ($p$ = 0.6415), evident by Picrosirius Red staining (**G**) and quantification (**H**). **I, J, K, L,** Intraplaque calcification was not altered after colchicine intervention in the BCA ($p$ = 0.6673) or aortic root ($p$ = 0.5068), as indicated by Alizarin Red staining (**I, K**), and its corresponding quantification (**J, L**). **M,** Representative image of TGFβR1 expression after colchicine intervention (n = 10 mice for each group). L, lumen; Med, tunica media; Pl, plaque. Scale bars: 200μm (**E, K**), 100μm (**A, C, G, I**), 20μm (**M**). Data was analyzed using a two-way ANOVA with Sidak correction and multiple comparisons. Individual dots represent biologically independent animals. Graphs show mean ± SEM. *P* values refer to two-way ANOVA between treatment conditions.

**Table S1. Differentially expressed genes regulated by cholesterol loading compared to control HASMCs.** HASMCs were cholesterol loaded (10µg/mL) for 120 hours.

| ENSEMBL | name | LogFC_Chol | padj_Chol |
|---|---|---|---|
| ENSG00000001630.17 | CYP51A1 | -1.75593 | ######## |
| ENSG00000003989.18 | SLC7A2 | 1.507128 | 1.14E-54 |
| ENSG00000006451.8 | RALA | 0.802731 | 3.59E-60 |
| ENSG00000007944.15 | MYLIP | 0.889714 | 2.99E-35 |
| ENSG00000008118.10 | CAMK1G | 1.099378 | 4.23E-19 |
| ENSG00000008256.16 | CYTH3 | -0.59329 | 3.64E-31 |
| ENSG00000011347.10 | SYT7 | 1.509904 | 1.07E-23 |
| ENSG00000011422.12 | PLAUR | 0.796832 | 4.08E-27 |
| ENSG00000012171.20 | SEMA3B | -0.61616 | 6.79E-06 |
| ENSG00000017427.17 | IGF1 | -2.18954 | 2.96E-19 |
| ENSG00000021645.20 | NRXN3 | -0.67308 | 3.45E-24 |
| ENSG00000025434.19 | NR1H3 | 1.426265 | 5.75E-50 |
| ENSG00000025708.14 | TYMP | 0.58989 | 0.002219 |
| ENSG00000026508.20 | CD44 | 0.715084 | 2.85E-63 |
| ENSG00000028137.19 | TNFRSF1B | 1.056843 | 9.01E-26 |
| ENSG00000028277.21 | POU2F2 | 0.612606 | 0.000151 |
| ENSG00000037897.17 | METTL1 | 0.729352 | 1.10E-07 |
| ENSG00000041353.10 | RAB27B | 2.089876 | 1.14E-54 |
| ENSG00000042062.13 | RIPOR3 | 1.671901 | 5.43E-28 |
| ENSG00000049540.18 | ELN | -0.83508 | 3.21E-14 |
| ENSG00000050730.16 | TNIP3 | 1.55653 | 1.53E-05 |
| ENSG00000051523.11 | CYBA | 1.819508 | ######## |
| ENSG00000052802.13 | MSMO1 | -2.73315 | 0 |
| ENSG00000054277.14 | OPN3 | -0.6244 | 1.01E-11 |
| ENSG00000054654.19 | SYNE2 | -0.73922 | 1.85E-22 |
| ENSG00000055813.6 | CCDC85A | -0.72502 | 1.60E-11 |
| ENSG00000056972.20 | TRAF3IP2 | 0.785732 | 1.05E-17 |
| ENSG00000058404.20 | CAMK2B | 1.83096 | 8.09E-16 |
| ENSG00000059728.11 | MXD1 | 0.677615 | 3.16E-21 |
| ENSG00000064205.11 | CCN5 | -0.63714 | 0.049028 |
| ENSG00000064692.20 | SNCAIP | -0.89218 | 4.63E-11 |
| ENSG00000065413.20 | ANKRD44 | -0.77985 | 5.87E-15 |
| ENSG00000065534.19 | MYLK | -0.71275 | 1.26E-36 |
| ENSG00000065911.13 | MTHFD2 | 0.852222 | 6.02E-33 |
| ENSG00000067064.12 | IDI1 | -2.04484 | ######## |
| ENSG00000068781.21 | STON1-GTF2A1L | -0.91592 | 1.09E-07 |
| ENSG00000069431.13 | ABCC9 | -0.77707 | 1.91E-18 |
| ENSG00000071282.12 | LMCD1 | -1.72859 | 1.70E-66 |
| ENSG00000072163.20 | LIMS2 | -0.8329 | 5.25E-13 |
| ENSG00000072201.14 | LNX1 | -0.5994 | 0.00017 |
| ENSG00000072310.18 | SREBF1 | 0.64891 | 1.54E-41 |
| ENSG00000072422.17 | RHOBTB1 | -0.70782 | 1.77E-21 |
| ENSG00000072952.20 | IRAG1 | -0.82541 | 4.10E-07 |
| ENSG00000073756.13 | PTGS2 | 2.00564 | 4.32E-89 |
| ENSG00000073910.23 | FRY | -1.02563 | 9.47E-38 |
| ENSG00000074410.14 | CA12 | 1.420379 | 2.12E-85 |
| ENSG00000074527.13 | NTN4 | 0.648495 | 0.005185 |
| ENSG00000074590.14 | NUAK1 | -0.8084 | 4.19E-44 |
| ENSG00000075223.14 | SEMA3C | -0.76872 | 1.76E-34 |
| ENSG00000075651.16 | PLD1 | 0.596142 | 9.19E-16 |
| ENSG00000077238.14 | IL4R | 0.712768 | 2.18E-28 |
| ENSG00000078018.21 | MAP2 | -0.60894 | 2.76E-05 |
| ENSG00000078081.8 | LAMP3 | 0.664373 | 0.020027 |
| ENSG00000079459.13 | FDFT1 | -1.97998 | 0 |

| | | | |
|---|---|---|---|
| ENSG00000080031.10 | PTPRH | 1.522995 | 2.22E-08 |
| ENSG00000080493.18 | SLC4A4 | -0.60357 | 5.77E-09 |
| ENSG00000081041.9 | CXCL2 | 2.527451 | 5.25E-58 |
| ENSG00000081059.20 | TCF7 | 0.818543 | 2.59E-05 |
| ENSG00000081320.11 | STK17B | -0.62832 | 1.82E-20 |
| ENSG00000082497.12 | SERTAD4 | -1.0862 | 9.33E-11 |
| ENSG00000085117.12 | CD82 | 0.588561 | 1.38E-06 |
| ENSG00000086991.13 | NOX4 | -1.974 | ######## |
| ENSG00000087074.8 | PPP1R15A | 0.603204 | 9.50E-25 |
| ENSG00000087301.9 | TXNDC16 | -0.75286 | 3.33E-21 |
| ENSG00000087303.18 | NID2 | -0.62145 | 3.64E-35 |
| ENSG00000088448.14 | ANKRD10 | 0.611192 | 9.62E-32 |
| ENSG00000090339.9 | ICAM1 | 1.499595 | 3.41E-30 |
| ENSG00000090376.11 | IRAK3 | 0.945783 | 1.95E-26 |
| ENSG00000091844.8 | RGS17 | 1.058998 | 4.19E-18 |
| ENSG00000091879.14 | ANGPT2 | 1.458962 | 3.99E-43 |
| ENSG00000091986.16 | CCDC80 | -0.73765 | 2.93E-64 |
| ENSG00000094804.12 | CDC6 | 0.787807 | 0.005317 |
| ENSG00000095303.17 | PTGS1 | -0.67666 | 8.16E-22 |
| ENSG00000097096.9 | SYDE2 | -0.88424 | 7.65E-12 |
| ENSG00000099194.6 | SCD | -0.94698 | ######## |
| ENSG00000099204.20 | ABLIM1 | -1.23588 | 4.12E-17 |
| ENSG00000099251.14 | HSD17B7P2 | -1.60828 | 1.32E-07 |
| ENSG00000099849.15 | RASSF7 | -0.68439 | 0.002828 |
| ENSG00000099860.9 | GADD45B | -0.88409 | 2.57E-28 |
| ENSG00000099998.19 | GGT5 | 1.463014 | 2.81E-15 |
| ENSG00000100092.24 | SH3BP1 | 0.738122 | 4.70E-31 |
| ENSG00000100100.13 | PIK3IP1 | -0.64902 | 3.63E-28 |
| ENSG00000100234.12 | TIMP3 | -0.84934 | 1.01E-58 |
| ENSG00000100344.11 | PNPLA3 | -1.49007 | 7.16E-13 |
| ENSG00000100490.10 | CDKL1 | -1.42081 | 2.06E-21 |
| ENSG00000100739.11 | BDKRB1 | 0.922451 | 1.99E-28 |
| ENSG00000100906.11 | NFKBIA | 0.635777 | 1.31E-26 |
| ENSG00000101057.16 | MYBL2 | 1.825633 | 3.57E-07 |
| ENSG00000101098.13 | RIMS4 | -3.46295 | 1.95E-71 |
| ENSG00000101255.12 | TRIB3 | 0.812678 | 2.85E-32 |
| ENSG00000101265.16 | RASSF2 | -0.68757 | 6.71E-37 |
| ENSG00000101311.16 | FERMT1 | -0.72898 | 1.40E-09 |
| ENSG00000101680.15 | LAMA1 | 0.664864 | 2.76E-12 |
| ENSG00000102452.18 | NALCN | -0.61187 | 0.000241 |
| ENSG00000102760.13 | RGCC | -0.6371 | 0.000449 |
| ENSG00000102802.10 | MEDAG | 0.830037 | 1.36E-49 |
| ENSG00000103044.11 | HAS3 | 1.337024 | 5.36E-09 |
| ENSG00000103241.7 | FOXF1 | 0.78647 | 1.31E-14 |
| ENSG00000103257.9 | SLC7A5 | 0.67673 | 2.83E-30 |
| ENSG00000103316.12 | CRYM | 1.406691 | 0.001977 |
| ENSG00000103485.19 | QPRT | -0.81175 | 1.23E-22 |
| ENSG00000103647.13 | CORO2B | 0.689196 | 1.36E-24 |
| ENSG00000104312.8 | RIPK2 | 0.617224 | 3.48E-12 |
| ENSG00000104368.19 | PLAT | 0.917672 | 7.20E-61 |
| ENSG00000104490.18 | NCALD | -0.96785 | 1.95E-24 |
| ENSG00000104549.12 | SQLE | -1.70351 | ######## |
| ENSG00000104635.15 | SLC39A14 | 1.376927 | 6.07E-88 |
| ENSG00000104856.14 | RELB | 0.585498 | 1.84E-09 |
| ENSG00000104951.16 | IL4I1 | 1.183607 | 6.75E-07 |
| ENSG00000105185.12 | PDCD5 | 0.726119 | 4.50E-09 |
| ENSG00000105792.19 | CFAP69 | 1.051793 | 1.06E-06 |
| ENSG00000105835.13 | NAMPT | 1.526776 | 4.30E-32 |
| ENSG00000105963.15 | ADAP1 | 0.804795 | 8.99E-13 |
| ENSG00000105976.16 | MET | -0.64387 | 1.64E-25 |
| ENSG00000105989.10 | WNT2 | -1.30789 | 2.90E-11 |
| ENSG00000106003.13 | LFNG | -0.74776 | 0.003649 |
| ENSG00000106366.9 | SERPINE1 | 0.825275 | 2.22E-10 |

| | | | |
|---|---|---|---|
| ENSG00000106483.12 | SFRP4 | -0.61958 | 2.96E-22 |
| ENSG00000106511.6 | MEOX2 | -0.60306 | 0.000898 |
| ENSG00000106538.10 | RARRES2 | -1.55181 | 1.40E-12 |
| ENSG00000106772.18 | PRUNE2 | -0.74575 | 1.66E-08 |
| ENSG00000106809.11 | OGN | -0.99438 | 5.99E-39 |
| ENSG00000106819.13 | ASPN | -1.7242 | 1.07E-82 |
| ENSG00000106823.12 | ECM2 | -1.33418 | 3.38E-19 |
| ENSG00000107104.20 | KANK1 | -0.94084 | 2.73E-47 |
| ENSG00000107796.13 | ACTA2 | -1.04457 | ######## |
| ENSG00000107968.10 | MAP3K8 | 0.742971 | 5.91E-17 |
| ENSG00000108342.13 | CSF3 | 6.062417 | 2.65E-51 |
| ENSG00000108576.10 | SLC6A4 | -1.48366 | 1.67E-13 |
| ENSG00000108688.11 | CCL7 | 2.728612 | 1.70E-10 |
| ENSG00000108691.9 | CCL2 | 1.530104 | 8.03E-23 |
| ENSG00000108798.9 | ABI3 | 0.721455 | 0.048259 |
| ENSG00000108821.14 | COL1A1 | -0.89182 | 2.36E-69 |
| ENSG00000108950.12 | FAM20A | 1.088559 | 2.11E-07 |
| ENSG00000109084.14 | TMEM97 | -1.46983 | 1.81E-75 |
| ENSG00000109107.14 | ALDOC | -1.03951 | 5.41E-34 |
| ENSG00000109686.19 | SH3D19 | -0.6307 | 1.44E-38 |
| ENSG00000109705.8 | NKX3-2 | -1.1929 | 3.06E-07 |
| ENSG00000109846.9 | CRYAB | -0.72138 | 1.53E-09 |
| ENSG00000109929.10 | SC5D | -1.41539 | ######## |
| ENSG00000110031.13 | LPXN | 0.708657 | 3.23E-09 |
| ENSG00000110921.14 | MVK | -1.85427 | ######## |
| ENSG00000111057.11 | KRT18 | -0.7711 | 1.28E-46 |
| ENSG00000111087.10 | GLI1 | 1.542 | 5.96E-21 |
| ENSG00000111341.10 | MGP | -0.73999 | 5.46E-53 |
| ENSG00000111452.13 | ADGRD1 | 1.029867 | 0.000149 |
| ENSG00000111799.22 | COL12A1 | -0.66398 | 1.01E-43 |
| ENSG00000111859.17 | NEDD9 | -0.60729 | 0.000405 |
| ENSG00000112096.19 | SOD2 | 1.941202 | 6.88E-56 |
| ENSG00000112139.17 | MDGA1 | -0.87284 | 2.57E-16 |
| ENSG00000112182.15 | BACH2 | -0.656 | 4.08E-06 |
| ENSG00000112559.15 | MDFI | -0.59854 | 0.0002 |
| ENSG00000112715.25 | VEGFA | 0.999085 | 5.65E-55 |
| ENSG00000112759.19 | SLC29A1 | -0.71286 | 3.97E-15 |
| ENSG00000112972.15 | HMGCS1 | -2.8746 | 0 |
| ENSG00000113140.11 | SPARC | -0.9505 | 3.69E-71 |
| ENSG00000113161.17 | HMGCR | -2.0472 | 0 |
| ENSG00000113389.16 | NPR3 | -1.203 | 1.91E-41 |
| ENSG00000113448.19 | PDE4D | 0.612913 | 4.51E-10 |
| ENSG00000113645.15 | WWC1 | 0.812314 | 1.37E-10 |
| ENSG00000113739.11 | STC2 | 0.946871 | 3.10E-61 |
| ENSG00000114200.10 | BCHE | -1.0541 | 9.18E-28 |
| ENSG00000114698.15 | PLSCR4 | -0.62843 | 8.32E-42 |
| ENSG00000115008.6 | IL1A | 2.22202 | 1.32E-60 |
| ENSG00000115009.13 | CCL20 | 3.475487 | 1.82E-08 |
| ENSG00000115232.14 | ITGA4 | -0.95829 | 7.79E-60 |
| ENSG00000115290.10 | GRB14 | -0.99726 | 7.53E-05 |
| ENSG00000115504.15 | EHBP1 | -1.00843 | 2.19E-45 |
| ENSG00000115525.18 | ST3GAL5 | 0.601541 | 4.10E-31 |
| ENSG00000115556.14 | PLCD4 | -0.76167 | 0.000172 |
| ENSG00000115594.12 | IL1R1 | 0.960527 | 2.72E-92 |
| ENSG00000116032.5 | GRIN3B | -0.5908 | 3.88E-06 |
| ENSG00000116133.13 | DHCR24 | -2.23179 | 0 |
| ENSG00000116183.11 | PAPPA2 | 0.684567 | 0.001114 |
| ENSG00000116194.13 | ANGPTL1 | 0.875898 | 3.56E-09 |
| ENSG00000116711.10 | PLA2G4A | 0.700729 | 3.78E-13 |
| ENSG00000116774.12 | OLFML3 | -0.78713 | 6.56E-53 |
| ENSG00000117115.13 | PADI2 | -1.99345 | 1.13E-56 |
| ENSG00000117461.15 | PIK3R3 | -0.72391 | 5.83E-21 |
| ENSG00000117525.14 | F3 | 1.113438 | 3.58E-14 |

| | | | |
|---|---|---|---|
| ENSG00000117594.10 | HSD11B1 | 1.664668 | 2.30E-41 |
| ENSG00000118369.13 | USP35 | -0.72378 | 3.94E-17 |
| ENSG00000118503.15 | TNFAIP3 | 1.311264 | 2.08E-95 |
| ENSG00000118523.6 | CCN2 | -0.68633 | 5.00E-38 |
| ENSG00000118526.7 | TCF21 | 0.609618 | 9.19E-09 |
| ENSG00000119280.17 | C1orf198 | -0.65042 | 1.00E-46 |
| ENSG00000119681.12 | LTBP2 | -0.93203 | 3.87E-93 |
| ENSG00000119714.11 | GPR68 | 1.625589 | 2.37E-27 |
| ENSG00000119900.9 | OGFRL1 | 0.75711 | 1.11E-29 |
| ENSG00000119938.9 | PPP1R3C | -1.03806 | 4.78E-50 |
| ENSG00000120437.9 | ACAT2 | -2.42795 | 0 |
| ENSG00000120802.14 | TMPO | -0.83117 | 7.27E-41 |
| ENSG00000121039.10 | RDH10 | -0.62829 | 1.95E-40 |
| ENSG00000121769.8 | FABP3 | -2.04118 | ######## |
| ENSG00000121858.11 | TNFSF10 | 1.156083 | 6.20E-07 |
| ENSG00000122121.12 | XPNPEP2 | 1.237983 | 8.05E-14 |
| ENSG00000122378.14 | PRXL2A | -1.064 | 3.35E-22 |
| ENSG00000122694.16 | GLIPR2 | -0.6603 | 5.85E-21 |
| ENSG00000122966.17 | CIT | 1.062856 | 1.25E-12 |
| ENSG00000123610.5 | TNFAIP6 | 2.03263 | 4.50E-78 |
| ENSG00000123689.6 | G0S2 | 1.407738 | 5.00E-60 |
| ENSG00000124249.7 | KCNK15 | -2.59874 | 8.54E-33 |
| ENSG00000124440.16 | HIF3A | 0.723328 | 2.16E-05 |
| ENSG00000124875.10 | CXCL6 | 2.285527 | 1.05E-48 |
| ENSG00000124920.14 | MYRF | -0.85324 | 3.54E-41 |
| ENSG00000125148.7 | MT2A | 0.783212 | 1.95E-05 |
| ENSG00000125266.8 | EFNB2 | -0.61711 | 5.13E-25 |
| ENSG00000125347.15 | IRF1 | 1.172382 | 6.22E-37 |
| ENSG00000125510.18 | OPRL1 | -0.66296 | 0.000238 |
| ENSG00000125538.12 | IL1B | 2.971414 | 7.39E-54 |
| ENSG00000125730.17 | C3 | 2.355263 | 3.33E-44 |
| ENSG00000125965.9 | GDF5 | 0.748869 | 2.93E-36 |
| ENSG00000126016.17 | AMOT | -0.64034 | 1.77E-20 |
| ENSG00000126218.12 | F10 | -0.9326 | 9.32E-14 |
| ENSG00000126861.5 | OMG | -0.73957 | 0.007305 |
| ENSG00000127863.16 | TNFRSF19 | 0.636232 | 3.88E-05 |
| ENSG00000127951.8 | FGL2 | -0.58812 | 5.74E-09 |
| ENSG00000128045.7 | RASL11B | -3.2526 | 2.14E-48 |
| ENSG00000128271.22 | ADORA2A | 0.685555 | 3.04E-07 |
| ENSG00000128284.19 | APOL3 | 0.70035 | 4.26E-05 |
| ENSG00000128340.15 | RAC2 | 0.687454 | 1.44E-30 |
| ENSG00000128342.5 | LIF | 1.804684 | 1.01E-24 |
| ENSG00000128482.16 | RNF112 | -0.61922 | 2.35E-07 |
| ENSG00000128590.5 | DNAJB9 | 0.720947 | 1.38E-14 |
| ENSG00000128594.8 | LRRC4 | -1.39821 | 7.67E-21 |
| ENSG00000128965.13 | CHAC1 | 0.86195 | 9.04E-19 |
| ENSG00000129009.13 | ISLR | -0.67809 | 3.36E-07 |
| ENSG00000129048.7 | ACKR4 | -0.70082 | 5.39E-11 |
| ENSG00000129226.14 | CD68 | 0.608218 | 1.92E-13 |
| ENSG00000130147.16 | SH3BP4 | -0.60319 | 4.19E-30 |
| ENSG00000130164.14 | LDLR | -0.90104 | 3.12E-65 |
| ENSG00000130176.8 | CNN1 | -1.03694 | 8.01E-35 |
| ENSG00000130203.10 | APOE | 0.905296 | 1.08E-11 |
| ENSG00000130513.6 | GDF15 | 1.286222 | 6.44E-69 |
| ENSG00000130600.19 | H19 | 0.778182 | 4.18E-23 |
| ENSG00000130775.16 | THEMIS2 | 0.643348 | 0.002448 |
| ENSG00000131069.20 | ACSS2 | -0.91626 | 2.54E-44 |
| ENSG00000131473.17 | ACLY | -0.79583 | 1.32E-87 |
| ENSG00000131747.15 | TOP2A | 1.667963 | 1.57E-29 |
| ENSG00000131979.20 | GCH1 | 1.520277 | 4.80E-37 |
| ENSG00000132000.14 | PODNL1 | -1.30717 | 8.35E-20 |
| ENSG00000132031.13 | MATN3 | -0.97953 | 0.000131 |
| ENSG00000132170.24 | PPARG | 0.919116 | 6.55E-05 |

| | | | |
|---|---|---|---|
| ENSG00000132196.16 | HSD17B7 | -1.72985 | 4.72E-40 |
| ENSG00000132199.20 | ENOSF1 | 0.590583 | 1.51E-12 |
| ENSG00000132205.11 | EMILIN2 | 0.864479 | 1.36E-13 |
| ENSG00000132329.11 | RAMP1 | -1.19011 | 0.000145 |
| ENSG00000133026.13 | MYH10 | -0.849 | ######## |
| ENSG00000133048.13 | CHI3L1 | 2.706199 | 6.72E-23 |
| ENSG00000133805.15 | AMPD3 | 1.239033 | 6.84E-25 |
| ENSG00000133935.7 | ERG28 | -1.55278 | ######## |
| ENSG00000134070.5 | IRAK2 | 1.168467 | 3.46E-42 |
| ENSG00000134107.5 | BHLHE40 | -0.60645 | 3.82E-24 |
| ENSG00000134324.12 | LPIN1 | -0.72171 | 1.43E-61 |
| ENSG00000134470.21 | IL15RA | 1.119361 | 3.88E-11 |
| ENSG00000134668.12 | SPOCD1 | 1.406136 | 6.37E-39 |
| ENSG00000134802.18 | SLC43A3 | 1.39887 | 2.86E-64 |
| ENSG00000134824.14 | FADS2 | -1.3165 | ######## |
| ENSG00000134871.19 | COL4A2 | -0.60284 | 7.31E-41 |
| ENSG00000135363.12 | LMO2 | 0.62014 | 0.000116 |
| ENSG00000135472.9 | FAIM2 | 0.602682 | 1.54E-20 |
| ENSG00000135547.9 | HEY2 | 0.629283 | 0.030533 |
| ENSG00000135636.15 | DYSF | -0.6354 | 8.86E-07 |
| ENSG00000135919.14 | SERPINE2 | -1.4052 | ######## |
| ENSG00000135932.11 | CAB39 | 0.755195 | 3.75E-39 |
| ENSG00000136040.9 | PLXNC1 | -1.56886 | 1.08E-60 |
| ENSG00000136048.14 | DRAM1 | 0.737987 | 5.71E-27 |
| ENSG00000136153.20 | LMO7 | -0.62732 | 5.02E-41 |
| ENSG00000136244.12 | IL6 | 2.668888 | 7.37E-19 |
| ENSG00000136546.16 | SCN7A | -0.96463 | 9.60E-17 |
| ENSG00000136689.19 | IL1RN | 2.523737 | 4.65E-15 |
| ENSG00000136960.13 | ENPP2 | 1.33931 | 1.47E-27 |
| ENSG00000136999.5 | CCN3 | 1.156622 | 3.59E-05 |
| ENSG00000137203.15 | TFAP2A | 0.77005 | 6.37E-07 |
| ENSG00000137266.15 | SLC22A23 | 0.836553 | 5.90E-20 |
| ENSG00000137309.20 | HMGA1 | 0.779718 | 5.35E-34 |
| ENSG00000137463.5 | MGARP | -0.82357 | 1.13E-17 |
| ENSG00000137801.11 | THBS1 | -0.76953 | 7.08E-67 |
| ENSG00000137804.14 | NUSAP1 | 0.723491 | 0.014125 |
| ENSG00000137872.17 | SEMA6D | 0.697366 | 0.000246 |
| ENSG00000137975.8 | CLCA2 | 0.649334 | 1.15E-07 |
| ENSG00000138061.12 | CYP1B1 | 0.702625 | 1.68E-19 |
| ENSG00000138316.11 | ADAMTS14 | 1.03124 | 1.22E-23 |
| ENSG00000138413.14 | IDH1 | -0.62851 | 1.52E-35 |
| ENSG00000138449.11 | SLC40A1 | -1.60488 | 2.69E-39 |
| ENSG00000138650.9 | PCDH10 | -0.62783 | 6.28E-35 |
| ENSG00000138735.16 | PDE5A | -0.75671 | 3.35E-33 |
| ENSG00000138759.20 | FRAS1 | -0.83016 | 4.32E-29 |
| ENSG00000138771.16 | SHROOM3 | -1.23814 | 3.11E-36 |
| ENSG00000138821.14 | SLC39A8 | 1.180203 | 1.97E-34 |
| ENSG00000139174.12 | PRICKLE1 | -0.77659 | 3.69E-13 |
| ENSG00000139269.3 | INHBE | 1.560953 | 6.51E-05 |
| ENSG00000139318.8 | DUSP6 | 1.176945 | 9.50E-39 |
| ENSG00000139364.11 | TMEM132B | 1.05434 | 1.09E-14 |
| ENSG00000139428.12 | MMAB | -1.66591 | ######## |
| ENSG00000139514.13 | SLC7A1 | 0.742435 | 1.56E-40 |
| ENSG00000139874.6 | SSTR1 | 1.440769 | 7.95E-11 |
| ENSG00000140092.15 | FBLN5 | -1.40564 | ######## |
| ENSG00000140450.9 | ARRDC4 | 0.63062 | 9.50E-07 |
| ENSG00000140538.16 | NTRK3 | 0.794342 | 1.15E-10 |
| ENSG00000141338.14 | ABCA8 | -0.60647 | 1.28E-46 |
| ENSG00000141469.18 | SLC14A1 | 1.330104 | ######## |
| ENSG00000141682.12 | PMAIP1 | 1.022184 | 1.85E-16 |
| ENSG00000142178.9 | SIK1 | -1.00436 | 1.32E-21 |
| ENSG00000143341.12 | HMCN1 | -0.66526 | 3.97E-26 |
| ENSG00000143355.16 | LHX9 | -0.75127 | 1.01E-05 |

| | | | |
|---|---|---|---|
| ENSG00000143479.18 | DYRK3 | 0.608529 | 1.23E-18 |
| ENSG00000143494.16 | VASH2 | -0.76782 | 0.000154 |
| ENSG00000143786.8 | CNIH3 | 0.706416 | 1.29E-31 |
| ENSG00000143867.7 | OSR1 | -0.62827 | 0.000509 |
| ENSG00000143878.10 | RHOB | -0.65294 | 1.11E-28 |
| ENSG00000143924.19 | EML4 | -0.64607 | 1.44E-29 |
| ENSG00000144136.11 | SLC20A1 | 0.618776 | 9.15E-39 |
| ENSG00000144802.11 | NFKBIZ | 1.282408 | 9.56E-94 |
| ENSG00000144810.16 | COL8A1 | -0.92445 | ######## |
| ENSG00000144857.15 | BOC | -0.6704 | 5.02E-05 |
| ENSG00000144891.19 | AGTR1 | 0.698386 | 0.005558 |
| ENSG00000144959.11 | NCEH1 | 0.883719 | 9.44E-21 |
| ENSG00000145358.6 | DDIT4L | 1.050706 | 1.55E-20 |
| ENSG00000145506.14 | NKD2 | 0.938855 | 9.63E-05 |
| ENSG00000145632.15 | PLK2 | -0.80858 | 4.02E-39 |
| ENSG00000145685.14 | LHFPL2 | 0.675744 | 2.85E-37 |
| ENSG00000145860.12 | RNF145 | 0.78645 | 5.81E-66 |
| ENSG00000145901.16 | TNIP1 | 0.642649 | 3.16E-24 |
| ENSG00000146122.17 | DAAM2 | -0.99319 | 1.26E-22 |
| ENSG00000146374.14 | RSPO3 | -1.13413 | 1.73E-17 |
| ENSG00000146411.6 | SLC2A12 | -0.89094 | 4.18E-15 |
| ENSG00000146457.16 | WTAP | 0.924792 | 2.55E-48 |
| ENSG00000146555.19 | SDK1 | 1.651637 | 4.42E-25 |
| ENSG00000146592.17 | CREB5 | 0.759901 | 8.74E-15 |
| ENSG00000146648.20 | EGFR | 0.741405 | 2.55E-53 |
| ENSG00000146674.16 | IGFBP3 | -0.73943 | 1.93E-44 |
| ENSG00000147100.11 | SLC16A2 | -0.63688 | 2.72E-23 |
| ENSG00000147155.11 | EBP | -1.57882 | 1.43E-86 |
| ENSG00000147383.11 | NSDHL | -1.49573 | 1.05E-73 |
| ENSG00000147872.10 | PLIN2 | 0.773743 | 1.11E-42 |
| ENSG00000148344.11 | PTGES | 1.02078 | 2.67E-30 |
| ENSG00000148541.13 | FAM13C | -0.87947 | 1.74E-12 |
| ENSG00000148677.7 | ANKRD1 | -0.68337 | 0.00013 |
| ENSG00000148680.16 | HTR7 | 1.192679 | 4.18E-08 |
| ENSG00000149212.12 | SESN3 | -0.7662 | 2.36E-28 |
| ENSG00000149289.11 | ZC3H12C | 0.765731 | 8.20E-31 |
| ENSG00000149294.17 | NCAM1 | -0.86748 | 1.11E-53 |
| ENSG00000149485.19 | FADS1 | -0.79007 | 3.42E-57 |
| ENSG00000149571.12 | KIRREL3 | 0.66505 | 0.000767 |
| ENSG00000149591.17 | TAGLN | -0.68124 | 4.61E-36 |
| ENSG00000149809.17 | TM7SF2 | -2.02635 | ######## |
| ENSG00000149968.12 | MMP3 | 0.953487 | 0.009215 |
| ENSG00000150636.17 | CCDC102B | 0.67015 | 6.98E-05 |
| ENSG00000150687.12 | PRSS23 | -1.16335 | 5.96E-67 |
| ENSG00000150907.10 | FOXO1 | -0.70384 | 6.70E-26 |
| ENSG00000151012.13 | SLC7A11 | 0.632478 | 4.03E-05 |
| ENSG00000151276.23 | MAGI1 | -0.61935 | 5.82E-23 |
| ENSG00000151474.23 | FRMD4A | 0.588224 | 8.01E-32 |
| ENSG00000151632.17 | AKR1C2 | -0.69231 | 3.33E-15 |
| ENSG00000151725.12 | CENPU | 0.664197 | 0.019162 |
| ENSG00000152217.20 | SETBP1 | -0.62397 | 1.58E-10 |
| ENSG00000153162.9 | BMP6 | -1.20131 | 4.30E-61 |
| ENSG00000153993.14 | SEMA3D | 0.68761 | 7.89E-23 |
| ENSG00000154027.19 | AK5 | -0.72395 | 4.31E-32 |
| ENSG00000154040.21 | CABYR | -0.68684 | 0.002703 |
| ENSG00000154175.18 | ABI3BP | 0.637674 | 8.27E-06 |
| ENSG00000154310.17 | TNIK | 0.658958 | 1.13E-40 |
| ENSG00000154319.16 | FAM167A | 0.862733 | 1.19E-08 |
| ENSG00000154511.12 | DIPK1A | -0.67248 | 9.07E-07 |
| ENSG00000154553.16 | PDLIM3 | -0.99389 | 6.33E-91 |
| ENSG00000154734.16 | ADAMTS1 | -1.03425 | 3.55E-12 |
| ENSG00000154839.10 | SKA1 | 0.736096 | 0.001042 |
| ENSG00000154864.13 | PIEZO2 | 0.70438 | 1.03E-08 |

| | | | |
|---|---|---|---|
| ENSG00000155304.6 | HSPA13 | 0.776903 | 1.59E-28 |
| ENSG00000155324.10 | GRAMD2B | -0.80594 | 3.39E-36 |
| ENSG00000156026.14 | MCU | -0.60328 | 2.80E-24 |
| ENSG00000156113.24 | KCNMA1 | -0.59399 | 3.77E-13 |
| ENSG00000156381.9 | ANKRD9 | -0.61865 | 1.16E-06 |
| ENSG00000156453.14 | PCDH1 | 0.754198 | 8.42E-17 |
| ENSG00000156463.18 | SH3RF2 | 0.722697 | 6.21E-12 |
| ENSG00000157168.20 | NRG1 | -1.08865 | 1.71E-08 |
| ENSG00000157214.14 | STEAP2 | 1.42068 | 5.22E-16 |
| ENSG00000157240.4 | FZD1 | -0.83469 | 2.48E-42 |
| ENSG00000157399.17 | ARSL | -0.86693 | 1.08E-05 |
| ENSG00000157510.14 | AFAP1L1 | 0.646192 | 5.57E-17 |
| ENSG00000157514.17 | TSC22D3 | -0.94319 | 1.36E-50 |
| ENSG00000157637.13 | SLC38A10 | -0.72094 | 5.27E-54 |
| ENSG00000157680.16 | DGKI | 0.639692 | 8.71E-10 |
| ENSG00000157927.17 | RADIL | -0.62382 | 1.47E-06 |
| ENSG00000158246.8 | TENT5B | -2.01715 | 1.59E-29 |
| ENSG00000158470.5 | B4GALT5 | 0.641716 | 3.86E-32 |
| ENSG00000158555.15 | GDPD5 | -0.6906 | 2.68E-14 |
| ENSG00000158966.16 | CACHD1 | -0.63675 | 5.85E-17 |
| ENSG00000158987.22 | RAPGEF6 | -0.73104 | 5.25E-25 |
| ENSG00000159167.12 | STC1 | 2.521106 | 7.21E-65 |
| ENSG00000159261.12 | CLDN14 | 1.678157 | 8.17E-08 |
| ENSG00000159399.10 | HK2 | 0.689188 | 1.09E-29 |
| ENSG00000160111.15 | CPAMD8 | -0.87163 | 6.06E-05 |
| ENSG00000160179.19 | ABCG1 | 4.263724 | 0 |
| ENSG00000160285.15 | LSS | -0.93852 | ######## |
| ENSG00000160752.15 | FDPS | -2.11272 | 0 |
| ENSG00000161544.10 | CYGB | 0.933762 | 3.34E-29 |
| ENSG00000161682.15 | FAM171A2 | -0.59852 | 2.11E-15 |
| ENSG00000161888.11 | SPC24 | 0.606498 | 0.005308 |
| ENSG00000162267.12 | ITIH3 | -1.45619 | 4.36E-07 |
| ENSG00000162490.7 | DRAXIN | 1.479306 | 9.57E-35 |
| ENSG00000162591.16 | MEGF6 | -0.84873 | 1.33E-19 |
| ENSG00000162614.19 | NEXN | -1.05082 | 7.09E-65 |
| ENSG00000162630.6 | B3GALT2 | -1.27454 | 9.95E-33 |
| ENSG00000162692.12 | VCAM1 | 0.625054 | 1.28E-20 |
| ENSG00000162772.17 | ATF3 | 1.366992 | 5.84E-15 |
| ENSG00000162849.16 | KIF26B | -0.77352 | 2.65E-53 |
| ENSG00000163131.12 | CTSS | 0.678931 | 4.11E-06 |
| ENSG00000163347.6 | CLDN1 | 2.381586 | 3.15E-33 |
| ENSG00000163376.11 | KBTBD8 | 0.684963 | 1.30E-12 |
| ENSG00000163431.13 | LMOD1 | -1.33504 | 1.08E-25 |
| ENSG00000163453.11 | IGFBP7 | -0.70568 | 5.59E-49 |
| ENSG00000163512.14 | AZI2 | 0.604325 | 2.43E-21 |
| ENSG00000163661.4 | PTX3 | 0.808191 | 3.34E-12 |
| ENSG00000163734.4 | CXCL3 | 2.467289 | 2.57E-84 |
| ENSG00000163735.7 | CXCL5 | 2.339163 | 4.18E-37 |
| ENSG00000163739.5 | CXCL1 | 2.876684 | 1.23E-61 |
| ENSG00000163814.8 | CDCP1 | 0.860829 | 7.87E-36 |
| ENSG00000163874.11 | ZC3H12A | 1.098032 | 5.89E-44 |
| ENSG00000163975.12 | MELTF | 0.77446 | 1.30E-06 |
| ENSG00000164035.10 | EMCN | 0.67814 | 0.000787 |
| ENSG00000164093.17 | PITX2 | -0.6179 | 1.75E-17 |
| ENSG00000164171.11 | ITGA2 | 0.893938 | 6.03E-71 |
| ENSG00000164176.13 | EDIL3 | -0.63598 | 4.57E-27 |
| ENSG00000164211.13 | STARD4 | -0.92183 | 1.47E-34 |
| ENSG00000164283.13 | ESM1 | 1.587367 | 5.83E-29 |
| ENSG00000164292.13 | RHOBTB3 | -0.66641 | 2.93E-42 |
| ENSG00000164574.16 | GALNT10 | -0.76067 | 3.61E-61 |
| ENSG00000164611.13 | PTTG1 | 0.755642 | 0.000695 |
| ENSG00000164619.10 | BMPER | 0.627302 | 1.16E-21 |
| ENSG00000164647.9 | STEAP1 | 1.662663 | 1.35E-51 |

| | | | |
|---|---|---|---|
| ENSG00000164659.15 | ELAPOR2 | -0.69425 | 0.002008 |
| ENSG00000164683.18 | HEY1 | 1.811527 | 5.88E-63 |
| ENSG00000164687.11 | FABP5 | -0.59727 | 9.34E-15 |
| ENSG00000164920.9 | OSR2 | -1.22428 | 9.54E-13 |
| ENSG00000164932.13 | CTHRC1 | -0.79922 | 2.39E-37 |
| ENSG00000164949.8 | GEM | 0.640261 | 6.10E-17 |
| ENSG00000165029.17 | ABCA1 | 3.209981 | 0 |
| ENSG00000165030.4 | NFIL3 | 0.681324 | 1.18E-21 |
| ENSG00000165072.10 | MAMDC2 | -0.60337 | 0.000654 |
| ENSG00000165338.17 | HECTD2 | -0.64641 | 5.06E-18 |
| ENSG00000165474.8 | GJB2 | 1.333203 | 7.32E-06 |
| ENSG00000165617.15 | DACT1 | -1.0453 | 5.48E-74 |
| ENSG00000165655.17 | ZNF503 | -0.83477 | 1.03E-48 |
| ENSG00000165868.16 | HSPA12A | -0.6526 | 9.02E-23 |
| ENSG00000165891.16 | E2F7 | 0.702414 | 6.03E-32 |
| ENSG00000166165.13 | CKB | -0.70338 | 3.03E-13 |
| ENSG00000166401.15 | SERPINB8 | 0.766112 | 1.18E-14 |
| ENSG00000166448.15 | TMEM130 | -0.78865 | 0.000135 |
| ENSG00000166803.14 | PCLAF | 1.207944 | 0.003239 |
| ENSG00000166851.15 | PLK1 | 0.814467 | 0.007648 |
| ENSG00000166897.16 | ELFN2 | -0.69893 | 7.97E-10 |
| ENSG00000167034.10 | NKX3-1 | 0.660882 | 8.39E-05 |
| ENSG00000167081.18 | PBX3 | -0.78666 | 1.34E-32 |
| ENSG00000167191.12 | GPRC5B | 0.775595 | 5.80E-30 |
| ENSG00000167508.12 | MVD | -2.23753 | 0 |
| ENSG00000167513.9 | CDT1 | 0.65067 | 0.002301 |
| ENSG00000167552.15 | TUBA1A | -0.59526 | 1.62E-18 |
| ENSG00000167799.10 | NUDT8 | -0.60079 | 0.04382 |
| ENSG00000167900.12 | TK1 | 1.087 | 0.000483 |
| ENSG00000168003.18 | SLC3A2 | 0.805711 | 5.27E-47 |
| ENSG00000168209.6 | DDIT4 | 0.726958 | 7.28E-30 |
| ENSG00000168389.18 | MFSD2A | 1.555709 | 1.27E-06 |
| ENSG00000168398.7 | BDKRB2 | 0.913665 | 1.17E-28 |
| ENSG00000168477.19 | TNXB | 0.900641 | 7.25E-25 |
| ENSG00000168497.5 | CAVIN2 | -1.03197 | 4.32E-21 |
| ENSG00000168646.13 | AXIN2 | -1.0153 | 8.11E-30 |
| ENSG00000168672.4 | LRATD2 | -1.55171 | 2.62E-50 |
| ENSG00000168679.18 | SLC16A4 | -0.68835 | 1.63E-05 |
| ENSG00000169184.6 | MN1 | -0.8446 | 1.15E-40 |
| ENSG00000169429.11 | CXCL8 | 3.825783 | 6.39E-73 |
| ENSG00000169710.9 | FASN | -0.92729 | ######## |
| ENSG00000169946.14 | ZFPM2 | -0.81951 | 2.17E-32 |
| ENSG00000170276.6 | HSPB2 | -0.69005 | 0.006219 |
| ENSG00000170312.16 | CDK1 | 1.069184 | 0.00051 |
| ENSG00000170323.9 | FABP4 | -1.10753 | 7.65E-05 |
| ENSG00000170340.11 | B3GNT2 | -0.61777 | 2.33E-11 |
| ENSG00000170390.16 | DCLK2 | -1.05852 | 1.41E-13 |
| ENSG00000170522.10 | ELOVL6 | -0.9725 | 1.13E-52 |
| ENSG00000170915.9 | PAQR8 | 0.610009 | 2.24E-11 |
| ENSG00000171241.9 | SHCBP1 | 0.854327 | 0.010146 |
| ENSG00000171246.6 | NPTX1 | 2.587191 | 4.50E-78 |
| ENSG00000171388.12 | APLN | 0.926708 | 3.80E-17 |
| ENSG00000171621.14 | SPSB1 | 0.837167 | 2.46E-44 |
| ENSG00000171729.14 | TMEM51 | 0.966574 | 2.50E-18 |
| ENSG00000171812.13 | COL8A2 | -1.06428 | 3.77E-33 |
| ENSG00000171848.16 | RRM2 | 1.716117 | 1.23E-11 |
| ENSG00000171951.5 | SCG2 | 1.099646 | 5.68E-13 |
| ENSG00000172061.9 | LRRC15 | -1.34153 | 1.25E-41 |
| ENSG00000172137.19 | CALB2 | 1.378056 | 1.44E-06 |
| ENSG00000172156.4 | CCL11 | 2.146023 | 4.68E-08 |
| ENSG00000172183.16 | ISG20 | 0.78994 | 0.000402 |
| ENSG00000172201.12 | ID4 | -0.76421 | 5.36E-06 |
| ENSG00000172432.19 | GTPBP2 | 0.670715 | 6.60E-29 |

| | | | |
|---|---|---|---|
| ENSG00000172458.5 | IL17D | -0.80379 | 2.79E-17 |
| ENSG00000172572.7 | PDE3A | -0.64295 | 1.53E-12 |
| ENSG00000172893.17 | DHCR7 | -1.99778 | ######## |
| ENSG00000173210.20 | ABLIM3 | 0.644866 | 6.90E-27 |
| ENSG00000173334.4 | TRIB1 | 0.612489 | 2.27E-07 |
| ENSG00000173918.15 | C1QTNF1 | 0.972845 | 1.05E-37 |
| ENSG00000173926.6 | MARCHF3 | 0.7461 | 1.68E-15 |
| ENSG00000174004.6 | NRROS | 0.886225 | 3.61E-08 |
| ENSG00000174028.6 | FAM3C2P | -0.85686 | 0.013899 |
| ENSG00000174721.10 | FGFBP3 | -1.15899 | 1.56E-08 |
| ENSG00000174791.11 | RIN1 | 0.614413 | 1.99E-20 |
| ENSG00000175063.17 | UBE2C | 1.169785 | 0.003787 |
| ENSG00000175183.10 | CSRP2 | -1.02287 | 6.48E-24 |
| ENSG00000175197.13 | DDIT3 | 0.754528 | 1.43E-29 |
| ENSG00000175471.19 | MCTP1 | 1.051675 | 4.09E-13 |
| ENSG00000175567.11 | UCP2 | -1.02984 | 1.27E-07 |
| ENSG00000175592.9 | FOSL1 | 1.057885 | 4.09E-26 |
| ENSG00000175832.13 | ETV4 | 0.616861 | 1.85E-07 |
| ENSG00000175874.10 | CREG2 | 2.11298 | 3.40E-14 |
| ENSG00000175906.5 | ARL4D | -0.73796 | 3.82E-10 |
| ENSG00000176406.23 | RIMS2 | -1.16969 | 8.63E-07 |
| ENSG00000176597.12 | B3GNT5 | 0.663596 | 1.09E-18 |
| ENSG00000176907.5 | TCIM | 0.860578 | 8.85E-07 |
| ENSG00000176971.4 | FIBIN | -1.1018 | 1.01E-56 |
| ENSG00000177191.2 | B3GNT8 | -0.75419 | 9.01E-05 |
| ENSG00000177374.13 | HIC1 | -0.65068 | 2.94E-25 |
| ENSG00000177426.22 | TGIF1 | 0.616118 | 4.52E-16 |
| ENSG00000177606.8 | JUN | 0.712746 | 6.41E-24 |
| ENSG00000178607.17 | ERN1 | 1.10044 | 2.04E-45 |
| ENSG00000178662.16 | CSRNP3 | -0.67791 | 8.44E-07 |
| ENSG00000178726.7 | THBD | 0.875396 | 3.94E-45 |
| ENSG00000178860.9 | MSC | 0.755244 | 6.76E-14 |
| ENSG00000179403.12 | VWA1 | -0.74378 | 2.15E-11 |
| ENSG00000180155.20 | LYNX1 | -0.92863 | 1.34E-25 |
| ENSG00000180340.7 | FZD2 | -0.8409 | 1.38E-35 |
| ENSG00000180440.4 | SERTM1 | -0.86838 | 3.04E-07 |
| ENSG00000180694.14 | TMEM64 | -0.64296 | 1.57E-17 |
| ENSG00000180914.11 | OXTR | -1.64294 | 1.74E-78 |
| ENSG00000180998.12 | GPR137C | -0.6287 | 0.00095 |
| ENSG00000182134.16 | TDRKH | -0.76389 | 0.00079 |
| ENSG00000182175.14 | RGMA | -0.5856 | 7.83E-09 |
| ENSG00000182263.14 | FIGN | -0.68106 | 4.27E-06 |
| ENSG00000182492.16 | BGN | -0.60311 | 3.80E-20 |
| ENSG00000182580.3 | EPHB3 | -1.28588 | 1.54E-55 |
| ENSG00000182752.10 | PAPPA | 0.77867 | 2.64E-52 |
| ENSG00000183087.15 | GAS6 | -0.63739 | 4.23E-51 |
| ENSG00000183111.12 | ARHGEF37 | -0.68014 | 3.14E-07 |
| ENSG00000183160.9 | TMEM119 | -0.58524 | 3.59E-21 |
| ENSG00000183570.17 | PCBP3 | 0.92212 | 1.33E-06 |
| ENSG00000183578.8 | TNFAIP8L3 | 0.592353 | 4.76E-14 |
| ENSG00000183598.4 | H3C13 | 0.799686 | 0.03665 |
| ENSG00000183715.14 | OPCML | 0.703781 | 2.02E-05 |
| ENSG00000183876.9 | ARSI | -0.5966 | 0.000145 |
| ENSG00000184227.8 | ACOT1 | -0.72386 | 9.38E-08 |
| ENSG00000184254.17 | ALDH1A3 | 0.65703 | 2.04E-35 |
| ENSG00000184357.5 | H1-5 | 1.002201 | 1.07E-14 |
| ENSG00000184524.6 | CEND1 | 0.636873 | 0.0225 |
| ENSG00000184702.20 | SEPTIN5 | -0.731 | 1.20E-15 |
| ENSG00000185022.12 | MAFF | 0.740065 | 4.22E-17 |
| ENSG00000185215.11 | TNFAIP2 | 1.261229 | 1.98E-64 |
| ENSG00000185338.7 | SOCS1 | 1.104497 | 2.86E-25 |
| ENSG00000185483.12 | ROR1 | -0.73269 | 1.73E-20 |
| ENSG00000185585.20 | OLFML2A | 0.591902 | 2.31E-09 |

| | | | |
|---|---|---|---|
| ENSG00000185813.11 | PCYT2 | -0.75061 | 2.41E-34 |
| ENSG00000185862.7 | EVI2B | 0.68072 | 0.033584 |
| ENSG00000186480.13 | INSIG1 | -1.96391 | 0 |
| ENSG00000186765.12 | FSCN2 | -0.7763 | 0.014295 |
| ENSG00000186907.8 | RTN4RL2 | 1.464744 | 1.86E-20 |
| ENSG00000187479.8 | C11orf96 | 1.137426 | 0.000424 |
| ENSG00000187498.16 | COL4A1 | -0.64679 | 1.57E-46 |
| ENSG00000187634.13 | SAMD11 | -1.19461 | 8.18E-06 |
| ENSG00000188312.14 | CENPP | -0.59762 | 4.01E-12 |
| ENSG00000189184.12 | PCDH18 | -0.69293 | 5.26E-37 |
| ENSG00000189337.17 | KAZN | 0.638579 | 4.67E-14 |
| ENSG00000189367.15 | KIAA0408 | -0.63756 | 0.007903 |
| ENSG00000189409.14 | MMP23B | -0.81958 | 0.002283 |
| ENSG00000196139.14 | AKR1C3 | -0.81534 | 2.85E-09 |
| ENSG00000196155.13 | PLEKHG4 | 0.597037 | 0.000435 |
| ENSG00000196159.14 | FAT4 | -0.63276 | 5.53E-23 |
| ENSG00000196460.14 | RFX8 | 1.170314 | 9.66E-09 |
| ENSG00000196549.13 | MME | 1.552928 | ######## |
| ENSG00000196562.14 | SULF2 | -0.75425 | 1.41E-33 |
| ENSG00000196611.6 | MMP1 | 1.480193 | ######## |
| ENSG00000196639.7 | HRH1 | 0.597535 | 1.55E-19 |
| ENSG00000196878.15 | LAMB3 | 1.191185 | 1.26E-25 |
| ENSG00000196972.9 | SMIM10L2B | -0.85601 | 0.00017 |
| ENSG00000197442.10 | MAP3K5 | 0.70632 | 1.43E-12 |
| ENSG00000197461.13 | PDGFA | -0.6082 | 4.41E-06 |
| ENSG00000197467.17 | COL13A1 | 0.943588 | 1.60E-34 |
| ENSG00000197614.11 | MFAP5 | -0.61711 | 4.49E-25 |
| ENSG00000197632.9 | SERPINB2 | 2.79964 | 9.47E-38 |
| ENSG00000197646.8 | PDCD1LG2 | 0.605517 | 4.42E-06 |
| ENSG00000197696.10 | NMB | 0.69737 | 0.001434 |
| ENSG00000197977.4 | ELOVL2 | 0.697981 | 1.20E-12 |
| ENSG00000198121.14 | LPAR1 | 0.631886 | 1.72E-41 |
| ENSG00000198429.10 | ZNF69 | -0.65496 | 1.75E-05 |
| ENSG00000198523.6 | PLN | -0.82039 | 0.000156 |
| ENSG00000198542.14 | ITGBL1 | -0.76046 | 9.91E-56 |
| ENSG00000198814.13 | GK | 0.711585 | 4.56E-10 |
| ENSG00000198855.7 | FICD | 0.670321 | 2.91E-10 |
| ENSG00000198911.12 | SREBF2 | -0.91233 | 8.61E-76 |
| ENSG00000198947.18 | DMD | -0.91024 | 9.22E-28 |
| ENSG00000202538.1 | RNU4-2 | -0.64154 | 0.0016 |
| ENSG00000203618.6 | GP1BB | -1.67887 | 9.92E-05 |
| ENSG00000203668.3 | CHML | -0.59114 | 2.70E-10 |
| ENSG00000203761.5 | MSTO2P | 0.968178 | 0.002695 |
| ENSG00000203811.1 | H3C14 | 2.280626 | 1.15E-15 |
| ENSG00000203852.3 | H3C15 | 2.280626 | 1.15E-15 |
| ENSG00000204257.15 | HLA-DMA | -0.94754 | 0.003956 |
| ENSG00000204520.14 | MICA | -0.89121 | 2.51E-28 |
| ENSG00000204682.8 | MIR1915HG | -0.62456 | 0.002408 |
| ENSG00000205213.14 | LGR4 | -0.63682 | 1.31E-22 |
| ENSG00000211445.12 | GPX3 | -0.59839 | 0.036085 |
| ENSG00000213626.13 | LBH | -1.26075 | 1.13E-14 |
| ENSG00000213694.6 | S1PR3 | -0.66838 | 1.97E-20 |
| ENSG00000214967.5 | NPIPA7 | 0.751922 | 0.047691 |
| ENSG00000215424.10 | MCM3AP-AS1 | -0.77006 | 0.000121 |
| ENSG00000219438.9 | TAFA5 | 0.625537 | 0.002451 |
| ENSG00000221818.9 | EBF2 | -0.62142 | 3.22E-12 |
| ENSG00000221963.6 | APOL6 | 0.731972 | 6.88E-21 |
| ENSG00000225177.6 | RP11-390P2.4 | 0.875286 | 0.004336 |
| ENSG00000226380.9 | AC058791.1 | 0.687358 | 0.003761 |
| ENSG00000226964.1 | RHEBP2 | -0.97867 | 0.016928 |
| ENSG00000227544.10 | AC018647.3 | -1.17082 | 0.005864 |
| ENSG00000228649.9 | SNHG26 | 0.70307 | 0.008898 |
| ENSG00000229152.2 | ANKRD10-IT1 | 0.858147 | 0.003437 |

| | | | |
|---|---|---|---|
| ENSG00000229644.6 | NAMPTP1 | 1.329307 | 3.19E-13 |
| ENSG00000230615.7 | RP5-1198O20.4 | 0.65484 | 0.029878 |
| ENSG00000231187.3 | RP11-38L15.3 | -1.18822 | 4.55E-22 |
| ENSG00000232679.2 | LINC01705 | 2.089961 | 0.000229 |
| ENSG00000233098.9 | CCDC144NL-AS1 | -0.71204 | 0.020825 |
| ENSG00000235314.1 | LINC00957 | -0.7434 | 0.00869 |
| ENSG00000235513.2 | L3MBTL2-AS1 | 1.017436 | 8.94E-10 |
| ENSG00000236824.2 | BCYRN1 | 0.912232 | 0.04691 |
| ENSG00000237649.8 | KIFC1 | 0.863714 | 0.011534 |
| ENSG00000239521.9 | CASTOR3 | -0.61118 | 4.37E-16 |
| ENSG00000239704.11 | CDRT4 | 2.369965 | 1.32E-08 |
| ENSG00000240065.8 | PSMB9 | 0.606807 | 0.015796 |
| ENSG00000240694.9 | PNMA2 | 0.587314 | 2.95E-07 |
| ENSG00000241644.2 | INMT | -1.96984 | 3.30E-15 |
| ENSG00000242759.8 | LINC00882 | -0.82146 | 7.63E-05 |
| ENSG00000243244.7 | STON1 | -1.08213 | 2.83E-56 |
| ENSG00000243649.9 | CFB | 1.670554 | 3.99E-16 |
| ENSG00000243742.5 | RPLP0P2 | 2.106738 | 5.69E-17 |
| ENSG00000248869.7 | LINC02511 | -0.69452 | 0.014015 |
| ENSG00000249624.10 | AP000295.9 | 0.93772 | 0.035354 |
| ENSG00000249992.2 | TMEM158 | 0.879933 | 6.20E-44 |
| ENSG00000250273.1 | PSMC1P5 | -2.02946 | 0.003437 |
| ENSG00000250320.6 | EDIL3-DT | -0.86843 | 6.68E-09 |
| ENSG00000253161.5 | LINC01605 | 0.822514 | 5.13E-05 |
| ENSG00000253227.2 | RP11-383J24.1 | 2.035957 | 7.57E-06 |
| ENSG00000254535.4 | PABPC4L | -0.65615 | 9.55E-07 |
| ENSG00000254682.1 | RP11-660L16.2 | -0.72035 | 0.009771 |
| ENSG00000254806.5 | SYS1-DBNDD2 | 1.452949 | 0.001167 |
| ENSG00000255284.2 | AP006621.5 | -0.65694 | 0.001416 |
| ENSG00000255690.3 | TRIL | -1.62381 | 4.25E-41 |
| ENSG00000258791.9 | LINC00520 | 1.621374 | 0.000454 |
| ENSG00000258864.1 | CTC-554D6.1 | 0.738525 | 0.047126 |
| ENSG00000259171.1 | RP11-903H12.5 | -0.90005 | 0.002646 |
| ENSG00000259316.12 | CTD-2116N17.1 | 1.348003 | 0.012788 |
| ENSG00000260628.5 | RP11-1166P10.1 | -0.81126 | 0.028129 |
| ENSG00000260630.7 | SNAI3-AS1 | -1.33948 | 2.63E-58 |
| ENSG00000261040.7 | WFDC21P | 0.631822 | 0.001866 |
| ENSG00000261064.1 | LINC02256 | -1.43078 | 0.012048 |
| ENSG00000261114.1 | RP11-325K4.2 | 1.215006 | 0.017606 |
| ENSG00000261327.5 | RP11-863P13.3 | 0.597047 | 0.000669 |
| ENSG00000261371.6 | PECAM1 | 0.762234 | 4.60E-11 |
| ENSG00000262001.1 | DLGAP1-AS2 | 1.032103 | 4.60E-06 |
| ENSG00000262580.5 | RP11-334C17.5 | 0.927603 | 0.00121 |
| ENSG00000263155.6 | MYZAP | -1.47025 | 1.39E-10 |
| ENSG00000263528.8 | IKBKE | 0.77501 | 3.25E-32 |
| ENSG00000267339.7 | LINC00906 | -0.73953 | 8.35E-07 |
| ENSG00000268105.1 | RP11-369G6.2 | 0.904425 | 0.011348 |
| ENSG00000268388.6 | FENDRR | 1.49897 | 1.47E-08 |
| ENSG00000268894.7 | PLCE1-AS1 | -1.233 | 9.80E-08 |
| ENSG00000269937.1 | RP11-20I23.8 | 0.65855 | 2.71E-13 |
| ENSG00000271605.6 | MILR1 | 0.716801 | 2.53E-08 |
| ENSG00000272695.2 | GAS6-DT | -0.90887 | 2.58E-10 |
| ENSG00000272711.1 | HK2-DT | 0.665218 | 0.007869 |
| ENSG00000273117.1 | INSIG1-DT | -0.98544 | 8.03E-05 |
| ENSG00000274070.2 | CASTOR2 | -0.68327 | 3.84E-35 |
| ENSG00000274641.2 | H2BC17 | 1.023949 | 9.25E-06 |
| ENSG00000275221.2 | H2AC15 | 0.720071 | 0.001557 |
| ENSG00000275993.3 | SIK1B | -1.1355 | 6.34E-15 |
| ENSG00000276107.1 | THBS1-IT1 | -0.90212 | 2.90E-09 |
| ENSG00000276368.2 | H2AC14 | 0.987981 | 4.14E-06 |
| ENSG00000276903.2 | H2AC16 | 1.118382 | 7.65E-06 |
| ENSG00000277304.1 | RP11-989E6.13 | -0.67148 | 0.000243 |
| ENSG00000277443.3 | MARCKS | -0.7472 | 1.02E-56 |

| | | | |
|---|---|---|---|
| ENSG00000277775.2 | H3C7 | 0.683132 | 0.000507 |
| ENSG00000278291.1 | RP11-172H24.4 | -0.79511 | 1.40E-17 |
| ENSG00000278588.2 | H2BC10 | 0.650326 | 0.000287 |
| ENSG00000278903.3 | CH507-145C22.1 | 0.957018 | 0.026347 |
| ENSG00000279118.1 | RP11-517I3.2 | -0.91798 | 4.77E-08 |
| ENSG00000279407.1 | AC007191.4 | -0.6742 | 0.002796 |
| ENSG00000282278.1 | RP11-231C18.3 | -1.08049 | 0.001176 |
| ENSG00000283154.2 | IQCJ-SCHIP1 | -0.62403 | 2.05E-06 |
| ENSG00000285006.1 | RP1-240K6.5 | 1.594114 | 0.019953 |
| ENSG00000285053.1 | TBCE | 0.644602 | 0.001164 |
| ENSG00000285106.2 | RP11-36B6.2 | 0.606771 | 0.005365 |
| ENSG00000285283.1 | RP1-65P5.6 | -0.98235 | 0.001057 |
| ENSG00000285517.1 | LINC00941 | 0.894131 | 5.70E-32 |
| ENSG00000286522.2 | H3C2 | 1.264154 | 9.15E-12 |
| ENSG00000287774.1 | RP11-69K20.1 | 1.861936 | 6.00E-05 |
| ENSG00000287839.1 | RP11-29H23.8 | 0.905464 | 2.57E-05 |
| ENSG00000288632.1 | RP11-345J4.11 | 0.615682 | 0.000875 |

**Table S2. Differentially expressed genes regulated by colchicine intervention compared to cholesterol loaded HASMCs.** HASMCs were cholesterol loaded (10μg/mL) for 72 hours, followed by treatment with colchicine (50nM) for 48 hours.

| ENSEMBL | name | LogFC_Reg_vs_Chol | padj_Reg_vs_Chol |
|---|---|---|---|
| ENSG00000006327.14 | TNFRSF12A | 1.494321 | ######## |
| ENSG00000006468.14 | ETV1 | -1.20268 | 5.10E-74 |
| ENSG00000007237.19 | GAS7 | -0.98467 | 3.51E-12 |
| ENSG00000008118.10 | CAMK1G | -1.45469 | 3.23E-31 |
| ENSG00000008517.19 | IL32 | 1.126674 | 1.32E-27 |
| ENSG00000010818.10 | HIVEP2 | 0.75789 | 1.03E-32 |
| ENSG00000011422.12 | PLAUR | 1.131205 | 5.66E-74 |
| ENSG00000011426.11 | ANLN | 0.714522 | 1.76E-08 |
| ENSG00000011465.18 | DCN | -0.58969 | 5.27E-32 |
| ENSG00000013375.16 | PGM3 | 0.666472 | 2.75E-30 |
| ENSG00000013588.9 | GPRC5A | 1.132195 | 1.08E-43 |
| ENSG00000017427.17 | IGF1 | 1.406263 | 1.32E-06 |
| ENSG00000018408.15 | WWTR1 | -0.8184 | 1.30E-94 |
| ENSG00000019991.18 | HGF | -0.70395 | 4.35E-27 |
| ENSG00000020577.14 | SAMD4A | 0.773579 | 2.28E-58 |
| ENSG00000022267.19 | FHL1 | 0.599613 | 4.88E-22 |
| ENSG00000023902.14 | PLEKHO1 | 0.832269 | 2.65E-45 |
| ENSG00000023909.10 | GCLM | 0.592657 | 6.08E-12 |
| ENSG00000025434.19 | NR1H3 | -1.23024 | 1.39E-43 |
| ENSG00000028137.19 | TNFRSF1B | -1.23631 | 3.23E-31 |
| ENSG00000034152.19 | MAP2K3 | 0.645756 | 1.49E-23 |
| ENSG00000038382.20 | TRIO | 0.613135 | 1.18E-38 |
| ENSG00000039560.14 | RAI14 | 1.034381 | 4.86E-96 |
| ENSG00000041982.16 | TNC | 2.685844 | 0 |
| ENSG00000042062.13 | RIPOR3 | -1.79566 | 8.08E-24 |
| ENSG00000044574.9 | HSPA5 | 0.821332 | 9.66E-45 |
| ENSG00000046604.14 | DSG2 | 0.887974 | 7.07E-40 |
| ENSG00000048052.23 | HDAC9 | 0.706195 | 6.50E-23 |
| ENSG00000048162.21 | NOP16 | 0.625832 | 0.000153 |
| ENSG00000049246.15 | PER3 | -0.67707 | 2.29E-19 |
| ENSG00000049449.10 | RCN1 | 0.739908 | 2.09E-28 |
| ENSG00000049759.20 | NEDD4L | 0.760118 | 6.85E-11 |
| ENSG00000050405.13 | LIMA1 | 0.737083 | 1.18E-58 |
| ENSG00000050730.16 | TNIP3 | 1.080513 | 2.11E-05 |
| ENSG00000051108.15 | HERPUD1 | 0.867599 | 1.59E-70 |
| ENSG00000051180.17 | RAD51 | 0.82623 | 0.001075 |
| ENSG00000058404.20 | CAMK2B | 0.970065 | 9.35E-10 |
| ENSG00000058668.15 | ATP2B4 | 0.714402 | 5.20E-35 |
| ENSG00000059728.11 | MXD1 | 0.639888 | 2.47E-18 |
| ENSG00000063180.9 | CA11 | -0.63296 | 7.78E-05 |
| ENSG00000064205.11 | CCN5 | -1.21072 | 0.002743 |
| ENSG00000064651.14 | SLC12A2 | -0.71367 | 4.00E-49 |
| ENSG00000064652.11 | SNX24 | 0.706081 | 2.33E-10 |
| ENSG00000064687.13 | ABCA7 | -0.79649 | 2.47E-17 |
| ENSG00000064763.11 | FAR2 | 0.693057 | 1.29E-08 |
| ENSG00000064989.13 | CALCRL | -0.85251 | 1.71E-22 |
| ENSG00000065802.12 | ASB1 | 0.969138 | 3.77E-60 |
| ENSG00000067057.18 | PFKP | 0.971664 | 1.34E-72 |
| ENSG00000067082.15 | KLF6 | 0.889318 | 1.19E-89 |
| ENSG00000067113.17 | PLPP1 | 0.660861 | 1.53E-22 |
| ENSG00000067715.14 | SYT1 | -0.89618 | 9.42E-49 |
| ENSG00000068366.21 | ACSL4 | 0.744814 | 2.31E-37 |
| ENSG00000068650.19 | ATP11A | 0.657433 | 1.02E-37 |
| ENSG00000068971.14 | PPP2R5B | 0.621884 | 8.00E-17 |
| ENSG00000069399.15 | BCL3 | -0.70097 | 2.89E-19 |
| ENSG00000069702.11 | TGFBR3 | -0.64746 | 8.08E-30 |

| | | | |
|---|---|---|---|
| ENSG00000070081.17 | NUCB2 | 0.611903 | 4.08E-18 |
| ENSG00000070404.10 | FSTL3 | 1.184395 | 6.58E-55 |
| ENSG00000070495.15 | JMJD6 | 0.710153 | 1.91E-16 |
| ENSG00000070669.17 | ASNS | 0.67475 | 7.30E-28 |
| ENSG00000070961.16 | ATP2B1 | 0.66411 | 1.13E-05 |
| ENSG00000071127.17 | WDR1 | 0.768617 | 1.98E-81 |
| ENSG00000071282.12 | LMCD1 | 1.338217 | 6.87E-35 |
| ENSG00000072110.16 | ACTN1 | 0.812997 | 9.85E-69 |
| ENSG00000072163.20 | LIMS2 | 1.205873 | 2.74E-21 |
| ENSG00000072952.20 | IRAG1 | 1.247787 | 9.66E-17 |
| ENSG00000072954.7 | TMEM38A | -0.86568 | 1.60E-06 |
| ENSG00000073008.15 | PVR | 1.294457 | ######## |
| ENSG00000073605.19 | GSDMB | 0.600904 | 0.00019 |
| ENSG00000073712.15 | FERMT2 | 0.599963 | 2.57E-38 |
| ENSG00000073756.13 | PTGS2 | 1.713019 | 2.68E-13 |
| ENSG00000074416.15 | MGLL | 1.215177 | 9.39E-83 |
| ENSG00000074527.13 | NTN4 | 0.652708 | 0.000419 |
| ENSG00000074590.14 | NUAK1 | 0.664007 | 7.59E-25 |
| ENSG00000074800.16 | ENO1 | 0.614408 | 3.43E-48 |
| ENSG00000075426.12 | FOSL2 | 0.647801 | 8.20E-37 |
| ENSG00000075624.17 | ACTB | 0.773356 | 7.16E-45 |
| ENSG00000075711.21 | DLG1 | 0.658286 | 2.48E-35 |
| ENSG00000076356.7 | PLXNA2 | -0.66371 | 1.66E-21 |
| ENSG00000076706.17 | MCAM | 0.982963 | 1.01E-32 |
| ENSG00000077942.19 | FBLN1 | -0.71217 | 9.55E-56 |
| ENSG00000078401.7 | EDN1 | -0.78862 | 1.88E-16 |
| ENSG00000079931.15 | MOXD1 | 0.641154 | 4.71E-21 |
| ENSG00000080573.7 | COL5A3 | 1.266966 | ######## |
| ENSG00000081051.8 | AFP | 1.776257 | 0.004621 |
| ENSG00000081059.20 | TCF7 | -0.75186 | 0.000134 |
| ENSG00000081181.8 | ARG2 | 0.846178 | 2.84E-06 |
| ENSG00000081189.16 | MEF2C | -0.61595 | 9.96E-16 |
| ENSG00000081320.11 | STK17B | -0.60404 | 1.00E-21 |
| ENSG00000081923.15 | ATP8B1 | 1.528308 | ######## |
| ENSG00000082126.18 | MPP4 | 1.458712 | 4.51E-08 |
| ENSG00000082482.14 | KCNK2 | -0.87418 | 1.93E-38 |
| ENSG00000082512.15 | TRAF5 | 0.586785 | 2.52E-12 |
| ENSG00000087116.16 | ADAMTS2 | 0.738393 | 2.37E-35 |
| ENSG00000088305.18 | DNMT3B | 0.660404 | 0.00017 |
| ENSG00000088451.11 | TGDS | 0.594493 | 5.24E-05 |
| ENSG00000090447.12 | TFAP4 | -0.60321 | 3.56E-11 |
| ENSG00000090520.12 | DNAJB11 | 0.756371 | 4.52E-16 |
| ENSG00000090530.10 | P3H2 | 0.644042 | 1.23E-12 |
| ENSG00000090539.15 | CHRD | 0.626639 | 0.000756 |
| ENSG00000092068.20 | SLC7A8 | -0.97921 | 4.46E-28 |
| ENSG00000092758.18 | COL9A3 | -0.65632 | 0.021089 |
| ENSG00000092841.19 | MYL6 | 0.833604 | 1.38E-09 |
| ENSG00000094804.12 | CDC6 | 0.697445 | 0.001594 |
| ENSG00000095303.17 | PTGS1 | 0.857737 | 2.43E-22 |
| ENSG00000095380.11 | NANS | 0.608666 | 8.50E-16 |
| ENSG00000095752.7 | IL11 | 0.920054 | 1.05E-07 |
| ENSG00000099337.5 | KCNK6 | 1.185321 | 9.28E-41 |
| ENSG00000099849.15 | RASSF7 | 0.786651 | 0.000293 |
| ENSG00000099860.9 | GADD45B | 0.739286 | 1.18E-16 |
| ENSG00000099953.10 | MMP11 | -0.70988 | 7.45E-12 |
| ENSG00000099957.16 | P2RX6 | -1.09319 | 2.08E-06 |
| ENSG00000100027.17 | YPEL1 | -0.71979 | 0.000306 |
| ENSG00000100092.24 | SH3BP1 | -0.93173 | 1.44E-41 |
| ENSG00000100139.14 | MICALL1 | 0.771485 | 1.78E-50 |
| ENSG00000100196.11 | KDELR3 | 0.822692 | 1.51E-58 |
| ENSG00000100234.12 | TIMP3 | -0.7894 | 1.09E-42 |
| ENSG00000100342.21 | APOL1 | -0.72844 | 0.000324 |
| ENSG00000100345.22 | MYH9 | 1.24657 | ######## |

| | | | |
|---|---|---|---|
| ENSG00000100596.7 | SPTLC2 | 0.595541 | 3.34E-33 |
| ENSG00000100767.17 | PAPLN | -0.64964 | 0.000183 |
| ENSG00000100889.12 | PCK2 | 0.69396 | 1.83E-21 |
| ENSG00000100934.15 | SEC23A | 0.598686 | 7.10E-37 |
| ENSG00000100979.15 | PLTP | -0.69017 | 7.38E-31 |
| ENSG00000101057.16 | MYBL2 | 0.875185 | 0.000169 |
| ENSG00000101098.13 | RIMS4 | -2.09424 | 0.000104 |
| ENSG00000101224.18 | CDC25B | -0.74638 | 1.80E-56 |
| ENSG00000101310.17 | SEC23B | 0.606831 | 6.76E-18 |
| ENSG00000101335.10 | MYL9 | 1.141636 | 5.18E-82 |
| ENSG00000101608.13 | MYL12A | 0.752413 | 1.06E-55 |
| ENSG00000101680.15 | LAMA1 | 0.803195 | 2.62E-19 |
| ENSG00000101928.13 | MOSPD1 | 0.708763 | 4.51E-29 |
| ENSG00000101955.15 | SRPX | -0.78902 | 3.12E-63 |
| ENSG00000102057.10 | KCND1 | -0.97515 | 1.74E-14 |
| ENSG00000102096.9 | PIM2 | -0.62447 | 5.76E-05 |
| ENSG00000102174.10 | PHEX | -1.141 | 4.03E-09 |
| ENSG00000102271.14 | KLHL4 | -0.76092 | 4.33E-09 |
| ENSG00000102359.9 | SRPX2 | 0.609251 | 8.71E-34 |
| ENSG00000102760.13 | RGCC | 1.014437 | 5.75E-11 |
| ENSG00000102802.10 | MEDAG | 0.884752 | 1.93E-66 |
| ENSG00000103187.8 | COTL1 | 0.713586 | 2.74E-47 |
| ENSG00000103241.7 | FOXF1 | 0.596421 | 1.64E-11 |
| ENSG00000103257.9 | SLC7A5 | 0.786196 | 1.28E-43 |
| ENSG00000103485.19 | QPRT | -0.97566 | 2.53E-23 |
| ENSG00000103742.12 | IGDCC4 | -0.70057 | 1.59E-18 |
| ENSG00000103888.17 | CEMIP | 0.896412 | ######## |
| ENSG00000104177.18 | MYEF2 | 0.773677 | 1.14E-17 |
| ENSG00000104356.11 | POP1 | 0.635573 | 3.62E-07 |
| ENSG00000104368.19 | PLAT | -0.83017 | 1.66E-58 |
| ENSG00000104812.15 | GYS1 | 0.770567 | 1.55E-40 |
| ENSG00000104881.16 | PPP1R13L | 1.001352 | 5.26E-32 |
| ENSG00000104951.16 | IL4I1 | -0.92871 | 0.000106 |
| ENSG00000105088.9 | OLFM2 | 1.067306 | 1.90E-25 |
| ENSG00000105516.11 | DBP | -1.02748 | 4.11E-13 |
| ENSG00000105755.8 | ETHE1 | 0.65804 | 1.17E-14 |
| ENSG00000105928.16 | GSDME | 0.621484 | 5.15E-28 |
| ENSG00000105996.7 | HOXA2 | -1.2506 | 2.86E-12 |
| ENSG00000106034.18 | CPED1 | 0.590865 | 1.38E-21 |
| ENSG00000106080.11 | FKBP14 | 0.739497 | 4.77E-30 |
| ENSG00000106278.12 | PTPRZ1 | 0.788397 | 6.72E-12 |
| ENSG00000106351.13 | AGFG2 | -0.62957 | 3.78E-09 |
| ENSG00000106366.9 | SERPINE1 | 2.325334 | 2.37E-79 |
| ENSG00000106511.6 | MEOX2 | -1.42584 | 1.63E-09 |
| ENSG00000106617.15 | PRKAG2 | 0.856609 | 9.69E-26 |
| ENSG00000106688.12 | SLC1A1 | 0.800326 | 1.31E-12 |
| ENSG00000106799.13 | TGFBR1 | 1.700027 | ######## |
| ENSG00000106976.21 | DNM1 | -0.65364 | 5.46E-25 |
| ENSG00000107317.13 | PTGDS | -0.63651 | 1.76E-05 |
| ENSG00000107562.16 | CXCL12 | -0.83095 | 6.62E-27 |
| ENSG00000107796.13 | ACTA2 | 1.44695 | ######## |
| ENSG00000107821.15 | KAZALD1 | -0.6825 | 6.14E-09 |
| ENSG00000107960.11 | STN1 | -0.69397 | 3.79E-14 |
| ENSG00000108576.10 | SLC6A4 | -1.41269 | 2.47E-10 |
| ENSG00000108688.11 | CCL7 | -1.52212 | 0.000427 |
| ENSG00000108821.14 | COL1A1 | 0.673472 | 2.86E-32 |
| ENSG00000108829.10 | LRRC59 | 0.777222 | 1.43E-57 |
| ENSG00000108854.16 | SMURF2 | 1.371057 | ######## |
| ENSG00000108947.5 | EFNB3 | -0.59014 | 9.67E-11 |
| ENSG00000108950.12 | FAM20A | -2.43423 | 7.83E-23 |
| ENSG00000108984.15 | MAP2K6 | -1.44566 | 1.50E-42 |
| ENSG00000109089.7 | CDR2L | 0.743429 | 9.32E-29 |
| ENSG00000109339.24 | MAPK10 | -0.66613 | 8.31E-05 |

| | | | |
|---|---|---|---|
| ENSG00000109667.12 | SLC2A9 | -0.65532 | 0.038354 |
| ENSG00000109705.8 | NKX3-2 | -0.81694 | 0.021799 |
| ENSG00000109743.11 | BST1 | 0.765647 | 1.73E-18 |
| ENSG00000109814.12 | UGDH | 0.68231 | 3.00E-39 |
| ENSG00000110031.13 | LPXN | 0.813898 | 3.59E-17 |
| ENSG00000110203.9 | FOLR3 | 1.232482 | 0.019021 |
| ENSG00000110852.5 | CLEC2B | -0.98591 | 4.60E-09 |
| ENSG00000110876.10 | SELPLG | 0.860943 | 6.62E-13 |
| ENSG00000110880.11 | CORO1C | 0.763226 | 4.07E-63 |
| ENSG00000110881.12 | ASIC1 | -0.77657 | 1.38E-28 |
| ENSG00000111057.11 | KRT18 | 0.59002 | 5.63E-19 |
| ENSG00000111077.18 | TNS2 | -0.59473 | 1.02E-21 |
| ENSG00000111087.10 | GLI1 | -1.10164 | 4.68E-12 |
| ENSG00000111186.13 | WNT5B | 0.751722 | 5.13E-05 |
| ENSG00000111266.9 | DUSP16 | 0.990238 | 2.90E-51 |
| ENSG00000111452.13 | ADGRD1 | 0.776295 | 0.000231 |
| ENSG00000111711.10 | GOLT1B | 0.634477 | 1.42E-21 |
| ENSG00000111799.22 | COL12A1 | 0.871488 | 3.41E-88 |
| ENSG00000111859.17 | NEDD9 | 2.477722 | ######## |
| ENSG00000111981.5 | ULBP1 | 0.76841 | 3.46E-06 |
| ENSG00000112208.11 | BAG2 | 0.692288 | 1.00E-15 |
| ENSG00000112210.12 | RAB23 | 1.161177 | 2.08E-64 |
| ENSG00000112419.14 | PHACTR2 | 0.842243 | 1.42E-70 |
| ENSG00000112658.8 | SRF | 0.588412 | 1.48E-24 |
| ENSG00000112773.16 | TENT5A | -0.65507 | 6.00E-26 |
| ENSG00000112893.10 | MAN2A1 | 0.648171 | 1.12E-34 |
| ENSG00000112936.19 | C7 | -1.25086 | 7.47E-38 |
| ENSG00000113070.8 | HBEGF | 2.841591 | ######## |
| ENSG00000113083.15 | LOX | 1.295854 | ######## |
| ENSG00000113140.11 | SPARC | 0.773365 | 8.74E-40 |
| ENSG00000113578.18 | FGF1 | 1.787844 | 1.16E-08 |
| ENSG00000113722.17 | CDX1 | -0.92608 | 0.011921 |
| ENSG00000113739.11 | STC2 | 1.333721 | ######## |
| ENSG00000114115.10 | RBP1 | -0.71449 | 8.11E-12 |
| ENSG00000114251.15 | WNT5A | 1.170019 | 2.98E-67 |
| ENSG00000114529.13 | C3orf52 | 0.593246 | 0.000206 |
| ENSG00000114850.7 | SSR3 | 0.778302 | 8.89E-62 |
| ENSG00000115009.13 | CCL20 | 0.830228 | 0.012448 |
| ENSG00000115091.12 | ACTR3 | 0.720732 | 1.35E-55 |
| ENSG00000115232.14 | ITGA4 | 0.765524 | 4.21E-37 |
| ENSG00000115290.10 | GRB14 | -0.86158 | 0.020253 |
| ENSG00000115380.20 | EFEMP1 | 0.625074 | 4.65E-48 |
| ENSG00000115461.5 | IGFBP5 | 2.509723 | 0 |
| ENSG00000115504.15 | EHBP1 | -0.58637 | 2.16E-14 |
| ENSG00000115525.18 | ST3GAL5 | -1.20436 | 1.11E-94 |
| ENSG00000115641.19 | FHL2 | 0.645759 | 4.39E-28 |
| ENSG00000115758.13 | ODC1 | 0.88206 | 2.30E-55 |
| ENSG00000115896.16 | PLCL1 | -0.92974 | 1.06E-25 |
| ENSG00000115902.11 | SLC1A4 | 0.967374 | 3.46E-47 |
| ENSG00000115935.18 | WIPF1 | -0.59627 | 6.19E-29 |
| ENSG00000115963.14 | RND3 | 0.631509 | 1.99E-17 |
| ENSG00000116191.18 | RALGPS2 | 0.589 | 2.02E-29 |
| ENSG00000116741.8 | RGS2 | -1.23604 | 1.23E-52 |
| ENSG00000116761.12 | CTH | 0.625547 | 4.01E-06 |
| ENSG00000116991.11 | SIPA1L2 | -1.74186 | 1.60E-54 |
| ENSG00000117143.13 | UAP1 | 1.375744 | ######## |
| ENSG00000117152.14 | RGS4 | 1.079059 | 9.54E-79 |
| ENSG00000117228.11 | GBP1 | 0.636523 | 2.99E-16 |
| ENSG00000117318.9 | ID3 | -0.68743 | 2.49E-11 |
| ENSG00000117394.24 | SLC2A1 | 0.826501 | 4.72E-45 |
| ENSG00000117395.13 | EBNA1BP2 | 0.738923 | 4.81E-21 |
| ENSG00000117479.15 | SLC19A2 | 0.71736 | 8.15E-11 |
| ENSG00000117525.14 | F3 | 2.042509 | ######## |

| | | | |
|---|---|---|---|
| ENSG00000117594.10 | HSD11B1 | -1.01544 | 1.61E-05 |
| ENSG00000117758.14 | STX12 | 0.740613 | 1.10E-42 |
| ENSG00000118263.15 | KLF7 | 0.851421 | 1.64E-57 |
| ENSG00000118508.5 | RAB32 | 0.856439 | 5.47E-44 |
| ENSG00000118523.6 | CCN2 | 1.370036 | ######## |
| ENSG00000118526.7 | TCF21 | 0.946796 | 1.49E-27 |
| ENSG00000118596.12 | SLC16A7 | 0.652912 | 2.57E-21 |
| ENSG00000118804.9 | STBD1 | 1.031568 | 3.89E-21 |
| ENSG00000118898.16 | PPL | -1.72642 | 1.20E-54 |
| ENSG00000118985.16 | ELL2 | 0.819898 | 7.34E-64 |
| ENSG00000119812.19 | FAM98A | 0.64719 | 7.38E-31 |
| ENSG00000119922.11 | IFIT2 | -0.66385 | 1.68E-10 |
| ENSG00000120129.6 | DUSP1 | 1.816787 | ######## |
| ENSG00000120217.14 | CD274 | 1.759148 | 1.64E-49 |
| ENSG00000120594.17 | PLXDC2 | -1.04766 | 4.11E-51 |
| ENSG00000120738.8 | EGR1 | -0.69467 | 2.99E-25 |
| ENSG00000120832.10 | MTERF2 | -0.58845 | 0.002001 |
| ENSG00000120875.9 | DUSP4 | -0.71308 | 1.70E-27 |
| ENSG00000120915.14 | EPHX2 | -0.99693 | 0.001657 |
| ENSG00000121440.15 | PDZRN3 | -0.60759 | 9.32E-29 |
| ENSG00000121578.13 | B4GALT4 | 0.624276 | 1.01E-16 |
| ENSG00000121858.11 | TNFSF10 | -0.82335 | 0.000512 |
| ENSG00000121957.15 | GPSM2 | -0.70655 | 9.98E-18 |
| ENSG00000122420.10 | PTGFR | -0.92263 | 2.25E-13 |
| ENSG00000122547.11 | EEPD1 | -0.97344 | 1.61E-06 |
| ENSG00000122641.11 | INHBA | 1.428249 | ######## |
| ENSG00000122861.16 | PLAU | 0.657598 | 1.03E-18 |
| ENSG00000122862.5 | SRGN | 0.801234 | 1.08E-30 |
| ENSG00000122877.17 | EGR2 | -1.94524 | 1.74E-16 |
| ENSG00000122966.17 | CIT | -0.84647 | 2.69E-08 |
| ENSG00000122986.14 | HVCN1 | -0.62044 | 0.020879 |
| ENSG00000123213.23 | NLN | 0.662423 | 6.92E-25 |
| ENSG00000123219.13 | CENPK | 0.987086 | 0.00044 |
| ENSG00000123689.6 | G0S2 | -1.30891 | 9.33E-55 |
| ENSG00000123977.10 | DAW1 | 1.854037 | 1.80E-14 |
| ENSG00000123983.15 | ACSL3 | 0.592189 | 1.73E-25 |
| ENSG00000124212.6 | PTGIS | -1.08505 | 4.21E-49 |
| ENSG00000124216.4 | SNAI1 | 0.592766 | 6.40E-08 |
| ENSG00000124225.16 | PMEPA1 | 0.660868 | 8.45E-08 |
| ENSG00000124249.7 | KCNK15 | -1.54084 | 0.002547 |
| ENSG00000124257.7 | NEURL2 | -0.63031 | 0.00271 |
| ENSG00000124593.16 | RP11-298J23.10 | -0.76681 | 6.73E-12 |
| ENSG00000124831.19 | LRRFIP1 | 1.142042 | 4.56E-83 |
| ENSG00000125378.16 | BMP4 | -0.92569 | 0.002082 |
| ENSG00000125538.12 | IL1B | 0.923085 | 0.000366 |
| ENSG00000125730.17 | C3 | -1.05345 | 7.09E-07 |
| ENSG00000125753.14 | VASP | 0.721037 | 1.16E-44 |
| ENSG00000125827.9 | TMX4 | -0.74009 | 5.44E-43 |
| ENSG00000125845.7 | BMP2 | 0.910214 | 9.26E-21 |
| ENSG00000125864.14 | BFSP1 | -0.79131 | 1.38E-06 |
| ENSG00000125968.9 | ID1 | -1.28223 | 8.06E-21 |
| ENSG00000126016.17 | AMOT | -0.63819 | 8.29E-18 |
| ENSG00000126522.17 | ASL | 0.588551 | 2.15E-13 |
| ENSG00000126562.17 | WNK4 | 0.829099 | 1.05E-36 |
| ENSG00000126790.12 | L3HYPDH | 0.609547 | 6.65E-10 |
| ENSG00000126860.12 | EVI2A | -0.60036 | 7.24E-06 |
| ENSG00000126861.5 | OMG | -1.41338 | 0.00044 |
| ENSG00000127329.16 | PTPRB | 0.650143 | 2.76E-14 |
| ENSG00000127561.15 | SYNGR3 | 0.739574 | 7.41E-06 |
| ENSG00000127585.12 | FBXL16 | 1.047928 | 4.92E-09 |
| ENSG00000127589.4 | TUBBP1 | -0.76172 | 0.021241 |
| ENSG00000127951.8 | FGL2 | -1.23013 | 1.35E-30 |
| ENSG00000128052.10 | KDR | -1.16153 | ######## |

| | | | |
|---|---|---|---|
| ENSG00000128228.5 | SDF2L1 | 1.124561 | 9.63E-22 |
| ENSG00000128284.19 | APOL3 | -0.79407 | 3.43E-06 |
| ENSG00000128294.16 | TPST2 | 0.591821 | 6.52E-15 |
| ENSG00000128342.5 | LIF | 1.3282 | 5.93E-13 |
| ENSG00000128482.16 | RNF112 | -1.24012 | 1.69E-17 |
| ENSG00000128512.23 | DOCK4 | -0.67043 | 1.37E-22 |
| ENSG00000128590.5 | DNAJB9 | 0.647827 | 5.77E-05 |
| ENSG00000128591.16 | FLNC | 0.755249 | 2.42E-33 |
| ENSG00000128594.8 | LRRC4 | -0.98082 | 3.38E-05 |
| ENSG00000128595.17 | CALU | 0.694943 | 8.72E-45 |
| ENSG00000128849.11 | CGNL1 | -0.96874 | 4.31E-47 |
| ENSG00000129116.19 | PALLD | 0.84176 | 1.10E-45 |
| ENSG00000129128.13 | SPCS3 | 0.611864 | 6.82E-32 |
| ENSG00000130176.8 | CNN1 | 2.074297 | ######## |
| ENSG00000130283.9 | GDF1 | -0.65872 | 0.000108 |
| ENSG00000130402.13 | ACTN4 | 0.882878 | 2.38E-56 |
| ENSG00000130513.6 | GDF15 | 0.968377 | 1.75E-53 |
| ENSG00000130592.17 | LSP1 | -0.83165 | 0.000259 |
| ENSG00000130600.19 | H19 | -0.96849 | 1.02E-44 |
| ENSG00000130635.16 | COL5A1 | 0.606311 | 2.38E-38 |
| ENSG00000130653.16 | PNPLA7 | -0.86771 | 3.38E-09 |
| ENSG00000131015.5 | ULBP2 | 1.247972 | 9.65E-18 |
| ENSG00000131018.25 | SYNE1 | 0.944437 | 6.76E-79 |
| ENSG00000131019.11 | ULBP3 | 0.86219 | 7.46E-09 |
| ENSG00000131477.11 | RAMP2 | -0.61155 | 0.001658 |
| ENSG00000131773.14 | KHDRBS3 | 0.98667 | 4.81E-29 |
| ENSG00000131871.15 | SELENOS | 0.791117 | 2.12E-29 |
| ENSG00000131979.20 | GCH1 | 0.608071 | 0.002179 |
| ENSG00000132122.12 | SPATA6 | -0.60377 | 3.71E-09 |
| ENSG00000132205.11 | EMILIN2 | 1.141416 | 5.93E-44 |
| ENSG00000132386.11 | SERPINF1 | -0.63125 | 6.35E-43 |
| ENSG00000132432.14 | SEC61G | 0.789187 | 4.86E-20 |
| ENSG00000132481.7 | TRIM47 | -0.63389 | 5.78E-16 |
| ENSG00000133026.13 | MYH10 | 1.006993 | ######## |
| ENSG00000133107.15 | TRPC4 | 1.740707 | 2.32E-72 |
| ENSG00000133321.11 | PLAAT4 | -1.15345 | 7.42E-13 |
| ENSG00000133657.16 | ATP13A3 | 0.88665 | 2.74E-71 |
| ENSG00000133805.15 | AMPD3 | 0.82065 | 0.000285 |
| ENSG00000133816.18 | MICAL2 | 1.114488 | ######## |
| ENSG00000133818.14 | RRAS2 | 0.740091 | 1.36E-38 |
| ENSG00000134013.16 | LOXL2 | 0.919721 | 7.42E-98 |
| ENSG00000134049.6 | IER3IP1 | 0.600507 | 4.52E-15 |
| ENSG00000134259.6 | NGF | 0.884201 | 1.46E-18 |
| ENSG00000134285.11 | FKBP11 | 0.63725 | 1.62E-14 |
| ENSG00000134333.14 | LDHA | 0.871288 | 2.96E-83 |
| ENSG00000134352.20 | IL6ST | -0.61259 | 1.09E-29 |
| ENSG00000134375.11 | TIMM17A | 0.612239 | 1.59E-19 |
| ENSG00000134569.10 | LRP4 | -1.01488 | ######## |
| ENSG00000134668.12 | SPOCD1 | 1.528564 | 1.44E-90 |
| ENSG00000134769.23 | DTNA | -0.70943 | 3.21E-07 |
| ENSG00000134853.12 | PDGFRA | -0.80793 | 2.62E-87 |
| ENSG00000134871.19 | COL4A2 | 0.762346 | 1.14E-72 |
| ENSG00000135269.18 | TES | 0.737488 | 6.56E-25 |
| ENSG00000135362.14 | PRR5L | 1.194454 | 4.82E-12 |
| ENSG00000135363.12 | LMO2 | -0.677 | 5.61E-06 |
| ENSG00000135424.18 | ITGA7 | 0.887849 | 3.35E-15 |
| ENSG00000135472.9 | FAIM2 | -1.15916 | 1.63E-63 |
| ENSG00000135547.9 | HEY2 | 1.233892 | 1.06E-10 |
| ENSG00000135549.15 | PKIB | 0.810735 | 4.91E-12 |
| ENSG00000135698.10 | MPHOSPH6 | 0.594292 | 7.63E-07 |
| ENSG00000135778.12 | NTPCR | -0.58819 | 3.58E-14 |
| ENSG00000135914.6 | HTR2B | -1.62794 | 7.31E-12 |
| ENSG00000136010.14 | ALDH1L2 | 0.628023 | 2.56E-34 |

| | | | |
|---|---|---|---|
| ENSG00000136026.14 | CKAP4 | 0.628326 | 1.74E-48 |
| ENSG00000136040.9 | PLXNC1 | -0.76909 | 1.58E-07 |
| ENSG00000136068.16 | FLNB | 0.807672 | 4.62E-57 |
| ENSG00000136240.10 | KDELR2 | 0.604942 | 2.56E-43 |
| ENSG00000136546.16 | SCN7A | -0.79642 | 4.84E-06 |
| ENSG00000136574.19 | GATA4 | -0.75693 | 4.76E-33 |
| ENSG00000136689.19 | IL1RN | -0.69982 | 0.007865 |
| ENSG00000136826.15 | KLF4 | -0.63848 | 6.44E-26 |
| ENSG00000136928.7 | GABBR2 | 0.840619 | 2.58E-12 |
| ENSG00000136960.13 | ENPP2 | -0.80657 | 2.69E-10 |
| ENSG00000136997.21 | MYC | 0.760565 | 2.39E-37 |
| ENSG00000137124.8 | ALDH1B1 | 0.655292 | 6.76E-18 |
| ENSG00000137266.15 | SLC22A23 | 0.667682 | 2.35E-16 |
| ENSG00000137331.12 | IER3 | 1.390716 | 4.37E-96 |
| ENSG00000137393.10 | RNF144B | 0.858487 | 1.49E-16 |
| ENSG00000137486.17 | ARRB1 | -0.6355 | 4.21E-20 |
| ENSG00000137501.17 | SYTL2 | 0.791567 | 2.70E-07 |
| ENSG00000137563.13 | GGH | 0.934455 | 1.54E-19 |
| ENSG00000137727.13 | ARHGAP20 | -0.78407 | 2.77E-14 |
| ENSG00000137752.24 | CASP1 | -0.85599 | 1.09E-06 |
| ENSG00000137801.11 | THBS1 | 0.720416 | 1.32E-54 |
| ENSG00000137807.16 | KIF23 | 0.918093 | 7.35E-11 |
| ENSG00000137872.17 | SEMA6D | -0.60434 | 0.001218 |
| ENSG00000137880.6 | GCHFR | -0.79944 | 0.0038 |
| ENSG00000137941.17 | TTLL7 | 0.712914 | 1.64E-15 |
| ENSG00000137959.17 | IFI44L | -0.96693 | 0.001189 |
| ENSG00000137965.11 | IFI44 | -0.92206 | 2.53E-05 |
| ENSG00000137970.7 | RPL7P9 | 1.314016 | 0.022113 |
| ENSG00000138028.16 | CGREF1 | 0.787854 | 0.008961 |
| ENSG00000138061.12 | CYP1B1 | 0.62431 | 9.77E-15 |
| ENSG00000138134.12 | STAMBPL1 | -0.67637 | 1.18E-08 |
| ENSG00000138316.11 | ADAMTS14 | 0.926438 | 1.33E-30 |
| ENSG00000138356.14 | AOX1 | 1.155377 | 1.91E-17 |
| ENSG00000138378.19 | STAT4 | 0.671454 | 0.004479 |
| ENSG00000138449.11 | SLC40A1 | -1.70123 | 6.77E-81 |
| ENSG00000138495.7 | COX17 | 0.646764 | 2.12E-05 |
| ENSG00000138675.17 | FGF5 | 1.241501 | ######## |
| ENSG00000138685.17 | FGF2 | 0.660904 | 1.66E-19 |
| ENSG00000138758.12 | SEPTIN11 | 0.618124 | 5.29E-36 |
| ENSG00000138771.16 | SHROOM3 | 1.575498 | 2.86E-51 |
| ENSG00000138835.22 | RGS3 | 0.939808 | 4.21E-63 |
| ENSG00000139269.3 | INHBE | 0.883783 | 0.000539 |
| ENSG00000139278.10 | GLIPR1 | 0.716792 | 4.20E-58 |
| ENSG00000139289.14 | PHLDA1 | -0.82064 | 4.17E-95 |
| ENSG00000139318.8 | DUSP6 | -0.61746 | 2.81E-10 |
| ENSG00000139329.5 | LUM | -0.59508 | 9.94E-29 |
| ENSG00000139364.11 | TMEM132B | 0.97451 | 3.20E-16 |
| ENSG00000139514.13 | SLC7A1 | 0.619555 | 8.47E-30 |
| ENSG00000139597.18 | N4BP2L1 | -0.93354 | 1.40E-09 |
| ENSG00000139629.16 | GALNT6 | 0.747604 | 6.58E-10 |
| ENSG00000139679.16 | LPAR6 | -0.76675 | 1.59E-05 |
| ENSG00000139734.19 | DIAPH3 | 1.530865 | 3.75E-58 |
| ENSG00000139874.6 | SSTR1 | 0.984022 | 8.18E-11 |
| ENSG00000140092.15 | FBLN5 | 0.60264 | 1.67E-32 |
| ENSG00000140416.23 | TPM1 | 1.206462 | ######## |
| ENSG00000140859.16 | KIFC3 | 0.608497 | 1.95E-22 |
| ENSG00000140876.11 | NUDT7 | -0.59117 | 0.015165 |
| ENSG00000141338.14 | ABCA8 | -0.73982 | 3.16E-56 |
| ENSG00000141448.11 | GATA6 | 0.601164 | 6.20E-06 |
| ENSG00000141469.18 | SLC14A1 | -2.15142 | ######## |
| ENSG00000141540.11 | TTYH2 | -0.86538 | 6.33E-20 |
| ENSG00000141682.12 | PMAIP1 | 0.89552 | 4.16E-17 |
| ENSG00000142552.8 | RCN3 | 0.684212 | 1.86E-24 |

| | | | |
|---|---|---|---|
| ENSG00000142871.18 | CCN1 | 1.19531 | ######## |
| ENSG00000142949.17 | PTPRF | 1.017602 | ######## |
| ENSG00000143127.13 | ITGA10 | -0.63616 | 6.51E-11 |
| ENSG00000143158.11 | MPC2 | 0.627379 | 2.57E-18 |
| ENSG00000143179.16 | UCK2 | 0.793129 | 4.94E-37 |
| ENSG00000143320.9 | CRABP2 | -0.58914 | 8.15E-14 |
| ENSG00000143355.16 | LHX9 | -0.70978 | 0.001617 |
| ENSG00000143367.16 | TUFT1 | 0.72038 | 5.26E-19 |
| ENSG00000143387.14 | CTSK | -0.6552 | 4.44E-39 |
| ENSG00000143416.21 | SELENBP1 | -1.26349 | 6.69E-25 |
| ENSG00000143443.10 | C1orf56 | -0.59589 | 0.002899 |
| ENSG00000143479.18 | DYRK3 | 0.654271 | 1.53E-22 |
| ENSG00000143494.16 | VASH2 | -0.96062 | 0.001356 |
| ENSG00000143867.7 | OSR1 | -1.06904 | 1.43E-20 |
| ENSG00000143878.10 | RHOB | 0.621687 | 8.75E-20 |
| ENSG00000144339.12 | TMEFF2 | -0.89391 | 1.65E-06 |
| ENSG00000144583.5 | MARCHF4 | 0.911175 | 5.75E-36 |
| ENSG00000144655.15 | CSRNP1 | 0.776683 | 9.91E-15 |
| ENSG00000144681.11 | STAC | -0.64829 | 0.001134 |
| ENSG00000144730.19 | IL17RD | -0.5955 | 3.83E-18 |
| ENSG00000144824.21 | PHLDB2 | 0.971758 | 7.11E-64 |
| ENSG00000144891.19 | AGTR1 | -0.88918 | 0.000376 |
| ENSG00000144959.11 | NCEH1 | 0.797994 | 3.62E-20 |
| ENSG00000145012.14 | LPP | 0.592237 | 2.43E-35 |
| ENSG00000145050.19 | MANF | 0.988797 | 2.28E-41 |
| ENSG00000145246.14 | ATP10D | 0.622132 | 1.89E-33 |
| ENSG00000145247.12 | OCIAD2 | 0.627224 | 0.000287 |
| ENSG00000145358.6 | DDIT4L | -1.27004 | 4.32E-27 |
| ENSG00000145506.14 | NKD2 | -1.40711 | 1.89E-08 |
| ENSG00000145526.12 | CDH18 | -0.7038 | 0.018752 |
| ENSG00000145632.15 | PLK2 | -0.80295 | 1.82E-35 |
| ENSG00000145675.15 | PIK3R1 | -0.61289 | 4.06E-27 |
| ENSG00000145779.8 | TNFAIP8 | -0.66117 | 1.16E-14 |
| ENSG00000145934.16 | TENM2 | 2.360425 | 4.00E-98 |
| ENSG00000146072.6 | TNFRSF21 | -0.79478 | 1.30E-60 |
| ENSG00000146267.12 | FAXC | 0.662161 | 0.001393 |
| ENSG00000146374.14 | RSPO3 | 1.316245 | 8.79E-25 |
| ENSG00000146376.11 | ARHGAP18 | -0.64361 | 1.63E-20 |
| ENSG00000146386.8 | ABRACL | 0.643371 | 1.18E-10 |
| ENSG00000146592.17 | CREB5 | 1.829489 | ######## |
| ENSG00000147027.4 | TMEM47 | 0.632816 | 6.81E-26 |
| ENSG00000147251.16 | DOCK11 | -0.65301 | 7.91E-32 |
| ENSG00000147408.14 | CSGALNACT1 | -0.74303 | 3.35E-22 |
| ENSG00000147852.17 | VLDLR | 0.747896 | 1.28E-43 |
| ENSG00000148154.10 | UGCG | 1.323533 | ######## |
| ENSG00000148344.11 | PTGES | -1.14556 | 3.89E-38 |
| ENSG00000148541.13 | FAM13C | -1.00406 | 9.04E-09 |
| ENSG00000148677.7 | ANKRD1 | 1.757916 | 4.93E-43 |
| ENSG00000148680.16 | HTR7 | -0.60611 | 0.005162 |
| ENSG00000148700.15 | ADD3 | -0.87242 | 2.92E-92 |
| ENSG00000148848.15 | ADAM12 | 0.878636 | 1.40E-72 |
| ENSG00000148926.10 | ADM | -0.69972 | 1.04E-33 |
| ENSG00000148935.11 | GAS2 | -0.87022 | 2.27E-07 |
| ENSG00000149054.16 | ZNF215 | 0.682637 | 0.001341 |
| ENSG00000149212.12 | SESN3 | -0.6394 | 3.47E-18 |
| ENSG00000149256.16 | TENM4 | -0.76728 | 1.98E-39 |
| ENSG00000149428.19 | HYOU1 | 0.781002 | 2.32E-39 |
| ENSG00000149571.12 | KIRREL3 | 0.718502 | 1.56E-06 |
| ENSG00000149591.17 | TAGLN | 1.950316 | 0 |
| ENSG00000149633.12 | KIAA1755 | -0.85658 | 3.30E-10 |
| ENSG00000150394.14 | CDH8 | 0.664931 | 0.009539 |
| ENSG00000150551.11 | LYPD1 | 0.785969 | 4.49E-16 |
| ENSG00000150636.17 | CCDC102B | -0.98733 | 2.23E-08 |

| | | | |
|---|---|---|---|
| ENSG00000150907.10 | FOXO1 | -0.92211 | 1.97E-31 |
| ENSG00000150938.10 | CRIM1 | 0.970701 | 2.47E-90 |
| ENSG00000151012.13 | SLC7A11 | 0.618491 | 8.23E-22 |
| ENSG00000151136.15 | BTBD11 | -1.15736 | 2.74E-38 |
| ENSG00000151150.22 | ANK3 | -0.80888 | 2.43E-32 |
| ENSG00000151276.23 | MAGI1 | 0.603631 | 1.04E-18 |
| ENSG00000151327.13 | FAM177A1 | 0.586381 | 4.26E-18 |
| ENSG00000151458.12 | ANKRD50 | 0.738695 | 9.11E-52 |
| ENSG00000151632.17 | AKR1C2 | -0.87162 | 1.91E-15 |
| ENSG00000151692.15 | RNF144A | -0.60542 | 3.79E-22 |
| ENSG00000151835.17 | SACS | 0.978773 | 3.23E-89 |
| ENSG00000152056.17 | AP1S3 | 1.351864 | 5.20E-35 |
| ENSG00000152137.8 | HSPB8 | 0.587055 | 6.13E-13 |
| ENSG00000152256.14 | PDK1 | 0.614278 | 1.01E-14 |
| ENSG00000152402.11 | GUCY1A2 | 0.637116 | 2.57E-19 |
| ENSG00000152518.8 | ZFP36L2 | -0.85279 | 3.24E-08 |
| ENSG00000152952.12 | PLOD2 | 0.975725 | 9.34E-53 |
| ENSG00000153162.9 | BMP6 | 0.646048 | 9.58E-14 |
| ENSG00000153234.15 | NR4A2 | -0.66951 | 0.001951 |
| ENSG00000153395.10 | LPCAT1 | 0.78276 | 6.44E-54 |
| ENSG00000153904.21 | DDAH1 | 1.100507 | ######## |
| ENSG00000153989.8 | NUS1 | 0.630698 | 1.38E-27 |
| ENSG00000154096.14 | THY1 | 0.684801 | 8.86E-33 |
| ENSG00000154127.10 | UBASH3B | 0.735285 | 1.30E-23 |
| ENSG00000154262.13 | ABCA6 | -0.6039 | 1.12E-21 |
| ENSG00000154263.17 | ABCA10 | -0.98216 | 1.44E-06 |
| ENSG00000154319.16 | FAM167A | 0.66642 | 1.95E-07 |
| ENSG00000154380.17 | ENAH | 0.902646 | ######## |
| ENSG00000154511.12 | DIPK1A | 0.884802 | 6.70E-12 |
| ENSG00000154545.17 | MAGED4 | 0.839663 | 0.002075 |
| ENSG00000154553.16 | PDLIM3 | 1.738171 | ######## |
| ENSG00000154556.18 | SORBS2 | 0.934317 | 7.89E-11 |
| ENSG00000154639.19 | CXADR | 1.027888 | 3.85E-13 |
| ENSG00000154640.15 | BTG3 | 0.60902 | 1.15E-12 |
| ENSG00000154678.18 | PDE1C | 0.972915 | 7.85E-10 |
| ENSG00000154721.15 | JAM2 | -0.94003 | 1.70E-35 |
| ENSG00000155755.19 | TMEM237 | 0.686835 | 1.33E-34 |
| ENSG00000155760.3 | FZD7 | -0.92607 | 2.92E-19 |
| ENSG00000156140.10 | ADAMTS3 | 0.912999 | 2.82E-24 |
| ENSG00000156265.16 | MAP3K7CL | 2.062712 | ######## |
| ENSG00000156384.15 | SFR1 | -0.67813 | 6.87E-06 |
| ENSG00000156463.18 | SH3RF2 | -1.00663 | 2.47E-21 |
| ENSG00000156521.14 | TYSND1 | -0.67493 | 2.36E-06 |
| ENSG00000157368.11 | IL34 | -0.63234 | 1.21E-07 |
| ENSG00000157404.16 | KIT | 1.269764 | 2.12E-29 |
| ENSG00000157456.8 | CCNB2 | -0.65484 | 0.031112 |
| ENSG00000157557.13 | ETS2 | -0.6563 | 5.73E-32 |
| ENSG00000157680.16 | DGKI | 1.237096 | 1.93E-46 |
| ENSG00000158023.10 | CFAP251 | 0.803962 | 2.07E-05 |
| ENSG00000158292.7 | GPR153 | -0.67119 | 1.71E-18 |
| ENSG00000158710.15 | TAGLN2 | 0.680841 | 3.73E-33 |
| ENSG00000158747.15 | NBL1 | -0.66546 | 3.35E-31 |
| ENSG00000158859.10 | ADAMTS4 | 0.92378 | 6.48E-34 |
| ENSG00000159176.14 | CSRP1 | 0.915705 | 3.76E-69 |
| ENSG00000159200.18 | RCAN1 | 1.385037 | 1.21E-89 |
| ENSG00000159261.12 | CLDN14 | 1.604226 | 2.29E-19 |
| ENSG00000159674.12 | SPON2 | 1.123882 | 1.72E-10 |
| ENSG00000159713.11 | TPPP3 | 0.742714 | 0.043959 |
| ENSG00000159761.15 | C16orf86 | -0.88775 | 0.007102 |
| ENSG00000159840.16 | ZYX | 0.895422 | 1.45E-60 |
| ENSG00000160870.15 | CYP3A7 | -0.954 | 1.00E-11 |
| ENSG00000161647.19 | MPP3 | 0.587985 | 0.002473 |
| ENSG00000161888.11 | SPC24 | -0.61336 | 0.004091 |

| | | | |
|---|---|---|---|
| ENSG00000162073.14 | PAQR4 | 0.638659 | 0.000243 |
| ENSG00000162407.9 | PLPP3 | -0.77019 | 2.04E-57 |
| ENSG00000162433.15 | AK4 | 0.725984 | 3.19E-33 |
| ENSG00000162490.7 | DRAXIN | -1.15396 | 3.72E-23 |
| ENSG00000162493.17 | PDPN | 1.136771 | 0.000165 |
| ENSG00000162496.9 | DHRS3 | -0.75717 | 3.24E-14 |
| ENSG00000162520.15 | SYNC | 0.828853 | 2.16E-49 |
| ENSG00000162595.7 | DIRAS3 | -1.00968 | 1.51E-13 |
| ENSG00000162599.17 | NFIA | -0.69077 | 1.02E-25 |
| ENSG00000162614.19 | NEXN | 1.094639 | 7.78E-48 |
| ENSG00000162616.9 | DNAJB4 | 0.987093 | 2.45E-44 |
| ENSG00000162631.20 | NTNG1 | -0.9082 | 1.67E-26 |
| ENSG00000162654.9 | GBP4 | -0.9476 | 2.08E-26 |
| ENSG00000162692.12 | VCAM1 | -1.87959 | 8.03E-29 |
| ENSG00000162702.8 | ZNF281 | 0.720347 | 1.18E-32 |
| ENSG00000162704.16 | ARPC5 | 0.656592 | 3.29E-29 |
| ENSG00000162772.17 | ATF3 | 1.617617 | 6.03E-46 |
| ENSG00000162804.14 | SNED1 | -0.59043 | 8.10E-21 |
| ENSG00000162944.11 | RFTN2 | -0.70031 | 3.79E-10 |
| ENSG00000162946.23 | DISC1 | -0.61004 | 2.95E-07 |
| ENSG00000163017.14 | ACTG2 | 3.015935 | 1.89E-10 |
| ENSG00000163050.18 | COQ8A | -0.60504 | 1.57E-11 |
| ENSG00000163110.15 | PDLIM5 | 0.718617 | 2.19E-45 |
| ENSG00000163378.14 | EOGT | 0.598056 | 7.74E-21 |
| ENSG00000163430.12 | FSTL1 | 0.628104 | 1.73E-47 |
| ENSG00000163431.13 | LMOD1 | 1.528623 | 2.61E-34 |
| ENSG00000163485.17 | ADORA1 | -0.64233 | 0.006026 |
| ENSG00000163545.11 | NUAK2 | 0.648243 | 3.18E-09 |
| ENSG00000163644.15 | PPM1K | 0.601565 | 3.76E-13 |
| ENSG00000163661.4 | PTX3 | 0.587251 | 1.00E-17 |
| ENSG00000163814.8 | CDCP1 | 0.750464 | 2.80E-28 |
| ENSG00000163827.14 | LRRC2 | 1.171917 | 3.25E-16 |
| ENSG00000163888.4 | CAMK2N2 | 0.60859 | 0.009322 |
| ENSG00000163947.12 | ARHGEF3 | -1.01713 | 8.32E-58 |
| ENSG00000163975.12 | MELTF | 0.970938 | 5.71E-17 |
| ENSG00000164035.10 | EMCN | -1.98694 | 1.76E-18 |
| ENSG00000164056.11 | SPRY1 | -0.85094 | 6.98E-50 |
| ENSG00000164093.17 | PITX2 | -0.70336 | 1.03E-16 |
| ENSG00000164099.3 | PRSS12 | 0.778431 | 1.02E-33 |
| ENSG00000164136.17 | IL15 | 0.67075 | 0.031799 |
| ENSG00000164161.10 | HHIP | -1.15841 | 4.64E-76 |
| ENSG00000164236.12 | ANKRD33B | -1.07073 | 7.29E-28 |
| ENSG00000164440.15 | TXLNB | -1.17845 | 4.35E-11 |
| ENSG00000164442.10 | CITED2 | 0.588636 | 6.19E-30 |
| ENSG00000164483.17 | SAMD3 | -0.6867 | 3.15E-07 |
| ENSG00000164741.15 | DLC1 | 0.679986 | 2.57E-57 |
| ENSG00000164761.9 | TNFRSF11B | 1.165618 | 2.36E-31 |
| ENSG00000164849.10 | GPR146 | -0.64211 | 0.0311 |
| ENSG00000164920.9 | OSR2 | -1.57719 | 1.75E-08 |
| ENSG00000165195.16 | PIGA | 0.762212 | 6.78E-08 |
| ENSG00000165424.7 | ZCCHC24 | -0.91232 | 2.37E-70 |
| ENSG00000165434.8 | PGM2L1 | 0.6063 | 1.69E-15 |
| ENSG00000165527.7 | ARF6 | 0.589723 | 6.09E-37 |
| ENSG00000165617.15 | DACT1 | 1.364122 | ######## |
| ENSG00000165795.24 | NDRG2 | -0.69316 | 1.55E-10 |
| ENSG00000165996.14 | HACD1 | 0.742527 | 7.57E-09 |
| ENSG00000166106.4 | ADAMTS15 | -0.61659 | 2.96E-15 |
| ENSG00000166123.14 | GPT2 | 0.684198 | 3.39E-17 |
| ENSG00000166396.13 | SERPINB7 | 1.185035 | 2.91E-08 |
| ENSG00000166401.15 | SERPINB8 | 0.643431 | 3.72E-12 |
| ENSG00000166483.12 | WEE1 | 0.614686 | 9.00E-26 |
| ENSG00000166562.9 | SEC11C | 1.245387 | 1.26E-21 |
| ENSG00000166582.10 | CENPV | 0.982665 | 6.06E-15 |

| | | | |
|---|---|---|---|
| ENSG00000166598.16 | HSP90B1 | 0.611153 | 2.02E-22 |
| ENSG00000166670.10 | MMP10 | -1.12211 | 2.00E-05 |
| ENSG00000166741.8 | NNMT | 0.587183 | 4.60E-31 |
| ENSG00000166780.11 | BMERB1 | -0.90727 | 1.72E-56 |
| ENSG00000166900.17 | STX3 | 0.654334 | 2.68E-22 |
| ENSG00000166920.13 | C15orf48 | 1.378043 | 2.02E-07 |
| ENSG00000166922.8 | SCG5 | 1.542928 | 2.82E-20 |
| ENSG00000166923.12 | GREM1 | 1.688809 | 0 |
| ENSG00000166949.17 | SMAD3 | -0.67479 | 4.25E-47 |
| ENSG00000167371.21 | PRRT2 | -1.01836 | 2.86E-16 |
| ENSG00000167460.17 | TPM4 | 0.88377 | ######## |
| ENSG00000167549.19 | CORO6 | -0.74334 | 1.28E-06 |
| ENSG00000167641.11 | PPP1R14A | 0.766755 | 0.009798 |
| ENSG00000167642.13 | SPINT2 | 0.742096 | 0.022503 |
| ENSG00000167645.17 | YIF1B | 0.766564 | 5.97E-23 |
| ENSG00000167657.14 | DAPK3 | 0.8401 | 4.24E-51 |
| ENSG00000167703.15 | SLC43A2 | -0.67037 | 1.05E-13 |
| ENSG00000167797.8 | CDK2AP2 | 0.940408 | 1.07E-40 |
| ENSG00000167972.14 | ABCA3 | 0.806429 | 1.60E-09 |
| ENSG00000167992.13 | VWCE | -0.97276 | 5.56E-09 |
| ENSG00000168374.11 | ARF4 | 0.6324 | 3.39E-36 |
| ENSG00000168386.19 | FILIP1L | 1.232339 | ######## |
| ENSG00000168389.18 | MFSD2A | 1.102757 | 4.46E-08 |
| ENSG00000168497.5 | CAVIN2 | -0.7311 | 1.27E-06 |
| ENSG00000168575.10 | SLC20A2 | 1.267991 | ######## |
| ENSG00000168621.15 | GDNF | 1.709868 | 2.73E-84 |
| ENSG00000168672.4 | LRATD2 | 0.624129 | 2.36E-06 |
| ENSG00000168916.16 | ZNF608 | -0.87377 | 1.79E-20 |
| ENSG00000168961.17 | LGALS9 | -0.70052 | 0.001846 |
| ENSG00000168994.14 | PXDC1 | 0.867626 | 3.96E-52 |
| ENSG00000169122.11 | FAM110B | -0.69372 | 5.39E-20 |
| ENSG00000169359.16 | SLC33A1 | 0.666349 | 3.33E-26 |
| ENSG00000169429.11 | CXCL8 | 1.043697 | 0.014513 |
| ENSG00000169439.12 | SDC2 | 0.689286 | 8.12E-31 |
| ENSG00000169744.13 | LDB2 | -0.69731 | 1.17E-29 |
| ENSG00000169756.16 | LIMS1 | 0.599357 | 2.71E-27 |
| ENSG00000169851.15 | PCDH7 | -0.70217 | 2.14E-28 |
| ENSG00000169857.9 | AVEN | 0.799337 | 1.89E-25 |
| ENSG00000170006.12 | TMEM154 | 0.598529 | 3.72E-13 |
| ENSG00000170017.12 | ALCAM | 0.617565 | 1.17E-31 |
| ENSG00000170153.11 | RNF150 | -0.70589 | 1.89E-14 |
| ENSG00000170271.11 | FAXDC2 | -0.6041 | 3.73E-12 |
| ENSG00000170323.9 | FABP4 | -2.07046 | 2.61E-06 |
| ENSG00000170485.17 | NPAS2 | 0.788275 | 1.17E-19 |
| ENSG00000170522.10 | ELOVL6 | 0.601384 | 5.69E-18 |
| ENSG00000170542.6 | SERPINB9 | -0.61733 | 5.54E-18 |
| ENSG00000170579.17 | DLGAP1 | -0.66307 | 0.000456 |
| ENSG00000170801.10 | HTRA3 | -0.73361 | 5.57E-16 |
| ENSG00000170962.13 | PDGFD | -0.72247 | 2.98E-33 |
| ENSG00000171067.11 | C11orf24 | 0.65299 | 1.94E-32 |
| ENSG00000171241.9 | SHCBP1 | 0.609291 | 0.022243 |
| ENSG00000171246.6 | NPTX1 | -0.87901 | 1.88E-14 |
| ENSG00000171316.12 | CHD7 | 0.65662 | 1.47E-14 |
| ENSG00000171388.12 | APLN | 0.669544 | 7.24E-12 |
| ENSG00000171522.6 | PTGER4 | -0.90514 | 1.92E-19 |
| ENSG00000171617.15 | ENC1 | 1.400833 | ######## |
| ENSG00000171724.3 | VAT1L | 0.643589 | 6.64E-41 |
| ENSG00000171793.16 | CTPS1 | 0.797483 | 9.16E-30 |
| ENSG00000171848.16 | RRM2 | 1.051147 | 4.66E-10 |
| ENSG00000171951.5 | SCG2 | 0.723641 | 8.90E-09 |
| ENSG00000172020.13 | GAP43 | -0.75853 | 0.001578 |
| ENSG00000172057.10 | ORMDL3 | 0.644546 | 9.23E-31 |
| ENSG00000172071.15 | EIF2AK3 | 0.619622 | 2.00E-22 |

| | | | |
|---|---|---|---|
| ENSG00000172115.9 | CYCS | 0.686953 | 4.63E-18 |
| ENSG00000172137.19 | CALB2 | 1.365078 | 1.54E-14 |
| ENSG00000172159.16 | FRMD3 | -1.69245 | 1.70E-44 |
| ENSG00000172260.15 | NEGR1 | 1.590282 | 6.00E-91 |
| ENSG00000172296.13 | SPTLC3 | -0.81626 | 7.62E-09 |
| ENSG00000172399.6 | MYOZ2 | 0.735313 | 1.09E-06 |
| ENSG00000172738.12 | TMEM217 | 1.274903 | 3.86E-17 |
| ENSG00000172803.18 | SNX32 | -0.77304 | 0.019297 |
| ENSG00000172965.17 | MIR4435-2HG | 0.618121 | 3.68E-27 |
| ENSG00000173210.20 | ABLIM3 | -0.83858 | 1.98E-42 |
| ENSG00000173221.14 | GLRX | 0.592554 | 1.12E-11 |
| ENSG00000173276.14 | ZBTB21 | 0.634843 | 1.65E-21 |
| ENSG00000173334.4 | TRIB1 | 1.332364 | 1.09E-51 |
| ENSG00000173531.15 | MST1 | -0.66953 | 3.31E-07 |
| ENSG00000173641.18 | HSPB7 | 0.903677 | 2.65E-31 |
| ENSG00000173706.14 | HEG1 | 0.669667 | 3.07E-66 |
| ENSG00000173905.9 | GOLIM4 | 0.681469 | 1.13E-23 |
| ENSG00000174059.17 | CD34 | -1.169 | 7.39E-09 |
| ENSG00000174099.12 | MSRB3 | 0.711515 | 3.94E-50 |
| ENSG00000174136.13 | RGMB | 0.741112 | 1.93E-38 |
| ENSG00000174437.18 | ATP2A2 | 0.588806 | 9.66E-49 |
| ENSG00000174804.4 | FZD4 | -0.76641 | 6.77E-46 |
| ENSG00000174851.16 | YIF1A | 0.59043 | 2.27E-19 |
| ENSG00000174939.11 | ASPHD1 | 0.736179 | 5.80E-06 |
| ENSG00000175274.19 | TP53I11 | 0.826756 | 1.14E-31 |
| ENSG00000175287.19 | PHYHD1 | -0.66044 | 2.91E-05 |
| ENSG00000175592.9 | FOSL1 | 0.789004 | 1.22E-24 |
| ENSG00000175745.14 | NR2F1 | -0.73295 | 8.00E-16 |
| ENSG00000175772.11 | LINC01106 | 0.82305 | 0.000294 |
| ENSG00000175874.10 | CREG2 | -0.89504 | 0.000147 |
| ENSG00000176020.9 | AMIGO3 | -1.09555 | 0.003275 |
| ENSG00000176125.7 | UFSP1 | -0.73809 | 0.021438 |
| ENSG00000176170.14 | SPHK1 | 0.922397 | 6.83E-43 |
| ENSG00000176171.11 | BNIP3 | 0.627338 | 8.40E-28 |
| ENSG00000176438.13 | SYNE3 | -1.34453 | ######## |
| ENSG00000176463.14 | SLCO3A1 | 0.602474 | 1.54E-18 |
| ENSG00000176641.11 | RNF152 | -0.72999 | 1.83E-38 |
| ENSG00000176692.8 | FOXC2 | 1.132522 | 1.07E-14 |
| ENSG00000176697.20 | BDNF | 1.802948 | ######## |
| ENSG00000176909.12 | MAMSTR | -0.6348 | 0.007173 |
| ENSG00000176971.4 | FIBIN | 1.012501 | 1.60E-45 |
| ENSG00000177425.11 | PAWR | 0.700018 | 4.37E-30 |
| ENSG00000178031.18 | ADAMTSL1 | -0.86798 | 1.04E-31 |
| ENSG00000178662.16 | CSRNP3 | -0.8535 | 1.40E-07 |
| ENSG00000178695.6 | KCTD12 | -0.60318 | 9.25E-32 |
| ENSG00000178726.7 | THBD | -1.19498 | 1.71E-68 |
| ENSG00000178773.15 | CPNE7 | 0.600471 | 0.000153 |
| ENSG00000178860.9 | MSC | 0.619704 | 2.81E-12 |
| ENSG00000178878.13 | APOLD1 | 0.825166 | 2.14E-15 |
| ENSG00000178922.18 | HYI | 0.933822 | 1.01E-35 |
| ENSG00000179218.15 | CALR | 0.725366 | 1.07E-69 |
| ENSG00000179388.9 | EGR3 | -1.75848 | 5.45E-13 |
| ENSG00000179476.8 | C14orf28 | 0.588244 | 1.83E-06 |
| ENSG00000179604.10 | CDC42EP4 | -0.63443 | 4.38E-22 |
| ENSG00000179776.19 | CDH5 | -0.78056 | 1.20E-39 |
| ENSG00000179820.16 | MYADM | 0.895064 | 2.64E-89 |
| ENSG00000180096.12 | SEPTIN1 | -0.69 | 0.024365 |
| ENSG00000180440.4 | SERTM1 | -1.29707 | 3.29E-08 |
| ENSG00000180537.13 | RNF182 | 0.754768 | 0.00032 |
| ENSG00000180769.10 | WDFY3-AS2 | -0.78845 | 8.29E-08 |
| ENSG00000180801.14 | ARSJ | 1.524219 | ######## |
| ENSG00000180875.5 | GREM2 | -1.19497 | 2.08E-18 |
| ENSG00000180881.19 | CAPS2 | -0.6416 | 0.046886 |

| | | | |
|---|---|---|---|
| ENSG00000180914.11 | OXTR | 1.584247 | 9.28E-55 |
| ENSG00000181444.13 | ZNF467 | -0.78896 | 4.35E-12 |
| ENSG00000181649.8 | PHLDA2 | 0.831622 | 0.002703 |
| ENSG00000181804.15 | SLC9A9 | -1.38842 | 1.31E-41 |
| ENSG00000182054.10 | IDH2 | 0.694643 | 4.99E-29 |
| ENSG00000182175.14 | RGMA | -1.02561 | 7.46E-16 |
| ENSG00000182621.18 | PLCB1 | -0.71219 | 4.46E-25 |
| ENSG00000182752.10 | PAPPA | 1.2029 | ######## |
| ENSG00000183010.17 | PYCR1 | 0.770661 | 5.74E-41 |
| ENSG00000183087.15 | GAS6 | 0.703216 | 2.79E-54 |
| ENSG00000183098.11 | GPC6 | -0.59085 | 4.54E-29 |
| ENSG00000183287.14 | CCBE1 | -0.86903 | 4.87E-29 |
| ENSG00000183696.14 | UPP1 | 0.704 | 6.65E-07 |
| ENSG00000183715.14 | OPCML | 0.703475 | 8.49E-08 |
| ENSG00000183876.9 | ARSI | 0.744676 | 1.86E-06 |
| ENSG00000184164.15 | CRELD2 | 0.762594 | 2.32E-17 |
| ENSG00000184226.15 | PCDH9 | -0.62635 | 5.22E-18 |
| ENSG00000184254.17 | ALDH1A3 | 0.69329 | 4.07E-33 |
| ENSG00000184545.11 | DUSP8 | 1.913592 | 3.56E-67 |
| ENSG00000184916.9 | JAG2 | -0.60909 | 0.000138 |
| ENSG00000184988.8 | TMEM106A | 0.915184 | 2.56E-12 |
| ENSG00000185015.8 | CA13 | 0.871373 | 7.76E-10 |
| ENSG00000185022.12 | MAFF | 0.635438 | 9.74E-16 |
| ENSG00000185215.11 | TNFAIP2 | -0.8605 | 5.71E-09 |
| ENSG00000185345.23 | PRKN | -0.81101 | 0.023513 |
| ENSG00000185432.12 | METTL7A | -1.60385 | 1.15E-33 |
| ENSG00000185499.16 | MUC1 | 0.623254 | 0.000338 |
| ENSG00000185614.7 | INKA1 | -0.67188 | 4.65E-08 |
| ENSG00000185634.12 | SHC4 | -1.17079 | 1.04E-16 |
| ENSG00000185745.10 | IFIT1 | -0.87164 | 3.45E-09 |
| ENSG00000185862.7 | EVI2B | -1.46708 | 2.02E-06 |
| ENSG00000185885.17 | IFITM1 | -0.70517 | 3.85E-15 |
| ENSG00000185920.17 | PTCH1 | -0.77759 | 1.93E-17 |
| ENSG00000186340.17 | THBS2 | 0.874549 | 1.60E-94 |
| ENSG00000186352.9 | ANKRD37 | 0.746267 | 0.004929 |
| ENSG00000186567.13 | CEACAM19 | 0.752786 | 7.32E-05 |
| ENSG00000186575.19 | NF2 | 0.791531 | 9.24E-72 |
| ENSG00000186765.12 | FSCN2 | 0.663921 | 0.036244 |
| ENSG00000187134.14 | AKR1C1 | -0.7574 | 7.22E-20 |
| ENSG00000187164.20 | SHTN1 | 0.743026 | 2.90E-11 |
| ENSG00000187479.8 | C11orf96 | 1.453068 | 4.77E-12 |
| ENSG00000187498.16 | COL4A1 | 1.057038 | ######## |
| ENSG00000187720.14 | THSD4 | 1.578137 | ######## |
| ENSG00000187741.15 | FANCA | 0.611102 | 0.012167 |
| ENSG00000187955.12 | COL14A1 | -0.94645 | 1.51E-66 |
| ENSG00000188042.8 | ARL4C | 0.740494 | 5.94E-51 |
| ENSG00000188112.9 | C6orf132 | 1.070181 | 4.63E-36 |
| ENSG00000188223.9 | AC002398.9 | -3.09333 | 8.79E-07 |
| ENSG00000188266.14 | HYKK | -0.58983 | 0.019425 |
| ENSG00000188290.11 | HES4 | 1.793056 | 8.12E-06 |
| ENSG00000188312.14 | CENPP | -0.93193 | 1.23E-20 |
| ENSG00000188522.15 | FAM83G | 0.633908 | 1.36E-23 |
| ENSG00000188641.14 | DPYD | -0.69543 | 1.29E-41 |
| ENSG00000189184.12 | PCDH18 | -0.82994 | 3.53E-47 |
| ENSG00000189320.9 | FAM180A | -0.74483 | 0.001334 |
| ENSG00000189410.12 | SH2D5 | 1.607929 | 8.22E-52 |
| ENSG00000196139.14 | AKR1C3 | -0.86013 | 1.51E-06 |
| ENSG00000196352.16 | CD55 | 0.684837 | 5.34E-21 |
| ENSG00000196616.14 | ADH1B | -2.4158 | 1.10E-31 |
| ENSG00000196628.20 | TCF4 | -0.82243 | 5.66E-68 |
| ENSG00000196782.12 | MAML3 | -0.91206 | 1.94E-39 |
| ENSG00000196839.13 | ADA | -0.7416 | 3.72E-11 |
| ENSG00000196843.17 | ARID5A | 1.187385 | 5.24E-36 |

| | | | |
|---|---|---|---|
| ENSG00000196878.15 | LAMB3 | -0.70603 | 2.27E-10 |
| ENSG00000196923.14 | PDLIM7 | 1.045607 | 2.55E-90 |
| ENSG00000196924.19 | FLNA | 0.779695 | 1.43E-52 |
| ENSG00000197301.7 | HMGA2-AS1 | -0.79137 | 0.006901 |
| ENSG00000197321.15 | SVIL | -0.63578 | 2.71E-48 |
| ENSG00000197381.17 | ADARB1 | 0.622862 | 3.83E-25 |
| ENSG00000197558.13 | SSPOP | -0.88139 | 3.07E-16 |
| ENSG00000197632.9 | SERPINB2 | 2.053491 | 8.62E-23 |
| ENSG00000197646.8 | PDCD1LG2 | 2.177804 | ######## |
| ENSG00000197785.14 | ATAD3A | 0.728119 | 6.80E-24 |
| ENSG00000197860.10 | SGTB | 0.853305 | 2.64E-40 |
| ENSG00000197977.4 | ELOVL2 | -0.80908 | 3.41E-15 |
| ENSG00000198018.7 | ENTPD7 | 0.727895 | 2.87E-22 |
| ENSG00000198121.14 | LPAR1 | -0.80226 | 2.55E-39 |
| ENSG00000198380.13 | GFPT1 | 0.642366 | 6.37E-28 |
| ENSG00000198431.16 | TXNRD1 | 0.691632 | 4.56E-52 |
| ENSG00000198467.16 | TPM2 | 0.717948 | 2.36E-05 |
| ENSG00000198483.13 | ANKRD35 | -0.64044 | 0.007263 |
| ENSG00000198517.10 | MAFK | 0.670517 | 5.65E-28 |
| ENSG00000198682.13 | PAPSS2 | 0.747354 | 9.13E-60 |
| ENSG00000198796.7 | ALPK2 | 0.98684 | 1.30E-43 |
| ENSG00000198805.12 | PNP | 0.748285 | 1.15E-13 |
| ENSG00000198814.13 | GK | 0.689292 | 1.54E-10 |
| ENSG00000198826.11 | ARHGAP11A | 0.618864 | 2.34E-07 |
| ENSG00000198855.7 | FICD | 0.838779 | 3.92E-20 |
| ENSG00000203497.2 | PDCD4-AS1 | -0.90411 | 0.046763 |
| ENSG00000203618.6 | GP1BB | 0.93195 | 0.039657 |
| ENSG00000204131.9 | NHSL2 | -0.95828 | 1.30E-26 |
| ENSG00000204228.4 | HSD17B8 | -1.05064 | 0.002121 |
| ENSG00000204314.12 | PRRT1 | -0.75192 | 0.000995 |
| ENSG00000204682.8 | MIR1915HG | -0.60265 | 0.020169 |
| ENSG00000205978.6 | NYNRIN | -0.72456 | 4.46E-20 |
| ENSG00000206190.12 | ATP10A | 1.149432 | 5.22E-16 |
| ENSG00000206538.9 | VGLL3 | 0.652092 | 8.50E-31 |
| ENSG00000212724.3 | KRTAP2-3 | 2.844297 | 4.91E-08 |
| ENSG00000212864.3 | RNF208 | -0.6707 | 0.0002 |
| ENSG00000213190.4 | MLLT11 | 0.597742 | 6.36E-14 |
| ENSG00000213626.13 | LBH | -0.81074 | 1.59E-05 |
| ENSG00000213742.7 | ZNF337-AS1 | -0.60755 | 0.032494 |
| ENSG00000213760.11 | ATP6V1G2 | -0.60802 | 1.97E-05 |
| ENSG00000213930.12 | GALT | -0.6175 | 7.81E-08 |
| ENSG00000214212.9 | C19orf38 | -1.23556 | 4.20E-07 |
| ENSG00000214338.10 | SOGA3 | 0.766821 | 2.74E-05 |
| ENSG00000215861.6 | WI2-1896O14.1 | 1.096686 | 5.26E-25 |
| ENSG00000215883.11 | CYB5RL | -0.63405 | 8.18E-11 |
| ENSG00000217801.10 | RP11-465B22.3 | 1.538258 | 0.005826 |
| ENSG00000221818.9 | EBF2 | -1.29279 | 1.30E-29 |
| ENSG00000221990.5 | EXOC3-AS1 | -0.60714 | 0.00893 |
| ENSG00000222041.11 | CYTOR | 0.657953 | 7.04E-26 |
| ENSG00000223403.6 | MEG9 | 0.621407 | 0.000619 |
| ENSG00000223478.1 | RP11-545E17.3 | -0.89173 | 0.043874 |
| ENSG00000223485.4 | LINC01615 | 0.816933 | 0.014724 |
| ENSG00000223802.9 | CERS1 | -0.66539 | 1.86E-13 |
| ENSG00000225614.4 | ZNF469 | 0.627351 | 1.44E-28 |
| ENSG00000226380.9 | AC058791.1 | 1.020337 | 1.28E-10 |
| ENSG00000227051.7 | C14orf132 | -0.71091 | 1.29E-32 |
| ENSG00000227268.5 | KLLN | -0.62326 | 0.003661 |
| ENSG00000228288.7 | PCAT6 | -0.69171 | 0.023335 |
| ENSG00000229689.3 | AC009237.8 | 0.626608 | 0.001751 |
| ENSG00000229915.1 | AC016999.2 | 1.851668 | 0.021968 |
| ENSG00000232533.1 | AC093673.5 | 0.776136 | 1.12E-05 |
| ENSG00000233024.7 | NPIPA9 | 0.63621 | 1.11E-08 |
| ENSG00000233058.2 | LINC00884 | 0.601388 | 1.93E-07 |

| | | | |
|---|---|---|---|
| ENSG00000233098.9 | CCDC144NL-AS1 | 1.447091 | 1.10E-09 |
| ENSG00000233117.4 | LINC00702 | 1.14553 | 1.01E-06 |
| ENSG00000233223.3 | AC113189.5 | -0.76424 | 0.002542 |
| ENSG00000233695.2 | GAS6-AS1 | -1.2456 | 4.63E-12 |
| ENSG00000235162.9 | C12orf75 | 0.889534 | 1.92E-55 |
| ENSG00000235169.11 | SMIM1 | -0.6956 | 0.033784 |
| ENSG00000235217.6 | TSPY26P | -0.60789 | 1.58E-05 |
| ENSG00000235505.7 | CASP4LP | 0.869063 | 0.031998 |
| ENSG00000235863.4 | B3GALT4 | -0.66175 | 0.004645 |
| ENSG00000237352.4 | LINC01358 | -0.64656 | 0.021715 |
| ENSG00000240065.8 | PSMB9 | -0.90795 | 0.000125 |
| ENSG00000241399.7 | CD302 | -0.65991 | 3.13E-20 |
| ENSG00000241749.4 | RPSAP52 | 1.370922 | 2.32E-05 |
| ENSG00000242265.6 | PEG10 | -0.79265 | 1.38E-69 |
| ENSG00000243056.2 | EIF4EBP3 | -0.73034 | 0.000113 |
| ENSG00000243137.8 | PSG4 | 1.707414 | 3.42E-27 |
| ENSG00000243244.7 | STON1 | -0.64148 | 1.56E-16 |
| ENSG00000243811.12 | APOBEC3D | -0.76714 | 0.004158 |
| ENSG00000245573.9 | BDNF-AS | -0.64342 | 0.010376 |
| ENSG00000245694.10 | CRNDE | 0.648884 | 0.000454 |
| ENSG00000246430.7 | LINC00968 | 1.281527 | 2.53E-10 |
| ENSG00000246898.1 | LINC00920 | -0.69866 | 0.001237 |
| ENSG00000247095.3 | MIR210HG | 0.772305 | 0.002162 |
| ENSG00000248441.7 | LINC01197 | -0.88754 | 0.001872 |
| ENSG00000248890.2 | HHIP-AS1 | -0.58679 | 0.002987 |
| ENSG00000249279.6 | LINC02057 | 1.393778 | 5.68E-05 |
| ENSG00000249669.10 | CARMN | 0.898078 | 3.47E-28 |
| ENSG00000250273.1 | PSMC1P5 | 1.932735 | 0.005937 |
| ENSG00000250303.4 | LINC02762 | 0.64068 | 0.00046 |
| ENSG00000250510.8 | GPR162 | -0.78108 | 4.89E-07 |
| ENSG00000253227.2 | RP11-383J24.1 | 0.811409 | 0.007493 |
| ENSG00000253414.3 | LINC01605 | -0.8915 | 0.022001 |
| ENSG00000253552.8 | HOXA-AS2 | -0.78789 | 0.000117 |
| ENSG00000254602.2 | AP000662.4 | -1.06658 | 0.000203 |
| ENSG00000255182.2 | CTD-2517M22.14 | -0.63214 | 0.008369 |
| ENSG00000255471.1 | RP11-736K20.5 | -1.09923 | 2.64E-07 |
| ENSG00000255690.3 | TRIL | -1.09165 | 2.75E-09 |
| ENSG00000256671.6 | LIMS4 | 0.996791 | 0.000117 |
| ENSG00000257108.2 | NHLRC4 | -1.06709 | 0.000244 |
| ENSG00000257219.6 | LNCOG | 0.925406 | 0.00431 |
| ENSG00000258643.6 | BCL2L2-PABPN1 | 0.595474 | 0.000344 |
| ENSG00000258644.6 | SYNJ2BP-COX16 | 1.264058 | 0.047303 |
| ENSG00000259207.9 | ITGB3 | 2.038566 | ######## |
| ENSG00000259426.6 | RP11-253M7.1 | 1.348447 | 8.95E-12 |
| ENSG00000259721.1 | RP11-758N13.1 | 0.979483 | 0.000934 |
| ENSG00000260293.2 | RP11-715J22.6 | -0.84816 | 1.34E-05 |
| ENSG00000261040.7 | WFDC21P | 0.597728 | 0.000537 |
| ENSG00000261371.6 | PECAM1 | -0.85538 | 2.30E-12 |
| ENSG00000261455.1 | LINC01003 | -0.68064 | 0.001752 |
| ENSG00000262049.1 | RP13-1032I1.7 | -0.61698 | 0.004898 |
| ENSG00000262074.7 | SNORD3B-2 | 0.690693 | 0.006712 |
| ENSG00000263155.6 | MYZAP | -0.85804 | 0.037716 |
| ENSG00000264501.2 | RN7SL731P | 1.170044 | 0.02617 |
| ENSG00000266094.8 | RASSF5 | -0.98759 | 1.29E-21 |
| ENSG00000266714.9 | MYO15B | -0.69558 | 0.010716 |
| ENSG00000268388.6 | FENDRR | 0.674301 | 0.00024 |
| ENSG00000270210.1 | RP11-373D23.3 | 0.952418 | 0.018379 |
| ENSG00000270757.1 | HSPE1-MOB4 | 0.673353 | 0.013949 |
| ENSG00000271303.2 | SRXN1 | 0.586751 | 1.14E-19 |
| ENSG00000272695.2 | GAS6-DT | 0.696409 | 1.18E-05 |
| ENSG00000273173.5 | SNURF | -0.97811 | 0.032251 |
| ENSG00000275131.3 | PDE4DIPP2 | 0.591604 | 2.94E-09 |
| ENSG00000276043.5 | UHRF1 | 0.793474 | 2.05E-10 |

| | | | |
|---|---|---|---|
| ENSG00000277639.2 | RP11-467J12.4 | -0.83428 | 0.001591 |
| ENSG00000277775.2 | H3C7 | -0.80131 | 0.000104 |
| ENSG00000277778.2 | PGM5P2 | 0.841578 | 0.006351 |
| ENSG00000277971.1 | XXbac-B562F10.12 | 1.360015 | 0.002387 |
| ENSG00000278224.7 | PRICKLE4 | -1.15943 | 0.040817 |
| ENSG00000278903.3 | CH507-145C22.1 | -1.22861 | 0.002644 |
| ENSG00000278934.1 | CTD-2006M22.2 | -0.90754 | 4.13E-05 |
| ENSG00000278948.1 | RP5-1039K5.12 | 0.636722 | 3.22E-12 |
| ENSG00000278962.1 | RP11-399B17.1 | 0.598708 | 0.004052 |
| ENSG00000279117.1 | CTD-2562J17.6 | -0.94222 | 0.002502 |
| ENSG00000279192.1 | PWAR5 | -0.81544 | 0.002075 |
| ENSG00000279821.1 | RP11-1334A24.5 | 0.983343 | 3.84E-05 |
| ENSG00000280339.1 | RP11-736K20.4 | -1.18892 | 7.93E-13 |
| ENSG00000280441.3 | CH507-528H12.1 | -0.62419 | 5.67E-07 |
| ENSG00000282057.1 | RP4-621F18.2 | 0.847449 | 1.08E-12 |
| ENSG00000283154.2 | IQCJ-SCHIP1 | 1.32861 | 3.61E-28 |
| ENSG00000285006.1 | RP1-240K6.5 | -1.6033 | 0.017279 |
| ENSG00000285106.2 | RP11-36B6.2 | 0.680322 | 5.32E-05 |
| ENSG00000285347.1 | CTD-2308N23.4 | -1.61738 | 0.00305 |
| ENSG00000285508.1 | RP5-931K24.3 | 6.511029 | 2.16E-06 |
| ENSG00000285901.2 | RP11-388F6.5 | 0.855349 | 2.92E-06 |
| ENSG00000285976.2 | RP5-1148A21.4 | -3.49145 | 0.00538 |
| ENSG00000286190.2 | RP11-807C20.3 | 0.644328 | 1.97E-16 |
| ENSG00000287839.1 | RP11-29H23.8 | -0.69332 | 0.001862 |
| ENSG00000287979.1 | CH17-3B23.3 | 1.16433 | 0.00127 |
| ENSG00000288620.1 | RNF216P1 | 0.737218 | 0.011097 |

**Table S3. Gene Ontology analysis of genes regulated by colchicine intervention compared to cholesterol loaded HASMCs.** HASMCs were cholesterol loaded (10µg/mL) for 72 hours, followed by treatment with colchicine (50nM) for 48 hours.

| GO.ID | Term | Annotated | Significant | Expected | classicFisher |
|---|---|---|---|---|---|
| GO:0032501 | multicellular organismal process | 4083 | 473 | 305.91 | < 1e-30 |
| GO:0007155 | cell adhesion | 824 | 165 | 61.74 | < 1e-30 |
| GO:0022610 | biological adhesion | 827 | 165 | 61.96 | < 1e-30 |
| GO:0048731 | system development | 2858 | 368 | 214.13 | < 1e-30 |
| GO:0048856 | anatomical structure development | 3469 | 416 | 259.91 | < 1e-30 |
| GO:0009653 | anatomical structure morphogenesis | 1669 | 250 | 125.05 | < 1e-30 |
| GO:0007584 | response to nutrient | 106 | 20 | 7.94 | 0.0001 |
| GO:0032944 | regulation of mononuclear cell proliferation | 106 | 20 | 7.94 | 0.0001 |
| GO:0032270 | positive regulation of cellular protein metabolic process | 1008 | 107 | 75.52 | 0.0001 |
| GO:0055021 | regulation of cardiac muscle tissue growth | 46 | 12 | 3.45 | 0.0001 |
| GO:0018108 | peptidyl-tyrosine phosphorylation | 212 | 32 | 15.88 | 0.00011 |
| GO:0006690 | icosanoid metabolic process | 53 | 13 | 3.97 | 0.00011 |
| GO:0015732 | prostaglandin transport | 8 | 5 | 0.6 | 0.00011 |
| GO:0032354 | response to follicle-stimulating hormone | 8 | 5 | 0.6 | 0.00011 |
| GO:0042448 | progesterone metabolic process | 8 | 5 | 0.6 | 0.00011 |
| GO:0072102 | glomerulus morphogenesis | 8 | 5 | 0.6 | 0.00011 |
| GO:0001759 | organ induction | 12 | 6 | 0.9 | 0.00011 |
| GO:0034762 | regulation of transmembrane transport | 270 | 38 | 20.23 | 0.00011 |
| GO:0030038 | contractile actin filament bundle assembly | 83 | 17 | 6.22 | 0.00012 |
| GO:0043149 | stress fiber assembly | 83 | 17 | 6.22 | 0.00012 |
| GO:0003158 | endothelium development | 91 | 18 | 6.82 | 0.00012 |
| GO:1905207 | regulation of cardiocyte differentiation | 22 | 8 | 1.65 | 0.00012 |
| GO:1901888 | regulation of cell junction assembly | 124 | 22 | 9.29 | 0.00012 |
| GO:0002062 | chondrocyte differentiation | 68 | 15 | 5.09 | 0.00012 |
| GO:0046620 | regulation of organ growth | 68 | 15 | 5.09 | 0.00012 |
| GO:0050728 | negative regulation of inflammatory response | 68 | 15 | 5.09 | 0.00012 |
| GO:0003208 | cardiac ventricle morphogenesis | 40 | 11 | 3 | 0.00012 |
| GO:0070527 | platelet aggregation | 40 | 11 | 3 | 0.00012 |
| GO:0001817 | regulation of cytokine production | 411 | 52 | 30.79 | 0.00012 |
| GO:0052548 | regulation of endopeptidase activity | 242 | 35 | 18.13 | 0.00012 |
| GO:0032496 | response to lipopolysaccharide | 186 | 29 | 13.94 | 0.00012 |
| GO:0051046 | regulation of secretion | 330 | 44 | 24.72 | 0.00012 |
| GO:0030510 | regulation of BMP signaling pathway | 61 | 14 | 4.57 | 0.00013 |
| GO:0043269 | regulation of ion transport | 733 | 82 | 54.92 | 0.00013 |
| GO:0002673 | regulation of acute inflammatory response | 17 | 7 | 1.27 | 0.00013 |
| GO:0010875 | positive regulation of cholesterol efflux | 17 | 7 | 1.27 | 0.00013 |
| GO:0045214 | sarcomere organization | 17 | 7 | 1.27 | 0.00013 |
| GO:0003231 | cardiac ventricle development | 76 | 16 | 5.69 | 0.00013 |
| GO:0043535 | regulation of blood vessel endothelial cell migration | 76 | 16 | 5.69 | 0.00013 |
| GO:0035924 | cellular response to vascular endothelial growth factor stimulus | 47 | 12 | 3.52 | 0.00013 |
| GO:0060191 | regulation of lipase activity | 47 | 12 | 3.52 | 0.00013 |
| GO:0060395 | SMAD protein signal transduction | 47 | 12 | 3.52 | 0.00013 |
| GO:2000106 | regulation of leukocyte apoptotic process | 47 | 12 | 3.52 | 0.00013 |
| GO:0006873 | cellular ion homeostasis | 341 | 45 | 25.55 | 0.00013 |
| GO:0060193 | positive regulation of lipase activity | 34 | 10 | 2.55 | 0.00013 |
| GO:0010518 | positive regulation of phospholipase activity | 28 | 9 | 2.1 | 0.00013 |
| GO:0048546 | digestive tract morphogenesis | 28 | 9 | 2.1 | 0.00013 |
| GO:0110151 | positive regulation of biomineralization | 28 | 9 | 2.1 | 0.00013 |
| GO:2000242 | negative regulation of reproductive process | 28 | 9 | 2.1 | 0.00013 |
| GO:0007565 | female pregnancy | 100 | 19 | 7.49 | 0.00013 |
| GO:0031399 | regulation of protein modification process | 1142 | 118 | 85.56 | 0.00014 |
| GO:0030168 | platelet activation | 92 | 18 | 6.89 | 0.00014 |
| GO:0046661 | male sex differentiation | 92 | 18 | 6.89 | 0.00014 |
| GO:0009628 | response to abiotic stimulus | 858 | 93 | 64.28 | 0.00014 |
| GO:0045321 | leukocyte activation | 736 | 82 | 55.14 | 0.00015 |
| GO:0045187 | regulation of circadian sleep/wake cycle, sleep | 5 | 4 | 0.37 | 0.00015 |

| | | | | | |
|---|---|---|---|---|---|
| GO:0035265 | organ growth | 109 | 20 | 8.17 | 0.00015 |
| GO:0007548 | sex differentiation | 152 | 25 | 11.39 | 0.00015 |
| GO:0048565 | digestive tract development | 77 | 16 | 5.77 | 0.00015 |
| GO:0060389 | pathway-restricted SMAD protein phosphorylation | 41 | 11 | 3.07 | 0.00015 |
| GO:1905954 | positive regulation of lipid localization | 62 | 14 | 4.65 | 0.00015 |
| GO:0051246 | regulation of protein metabolic process | 1821 | 175 | 136.43 | 0.00016 |
| GO:0051049 | regulation of transport | 1008 | 106 | 75.52 | 0.00016 |
| GO:0071407 | cellular response to organic cyclic compound | 354 | 46 | 26.52 | 0.00016 |
| GO:0051153 | regulation of striated muscle cell differentiation | 55 | 13 | 4.12 | 0.00016 |
| GO:0051781 | positive regulation of cell division | 48 | 12 | 3.6 | 0.00016 |
| GO:0016485 | protein processing | 144 | 24 | 10.79 | 0.00016 |
| GO:0016310 | phosphorylation | 1452 | 144 | 108.79 | 0.00016 |
| GO:0034765 | regulation of ion transmembrane transport | 265 | 37 | 19.85 | 0.00017 |
| GO:0031344 | regulation of cell projection organization | 437 | 54 | 32.74 | 0.00017 |
| GO:0002521 | leukocyte differentiation | 285 | 39 | 21.35 | 0.00017 |
| GO:0120035 | regulation of plasma membrane bounded cell projection organization | 427 | 53 | 31.99 | 0.00017 |
| GO:0010517 | regulation of phospholipase activity | 35 | 10 | 2.62 | 0.00017 |
| GO:0048732 | gland development | 266 | 37 | 19.93 | 0.00018 |
| GO:0031641 | regulation of myelination | 29 | 9 | 2.17 | 0.00018 |
| GO:0045744 | negative regulation of G protein-coupled receptor signaling pathway | 29 | 9 | 2.17 | 0.00018 |
| GO:0002682 | regulation of immune system process | 864 | 93 | 64.73 | 0.00018 |
| GO:0014743 | regulation of muscle hypertrophy | 42 | 11 | 3.15 | 0.00019 |
| GO:0045860 | positive regulation of protein kinase activity | 347 | 45 | 26 | 0.0002 |
| GO:0001709 | cell fate determination | 18 | 7 | 1.35 | 0.0002 |
| GO:0002691 | regulation of cellular extravasation | 18 | 7 | 1.35 | 0.0002 |
| GO:0010092 | specification of animal organ identity | 18 | 7 | 1.35 | 0.0002 |
| GO:1901615 | organic hydroxy compound metabolic process | 327 | 43 | 24.5 | 0.0002 |
| GO:0070665 | positive regulation of leukocyte proliferation | 71 | 15 | 5.32 | 0.0002 |
| GO:0035966 | response to topologically incorrect protein | 182 | 28 | 13.64 | 0.0002 |
| GO:0022408 | negative regulation of cell-cell adhesion | 103 | 19 | 7.72 | 0.0002 |
| GO:0044283 | small molecule biosynthetic process | 462 | 56 | 34.61 | 0.00021 |
| GO:0048667 | cell morphogenesis involved in neuron differentiation | 379 | 48 | 28.4 | 0.00022 |
| GO:0002237 | response to molecule of bacterial origin | 192 | 29 | 14.39 | 0.00022 |
| GO:0008406 | gonad development | 129 | 22 | 9.66 | 0.00022 |
| GO:0007498 | mesoderm development | 64 | 14 | 4.8 | 0.00022 |
| GO:0050671 | positive regulation of lymphocyte proliferation | 64 | 14 | 4.8 | 0.00022 |
| GO:0014015 | positive regulation of gliogenesis | 36 | 10 | 2.7 | 0.00022 |
| GO:0072577 | endothelial cell apoptotic process | 36 | 10 | 2.7 | 0.00022 |
| GO:1905330 | regulation of morphogenesis of an epithelium | 36 | 10 | 2.7 | 0.00022 |
| GO:0043281 | regulation of cysteine-type endopeptidase activity involved in apoptotic process | 147 | 24 | 11.01 | 0.00023 |
| GO:0072224 | metanephric glomerulus development | 9 | 5 | 0.67 | 0.00023 |
| GO:1901722 | regulation of cell proliferation involved in kidney development | 9 | 5 | 0.67 | 0.00023 |
| GO:0050865 | regulation of cell activation | 299 | 40 | 22.4 | 0.00023 |
| GO:0044093 | positive regulation of molecular function | 1180 | 120 | 88.41 | 0.00023 |
| GO:0032963 | collagen metabolic process | 72 | 15 | 5.39 | 0.00024 |
| GO:0070167 | regulation of biomineral tissue development | 57 | 13 | 4.27 | 0.00024 |
| GO:0010604 | positive regulation of macromolecule metabolic process | 2363 | 218 | 177.04 | 0.00024 |
| GO:0032268 | regulation of cellular protein metabolic process | 1703 | 164 | 127.59 | 0.00024 |
| GO:0048146 | positive regulation of fibroblast proliferation | 30 | 9 | 2.25 | 0.00024 |
| GO:0061383 | trabecula morphogenesis | 30 | 9 | 2.25 | 0.00024 |
| GO:0001523 | retinoid metabolic process | 43 | 11 | 3.22 | 0.00024 |
| GO:0051145 | smooth muscle cell differentiation | 43 | 11 | 3.22 | 0.00024 |
| GO:0007159 | leukocyte cell-cell adhesion | 184 | 28 | 13.79 | 0.00024 |
| GO:0048660 | regulation of smooth muscle cell proliferation | 88 | 17 | 6.59 | 0.00024 |
| GO:0014013 | regulation of gliogenesis | 50 | 12 | 3.75 | 0.00025 |
| GO:0001819 | positive regulation of cytokine production | 241 | 34 | 18.06 | 0.00025 |
| GO:0042391 | regulation of membrane potential | 203 | 30 | 15.21 | 0.00025 |
| GO:0032946 | positive regulation of mononuclear cell proliferation | 65 | 14 | 4.87 | 0.00026 |
| GO:0071560 | cellular response to transforming growth factor beta stimulus | 185 | 28 | 13.86 | 0.00027 |
| GO:0007611 | learning or memory | 131 | 22 | 9.81 | 0.00027 |
| GO:0009792 | embryo development ending in birth or egg hatching | 414 | 51 | 31.02 | 0.00027 |
| GO:0007411 | axon guidance | 167 | 26 | 12.51 | 0.00028 |
| GO:0055123 | digestive system development | 81 | 16 | 6.07 | 0.00028 |

| | | | | | |
|---|---|---|---|---|---|
| GO:0110149 | regulation of biomineralization | 58 | 13 | 4.35 | 0.00028 |
| GO:0048545 | response to steroid hormone | 214 | 31 | 16.03 | 0.00028 |
| GO:0001706 | endoderm formation | 37 | 10 | 2.77 | 0.00028 |
| GO:0055078 | sodium ion homeostasis | 19 | 7 | 1.42 | 0.00029 |
| GO:0090050 | positive regulation of cell migration involved in sprouting angiogenesis | 19 | 7 | 1.42 | 0.00029 |
| GO:0034308 | primary alcohol metabolic process | 51 | 12 | 3.82 | 0.0003 |
| GO:1903530 | regulation of secretion by cell | 313 | 41 | 23.45 | 0.0003 |
| GO:0045137 | development of primary sexual characteristics | 132 | 22 | 9.89 | 0.0003 |
| GO:1902903 | regulation of supramolecular fiber organization | 283 | 38 | 21.2 | 0.0003 |
| GO:0097485 | neuron projection guidance | 168 | 26 | 12.59 | 0.00031 |
| GO:2000116 | regulation of cysteine-type endopeptidase activity | 159 | 25 | 11.91 | 0.00031 |
| GO:0003299 | muscle hypertrophy in response to stress | 14 | 6 | 1.05 | 0.00031 |
| GO:0014887 | cardiac muscle adaptation | 14 | 6 | 1.05 | 0.00031 |
| GO:0014898 | cardiac muscle hypertrophy in response to stress | 14 | 6 | 1.05 | 0.00031 |
| GO:0034698 | response to gonadotropin | 14 | 6 | 1.05 | 0.00031 |
| GO:0035967 | cellular response to topologically incorrect protein | 150 | 24 | 11.24 | 0.00031 |
| GO:0070661 | leukocyte proliferation | 150 | 24 | 11.24 | 0.00031 |
| GO:0051147 | regulation of muscle cell differentiation | 98 | 18 | 7.34 | 0.00031 |
| GO:0043009 | chordate embryonic development | 406 | 50 | 30.42 | 0.00032 |
| GO:0043090 | amino acid import | 31 | 9 | 2.32 | 0.00032 |
| GO:0048659 | smooth muscle cell proliferation | 90 | 17 | 6.74 | 0.00032 |
| GO:0007229 | integrin-mediated signaling pathway | 74 | 15 | 5.54 | 0.00032 |
| GO:0110020 | regulation of actomyosin structure organization | 74 | 15 | 5.54 | 0.00032 |
| GO:0002683 | negative regulation of immune system process | 226 | 32 | 16.93 | 0.00035 |
| GO:0001933 | negative regulation of protein phosphorylation | 275 | 37 | 20.6 | 0.00035 |
| GO:0048645 | animal organ formation | 38 | 10 | 2.85 | 0.00036 |
| GO:0060393 | regulation of pathway-restricted SMAD protein phosphorylation | 38 | 10 | 2.85 | 0.00036 |
| GO:0060048 | cardiac muscle contraction | 67 | 14 | 5.02 | 0.00037 |
| GO:0090066 | regulation of anatomical structure size | 326 | 42 | 24.42 | 0.00037 |
| GO:0001952 | regulation of cell-matrix adhesion | 91 | 17 | 6.82 | 0.00037 |
| GO:0002088 | lens development in camera-type eye | 45 | 11 | 3.37 | 0.00037 |
| GO:0042102 | positive regulation of T cell proliferation | 45 | 11 | 3.37 | 0.00037 |
| GO:0010975 | regulation of neuron projection development | 296 | 39 | 22.18 | 0.00037 |
| GO:1903409 | reactive oxygen species biosynthetic process | 75 | 15 | 5.62 | 0.00038 |
| GO:0071559 | response to transforming growth factor beta | 189 | 28 | 14.16 | 0.00038 |
| GO:0042098 | T cell proliferation | 100 | 18 | 7.49 | 0.0004 |
| GO:0051962 | positive regulation of nervous system development | 171 | 26 | 12.81 | 0.0004 |
| GO:0030514 | negative regulation of BMP signaling pathway | 32 | 9 | 2.4 | 0.00041 |
| GO:0032330 | regulation of chondrocyte differentiation | 32 | 9 | 2.4 | 0.00041 |
| GO:0048662 | negative regulation of smooth muscle cell proliferation | 32 | 9 | 2.4 | 0.00041 |
| GO:0060322 | head development | 485 | 57 | 36.34 | 0.00041 |
| GO:0007567 | parturition | 6 | 4 | 0.45 | 0.00042 |
| GO:0031650 | regulation of heat generation | 6 | 4 | 0.45 | 0.00042 |
| GO:0032306 | regulation of prostaglandin secretion | 6 | 4 | 0.45 | 0.00042 |
| GO:0032308 | positive regulation of prostaglandin secretion | 6 | 4 | 0.45 | 0.00042 |
| GO:0042749 | regulation of circadian sleep/wake cycle | 6 | 4 | 0.45 | 0.00042 |
| GO:0051956 | negative regulation of amino acid transport | 6 | 4 | 0.45 | 0.00042 |
| GO:0061304 | retinal blood vessel morphogenesis | 6 | 4 | 0.45 | 0.00042 |
| GO:0072007 | mesangial cell differentiation | 6 | 4 | 0.45 | 0.00042 |
| GO:1901724 | positive regulation of cell proliferation involved in kidney development | 6 | 4 | 0.45 | 0.00042 |
| GO:0003298 | physiological muscle hypertrophy | 20 | 7 | 1.5 | 0.00042 |
| GO:0003301 | physiological cardiac muscle hypertrophy | 20 | 7 | 1.5 | 0.00042 |
| GO:0034694 | response to prostaglandin | 20 | 7 | 1.5 | 0.00042 |
| GO:0035886 | vascular associated smooth muscle cell differentiation | 20 | 7 | 1.5 | 0.00042 |
| GO:0061036 | positive regulation of cartilage development | 20 | 7 | 1.5 | 0.00042 |
| GO:0061049 | cell growth involved in cardiac muscle cell development | 20 | 7 | 1.5 | 0.00042 |
| GO:0070306 | lens fiber cell differentiation | 20 | 7 | 1.5 | 0.00042 |
| GO:0014745 | negative regulation of muscle adaptation | 10 | 5 | 0.75 | 0.00043 |
| GO:0072087 | renal vesicle development | 10 | 5 | 0.75 | 0.00043 |
| GO:0008584 | male gonad development | 84 | 16 | 6.29 | 0.00043 |
| GO:0098739 | import across plasma membrane | 84 | 16 | 6.29 | 0.00043 |
| GO:0015844 | monoamine transport | 26 | 8 | 1.95 | 0.00044 |
| GO:0055023 | positive regulation of cardiac muscle tissue growth | 26 | 8 | 1.95 | 0.00044 |

| | | | | | |
|---|---|---|---|---|---|
| GO:1904407 | positive regulation of nitric oxide metabolic process | 26 | 8 | 1.95 | 0.00044 |
| GO:0006812 | cation transport | 573 | 65 | 42.93 | 0.00045 |
| GO:0021675 | nerve development | 39 | 10 | 2.92 | 0.00045 |
| GO:0045428 | regulation of nitric oxide biosynthetic process | 39 | 10 | 2.92 | 0.00045 |
| GO:1903037 | regulation of leukocyte cell-cell adhesion | 163 | 25 | 12.21 | 0.00045 |
| GO:0002690 | positive regulation of leukocyte chemotaxis | 46 | 11 | 3.45 | 0.00046 |
| GO:0014910 | regulation of smooth muscle cell migration | 46 | 11 | 3.45 | 0.00046 |
| GO:0030500 | regulation of bone mineralization | 46 | 11 | 3.45 | 0.00046 |
| GO:0030217 | T cell differentiation | 127 | 21 | 9.52 | 0.00046 |
| GO:0043588 | skin development | 136 | 22 | 10.19 | 0.00047 |
| GO:0003006 | developmental process involved in reproduction | 520 | 60 | 38.96 | 0.00048 |
| GO:0001676 | long-chain fatty acid metabolic process | 61 | 13 | 4.57 | 0.00048 |
| GO:0016525 | negative regulation of angiogenesis | 61 | 13 | 4.57 | 0.00048 |
| GO:0150116 | regulation of cell-substrate junction organization | 61 | 13 | 4.57 | 0.00048 |
| GO:1901343 | negative regulation of vasculature development | 61 | 13 | 4.57 | 0.00048 |
| GO:2000181 | negative regulation of blood vessel morphogenesis | 61 | 13 | 4.57 | 0.00048 |
| GO:0060343 | trabecula formation | 15 | 6 | 1.12 | 0.00048 |
| GO:0097191 | extrinsic apoptotic signaling pathway | 173 | 26 | 12.96 | 0.00049 |
| GO:0006469 | negative regulation of protein kinase activity | 192 | 28 | 14.39 | 0.00049 |
| GO:0010469 | regulation of signaling receptor activity | 85 | 16 | 6.37 | 0.0005 |
| GO:0046546 | development of primary male sexual characteristics | 85 | 16 | 6.37 | 0.0005 |
| GO:2001236 | regulation of extrinsic apoptotic signaling pathway | 119 | 20 | 8.92 | 0.0005 |
| GO:0008277 | regulation of G protein-coupled receptor signaling pathway | 77 | 15 | 5.77 | 0.00051 |
| GO:0030148 | sphingolipid biosynthetic process | 77 | 15 | 5.77 | 0.00051 |
| GO:0044281 | small molecule metabolic process | 1298 | 128 | 97.25 | 0.00051 |
| GO:0071496 | cellular response to external stimulus | 231 | 32 | 17.31 | 0.00051 |
| GO:0043270 | positive regulation of ion transport | 394 | 48 | 29.52 | 0.00053 |
| GO:0060425 | lung morphogenesis | 33 | 9 | 2.47 | 0.00053 |
| GO:0051130 | positive regulation of cellular component organization | 835 | 88 | 62.56 | 0.00055 |
| GO:0016101 | diterpenoid metabolic process | 47 | 11 | 3.52 | 0.00056 |
| GO:0060038 | cardiac muscle cell proliferation | 40 | 10 | 3 | 0.00056 |
| GO:2001238 | positive regulation of extrinsic apoptotic signaling pathway | 40 | 10 | 3 | 0.00056 |
| GO:0060041 | retina development in camera-type eye | 78 | 15 | 5.84 | 0.00058 |
| GO:0051492 | regulation of stress fiber assembly | 70 | 14 | 5.24 | 0.00058 |
| GO:0016053 | organic acid biosynthetic process | 194 | 28 | 14.53 | 0.00059 |
| GO:0002686 | negative regulation of leukocyte migration | 27 | 8 | 2.02 | 0.00059 |
| GO:0010862 | positive regulation of pathway-restricted SMAD protein phosphorylation | 27 | 8 | 2.02 | 0.00059 |
| GO:0022602 | ovulation cycle process | 27 | 8 | 2.02 | 0.00059 |
| GO:0000188 | inactivation of MAPK activity | 21 | 7 | 1.57 | 0.00059 |
| GO:0008207 | C21-steroid hormone metabolic process | 21 | 7 | 1.57 | 0.00059 |
| GO:0036499 | PERK-mediated unfolded protein response | 21 | 7 | 1.57 | 0.00059 |
| GO:0061384 | heart trabecula morphogenesis | 21 | 7 | 1.57 | 0.00059 |
| GO:0050863 | regulation of T cell activation | 166 | 25 | 12.44 | 0.0006 |
| GO:0051493 | regulation of cytoskeleton organization | 386 | 47 | 28.92 | 0.00061 |
| GO:0090101 | negative regulation of transmembrane receptor protein serine/threonine kinase signaling pathway | 95 | 17 | 7.12 | 0.00062 |
| GO:0010466 | negative regulation of peptidase activity | 121 | 20 | 9.07 | 0.00063 |
| GO:0007613 | memory | 55 | 12 | 4.12 | 0.00063 |
| GO:0019725 | cellular homeostasis | 537 | 61 | 40.23 | 0.00064 |
| GO:0007420 | brain development | 451 | 53 | 33.79 | 0.00066 |
| GO:0046394 | carboxylic acid biosynthetic process | 186 | 27 | 13.94 | 0.00067 |
| GO:0043085 | positive regulation of catalytic activity | 955 | 98 | 71.55 | 0.00067 |
| GO:0033627 | cell adhesion mediated by integrin | 48 | 11 | 3.6 | 0.00068 |
| GO:0060411 | cardiac septum morphogenesis | 48 | 11 | 3.6 | 0.00068 |
| GO:0043393 | regulation of protein binding | 158 | 24 | 11.84 | 0.00068 |
| GO:0050848 | regulation of calcium-mediated signaling | 34 | 9 | 2.55 | 0.00068 |
| GO:1903531 | negative regulation of secretion by cell | 71 | 14 | 5.32 | 0.00068 |
| GO:0010611 | regulation of cardiac muscle hypertrophy | 41 | 10 | 3.07 | 0.0007 |
| GO:0031102 | neuron projection regeneration | 41 | 10 | 3.07 | 0.0007 |
| GO:0040007 | growth | 627 | 69 | 46.98 | 0.0007 |
| GO:0046890 | regulation of lipid biosynthetic process | 131 | 21 | 9.81 | 0.0007 |
| GO:0045944 | positive regulation of transcription by RNA polymerase II | 819 | 86 | 61.36 | 0.00072 |
| GO:0048333 | mesodermal cell differentiation | 16 | 6 | 1.2 | 0.00072 |
| GO:0050927 | positive regulation of positive chemotaxis | 16 | 6 | 1.2 | 0.00072 |

| | | | | | |
|---|---|---|---|---|---|
| GO:0070232 | regulation of T cell apoptotic process | 16 | 6 | 1.2 | 0.00072 |
| GO:0030431 | sleep | 11 | 5 | 0.82 | 0.00074 |
| GO:0060231 | mesenchymal to epithelial transition | 11 | 5 | 0.82 | 0.00074 |
| GO:0120255 | olefinic compound biosynthetic process | 11 | 5 | 0.82 | 0.00074 |
| GO:0051348 | negative regulation of transferase activity | 236 | 32 | 17.68 | 0.00075 |
| GO:0051346 | negative regulation of hydrolase activity | 266 | 35 | 19.93 | 0.00075 |
| GO:0030858 | positive regulation of epithelial cell differentiation | 28 | 8 | 2.1 | 0.00077 |
| GO:0044319 | wound healing, spreading of cells | 28 | 8 | 2.1 | 0.00077 |
| GO:0046456 | icosanoid biosynthetic process | 28 | 8 | 2.1 | 0.00077 |
| GO:0071634 | regulation of transforming growth factor beta production | 28 | 8 | 2.1 | 0.00077 |
| GO:0090505 | epiboly involved in wound healing | 28 | 8 | 2.1 | 0.00077 |
| GO:0046651 | lymphocyte proliferation | 132 | 21 | 9.89 | 0.00078 |
| GO:0048705 | skeletal system morphogenesis | 132 | 21 | 9.89 | 0.00078 |
| GO:0031644 | regulation of nervous system process | 64 | 13 | 4.8 | 0.00078 |
| GO:0090100 | positive regulation of transmembrane receptor protein serine/threonine kinase signaling pathway | 72 | 14 | 5.39 | 0.00079 |
| GO:0006633 | fatty acid biosynthetic process | 97 | 17 | 7.27 | 0.00079 |
| GO:0014741 | negative regulation of muscle hypertrophy | 22 | 7 | 1.65 | 0.00081 |
| GO:0032373 | positive regulation of sterol transport | 22 | 7 | 1.65 | 0.00081 |
| GO:0032376 | positive regulation of cholesterol transport | 22 | 7 | 1.65 | 0.00081 |
| GO:0032653 | regulation of interleukin-10 production | 22 | 7 | 1.65 | 0.00081 |
| GO:0048009 | insulin-like growth factor receptor signaling pathway | 22 | 7 | 1.65 | 0.00081 |
| GO:0055010 | ventricular cardiac muscle tissue morphogenesis | 22 | 7 | 1.65 | 0.00081 |
| GO:0051050 | positive regulation of transport | 542 | 61 | 40.61 | 0.00081 |
| GO:0002011 | morphogenesis of an epithelial sheet | 49 | 11 | 3.67 | 0.00081 |
| GO:0001659 | temperature homeostasis | 106 | 18 | 7.94 | 0.00083 |
| GO:0033673 | negative regulation of kinase activity | 208 | 29 | 15.58 | 0.00083 |
| GO:0045444 | fat cell differentiation | 151 | 23 | 11.31 | 0.00083 |
| GO:0071774 | response to fibroblast growth factor | 89 | 16 | 6.67 | 0.00084 |
| GO:0031325 | positive regulation of cellular metabolic process | 2266 | 206 | 169.77 | 0.00085 |
| GO:0010951 | negative regulation of endopeptidase activity | 115 | 19 | 8.62 | 0.00085 |
| GO:0033674 | positive regulation of kinase activity | 392 | 47 | 29.37 | 0.00085 |
| GO:0002027 | regulation of heart rate | 42 | 10 | 3.15 | 0.00086 |
| GO:0051248 | negative regulation of protein metabolic process | 744 | 79 | 55.74 | 0.00086 |
| GO:1901698 | response to nitrogen compound | 744 | 79 | 55.74 | 0.00086 |
| GO:0023061 | signal release | 248 | 33 | 18.58 | 0.00087 |
| GO:0051128 | regulation of cellular component organization | 1713 | 161 | 128.34 | 0.00087 |
| GO:0031668 | cellular response to extracellular stimulus | 180 | 26 | 13.49 | 0.00089 |
| GO:0006955 | immune response | 1127 | 112 | 84.44 | 0.0009 |
| GO:0003337 | mesenchymal to epithelial transition involved in metanephros morphogenesis | 7 | 4 | 0.52 | 0.00091 |
| GO:0022410 | circadian sleep/wake cycle process | 7 | 4 | 0.52 | 0.00091 |
| GO:0031649 | heat generation | 7 | 4 | 0.52 | 0.00091 |
| GO:0032303 | regulation of icosanoid secretion | 7 | 4 | 0.52 | 0.00091 |
| GO:0032305 | positive regulation of icosanoid secretion | 7 | 4 | 0.52 | 0.00091 |
| GO:0032310 | prostaglandin secretion | 7 | 4 | 0.52 | 0.00091 |
| GO:0033605 | positive regulation of catecholamine secretion | 7 | 4 | 0.52 | 0.00091 |
| GO:0038063 | collagen-activated tyrosine kinase receptor signaling pathway | 7 | 4 | 0.52 | 0.00091 |
| GO:0050802 | circadian sleep/wake cycle, sleep | 7 | 4 | 0.52 | 0.00091 |
| GO:0072124 | regulation of glomerular mesangial cell proliferation | 7 | 4 | 0.52 | 0.00091 |
| GO:0072203 | cell proliferation involved in metanephros development | 7 | 4 | 0.52 | 0.00091 |
| GO:1900038 | negative regulation of cellular response to hypoxia | 7 | 4 | 0.52 | 0.00091 |
| GO:1904238 | pericyte cell differentiation | 7 | 4 | 0.52 | 0.00091 |
| GO:0071248 | cellular response to metal ion | 107 | 18 | 8.02 | 0.00093 |
| GO:0032943 | mononuclear cell proliferation | 134 | 21 | 10.04 | 0.00095 |
| GO:0045927 | positive regulation of growth | 162 | 24 | 12.14 | 0.00097 |
| GO:0010038 | response to metal ion | 210 | 29 | 15.73 | 0.00097 |
| GO:0045936 | negative regulation of phosphate metabolic process | 384 | 46 | 28.77 | 0.00098 |
| GO:0060421 | positive regulation of heart growth | 29 | 8 | 2.17 | 0.001 |
| GO:0090504 | epiboly | 29 | 8 | 2.17 | 0.001 |
| GO:1903556 | negative regulation of tumor necrosis factor superfamily cytokine production | 29 | 8 | 2.17 | 0.001 |
| GO:0050769 | positive regulation of neurogenesis | 153 | 23 | 11.46 | 0.001 |
| GO:0015718 | monocarboxylic acid transport | 82 | 15 | 6.14 | 0.00101 |
| GO:0001889 | liver development | 99 | 17 | 7.42 | 0.00101 |
| GO:0010563 | negative regulation of phosphorus metabolic process | 385 | 46 | 28.85 | 0.00104 |

| | | | | | |
|---|---|---|---|---|---|
| GO:0051173 | positive regulation of nitrogen compound metabolic process | 2138 | 195 | 160.18 | 0.00104 |
| GO:0042149 | cellular response to glucose starvation | 43 | 10 | 3.22 | 0.00104 |
| GO:1904019 | epithelial cell apoptotic process | 74 | 14 | 5.54 | 0.00104 |
| GO:0014912 | negative regulation of smooth muscle cell migration | 17 | 6 | 1.27 | 0.00105 |
| GO:0035336 | long-chain fatty-acyl-CoA metabolic process | 17 | 6 | 1.27 | 0.00105 |
| GO:0040036 | regulation of fibroblast growth factor receptor signaling pathway | 17 | 6 | 1.27 | 0.00105 |
| GO:0050926 | regulation of positive chemotaxis | 17 | 6 | 1.27 | 0.00105 |
| GO:0060441 | epithelial tube branching involved in lung morphogenesis | 17 | 6 | 1.27 | 0.00105 |
| GO:0071887 | leukocyte apoptotic process | 66 | 13 | 4.94 | 0.00106 |
| GO:2000379 | positive regulation of reactive oxygen species metabolic process | 66 | 13 | 4.94 | 0.00106 |
| GO:2000027 | regulation of animal organ morphogenesis | 126 | 20 | 9.44 | 0.00106 |
| GO:0038066 | p38MAPK cascade | 36 | 9 | 2.7 | 0.00106 |
| GO:0031346 | positive regulation of cell projection organization | 251 | 33 | 18.81 | 0.00107 |
| GO:0002548 | monocyte chemotaxis | 23 | 7 | 1.72 | 0.00109 |
| GO:0007157 | heterophilic cell-cell adhesion via plasma membrane cell adhesion molecules | 23 | 7 | 1.72 | 0.00109 |
| GO:0010874 | regulation of cholesterol efflux | 23 | 7 | 1.72 | 0.00109 |
| GO:0035633 | maintenance of blood-brain barrier | 23 | 7 | 1.72 | 0.00109 |
| GO:1901031 | regulation of response to reactive oxygen species | 23 | 7 | 1.72 | 0.00109 |
| GO:2001233 | regulation of apoptotic signaling pathway | 272 | 35 | 20.38 | 0.00113 |
| GO:0030900 | forebrain development | 212 | 29 | 15.88 | 0.00113 |
| GO:0006979 | response to oxidative stress | 334 | 41 | 25.02 | 0.00114 |
| GO:0001701 | in utero embryonic development | 252 | 33 | 18.88 | 0.00114 |
| GO:0042326 | negative regulation of phosphorylation | 303 | 38 | 22.7 | 0.00115 |
| GO:0051047 | positive regulation of secretion | 164 | 24 | 12.29 | 0.00115 |
| GO:0006984 | ER-nucleus signaling pathway | 51 | 11 | 3.82 | 0.00116 |
| GO:0010927 | cellular component assembly involved in morphogenesis | 51 | 11 | 3.82 | 0.00116 |
| GO:0035282 | segmentation | 51 | 11 | 3.82 | 0.00116 |
| GO:0002064 | epithelial cell development | 127 | 20 | 9.52 | 0.00117 |
| GO:0032891 | negative regulation of organic acid transport | 12 | 5 | 0.9 | 0.00118 |
| GO:0040037 | negative regulation of fibroblast growth factor receptor signaling pathway | 12 | 5 | 0.9 | 0.00118 |
| GO:0043501 | skeletal muscle adaptation | 12 | 5 | 0.9 | 0.00118 |
| GO:0060602 | branch elongation of an epithelium | 12 | 5 | 0.9 | 0.00118 |
| GO:0090136 | epithelial cell-cell adhesion | 12 | 5 | 0.9 | 0.00118 |
| GO:1900037 | regulation of cellular response to hypoxia | 12 | 5 | 0.9 | 0.00118 |
| GO:2000725 | regulation of cardiac muscle cell differentiation | 12 | 5 | 0.9 | 0.00118 |
| GO:2001212 | regulation of vasculogenesis | 12 | 5 | 0.9 | 0.00118 |
| GO:0009799 | specification of symmetry | 75 | 14 | 5.62 | 0.0012 |
| GO:0071805 | potassium ion transmembrane transport | 75 | 14 | 5.62 | 0.0012 |
| GO:0007163 | establishment or maintenance of cell polarity | 155 | 23 | 11.61 | 0.0012 |
| GO:0061008 | hepaticobiliary system development | 101 | 17 | 7.57 | 0.00127 |
| GO:0033628 | regulation of cell adhesion mediated by integrin | 30 | 8 | 2.25 | 0.00127 |
| GO:0071604 | transforming growth factor beta production | 30 | 8 | 2.25 | 0.00127 |
| GO:0071241 | cellular response to inorganic substance | 128 | 20 | 9.59 | 0.00129 |
| GO:0048534 | hematopoietic or lymphoid organ development | 619 | 67 | 46.38 | 0.00129 |
| GO:0006813 | potassium ion transport | 84 | 15 | 6.29 | 0.0013 |
| GO:0015908 | fatty acid transport | 84 | 15 | 6.29 | 0.0013 |
| GO:0003073 | regulation of systemic arterial blood pressure | 37 | 9 | 2.77 | 0.00131 |
| GO:0006629 | lipid metabolic process | 917 | 93 | 68.7 | 0.00134 |
| GO:0120036 | plasma membrane bounded cell projection organization | 1047 | 104 | 78.44 | 0.00141 |
| GO:0014902 | myotube differentiation | 60 | 12 | 4.5 | 0.00142 |
| GO:2001234 | negative regulation of apoptotic signaling pathway | 176 | 25 | 13.19 | 0.00142 |
| GO:0031348 | negative regulation of defense response | 120 | 19 | 8.99 | 0.00144 |
| GO:0032613 | interleukin-10 production | 24 | 7 | 1.8 | 0.00144 |
| GO:0002694 | regulation of leukocyte activation | 276 | 35 | 20.68 | 0.00146 |
| GO:0001975 | response to amphetamine | 18 | 6 | 1.35 | 0.00147 |
| GO:0017145 | stem cell division | 18 | 6 | 1.35 | 0.00147 |
| GO:0031128 | developmental induction | 18 | 6 | 1.35 | 0.00147 |
| GO:0000302 | response to reactive oxygen species | 167 | 24 | 12.51 | 0.00149 |
| GO:0001818 | negative regulation of cytokine production | 167 | 24 | 12.51 | 0.00149 |
| GO:1903793 | positive regulation of anion transport | 318 | 39 | 23.83 | 0.00151 |
| GO:0009953 | dorsal/ventral pattern formation | 45 | 10 | 3.37 | 0.00151 |
| GO:0051899 | membrane depolarization | 45 | 10 | 3.37 | 0.00151 |
| GO:0070374 | positive regulation of ERK1 and ERK2 cascade | 94 | 16 | 7.04 | 0.00154 |

| | | | | | |
|---|---|---|---|---|---|
| GO:0046683 | response to organophosphorus | 77 | 14 | 5.77 | 0.00156 |
| GO:0010243 | response to organonitrogen compound | 692 | 73 | 51.85 | 0.0016 |
| GO:0042130 | negative regulation of T cell proliferation | 31 | 8 | 2.32 | 0.0016 |
| GO:0060043 | regulation of cardiac muscle cell proliferation | 31 | 8 | 2.32 | 0.0016 |
| GO:0070613 | regulation of protein processing | 38 | 9 | 2.85 | 0.0016 |
| GO:1902475 | L-alpha-amino acid transmembrane transport | 38 | 9 | 2.85 | 0.0016 |
| GO:0030162 | regulation of proteolysis | 491 | 55 | 36.79 | 0.00161 |
| GO:0006721 | terpenoid metabolic process | 53 | 11 | 3.97 | 0.00162 |
| GO:0021537 | telencephalon development | 149 | 22 | 11.16 | 0.00162 |
| GO:0097530 | granulocyte migration | 61 | 12 | 4.57 | 0.00164 |
| GO:1903426 | regulation of reactive oxygen species biosynthetic process | 61 | 12 | 4.57 | 0.00164 |
| GO:0044344 | cellular response to fibroblast growth factor stimulus | 86 | 15 | 6.44 | 0.00166 |
| GO:0045834 | positive regulation of lipid metabolic process | 86 | 15 | 6.44 | 0.00166 |
| GO:0010755 | regulation of plasminogen activation | 8 | 4 | 0.6 | 0.00171 |
| GO:0035739 | CD4-positive, alpha-beta T cell proliferation | 8 | 4 | 0.6 | 0.00171 |
| GO:0042574 | retinal metabolic process | 8 | 4 | 0.6 | 0.00171 |
| GO:0051953 | negative regulation of amine transport | 8 | 4 | 0.6 | 0.00171 |
| GO:0060713 | labyrinthine layer morphogenesis | 8 | 4 | 0.6 | 0.00171 |
| GO:0072110 | glomerular mesangial cell proliferation | 8 | 4 | 0.6 | 0.00171 |
| GO:0072283 | metanephric renal vesicle morphogenesis | 8 | 4 | 0.6 | 0.00171 |
| GO:1904181 | positive regulation of membrane depolarization | 8 | 4 | 0.6 | 0.00171 |
| GO:2000561 | regulation of CD4-positive, alpha-beta T cell proliferation | 8 | 4 | 0.6 | 0.00171 |
| GO:0009612 | response to mechanical stimulus | 131 | 20 | 9.81 | 0.00172 |
| GO:0044070 | regulation of anion transport | 537 | 59 | 40.23 | 0.00175 |
| GO:0051302 | regulation of cell division | 104 | 17 | 7.79 | 0.00176 |
| GO:0035725 | sodium ion transmembrane transport | 78 | 14 | 5.84 | 0.00177 |
| GO:2000177 | regulation of neural precursor cell proliferation | 46 | 10 | 3.45 | 0.0018 |
| GO:0010566 | regulation of ketone biosynthetic process | 13 | 5 | 0.97 | 0.00181 |
| GO:0010771 | negative regulation of cell morphogenesis involved in differentiation | 13 | 5 | 0.97 | 0.00181 |
| GO:0034114 | regulation of heterotypic cell-cell adhesion | 13 | 5 | 0.97 | 0.00181 |
| GO:0035338 | long-chain fatty-acyl-CoA biosynthetic process | 13 | 5 | 0.97 | 0.00181 |
| GO:0042474 | middle ear morphogenesis | 13 | 5 | 0.97 | 0.00181 |
| GO:0050901 | leukocyte tethering or rolling | 13 | 5 | 0.97 | 0.00181 |
| GO:0060384 | innervation | 13 | 5 | 0.97 | 0.00181 |
| GO:0070633 | transepithelial transport | 13 | 5 | 0.97 | 0.00181 |
| GO:0090025 | regulation of monocyte chemotaxis | 13 | 5 | 0.97 | 0.00181 |
| GO:1900025 | negative regulation of substrate adhesion-dependent cell spreading | 13 | 5 | 0.97 | 0.00181 |
| GO:1905209 | positive regulation of cardiocyte differentiation | 13 | 5 | 0.97 | 0.00181 |
| GO:0045862 | positive regulation of proteolysis | 280 | 35 | 20.98 | 0.00187 |
| GO:0003254 | regulation of membrane depolarization | 25 | 7 | 1.87 | 0.00188 |
| GO:0033555 | multicellular organismal response to stress | 25 | 7 | 1.87 | 0.00188 |
| GO:0045429 | positive regulation of nitric oxide biosynthetic process | 25 | 7 | 1.87 | 0.00188 |
| GO:0140467 | integrated stress response signaling | 25 | 7 | 1.87 | 0.00188 |
| GO:0150104 | transport across blood-brain barrier | 54 | 11 | 4.05 | 0.00189 |
| GO:0048588 | developmental cell growth | 151 | 22 | 11.31 | 0.00193 |
| GO:0072330 | monocarboxylic acid biosynthetic process | 123 | 19 | 9.22 | 0.00193 |
| GO:0071384 | cellular response to corticosteroid stimulus | 39 | 9 | 2.92 | 0.00195 |
| GO:1903317 | regulation of protein maturation | 39 | 9 | 2.92 | 0.00195 |
| GO:0035987 | endodermal cell differentiation | 32 | 8 | 2.4 | 0.002 |
| GO:0046888 | negative regulation of hormone secretion | 32 | 8 | 2.4 | 0.002 |
| GO:0090303 | positive regulation of wound healing | 32 | 8 | 2.4 | 0.002 |
| GO:1902895 | positive regulation of pri-miRNA transcription by RNA polymerase II | 32 | 8 | 2.4 | 0.002 |
| GO:0010769 | regulation of cell morphogenesis involved in differentiation | 79 | 14 | 5.92 | 0.00201 |
| GO:0045446 | endothelial cell differentiation | 79 | 14 | 5.92 | 0.00201 |
| GO:0042572 | retinol metabolic process | 19 | 6 | 1.42 | 0.00202 |
| GO:2000352 | negative regulation of endothelial cell apoptotic process | 19 | 6 | 1.42 | 0.00202 |
| GO:2000403 | positive regulation of lymphocyte migration | 19 | 6 | 1.42 | 0.00202 |
| GO:0007519 | skeletal muscle tissue development | 88 | 15 | 6.59 | 0.00211 |
| GO:0046660 | female sex differentiation | 71 | 13 | 5.32 | 0.00212 |
| GO:0043536 | positive regulation of blood vessel endothelial cell migration | 47 | 10 | 3.52 | 0.00214 |
| GO:0048511 | rhythmic process | 201 | 27 | 15.06 | 0.00215 |
| GO:0043280 | positive regulation of cysteine-type endopeptidase activity involved in apoptotic process | 97 | 16 | 7.27 | 0.00216 |
| GO:0010232 | vascular transport | 55 | 11 | 4.12 | 0.00221 |

| | | | | | |
|---|---|---|---|---|---|
| GO:0043406 | positive regulation of MAP kinase activity | 153 | 22 | 11.46 | 0.00228 |
| GO:0098655 | cation transmembrane transport | 444 | 50 | 33.27 | 0.00232 |
| GO:0050803 | regulation of synapse structure or activity | 125 | 19 | 9.37 | 0.00234 |
| GO:0010596 | negative regulation of endothelial cell migration | 40 | 9 | 3 | 0.00235 |
| GO:0035249 | synaptic transmission, glutamatergic | 40 | 9 | 3 | 0.00235 |
| GO:0070664 | negative regulation of leukocyte proliferation | 40 | 9 | 3 | 0.00235 |
| GO:1903036 | positive regulation of response to wounding | 40 | 9 | 3 | 0.00235 |
| GO:0150115 | cell-substrate junction organization | 89 | 15 | 6.67 | 0.00236 |
| GO:0045601 | regulation of endothelial cell differentiation | 26 | 7 | 1.95 | 0.0024 |
| GO:0150117 | positive regulation of cell-substrate junction organization | 26 | 7 | 1.95 | 0.0024 |
| GO:2000107 | negative regulation of leukocyte apoptotic process | 26 | 7 | 1.95 | 0.0024 |
| GO:0033344 | cholesterol efflux | 33 | 8 | 2.47 | 0.00247 |
| GO:0035337 | fatty-acyl-CoA metabolic process | 33 | 8 | 2.47 | 0.00247 |
| GO:0046622 | positive regulation of organ growth | 33 | 8 | 2.47 | 0.00247 |
| GO:0009064 | glutamine family amino acid metabolic process | 48 | 10 | 3.6 | 0.00253 |
| GO:0032233 | positive regulation of actin filament bundle assembly | 48 | 10 | 3.6 | 0.00253 |
| GO:0046889 | positive regulation of lipid biosynthetic process | 48 | 10 | 3.6 | 0.00253 |
| GO:0031099 | regeneration | 126 | 19 | 9.44 | 0.00257 |
| GO:0033273 | response to vitamin | 56 | 11 | 4.2 | 0.00257 |
| GO:0043407 | negative regulation of MAP kinase activity | 56 | 11 | 4.2 | 0.00257 |
| GO:0051893 | regulation of focal adhesion assembly | 56 | 11 | 4.2 | 0.00257 |
| GO:0090109 | regulation of cell-substrate junction assembly | 56 | 11 | 4.2 | 0.00257 |
| GO:0006066 | alcohol metabolic process | 244 | 31 | 18.28 | 0.00257 |
| GO:0046885 | regulation of hormone biosynthetic process | 14 | 5 | 1.05 | 0.00264 |
| GO:0070831 | basement membrane assembly | 14 | 5 | 1.05 | 0.00264 |
| GO:0048562 | embryonic organ morphogenesis | 155 | 22 | 11.61 | 0.00269 |
| GO:0032350 | regulation of hormone metabolic process | 20 | 6 | 1.5 | 0.0027 |
| GO:0060045 | positive regulation of cardiac muscle cell proliferation | 20 | 6 | 1.5 | 0.0027 |
| GO:0071295 | cellular response to vitamin | 20 | 6 | 1.5 | 0.0027 |
| GO:1905332 | positive regulation of morphogenesis of an epithelium | 20 | 6 | 1.5 | 0.0027 |
| GO:0019221 | cytokine-mediated signaling pathway | 492 | 54 | 36.86 | 0.00277 |
| GO:0006909 | phagocytosis | 175 | 24 | 13.11 | 0.00281 |
| GO:0002526 | acute inflammatory response | 41 | 9 | 3.07 | 0.00282 |
| GO:1902893 | regulation of pri-miRNA transcription by RNA polymerase II | 41 | 9 | 3.07 | 0.00282 |
| GO:0051341 | regulation of oxidoreductase activity | 65 | 12 | 4.87 | 0.00289 |
| GO:0032870 | cellular response to hormone stimulus | 416 | 47 | 31.17 | 0.0029 |
| GO:0002675 | positive regulation of acute inflammatory response | 9 | 4 | 0.67 | 0.0029 |
| GO:0007263 | nitric oxide mediated signal transduction | 9 | 4 | 0.67 | 0.0029 |
| GO:0009125 | nucleoside monophosphate catabolic process | 9 | 4 | 0.67 | 0.0029 |
| GO:0030049 | muscle filament sliding | 9 | 4 | 0.67 | 0.0029 |
| GO:0032026 | response to magnesium ion | 9 | 4 | 0.67 | 0.0029 |
| GO:0033275 | actin-myosin filament sliding | 9 | 4 | 0.67 | 0.0029 |
| GO:0035929 | steroid hormone secretion | 9 | 4 | 0.67 | 0.0029 |
| GO:0042745 | circadian sleep/wake cycle | 9 | 4 | 0.67 | 0.0029 |
| GO:0045986 | negative regulation of smooth muscle contraction | 9 | 4 | 0.67 | 0.0029 |
| GO:0046629 | gamma-delta T cell activation | 9 | 4 | 0.67 | 0.0029 |
| GO:0060347 | heart trabecula formation | 9 | 4 | 0.67 | 0.0029 |
| GO:0061299 | retina vasculature morphogenesis in camera-type eye | 9 | 4 | 0.67 | 0.0029 |
| GO:0072077 | renal vesicle morphogenesis | 9 | 4 | 0.67 | 0.0029 |
| GO:0072160 | nephron tubule epithelial cell differentiation | 9 | 4 | 0.67 | 0.0029 |
| GO:0150172 | regulation of phosphatidylcholine metabolic process | 9 | 4 | 0.67 | 0.0029 |
| GO:2000696 | regulation of epithelial cell differentiation involved in kidney development | 9 | 4 | 0.67 | 0.0029 |
| GO:2001214 | positive regulation of vasculogenesis | 9 | 4 | 0.67 | 0.0029 |
| GO:0051347 | positive regulation of transferase activity | 460 | 51 | 34.46 | 0.00292 |
| GO:0034220 | ion transmembrane transport | 720 | 74 | 53.94 | 0.00296 |
| GO:0003151 | outflow tract morphogenesis | 49 | 10 | 3.67 | 0.00296 |
| GO:0014855 | striated muscle cell proliferation | 49 | 10 | 3.67 | 0.00296 |
| GO:0051249 | regulation of lymphocyte activation | 236 | 30 | 17.68 | 0.00297 |
| GO:0032964 | collagen biosynthetic process | 34 | 8 | 2.55 | 0.00302 |
| GO:0045668 | negative regulation of osteoblast differentiation | 34 | 8 | 2.55 | 0.00302 |
| GO:0050805 | negative regulation of synaptic transmission | 34 | 8 | 2.55 | 0.00302 |
| GO:1990868 | response to chemokine | 34 | 8 | 2.55 | 0.00302 |
| GO:1990869 | cellular response to chemokine | 34 | 8 | 2.55 | 0.00302 |

| | | | | | |
|---|---|---|---|---|---|
| GO:0030030 | cell projection organization | 1072 | 104 | 80.32 | 0.00302 |
| GO:0015804 | neutral amino acid transport | 27 | 7 | 2.02 | 0.00304 |
| GO:0019229 | regulation of vasoconstriction | 27 | 7 | 2.02 | 0.00304 |
| GO:0050795 | regulation of behavior | 27 | 7 | 2.02 | 0.00304 |
| GO:0061098 | positive regulation of protein tyrosine kinase activity | 27 | 7 | 2.02 | 0.00304 |
| GO:0071548 | response to dexamethasone | 27 | 7 | 2.02 | 0.00304 |
| GO:0071902 | positive regulation of protein serine/threonine kinase activity | 216 | 28 | 16.18 | 0.00304 |
| GO:0032269 | negative regulation of cellular protein metabolic process | 698 | 72 | 52.3 | 0.00306 |
| GO:0071356 | cellular response to tumor necrosis factor | 196 | 26 | 14.68 | 0.00307 |
| GO:0009855 | determination of bilateral symmetry | 74 | 13 | 5.54 | 0.0031 |
| GO:1903532 | positive regulation of secretion by cell | 157 | 22 | 11.76 | 0.00316 |
| GO:0050807 | regulation of synapse organization | 119 | 18 | 8.92 | 0.00318 |
| GO:0040008 | regulation of growth | 440 | 49 | 32.97 | 0.0032 |
| GO:0002520 | immune system development | 665 | 69 | 49.82 | 0.00322 |
| GO:1905039 | carboxylic acid transmembrane transport | 92 | 15 | 6.89 | 0.00329 |
| GO:0043583 | ear development | 101 | 16 | 7.57 | 0.00329 |
| GO:1901655 | cellular response to ketone | 66 | 12 | 4.94 | 0.00329 |
| GO:0033077 | T cell differentiation in thymus | 42 | 9 | 3.15 | 0.00335 |
| GO:0061614 | pri-miRNA transcription by RNA polymerase II | 42 | 9 | 3.15 | 0.00335 |
| GO:0030183 | B cell differentiation | 75 | 13 | 5.62 | 0.0035 |
| GO:0001894 | tissue homeostasis | 139 | 20 | 10.41 | 0.00351 |
| GO:0031401 | positive regulation of protein modification process | 702 | 72 | 52.6 | 0.00353 |
| GO:0003209 | cardiac atrium morphogenesis | 21 | 6 | 1.57 | 0.00355 |
| GO:0010614 | negative regulation of cardiac muscle hypertrophy | 21 | 6 | 1.57 | 0.00355 |
| GO:0010863 | positive regulation of phospholipase C activity | 21 | 6 | 1.57 | 0.00355 |
| GO:0061311 | cell surface receptor signaling pathway involved in heart development | 21 | 6 | 1.57 | 0.00355 |
| GO:0070570 | regulation of neuron projection regeneration | 21 | 6 | 1.57 | 0.00355 |
| GO:1903055 | positive regulation of extracellular matrix organization | 21 | 6 | 1.57 | 0.00355 |
| GO:0006865 | amino acid transport | 84 | 14 | 6.29 | 0.00362 |
| GO:0034599 | cellular response to oxidative stress | 229 | 29 | 17.16 | 0.00365 |
| GO:0055085 | transmembrane transport | 854 | 85 | 63.98 | 0.00366 |
| GO:1903825 | organic acid transmembrane transport | 93 | 15 | 6.97 | 0.00366 |
| GO:0032371 | regulation of sterol transport | 35 | 8 | 2.62 | 0.00366 |
| GO:0032374 | regulation of cholesterol transport | 35 | 8 | 2.62 | 0.00366 |
| GO:1903428 | positive regulation of reactive oxygen species biosynthetic process | 35 | 8 | 2.62 | 0.00366 |
| GO:0021983 | pituitary gland development | 15 | 5 | 1.12 | 0.00371 |
| GO:0030220 | platelet formation | 15 | 5 | 1.12 | 0.00371 |
| GO:0032733 | positive regulation of interleukin-10 production | 15 | 5 | 1.12 | 0.00371 |
| GO:0051954 | positive regulation of amine transport | 15 | 5 | 1.12 | 0.00371 |
| GO:0051955 | regulation of amino acid transport | 15 | 5 | 1.12 | 0.00371 |
| GO:0060795 | cell fate commitment involved in formation of primary germ layer | 15 | 5 | 1.12 | 0.00371 |
| GO:0061298 | retina vasculature development in camera-type eye | 15 | 5 | 1.12 | 0.00371 |
| GO:0062009 | secondary palate development | 15 | 5 | 1.12 | 0.00371 |
| GO:0003129 | heart induction | 5 | 3 | 0.37 | 0.00374 |
| GO:0003223 | ventricular compact myocardium morphogenesis | 5 | 3 | 0.37 | 0.00374 |
| GO:0003308 | negative regulation of Wnt signaling pathway involved in heart development | 5 | 3 | 0.37 | 0.00374 |
| GO:0007620 | copulation | 5 | 3 | 0.37 | 0.00374 |
| GO:0009222 | pyrimidine ribonucleotide catabolic process | 5 | 3 | 0.37 | 0.00374 |
| GO:0010616 | negative regulation of cardiac muscle adaptation | 5 | 3 | 0.37 | 0.00374 |
| GO:0010749 | regulation of nitric oxide mediated signal transduction | 5 | 3 | 0.37 | 0.00374 |
| GO:0030322 | stabilization of membrane potential | 5 | 3 | 0.37 | 0.00374 |
| GO:0032914 | positive regulation of transforming growth factor beta1 production | 5 | 3 | 0.37 | 0.00374 |
| GO:0035630 | bone mineralization involved in bone maturation | 5 | 3 | 0.37 | 0.00374 |
| GO:0046886 | positive regulation of hormone biosynthetic process | 5 | 3 | 0.37 | 0.00374 |
| GO:0051917 | regulation of fibrinolysis | 5 | 3 | 0.37 | 0.00374 |
| GO:0060272 | embryonic skeletal joint morphogenesis | 5 | 3 | 0.37 | 0.00374 |
| GO:0060426 | lung vasculature development | 5 | 3 | 0.37 | 0.00374 |
| GO:0060638 | mesenchymal-epithelial cell signaling | 5 | 3 | 0.37 | 0.00374 |
| GO:0060670 | branching involved in labyrinthine layer morphogenesis | 5 | 3 | 0.37 | 0.00374 |
| GO:0061043 | regulation of vascular wound healing | 5 | 3 | 0.37 | 0.00374 |
| GO:0072008 | glomerular mesangial cell differentiation | 5 | 3 | 0.37 | 0.00374 |
| GO:0072103 | glomerulus vasculature morphogenesis | 5 | 3 | 0.37 | 0.00374 |
| GO:0072104 | glomerular capillary formation | 5 | 3 | 0.37 | 0.00374 |

| | | | | | |
|---|---|---|---|---|---|
| GO:0072126 | positive regulation of glomerular mesangial cell proliferation | 5 | 3 | 0.37 | 0.00374 |
| GO:0072143 | mesangial cell development | 5 | 3 | 0.37 | 0.00374 |
| GO:0072161 | mesenchymal cell differentiation involved in kidney development | 5 | 3 | 0.37 | 0.00374 |
| GO:0072172 | mesonephric tubule formation | 5 | 3 | 0.37 | 0.00374 |
| GO:0090027 | negative regulation of monocyte chemotaxis | 5 | 3 | 0.37 | 0.00374 |
| GO:0098713 | leucine import across plasma membrane | 5 | 3 | 0.37 | 0.00374 |
| GO:0099509 | regulation of presynaptic cytosolic calcium ion concentration | 5 | 3 | 0.37 | 0.00374 |
| GO:1903243 | negative regulation of cardiac muscle hypertrophy in response to stress | 5 | 3 | 0.37 | 0.00374 |
| GO:1903801 | L-leucine import across plasma membrane | 5 | 3 | 0.37 | 0.00374 |
| GO:2000121 | regulation of removal of superoxide radicals | 5 | 3 | 0.37 | 0.00374 |
| GO:2000562 | negative regulation of CD4-positive, alpha-beta T cell proliferation | 5 | 3 | 0.37 | 0.00374 |
| GO:2001012 | mesenchymal cell differentiation involved in renal system development | 5 | 3 | 0.37 | 0.00374 |
| GO:0009165 | nucleotide biosynthetic process | 199 | 26 | 14.91 | 0.00377 |
| GO:0010463 | mesenchymal cell proliferation | 28 | 7 | 2.1 | 0.00379 |
| GO:0032720 | negative regulation of tumor necrosis factor production | 28 | 7 | 2.1 | 0.00379 |
| GO:0043537 | negative regulation of blood vessel endothelial cell migration | 28 | 7 | 2.1 | 0.00379 |
| GO:0071901 | negative regulation of protein serine/threonine kinase activity | 112 | 17 | 8.39 | 0.00393 |
| GO:0097306 | cellular response to alcohol | 59 | 11 | 4.42 | 0.00393 |
| GO:1904035 | regulation of epithelial cell apoptotic process | 59 | 11 | 4.42 | 0.00393 |
| GO:0032649 | regulation of interferon-gamma production | 43 | 9 | 3.22 | 0.00396 |
| GO:0000187 | activation of MAPK activity | 103 | 16 | 7.72 | 0.00402 |
| GO:0045666 | positive regulation of neuron differentiation | 51 | 10 | 3.82 | 0.00402 |
| GO:0048678 | response to axon injury | 51 | 10 | 3.82 | 0.00402 |
| GO:0014074 | response to purine-containing compound | 85 | 14 | 6.37 | 0.00404 |
| GO:0032147 | activation of protein kinase activity | 221 | 28 | 16.56 | 0.00421 |
| GO:0008037 | cell recognition | 68 | 12 | 5.09 | 0.00425 |
| GO:0031669 | cellular response to nutrient levels | 161 | 22 | 12.06 | 0.00431 |
| GO:1901293 | nucleoside phosphate biosynthetic process | 201 | 26 | 15.06 | 0.00432 |
| GO:0051098 | regulation of binding | 284 | 34 | 21.28 | 0.00433 |
| GO:0048863 | stem cell differentiation | 181 | 24 | 13.56 | 0.00436 |
| GO:0009225 | nucleotide-sugar metabolic process | 36 | 8 | 2.7 | 0.00441 |
| GO:0032945 | negative regulation of mononuclear cell proliferation | 36 | 8 | 2.7 | 0.00441 |
| GO:0050672 | negative regulation of lymphocyte proliferation | 36 | 8 | 2.7 | 0.00441 |
| GO:0050886 | endocrine process | 36 | 8 | 2.7 | 0.00441 |
| GO:0071385 | cellular response to glucocorticoid stimulus | 36 | 8 | 2.7 | 0.00441 |
| GO:0010721 | negative regulation of cell development | 104 | 16 | 7.79 | 0.00443 |
| GO:0008585 | female gonad development | 60 | 11 | 4.5 | 0.0045 |
| GO:0002031 | G protein-coupled receptor internalization | 10 | 4 | 0.75 | 0.00456 |
| GO:0003128 | heart field specification | 10 | 4 | 0.75 | 0.00456 |
| GO:0009954 | proximal/distal pattern formation | 10 | 4 | 0.75 | 0.00456 |
| GO:0032908 | regulation of transforming growth factor beta1 production | 10 | 4 | 0.75 | 0.00456 |
| GO:0038065 | collagen-activated signaling pathway | 10 | 4 | 0.75 | 0.00456 |
| GO:0070278 | extracellular matrix constituent secretion | 10 | 4 | 0.75 | 0.00456 |
| GO:0072189 | ureter development | 10 | 4 | 0.75 | 0.00456 |
| GO:0048713 | regulation of oligodendrocyte differentiation | 22 | 6 | 1.65 | 0.00457 |
| GO:0051150 | regulation of smooth muscle cell differentiation | 22 | 6 | 1.65 | 0.00457 |
| GO:0051445 | regulation of meiotic cell cycle | 22 | 6 | 1.65 | 0.00457 |
| GO:0060317 | cardiac epithelial to mesenchymal transition | 22 | 6 | 1.65 | 0.00457 |
| GO:0060986 | endocrine hormone secretion | 22 | 6 | 1.65 | 0.00457 |
| GO:0070884 | regulation of calcineurin-NFAT signaling cascade | 22 | 6 | 1.65 | 0.00457 |
| GO:0106056 | regulation of calcineurin-mediated signaling | 22 | 6 | 1.65 | 0.00457 |
| GO:1903115 | regulation of actin filament-based movement | 22 | 6 | 1.65 | 0.00457 |
| GO:0009914 | hormone transport | 162 | 22 | 12.14 | 0.00465 |
| GO:0010633 | negative regulation of epithelial cell migration | 52 | 10 | 3.9 | 0.00466 |
| GO:0001953 | negative regulation of cell-matrix adhesion | 29 | 7 | 2.17 | 0.00468 |
| GO:0031670 | cellular response to nutrient | 29 | 7 | 2.17 | 0.00468 |
| GO:0045197 | establishment or maintenance of epithelial cell apical/basal polarity | 29 | 7 | 2.17 | 0.00468 |
| GO:0070228 | regulation of lymphocyte apoptotic process | 29 | 7 | 2.17 | 0.00468 |
| GO:1902414 | protein localization to cell junction | 69 | 12 | 5.17 | 0.0048 |
| GO:0042542 | response to hydrogen peroxide | 105 | 16 | 7.87 | 0.00487 |
| GO:2001056 | positive regulation of cysteine-type endopeptidase activity | 105 | 16 | 7.87 | 0.00487 |
| GO:0051604 | protein maturation | 203 | 26 | 15.21 | 0.00493 |
| GO:0007204 | positive regulation of cytosolic calcium ion concentration | 124 | 18 | 9.29 | 0.00496 |

| | | | | | |
|---|---|---|---|---|---|
| GO:0048015 | phosphatidylinositol-mediated signaling | 124 | 18 | 9.29 | 0.00496 |
| GO:0060538 | skeletal muscle organ development | 96 | 15 | 7.19 | 0.00497 |
| GO:0001702 | gastrulation with mouth forming second | 16 | 5 | 1.2 | 0.00507 |
| GO:0032331 | negative regulation of chondrocyte differentiation | 16 | 5 | 1.2 | 0.00507 |
| GO:0032689 | negative regulation of interferon-gamma production | 16 | 5 | 1.2 | 0.00507 |
| GO:0032892 | positive regulation of organic acid transport | 16 | 5 | 1.2 | 0.00507 |
| GO:0035137 | hindlimb morphogenesis | 16 | 5 | 1.2 | 0.00507 |
| GO:0036344 | platelet morphogenesis | 16 | 5 | 1.2 | 0.00507 |
| GO:0048745 | smooth muscle tissue development | 16 | 5 | 1.2 | 0.00507 |
| GO:0050433 | regulation of catecholamine secretion | 16 | 5 | 1.2 | 0.00507 |
| GO:0061037 | negative regulation of cartilage development | 16 | 5 | 1.2 | 0.00507 |
| GO:0070528 | protein kinase C signaling | 16 | 5 | 1.2 | 0.00507 |
| GO:0071379 | cellular response to prostaglandin stimulus | 16 | 5 | 1.2 | 0.00507 |
| GO:0090183 | regulation of kidney development | 16 | 5 | 1.2 | 0.00507 |
| GO:0090322 | regulation of superoxide metabolic process | 16 | 5 | 1.2 | 0.00507 |
| GO:0045893 | positive regulation of transcription, DNA-templated | 1139 | 108 | 85.34 | 0.00517 |
| GO:1903508 | positive regulation of nucleic acid-templated transcription | 1139 | 108 | 85.34 | 0.00517 |
| GO:0034612 | response to tumor necrosis factor | 214 | 27 | 16.03 | 0.00518 |
| GO:0031016 | pancreas development | 37 | 8 | 2.77 | 0.00526 |
| GO:0045599 | negative regulation of fat cell differentiation | 37 | 8 | 2.77 | 0.00526 |
| GO:0051353 | positive regulation of oxidoreductase activity | 37 | 8 | 2.77 | 0.00526 |
| GO:1902680 | positive regulation of RNA biosynthetic process | 1140 | 108 | 85.41 | 0.00531 |
| GO:0032612 | interleukin-1 production | 53 | 10 | 3.97 | 0.00536 |
| GO:0006720 | isoprenoid metabolic process | 70 | 12 | 5.24 | 0.00541 |
| GO:0035335 | peptidyl-tyrosine dephosphorylation | 70 | 12 | 5.24 | 0.00541 |
| GO:0008610 | lipid biosynthetic process | 497 | 53 | 37.24 | 0.00544 |
| GO:0080135 | regulation of cellular response to stress | 565 | 59 | 42.33 | 0.00545 |
| GO:0001890 | placenta development | 88 | 14 | 6.59 | 0.00556 |
| GO:0032103 | positive regulation of response to external stimulus | 278 | 33 | 20.83 | 0.00557 |
| GO:0072524 | pyridine-containing compound metabolic process | 30 | 7 | 2.25 | 0.00572 |
| GO:1900744 | regulation of p38MAPK cascade | 30 | 7 | 2.25 | 0.00572 |
| GO:0032467 | positive regulation of cytokinesis | 23 | 6 | 1.72 | 0.0058 |
| GO:0043900 | regulation of multi-organism process | 23 | 6 | 1.72 | 0.0058 |
| GO:0045332 | phospholipid translocation | 23 | 6 | 1.72 | 0.0058 |
| GO:0051937 | catecholamine transport | 23 | 6 | 1.72 | 0.0058 |
| GO:1900274 | regulation of phospholipase C activity | 23 | 6 | 1.72 | 0.0058 |
| GO:1901889 | negative regulation of cell junction assembly | 23 | 6 | 1.72 | 0.0058 |
| GO:0046545 | development of primary female sexual characteristics | 62 | 11 | 4.65 | 0.00582 |
| GO:0050770 | regulation of axonogenesis | 107 | 16 | 8.02 | 0.00587 |
| GO:0050870 | positive regulation of T cell activation | 107 | 16 | 8.02 | 0.00587 |
| GO:0009891 | positive regulation of biosynthetic process | 1399 | 129 | 104.82 | 0.00592 |
| GO:0097305 | response to alcohol | 136 | 19 | 10.19 | 0.00604 |
| GO:0007156 | homophilic cell adhesion via plasma membrane adhesion molecules | 71 | 12 | 5.32 | 0.00608 |
| GO:0048706 | embryonic skeletal system development | 71 | 12 | 5.32 | 0.00608 |
| GO:0062197 | cellular response to chemical stress | 269 | 32 | 20.15 | 0.00609 |
| GO:0097190 | apoptotic signaling pathway | 455 | 49 | 34.09 | 0.00613 |
| GO:0034341 | response to interferon-gamma | 117 | 17 | 8.77 | 0.00617 |
| GO:1903039 | positive regulation of leukocyte cell-cell adhesion | 117 | 17 | 8.77 | 0.00617 |
| GO:1902305 | regulation of sodium ion transmembrane transport | 38 | 8 | 2.85 | 0.00624 |
| GO:1904705 | regulation of vascular associated smooth muscle cell proliferation | 38 | 8 | 2.85 | 0.00624 |
| GO:1990874 | vascular associated smooth muscle cell proliferation | 38 | 8 | 2.85 | 0.00624 |
| GO:0034976 | response to endoplasmic reticulum stress | 259 | 31 | 19.4 | 0.00629 |
| GO:0002065 | columnar/cuboidal epithelial cell differentiation | 46 | 9 | 3.45 | 0.00633 |
| GO:0051963 | regulation of synapse assembly | 46 | 9 | 3.45 | 0.00633 |
| GO:0030301 | cholesterol transport | 63 | 11 | 4.72 | 0.00659 |
| GO:0009636 | response to toxic substance | 147 | 20 | 11.01 | 0.00663 |
| GO:0014065 | phosphatidylinositol 3-kinase signaling | 99 | 15 | 7.42 | 0.00664 |
| GO:0034614 | cellular response to reactive oxygen species | 118 | 17 | 8.84 | 0.00672 |
| GO:0002029 | desensitization of G protein-coupled receptor signaling pathway | 11 | 4 | 0.82 | 0.00674 |
| GO:0003222 | ventricular trabecula myocardium morphogenesis | 11 | 4 | 0.82 | 0.00674 |
| GO:0007187 | G protein-coupled receptor signaling pathway, coupled to cyclic nucleotide second messenger | 11 | 4 | 0.82 | 0.00674 |
| GO:0022401 | negative adaptation of signaling pathway | 11 | 4 | 0.82 | 0.00674 |
| GO:0023058 | adaptation of signaling pathway | 11 | 4 | 0.82 | 0.00674 |

| | | | | | |
|---|---|---|---|---|---|
| GO:0031643 | positive regulation of myelination | 11 | 4 | 0.82 | 0.00674 |
| GO:0032905 | transforming growth factor beta1 production | 11 | 4 | 0.82 | 0.00674 |
| GO:0032941 | secretion by tissue | 11 | 4 | 0.82 | 0.00674 |
| GO:0045176 | apical protein localization | 11 | 4 | 0.82 | 0.00674 |
| GO:0060669 | embryonic placenta morphogenesis | 11 | 4 | 0.82 | 0.00674 |
| GO:1902043 | positive regulation of extrinsic apoptotic signaling pathway via death domain receptors | 11 | 4 | 0.82 | 0.00674 |
| GO:2000651 | positive regulation of sodium ion transmembrane transporter activity | 11 | 4 | 0.82 | 0.00674 |
| GO:0043268 | positive regulation of potassium ion transport | 17 | 5 | 1.27 | 0.00675 |
| GO:0050432 | catecholamine secretion | 17 | 5 | 1.27 | 0.00675 |
| GO:0048017 | inositol lipid-mediated signaling | 128 | 18 | 9.59 | 0.00693 |
| GO:0008202 | steroid metabolic process | 188 | 24 | 14.09 | 0.00702 |
| GO:0009952 | anterior/posterior pattern specification | 109 | 16 | 8.17 | 0.00703 |
| GO:0003014 | renal system process | 55 | 10 | 4.12 | 0.00703 |
| GO:0003333 | amino acid transmembrane transport | 55 | 10 | 4.12 | 0.00703 |
| GO:0014068 | positive regulation of phosphatidylinositol 3-kinase signaling | 55 | 10 | 4.12 | 0.00703 |
| GO:0003139 | secondary heart field specification | 6 | 3 | 0.45 | 0.00706 |
| GO:0003307 | regulation of Wnt signaling pathway involved in heart development | 6 | 3 | 0.45 | 0.00706 |
| GO:0003330 | regulation of extracellular matrix constituent secretion | 6 | 3 | 0.45 | 0.00706 |
| GO:0006591 | ornithine metabolic process | 6 | 3 | 0.45 | 0.00706 |
| GO:0009158 | ribonucleoside monophosphate catabolic process | 6 | 3 | 0.45 | 0.00706 |
| GO:0009159 | deoxyribonucleoside monophosphate catabolic process | 6 | 3 | 0.45 | 0.00706 |
| GO:0010612 | regulation of cardiac muscle adaptation | 6 | 3 | 0.45 | 0.00706 |
| GO:0015820 | leucine transport | 6 | 3 | 0.45 | 0.00706 |
| GO:0032352 | positive regulation of hormone metabolic process | 6 | 3 | 0.45 | 0.00706 |
| GO:0032740 | positive regulation of interleukin-17 production | 6 | 3 | 0.45 | 0.00706 |
| GO:0035696 | monocyte extravasation | 6 | 3 | 0.45 | 0.00706 |
| GO:0035813 | regulation of renal sodium excretion | 6 | 3 | 0.45 | 0.00706 |
| GO:0042492 | gamma-delta T cell differentiation | 6 | 3 | 0.45 | 0.00706 |
| GO:0043129 | surfactant homeostasis | 6 | 3 | 0.45 | 0.00706 |
| GO:0045906 | negative regulation of vasoconstriction | 6 | 3 | 0.45 | 0.00706 |
| GO:0048103 | somatic stem cell division | 6 | 3 | 0.45 | 0.00706 |
| GO:0048617 | embryonic foregut morphogenesis | 6 | 3 | 0.45 | 0.00706 |
| GO:0051901 | positive regulation of mitochondrial depolarization | 6 | 3 | 0.45 | 0.00706 |
| GO:0055012 | ventricular cardiac muscle cell differentiation | 6 | 3 | 0.45 | 0.00706 |
| GO:0060947 | cardiac vascular smooth muscle cell differentiation | 6 | 3 | 0.45 | 0.00706 |
| GO:0061438 | renal system vasculature morphogenesis | 6 | 3 | 0.45 | 0.00706 |
| GO:0061439 | kidney vasculature morphogenesis | 6 | 3 | 0.45 | 0.00706 |
| GO:0070234 | positive regulation of T cell apoptotic process | 6 | 3 | 0.45 | 0.00706 |
| GO:0072079 | nephron tubule formation | 6 | 3 | 0.45 | 0.00706 |
| GO:0097048 | dendritic cell apoptotic process | 6 | 3 | 0.45 | 0.00706 |
| GO:0098856 | intestinal lipid absorption | 6 | 3 | 0.45 | 0.00706 |
| GO:1903242 | regulation of cardiac muscle hypertrophy in response to stress | 6 | 3 | 0.45 | 0.00706 |
| GO:2000668 | regulation of dendritic cell apoptotic process | 6 | 3 | 0.45 | 0.00706 |
| GO:2000826 | regulation of heart morphogenesis | 6 | 3 | 0.45 | 0.00706 |
| GO:0002228 | natural killer cell mediated immunity | 24 | 6 | 1.8 | 0.00725 |
| GO:0003230 | cardiac atrium development | 24 | 6 | 1.8 | 0.00725 |
| GO:0007622 | rhythmic behavior | 24 | 6 | 1.8 | 0.00725 |
| GO:0032232 | negative regulation of actin filament bundle assembly | 24 | 6 | 1.8 | 0.00725 |
| GO:0042267 | natural killer cell mediated cytotoxicity | 24 | 6 | 1.8 | 0.00725 |
| GO:0051155 | positive regulation of striated muscle cell differentiation | 24 | 6 | 1.8 | 0.00725 |
| GO:0140354 | lipid import into cell | 24 | 6 | 1.8 | 0.00725 |
| GO:0032609 | interferon-gamma production | 47 | 9 | 3.52 | 0.00732 |
| GO:0032611 | interleukin-1 beta production | 47 | 9 | 3.52 | 0.00732 |
| GO:0034332 | adherens junction organization | 47 | 9 | 3.52 | 0.00732 |
| GO:0046513 | ceramide biosynthetic process | 47 | 9 | 3.52 | 0.00732 |
| GO:1904377 | positive regulation of protein localization to cell periphery | 47 | 9 | 3.52 | 0.00732 |
| GO:0090199 | regulation of release of cytochrome c from mitochondria | 39 | 8 | 2.92 | 0.00734 |
| GO:1903959 | regulation of anion transmembrane transport | 73 | 12 | 5.47 | 0.00761 |
| GO:0007416 | synapse assembly | 82 | 13 | 6.14 | 0.00762 |
| GO:0014066 | regulation of phosphatidylinositol 3-kinase signaling | 82 | 13 | 6.14 | 0.00762 |
| GO:0050680 | negative regulation of epithelial cell proliferation | 82 | 13 | 6.14 | 0.00762 |
| GO:0006665 | sphingolipid metabolic process | 120 | 17 | 8.99 | 0.00794 |
| GO:0048144 | fibroblast proliferation | 56 | 10 | 4.2 | 0.008 |

| | | | | | |
|---|---|---|---|---|---|
| GO:0048145 | regulation of fibroblast proliferation | 56 | 10 | 4.2 | 0.008 |
| GO:1904062 | regulation of cation transmembrane transport | 170 | 22 | 12.74 | 0.0082 |
| GO:0008154 | actin polymerization or depolymerization | 160 | 21 | 11.99 | 0.00826 |
| GO:1902105 | regulation of leukocyte differentiation | 150 | 20 | 11.24 | 0.00827 |
| GO:0003197 | endocardial cushion development | 32 | 7 | 2.4 | 0.0083 |
| GO:0035088 | establishment or maintenance of apical/basal cell polarity | 32 | 7 | 2.4 | 0.0083 |
| GO:0051489 | regulation of filopodium assembly | 32 | 7 | 2.4 | 0.0083 |
| GO:0051785 | positive regulation of nuclear division | 32 | 7 | 2.4 | 0.0083 |
| GO:0061245 | establishment or maintenance of bipolar cell polarity | 32 | 7 | 2.4 | 0.0083 |
| GO:1900026 | positive regulation of substrate adhesion-dependent cell spreading | 32 | 7 | 2.4 | 0.0083 |
| GO:0030307 | positive regulation of cell growth | 111 | 16 | 8.32 | 0.00836 |
| GO:1903038 | negative regulation of leukocyte cell-cell adhesion | 65 | 11 | 4.87 | 0.00836 |
| GO:0032652 | regulation of interleukin-1 production | 48 | 9 | 3.6 | 0.00843 |
| GO:0007044 | cell-substrate junction assembly | 83 | 13 | 6.22 | 0.00843 |
| GO:0048041 | focal adhesion assembly | 74 | 12 | 5.54 | 0.00848 |
| GO:1905897 | regulation of response to endoplasmic reticulum stress | 74 | 12 | 5.54 | 0.00848 |
| GO:0051051 | negative regulation of transport | 254 | 30 | 19.03 | 0.00855 |
| GO:1901652 | response to peptide | 352 | 39 | 26.37 | 0.00871 |
| GO:0051345 | positive regulation of hydrolase activity | 532 | 55 | 39.86 | 0.00871 |
| GO:0050866 | negative regulation of cell activation | 102 | 15 | 7.64 | 0.00874 |
| GO:0003401 | axis elongation | 18 | 5 | 1.35 | 0.00878 |
| GO:0006670 | sphingosine metabolic process | 18 | 5 | 1.35 | 0.00878 |
| GO:0051882 | mitochondrial depolarization | 18 | 5 | 1.35 | 0.00878 |
| GO:0055093 | response to hyperoxia | 18 | 5 | 1.35 | 0.00878 |
| GO:0071715 | icosanoid transport | 18 | 5 | 1.35 | 0.00878 |
| GO:0150077 | regulation of neuroinflammatory response | 18 | 5 | 1.35 | 0.00878 |
| GO:0150105 | protein localization to cell-cell junction | 18 | 5 | 1.35 | 0.00878 |
| GO:0045055 | regulated exocytosis | 498 | 52 | 37.31 | 0.00879 |
| GO:0030097 | hemopoiesis | 590 | 60 | 44.2 | 0.00888 |
| GO:0030225 | macrophage differentiation | 25 | 6 | 1.87 | 0.00895 |
| GO:0034311 | diol metabolic process | 25 | 6 | 1.87 | 0.00895 |
| GO:0046949 | fatty-acyl-CoA biosynthetic process | 25 | 6 | 1.87 | 0.00895 |
| GO:0061756 | leukocyte adhesion to vascular endothelial cell | 25 | 6 | 1.87 | 0.00895 |
| GO:2000191 | regulation of fatty acid transport | 25 | 6 | 1.87 | 0.00895 |
| GO:0001570 | vasculogenesis | 57 | 10 | 4.27 | 0.00907 |
| GO:0006112 | energy reserve metabolic process | 57 | 10 | 4.27 | 0.00907 |
| GO:1901137 | carbohydrate derivative biosynthetic process | 499 | 52 | 37.39 | 0.00913 |
| GO:0071453 | cellular response to oxygen levels | 182 | 23 | 13.64 | 0.00923 |
| GO:0006898 | receptor-mediated endocytosis | 172 | 22 | 12.89 | 0.00937 |
| GO:0003279 | cardiac septum development | 75 | 12 | 5.62 | 0.00943 |
| GO:0006525 | arginine metabolic process | 12 | 4 | 0.9 | 0.00951 |
| GO:0014046 | dopamine secretion | 12 | 4 | 0.9 | 0.00951 |
| GO:0014059 | regulation of dopamine secretion | 12 | 4 | 0.9 | 0.00951 |
| GO:0030194 | positive regulation of blood coagulation | 12 | 4 | 0.9 | 0.00951 |
| GO:0043116 | negative regulation of vascular permeability | 12 | 4 | 0.9 | 0.00951 |
| GO:0043173 | nucleotide salvage | 12 | 4 | 0.9 | 0.00951 |
| GO:0045198 | establishment of epithelial cell apical/basal polarity | 12 | 4 | 0.9 | 0.00951 |
| GO:0051482 | positive regulation of cytosolic calcium ion concentration involved in phospholipase C-activating G protein-coupled signaling pathway | 12 | 4 | 0.9 | 0.00951 |
| GO:0060037 | pharyngeal system development | 12 | 4 | 0.9 | 0.00951 |
| GO:0070885 | negative regulation of calcineurin-NFAT signaling cascade | 12 | 4 | 0.9 | 0.00951 |
| GO:0106057 | negative regulation of calcineurin-mediated signaling | 12 | 4 | 0.9 | 0.00951 |
| GO:1900048 | positive regulation of hemostasis | 12 | 4 | 0.9 | 0.00951 |
| GO:0035050 | embryonic heart tube development | 49 | 9 | 3.67 | 0.00965 |
| GO:0031324 | negative regulation of cellular metabolic process | 1880 | 166 | 140.85 | 0.0098 |
| GO:0001756 | somitogenesis | 33 | 7 | 2.47 | 0.00987 |
| GO:0034656 | nucleobase-containing small molecule catabolic process | 33 | 7 | 2.47 | 0.00987 |
| GO:0046467 | membrane lipid biosynthetic process | 113 | 16 | 8.47 | 0.00988 |
| GO:0006081 | cellular aldehyde metabolic process | 41 | 8 | 3.07 | 0.00999 |
| GO:0035690 | cellular response to drug | 41 | 8 | 3.07 | 0.00999 |
| GO:0043266 | regulation of potassium ion transport | 41 | 8 | 3.07 | 0.00999 |
| GO:0046847 | filopodium assembly | 41 | 8 | 3.07 | 0.00999 |
| GO:1903078 | positive regulation of protein localization to plasma membrane | 41 | 8 | 3.07 | 0.00999 |
| GO:0007275 | multicellular organism development | 3175 | 387 | 237.88 | 1.80E-30 |

| | | | | | |
|---|---|---|---|---|---|
| GO:0048513 | animal organ development | 2046 | 284 | 153.29 | 1.00E-29 |
| GO:0040011 | locomotion | 1113 | 188 | 83.39 | 5.70E-29 |
| GO:0032502 | developmental process | 3791 | 433 | 284.03 | 1.20E-28 |
| GO:0009888 | tissue development | 1139 | 188 | 85.34 | 1.20E-27 |
| GO:0030154 | cell differentiation | 2403 | 309 | 180.04 | 1.20E-26 |
| GO:0016477 | cell migration | 928 | 162 | 69.53 | 2.40E-26 |
| GO:0051239 | regulation of multicellular organismal process | 1582 | 230 | 118.53 | 4.60E-26 |
| GO:0048870 | cell motility | 1008 | 170 | 75.52 | 6.30E-26 |
| GO:0051674 | localization of cell | 1008 | 170 | 75.52 | 6.30E-26 |
| GO:0048869 | cellular developmental process | 2471 | 313 | 185.13 | 7.00E-26 |
| GO:0003008 | system process | 860 | 153 | 64.43 | 1.00E-25 |
| GO:0006928 | movement of cell or subcellular component | 1285 | 198 | 96.28 | 3.00E-25 |
| GO:0050793 | regulation of developmental process | 1584 | 226 | 118.68 | 2.20E-24 |
| GO:0072359 | circulatory system development | 735 | 134 | 55.07 | 1.90E-23 |
| GO:0042221 | response to chemical | 2626 | 319 | 196.75 | 3.40E-23 |
| GO:0030334 | regulation of cell migration | 600 | 117 | 44.95 | 5.50E-23 |
| GO:0051270 | regulation of cellular component movement | 676 | 126 | 50.65 | 6.40E-23 |
| GO:0048646 | anatomical structure formation involved in morphogenesis | 685 | 127 | 51.32 | 6.90E-23 |
| GO:2000145 | regulation of cell motility | 633 | 120 | 47.43 | 1.80E-22 |
| GO:0023052 | signaling | 3678 | 405 | 275.56 | 2.20E-22 |
| GO:0040012 | regulation of locomotion | 653 | 122 | 48.92 | 2.80E-22 |
| GO:0007154 | cell communication | 3696 | 405 | 276.91 | 6.30E-22 |
| GO:0030198 | extracellular matrix organization | 247 | 67 | 18.51 | 2.40E-21 |
| GO:0043062 | extracellular structure organization | 247 | 67 | 18.51 | 2.40E-21 |
| GO:0045229 | external encapsulating structure organization | 247 | 67 | 18.51 | 2.40E-21 |
| GO:0050896 | response to stimulus | 5241 | 522 | 392.67 | 3.00E-21 |
| GO:0035295 | tube development | 682 | 123 | 51.1 | 4.30E-21 |
| GO:0009605 | response to external stimulus | 1535 | 210 | 115.01 | 3.80E-20 |
| GO:0022603 | regulation of anatomical structure morphogenesis | 676 | 119 | 50.65 | 1.60E-19 |
| GO:0051094 | positive regulation of developmental process | 777 | 130 | 58.21 | 2.40E-19 |
| GO:0070887 | cellular response to chemical stimulus | 2075 | 258 | 155.46 | 2.80E-19 |
| GO:0007165 | signal transduction | 3429 | 374 | 256.91 | 3.70E-19 |
| GO:0003012 | muscle system process | 232 | 61 | 17.38 | 8.70E-19 |
| GO:0098609 | cell-cell adhesion | 466 | 92 | 34.91 | 1.60E-18 |
| GO:0001944 | vasculature development | 497 | 95 | 37.24 | 4.20E-18 |
| GO:2000026 | regulation of multicellular organismal development | 845 | 134 | 63.31 | 7.20E-18 |
| GO:0042127 | regulation of cell population proliferation | 973 | 147 | 72.9 | 1.20E-17 |
| GO:0001568 | blood vessel development | 473 | 91 | 35.44 | 1.50E-17 |
| GO:0035239 | tube morphogenesis | 561 | 101 | 42.03 | 2.60E-17 |
| GO:0051241 | negative regulation of multicellular organismal process | 590 | 104 | 44.2 | 3.80E-17 |
| GO:0010033 | response to organic substance | 2109 | 254 | 158.01 | 4.60E-17 |
| GO:0061061 | muscle structure development | 381 | 78 | 28.55 | 9.50E-17 |
| GO:0030155 | regulation of cell adhesion | 439 | 85 | 32.89 | 1.30E-16 |
| GO:0060429 | epithelium development | 701 | 115 | 52.52 | 1.80E-16 |
| GO:0071310 | cellular response to organic substance | 1726 | 217 | 129.32 | 2.10E-16 |
| GO:0040017 | positive regulation of locomotion | 374 | 76 | 28.02 | 3.70E-16 |
| GO:0048514 | blood vessel morphogenesis | 416 | 81 | 31.17 | 5.00E-16 |
| GO:0030335 | positive regulation of cell migration | 361 | 74 | 27.05 | 5.70E-16 |
| GO:0048468 | cell development | 1236 | 169 | 92.6 | 5.90E-16 |
| GO:0010646 | regulation of cell communication | 2235 | 261 | 167.45 | 8.10E-16 |
| GO:0001525 | angiogenesis | 356 | 73 | 26.67 | 9.00E-16 |
| GO:0051240 | positive regulation of multicellular organismal process | 800 | 124 | 59.94 | 9.40E-16 |
| GO:0030029 | actin filament-based process | 534 | 94 | 40.01 | 1.60E-15 |
| GO:2000147 | positive regulation of cell motility | 368 | 74 | 27.57 | 1.70E-15 |
| GO:0009887 | animal organ morphogenesis | 623 | 104 | 46.68 | 1.80E-15 |
| GO:0048583 | regulation of response to stimulus | 2608 | 292 | 195.4 | 2.00E-15 |
| GO:0045595 | regulation of cell differentiation | 1021 | 146 | 76.5 | 2.10E-15 |
| GO:0008283 | cell population proliferation | 1153 | 159 | 86.39 | 2.50E-15 |
| GO:0032879 | regulation of localization | 1695 | 210 | 126.99 | 4.10E-15 |
| GO:0023051 | regulation of signaling | 2254 | 260 | 168.88 | 4.60E-15 |
| GO:0051272 | positive regulation of cellular component movement | 377 | 74 | 28.25 | 6.50E-15 |
| GO:0010941 | regulation of cell death | 1109 | 153 | 83.09 | 9.40E-15 |
| GO:0060537 | muscle tissue development | 227 | 54 | 17.01 | 9.70E-15 |

| | | | | | |
|---|---|---|---|---|---|
| GO:0042981 | regulation of apoptotic process | 1001 | 142 | 75 | 1.10E-14 |
| GO:0044057 | regulation of system process | 279 | 61 | 20.9 | 1.10E-14 |
| GO:0051716 | cellular response to stimulus | 4461 | 439 | 334.23 | 1.20E-14 |
| GO:0007166 | cell surface receptor signaling pathway | 1836 | 221 | 137.56 | 1.50E-14 |
| GO:0065008 | regulation of biological quality | 2460 | 276 | 184.31 | 1.50E-14 |
| GO:0043067 | regulation of programmed cell death | 1018 | 143 | 76.27 | 2.00E-14 |
| GO:0009611 | response to wounding | 403 | 76 | 30.19 | 2.60E-14 |
| GO:0030855 | epithelial cell differentiation | 347 | 69 | 26 | 2.90E-14 |
| GO:0048585 | negative regulation of response to stimulus | 1118 | 152 | 83.76 | 4.20E-14 |
| GO:0006935 | chemotaxis | 352 | 69 | 26.37 | 6.10E-14 |
| GO:0042330 | taxis | 353 | 69 | 26.45 | 7.00E-14 |
| GO:0030036 | actin cytoskeleton organization | 489 | 85 | 36.64 | 8.80E-14 |
| GO:0006936 | muscle contraction | 169 | 44 | 12.66 | 1.00E-13 |
| GO:0070848 | response to growth factor | 488 | 84 | 36.56 | 2.20E-13 |
| GO:0003013 | circulatory system process | 317 | 63 | 23.75 | 4.20E-13 |
| GO:0000902 | cell morphogenesis | 677 | 104 | 50.72 | 4.90E-13 |
| GO:0014706 | striated muscle tissue development | 212 | 49 | 15.88 | 5.60E-13 |
| GO:0010648 | negative regulation of cell communication | 957 | 132 | 71.7 | 9.80E-13 |
| GO:0023057 | negative regulation of signaling | 958 | 132 | 71.78 | 1.10E-12 |
| GO:0009719 | response to endogenous stimulus | 1043 | 140 | 78.14 | 1.50E-12 |
| GO:0008284 | positive regulation of cell population proliferation | 523 | 86 | 39.18 | 1.50E-12 |
| GO:0071363 | cellular response to growth factor stimulus | 470 | 80 | 35.21 | 1.60E-12 |
| GO:0009966 | regulation of signal transduction | 2039 | 231 | 152.77 | 2.50E-12 |
| GO:0097435 | supramolecular fiber organization | 510 | 84 | 38.21 | 2.60E-12 |
| GO:0048729 | tissue morphogenesis | 425 | 74 | 31.84 | 3.70E-12 |
| GO:0048518 | positive regulation of biological process | 4010 | 393 | 300.44 | 4.20E-12 |
| GO:0090130 | tissue migration | 232 | 50 | 17.38 | 5.30E-12 |
| GO:0031589 | cell-substrate adhesion | 249 | 52 | 18.66 | 7.10E-12 |
| GO:0042060 | wound healing | 331 | 62 | 24.8 | 9.60E-12 |
| GO:0008219 | cell death | 1457 | 176 | 109.16 | 1.40E-11 |
| GO:0007507 | heart development | 385 | 68 | 28.85 | 1.50E-11 |
| GO:0043542 | endothelial cell migration | 172 | 41 | 12.89 | 1.60E-11 |
| GO:0042692 | muscle cell differentiation | 224 | 48 | 16.78 | 1.80E-11 |
| GO:0045597 | positive regulation of cell differentiation | 520 | 83 | 38.96 | 1.90E-11 |
| GO:0048699 | generation of neurons | 936 | 126 | 70.13 | 2.00E-11 |
| GO:0051146 | striated muscle cell differentiation | 159 | 39 | 11.91 | 2.00E-11 |
| GO:0007167 | enzyme linked receptor protein signaling pathway | 699 | 102 | 52.37 | 2.00E-11 |
| GO:0048738 | cardiac muscle tissue development | 126 | 34 | 9.44 | 2.30E-11 |
| GO:0060541 | respiratory system development | 134 | 35 | 10.04 | 3.20E-11 |
| GO:0048598 | embryonic morphogenesis | 358 | 64 | 26.82 | 3.60E-11 |
| GO:0030324 | lung development | 115 | 32 | 8.62 | 3.70E-11 |
| GO:0060548 | negative regulation of cell death | 660 | 97 | 49.45 | 4.50E-11 |
| GO:0008015 | blood circulation | 261 | 52 | 19.55 | 4.60E-11 |
| GO:0090257 | regulation of muscle system process | 129 | 34 | 9.66 | 4.70E-11 |
| GO:0009968 | negative regulation of signal transduction | 909 | 122 | 68.1 | 5.30E-11 |
| GO:0022008 | neurogenesis | 1007 | 131 | 75.45 | 8.10E-11 |
| GO:0012501 | programmed cell death | 1360 | 164 | 101.89 | 9.90E-11 |
| GO:0030323 | respiratory tube development | 119 | 32 | 8.92 | 1.00E-10 |
| GO:0051093 | negative regulation of developmental process | 537 | 83 | 40.23 | 1.00E-10 |
| GO:0007399 | nervous system development | 1459 | 173 | 109.31 | 1.00E-10 |
| GO:0006915 | apoptotic process | 1319 | 160 | 98.82 | 1.10E-10 |
| GO:0006939 | smooth muscle contraction | 50 | 20 | 3.75 | 1.40E-10 |
| GO:0010631 | epithelial cell migration | 229 | 47 | 17.16 | 1.40E-10 |
| GO:0090132 | epithelium migration | 229 | 47 | 17.16 | 1.40E-10 |
| GO:0048523 | negative regulation of cellular process | 3384 | 336 | 253.54 | 1.50E-10 |
| GO:0007015 | actin filament organization | 311 | 57 | 23.3 | 1.70E-10 |
| GO:0072001 | renal system development | 192 | 42 | 14.39 | 1.70E-10 |
| GO:0009790 | embryo development | 667 | 96 | 49.97 | 1.90E-10 |
| GO:0034330 | cell junction organization | 427 | 70 | 31.99 | 2.50E-10 |
| GO:0071495 | cellular response to endogenous stimulus | 890 | 118 | 66.68 | 2.50E-10 |
| GO:0007162 | negative regulation of cell adhesion | 173 | 39 | 12.96 | 3.00E-10 |
| GO:0008285 | negative regulation of cell population proliferation | 438 | 71 | 32.82 | 3.00E-10 |
| GO:0045823 | positive regulation of heart contraction | 18 | 12 | 1.35 | 3.50E-10 |

| | | | | | |
|---|---|---|---|---|---|
| GO:1903524 | positive regulation of blood circulation | 18 | 12 | 1.35 | 3.50E-10 |
| GO:0001655 | urogenital system development | 212 | 44 | 15.88 | 3.80E-10 |
| GO:0030182 | neuron differentiation | 845 | 113 | 63.31 | 3.90E-10 |
| GO:0051271 | negative regulation of cellular component movement | 197 | 42 | 14.76 | 4.00E-10 |
| GO:0043066 | negative regulation of apoptotic process | 581 | 86 | 43.53 | 4.40E-10 |
| GO:0007369 | gastrulation | 119 | 31 | 8.92 | 4.70E-10 |
| GO:0048522 | positive regulation of cellular process | 3730 | 361 | 279.46 | 5.30E-10 |
| GO:0043069 | negative regulation of programmed cell death | 595 | 87 | 44.58 | 6.50E-10 |
| GO:0050789 | regulation of biological process | 7211 | 618 | 540.27 | 7.20E-10 |
| GO:0050794 | regulation of cellular process | 6856 | 593 | 513.67 | 8.40E-10 |
| GO:1901342 | regulation of vasculature development | 187 | 40 | 14.01 | 9.40E-10 |
| GO:0001667 | ameboidal-type cell migration | 308 | 55 | 23.08 | 9.50E-10 |
| GO:0022604 | regulation of cell morphogenesis | 235 | 46 | 17.61 | 1.10E-09 |
| GO:1901700 | response to oxygen-containing compound | 1038 | 130 | 77.77 | 1.20E-09 |
| GO:0006940 | regulation of smooth muscle contraction | 27 | 14 | 2.02 | 1.30E-09 |
| GO:0042476 | odontogenesis | 67 | 22 | 5.02 | 1.40E-09 |
| GO:2000146 | negative regulation of cell motility | 192 | 40 | 14.39 | 2.10E-09 |
| GO:0006950 | response to stress | 2632 | 269 | 197.2 | 2.20E-09 |
| GO:0050900 | leukocyte migration | 224 | 44 | 16.78 | 2.30E-09 |
| GO:0045765 | regulation of angiogenesis | 185 | 39 | 13.86 | 2.40E-09 |
| GO:0060326 | cell chemotaxis | 148 | 34 | 11.09 | 2.50E-09 |
| GO:0001932 | regulation of protein phosphorylation | 793 | 105 | 59.41 | 3.10E-09 |
| GO:0001822 | kidney development | 188 | 39 | 14.09 | 3.90E-09 |
| GO:0019220 | regulation of phosphate metabolic process | 1078 | 132 | 80.77 | 3.90E-09 |
| GO:0051174 | regulation of phosphorus metabolic process | 1078 | 132 | 80.77 | 3.90E-09 |
| GO:0048519 | negative regulation of biological process | 3764 | 359 | 282.01 | 4.40E-09 |
| GO:0072006 | nephron development | 83 | 24 | 6.22 | 4.60E-09 |
| GO:0045766 | positive regulation of angiogenesis | 109 | 28 | 8.17 | 4.60E-09 |
| GO:1904018 | positive regulation of vasculature development | 109 | 28 | 8.17 | 4.60E-09 |
| GO:0007423 | sensory organ development | 287 | 51 | 21.5 | 4.70E-09 |
| GO:0001704 | formation of primary germ layer | 71 | 22 | 5.32 | 4.80E-09 |
| GO:0071772 | response to BMP | 103 | 27 | 7.72 | 5.40E-09 |
| GO:0071773 | cellular response to BMP stimulus | 103 | 27 | 7.72 | 5.40E-09 |
| GO:0035051 | cardiocyte differentiation | 90 | 25 | 6.74 | 5.40E-09 |
| GO:0061448 | connective tissue development | 160 | 35 | 11.99 | 5.70E-09 |
| GO:0030336 | negative regulation of cell migration | 183 | 38 | 13.71 | 5.90E-09 |
| GO:0055001 | muscle cell development | 97 | 26 | 7.27 | 6.20E-09 |
| GO:0000904 | cell morphogenesis involved in differentiation | 480 | 72 | 35.96 | 7.30E-09 |
| GO:0006937 | regulation of muscle contraction | 79 | 23 | 5.92 | 8.30E-09 |
| GO:0002009 | morphogenesis of an epithelium | 371 | 60 | 27.8 | 8.50E-09 |
| GO:1903034 | regulation of response to wounding | 92 | 25 | 6.89 | 8.90E-09 |
| GO:0065007 | biological regulation | 7638 | 642 | 572.26 | 9.50E-09 |
| GO:0032102 | negative regulation of response to external stimulus | 210 | 41 | 15.73 | 9.90E-09 |
| GO:0003018 | vascular process in circulatory system | 141 | 32 | 10.56 | 1.00E-08 |
| GO:0048608 | reproductive structure development | 252 | 46 | 18.88 | 1.10E-08 |
| GO:0010460 | positive regulation of heart rate | 12 | 9 | 0.9 | 1.30E-08 |
| GO:0040013 | negative regulation of locomotion | 212 | 41 | 15.88 | 1.30E-08 |
| GO:0070371 | ERK1 and ERK2 cascade | 166 | 35 | 12.44 | 1.60E-08 |
| GO:0061458 | reproductive system development | 255 | 46 | 19.11 | 1.70E-08 |
| GO:0060047 | heart contraction | 130 | 30 | 9.74 | 1.90E-08 |
| GO:0061045 | negative regulation of wound healing | 37 | 15 | 2.77 | 2.30E-08 |
| GO:0150063 | visual system development | 208 | 40 | 15.58 | 2.30E-08 |
| GO:0033993 | response to lipid | 523 | 75 | 39.18 | 2.50E-08 |
| GO:0043010 | camera-type eye development | 177 | 36 | 13.26 | 2.60E-08 |
| GO:0003015 | heart process | 139 | 31 | 10.41 | 2.60E-08 |
| GO:0048880 | sensory system development | 209 | 40 | 15.66 | 2.70E-08 |
| GO:0060284 | regulation of cell development | 320 | 53 | 23.98 | 2.80E-08 |
| GO:0007267 | cell-cell signaling | 921 | 114 | 69 | 3.00E-08 |
| GO:0030199 | collagen fibril organization | 43 | 16 | 3.22 | 3.30E-08 |
| GO:1901701 | cellular response to oxygen-containing compound | 757 | 98 | 56.72 | 3.50E-08 |
| GO:0006869 | lipid transport | 261 | 46 | 19.55 | 3.50E-08 |
| GO:0061041 | regulation of wound healing | 72 | 21 | 5.39 | 3.60E-08 |
| GO:0030509 | BMP signaling pathway | 98 | 25 | 7.34 | 3.60E-08 |

| | | | | | |
|---|---|---|---|---|---|
| GO:0007517 | muscle organ development | 187 | 37 | 14.01 | 3.60E-08 |
| GO:0043408 | regulation of MAPK cascade | 423 | 64 | 31.69 | 3.70E-08 |
| GO:0031032 | actomyosin structure organization | 134 | 30 | 10.04 | 4.00E-08 |
| GO:0070482 | response to oxygen levels | 297 | 50 | 22.25 | 4.10E-08 |
| GO:0014070 | response to organic cyclic compound | 562 | 78 | 42.11 | 5.50E-08 |
| GO:0034446 | substrate adhesion-dependent cell spreading | 87 | 23 | 6.52 | 6.10E-08 |
| GO:1903522 | regulation of blood circulation | 129 | 29 | 9.66 | 6.20E-08 |
| GO:0001654 | eye development | 207 | 39 | 15.51 | 6.30E-08 |
| GO:0055007 | cardiac muscle cell differentiation | 68 | 20 | 5.09 | 6.50E-08 |
| GO:0001503 | ossification | 267 | 46 | 20 | 7.20E-08 |
| GO:0065009 | regulation of molecular function | 2079 | 215 | 155.76 | 8.10E-08 |
| GO:0042325 | regulation of phosphorylation | 939 | 114 | 70.35 | 8.60E-08 |
| GO:0003007 | heart morphogenesis | 154 | 32 | 11.54 | 9.30E-08 |
| GO:1903035 | negative regulation of response to wounding | 46 | 16 | 3.45 | 1.00E-07 |
| GO:0008016 | regulation of heart contraction | 110 | 26 | 8.24 | 1.00E-07 |
| GO:0032970 | regulation of actin filament-based process | 279 | 47 | 20.9 | 1.00E-07 |
| GO:0060485 | mesenchyme development | 195 | 37 | 14.61 | 1.10E-07 |
| GO:0090287 | regulation of cellular response to growth factor stimulus | 195 | 37 | 14.61 | 1.10E-07 |
| GO:0014897 | striated muscle hypertrophy | 58 | 18 | 4.35 | 1.20E-07 |
| GO:0006468 | protein phosphorylation | 1075 | 126 | 80.54 | 1.20E-07 |
| GO:0072012 | glomerulus vasculature development | 22 | 11 | 1.65 | 1.30E-07 |
| GO:0048584 | positive regulation of response to stimulus | 1341 | 150 | 100.47 | 1.30E-07 |
| GO:0001775 | cell activation | 831 | 103 | 62.26 | 1.40E-07 |
| GO:0010810 | regulation of cell-substrate adhesion | 157 | 32 | 11.76 | 1.50E-07 |
| GO:0061138 | morphogenesis of a branching epithelium | 112 | 26 | 8.39 | 1.50E-07 |
| GO:0014896 | muscle hypertrophy | 59 | 18 | 4.42 | 1.60E-07 |
| GO:0072073 | kidney epithelium development | 78 | 21 | 5.84 | 1.60E-07 |
| GO:0048878 | chemical homeostasis | 637 | 84 | 47.73 | 1.60E-07 |
| GO:0034329 | cell junction assembly | 257 | 44 | 19.26 | 1.70E-07 |
| GO:0061572 | actin filament bundle organization | 120 | 27 | 8.99 | 1.70E-07 |
| GO:0036293 | response to decreased oxygen levels | 275 | 46 | 20.6 | 1.80E-07 |
| GO:0045785 | positive regulation of cell adhesion | 249 | 43 | 18.66 | 1.80E-07 |
| GO:0010647 | positive regulation of cell communication | 1093 | 127 | 81.89 | 1.80E-07 |
| GO:0007178 | transmembrane receptor protein serine/threonine kinase signaling pathway | 241 | 42 | 18.06 | 1.90E-07 |
| GO:0061326 | renal tubule development | 48 | 16 | 3.6 | 2.00E-07 |
| GO:0007186 | G protein-coupled receptor signaling pathway | 330 | 52 | 24.72 | 2.00E-07 |
| GO:0072009 | nephron epithelium development | 60 | 18 | 4.5 | 2.10E-07 |
| GO:0009725 | response to hormone | 581 | 78 | 43.53 | 2.20E-07 |
| GO:0061437 | renal system vasculature development | 23 | 11 | 1.72 | 2.30E-07 |
| GO:0061440 | kidney vasculature development | 23 | 11 | 1.72 | 2.30E-07 |
| GO:0022407 | regulation of cell-cell adhesion | 234 | 41 | 17.53 | 2.30E-07 |
| GO:0032101 | regulation of response to external stimulus | 583 | 78 | 43.68 | 2.60E-07 |
| GO:0060419 | heart growth | 67 | 19 | 5.02 | 2.60E-07 |
| GO:0055006 | cardiac cell development | 49 | 16 | 3.67 | 2.70E-07 |
| GO:0120254 | olefinic compound metabolic process | 55 | 17 | 4.12 | 2.80E-07 |
| GO:0098742 | cell-cell adhesion via plasma-membrane adhesion molecules | 123 | 27 | 9.22 | 2.90E-07 |
| GO:0035556 | intracellular signal transduction | 1822 | 190 | 136.51 | 3.50E-07 |
| GO:0001656 | metanephros development | 50 | 16 | 3.75 | 3.70E-07 |
| GO:0003206 | cardiac chamber morphogenesis | 75 | 20 | 5.62 | 3.80E-07 |
| GO:0043409 | negative regulation of MAPK cascade | 125 | 27 | 9.37 | 4.10E-07 |
| GO:0043534 | blood vessel endothelial cell migration | 96 | 23 | 7.19 | 4.30E-07 |
| GO:0010594 | regulation of endothelial cell migration | 133 | 28 | 9.96 | 4.40E-07 |
| GO:0032989 | cellular component morphogenesis | 491 | 68 | 36.79 | 4.50E-07 |
| GO:0034097 | response to cytokine | 766 | 95 | 57.39 | 4.50E-07 |
| GO:0051017 | actin filament bundle assembly | 118 | 26 | 8.84 | 4.50E-07 |
| GO:0042592 | homeostatic process | 1156 | 131 | 86.61 | 4.70E-07 |
| GO:0023056 | positive regulation of signaling | 1101 | 126 | 82.49 | 4.70E-07 |
| GO:0055017 | cardiac muscle tissue growth | 63 | 18 | 4.72 | 4.80E-07 |
| GO:0048666 | neuron development | 715 | 90 | 53.57 | 4.80E-07 |
| GO:1902531 | regulation of intracellular signal transduction | 1224 | 137 | 91.71 | 5.00E-07 |
| GO:0003300 | cardiac muscle hypertrophy | 57 | 17 | 4.27 | 5.00E-07 |
| GO:0045933 | positive regulation of muscle contraction | 16 | 9 | 1.2 | 5.00E-07 |
| GO:0006952 | defense response | 853 | 103 | 63.91 | 5.10E-07 |

| | | | | | |
|---|---|---|---|---|---|
| GO:0000165 | MAPK cascade | 563 | 75 | 42.18 | 5.30E-07 |
| GO:0050673 | epithelial cell proliferation | 250 | 42 | 18.73 | 5.40E-07 |
| GO:0050877 | nervous system process | 416 | 60 | 31.17 | 5.50E-07 |
| GO:0010876 | lipid localization | 295 | 47 | 22.1 | 5.70E-07 |
| GO:0002685 | regulation of leukocyte migration | 112 | 25 | 8.39 | 5.80E-07 |
| GO:0061333 | renal tubule morphogenesis | 40 | 14 | 3 | 5.80E-07 |
| GO:0001935 | endothelial cell proliferation | 105 | 24 | 7.87 | 6.10E-07 |
| GO:0001763 | morphogenesis of a branching structure | 120 | 26 | 8.99 | 6.40E-07 |
| GO:0043500 | muscle adaptation | 71 | 19 | 5.32 | 7.00E-07 |
| GO:0050878 | regulation of body fluid levels | 280 | 45 | 20.98 | 7.70E-07 |
| GO:0045667 | regulation of osteoblast differentiation | 85 | 21 | 6.37 | 7.90E-07 |
| GO:0050920 | regulation of chemotaxis | 129 | 27 | 9.66 | 8.00E-07 |
| GO:0042493 | response to drug | 219 | 38 | 16.41 | 8.20E-07 |
| GO:0003002 | regionalization | 177 | 33 | 13.26 | 8.30E-07 |
| GO:0048762 | mesenchymal cell differentiation | 161 | 31 | 12.06 | 8.70E-07 |
| GO:0070372 | regulation of ERK1 and ERK2 cascade | 153 | 30 | 11.46 | 8.70E-07 |
| GO:0072080 | nephron tubule development | 47 | 15 | 3.52 | 9.00E-07 |
| GO:0046209 | nitric oxide metabolic process | 53 | 16 | 3.97 | 9.10E-07 |
| GO:0010035 | response to inorganic substance | 346 | 52 | 25.92 | 9.10E-07 |
| GO:0048568 | embryonic organ development | 255 | 42 | 19.11 | 9.30E-07 |
| GO:0001666 | response to hypoxia | 264 | 43 | 19.78 | 9.40E-07 |
| GO:0007610 | behavior | 282 | 45 | 21.13 | 9.50E-07 |
| GO:0045596 | negative regulation of cell differentiation | 394 | 57 | 29.52 | 9.70E-07 |
| GO:0002376 | immune system process | 1790 | 185 | 134.11 | 9.70E-07 |
| GO:0050678 | regulation of epithelial cell proliferation | 212 | 37 | 15.88 | 9.90E-07 |
| GO:0030195 | negative regulation of blood coagulation | 26 | 11 | 1.95 | 1.10E-06 |
| GO:1900047 | negative regulation of hemostasis | 26 | 11 | 1.95 | 1.10E-06 |
| GO:0072507 | divalent inorganic cation homeostasis | 230 | 39 | 17.23 | 1.10E-06 |
| GO:0006954 | inflammatory response | 367 | 54 | 27.5 | 1.10E-06 |
| GO:0008360 | regulation of cell shape | 116 | 25 | 8.69 | 1.20E-06 |
| GO:0051960 | regulation of nervous system development | 266 | 43 | 19.93 | 1.20E-06 |
| GO:0007596 | blood coagulation | 205 | 36 | 15.36 | 1.20E-06 |
| GO:0034109 | homotypic cell-cell adhesion | 54 | 16 | 4.05 | 1.20E-06 |
| GO:2001057 | reactive nitrogen species metabolic process | 54 | 16 | 4.05 | 1.20E-06 |
| GO:0140352 | export from cell | 847 | 101 | 63.46 | 1.20E-06 |
| GO:0009967 | positive regulation of signal transduction | 1010 | 116 | 75.67 | 1.20E-06 |
| GO:0050817 | coagulation | 206 | 36 | 15.43 | 1.30E-06 |
| GO:0055074 | calcium ion homeostasis | 206 | 36 | 15.43 | 1.30E-06 |
| GO:0050801 | ion homeostasis | 408 | 58 | 30.57 | 1.40E-06 |
| GO:0031214 | biomineral tissue development | 88 | 21 | 6.59 | 1.50E-06 |
| GO:0007160 | cell-matrix adhesion | 157 | 30 | 11.76 | 1.50E-06 |
| GO:0080134 | regulation of response to stress | 949 | 110 | 71.1 | 1.50E-06 |
| GO:0001823 | mesonephros development | 55 | 16 | 4.12 | 1.60E-06 |
| GO:0032835 | glomerulus development | 43 | 14 | 3.22 | 1.60E-06 |
| GO:0007599 | hemostasis | 208 | 36 | 15.58 | 1.70E-06 |
| GO:0110148 | biomineralization | 89 | 21 | 6.67 | 1.80E-06 |
| GO:0050679 | positive regulation of epithelial cell proliferation | 119 | 25 | 8.92 | 1.90E-06 |
| GO:0072111 | cell proliferation involved in kidney development | 14 | 8 | 1.05 | 1.90E-06 |
| GO:0045859 | regulation of protein kinase activity | 552 | 72 | 41.36 | 2.00E-06 |
| GO:0008544 | epidermis development | 159 | 30 | 11.91 | 2.00E-06 |
| GO:0046903 | secretion | 879 | 103 | 65.86 | 2.10E-06 |
| GO:0002040 | sprouting angiogenesis | 90 | 21 | 6.74 | 2.10E-06 |
| GO:0055013 | cardiac muscle cell development | 44 | 14 | 3.3 | 2.20E-06 |
| GO:0030856 | regulation of epithelial cell differentiation | 83 | 20 | 6.22 | 2.20E-06 |
| GO:0032940 | secretion by cell | 826 | 98 | 61.89 | 2.20E-06 |
| GO:0015849 | organic acid transport | 168 | 31 | 12.59 | 2.20E-06 |
| GO:0006874 | cellular calcium ion homeostasis | 202 | 35 | 15.13 | 2.30E-06 |
| GO:0071675 | regulation of mononuclear cell migration | 63 | 17 | 4.72 | 2.40E-06 |
| GO:0010632 | regulation of epithelial cell migration | 177 | 32 | 13.26 | 2.40E-06 |
| GO:0043086 | negative regulation of catalytic activity | 525 | 69 | 39.33 | 2.50E-06 |
| GO:0090092 | regulation of transmembrane receptor protein serine/threonine kinase signaling pathway | 169 | 31 | 12.66 | 2.60E-06 |
| GO:0050819 | negative regulation of coagulation | 28 | 11 | 2.1 | 2.60E-06 |
| GO:0010628 | positive regulation of gene expression | 680 | 84 | 50.95 | 2.60E-06 |

| | | | | | |
|---|---|---|---|---|---|
| GO:0042110 | T cell activation | 247 | 40 | 18.51 | 2.60E-06 |
| GO:0050921 | positive regulation of chemotaxis | 77 | 19 | 5.77 | 2.70E-06 |
| GO:0072088 | nephron epithelium morphogenesis | 39 | 13 | 2.92 | 2.80E-06 |
| GO:0042730 | fibrinolysis | 11 | 7 | 0.82 | 3.30E-06 |
| GO:0032368 | regulation of lipid transport | 78 | 19 | 5.84 | 3.30E-06 |
| GO:0051216 | cartilage development | 115 | 24 | 8.62 | 3.40E-06 |
| GO:0071345 | cellular response to cytokine stimulus | 706 | 86 | 52.9 | 3.40E-06 |
| GO:0001501 | skeletal system development | 305 | 46 | 22.85 | 3.50E-06 |
| GO:0055065 | metal ion homeostasis | 324 | 48 | 24.27 | 3.60E-06 |
| GO:0001936 | regulation of endothelial cell proliferation | 93 | 21 | 6.97 | 3.80E-06 |
| GO:0072028 | nephron morphogenesis | 40 | 13 | 3 | 3.80E-06 |
| GO:0071636 | positive regulation of transforming growth factor beta production | 15 | 8 | 1.12 | 3.80E-06 |
| GO:0072202 | cell differentiation involved in metanephros development | 15 | 8 | 1.12 | 3.80E-06 |
| GO:0046649 | lymphocyte activation | 373 | 53 | 27.95 | 4.00E-06 |
| GO:0042573 | retinoic acid metabolic process | 8 | 6 | 0.6 | 4.30E-06 |
| GO:0043549 | regulation of kinase activity | 626 | 78 | 46.9 | 4.40E-06 |
| GO:0034754 | cellular hormone metabolic process | 59 | 16 | 4.42 | 4.40E-06 |
| GO:0031175 | neuron projection development | 647 | 80 | 48.47 | 4.40E-06 |
| GO:0009617 | response to bacterium | 289 | 44 | 21.65 | 4.50E-06 |
| GO:0048754 | branching morphogenesis of an epithelial tube | 94 | 21 | 7.04 | 4.50E-06 |
| GO:0006793 | phosphorus metabolic process | 2143 | 211 | 160.56 | 4.70E-06 |
| GO:0072132 | mesenchyme morphogenesis | 35 | 12 | 2.62 | 4.80E-06 |
| GO:0001657 | ureteric bud development | 53 | 15 | 3.97 | 4.90E-06 |
| GO:0010812 | negative regulation of cell-substrate adhesion | 47 | 14 | 3.52 | 5.20E-06 |
| GO:0043491 | protein kinase B signaling | 150 | 28 | 11.24 | 5.50E-06 |
| GO:0010595 | positive regulation of endothelial cell migration | 88 | 20 | 6.59 | 5.70E-06 |
| GO:0061564 | axon development | 330 | 48 | 24.72 | 6.10E-06 |
| GO:0048871 | multicellular organismal homeostasis | 283 | 43 | 21.2 | 6.10E-06 |
| GO:0010573 | vascular endothelial growth factor production | 25 | 10 | 1.87 | 6.10E-06 |
| GO:0014888 | striated muscle adaptation | 25 | 10 | 1.87 | 6.10E-06 |
| GO:0050727 | regulation of inflammatory response | 176 | 31 | 13.19 | 6.10E-06 |
| GO:0072163 | mesonephric epithelium development | 54 | 15 | 4.05 | 6.30E-06 |
| GO:0072164 | mesonephric tubule development | 54 | 15 | 4.05 | 6.30E-06 |
| GO:0048589 | developmental growth | 408 | 56 | 30.57 | 6.30E-06 |
| GO:0032956 | regulation of actin cytoskeleton organization | 256 | 40 | 19.18 | 6.40E-06 |
| GO:0098657 | import into cell | 119 | 24 | 8.92 | 6.40E-06 |
| GO:1902532 | negative regulation of intracellular signal transduction | 379 | 53 | 28.4 | 6.50E-06 |
| GO:0010817 | regulation of hormone levels | 247 | 39 | 18.51 | 6.50E-06 |
| GO:0045216 | cell-cell junction organization | 127 | 25 | 9.52 | 6.50E-06 |
| GO:0007169 | transmembrane receptor protein tyrosine kinase signaling pathway | 479 | 63 | 35.89 | 6.80E-06 |
| GO:0031960 | response to corticosteroid | 89 | 20 | 6.67 | 6.90E-06 |
| GO:0032526 | response to retinoic acid | 61 | 16 | 4.57 | 7.00E-06 |
| GO:0072503 | cellular divalent inorganic cation homeostasis | 221 | 36 | 16.56 | 7.20E-06 |
| GO:0044706 | multi-multicellular organism process | 112 | 23 | 8.39 | 7.30E-06 |
| GO:0001938 | positive regulation of endothelial cell proliferation | 68 | 17 | 5.09 | 7.40E-06 |
| GO:0007268 | chemical synaptic transmission | 323 | 47 | 24.2 | 7.50E-06 |
| GO:0098916 | anterograde trans-synaptic signaling | 323 | 47 | 24.2 | 7.50E-06 |
| GO:2000377 | regulation of reactive oxygen species metabolic process | 128 | 25 | 9.59 | 7.60E-06 |
| GO:0019722 | calcium-mediated signaling | 97 | 21 | 7.27 | 7.60E-06 |
| GO:0048638 | regulation of developmental growth | 204 | 34 | 15.28 | 7.80E-06 |
| GO:0044092 | negative regulation of molecular function | 742 | 88 | 55.59 | 7.80E-06 |
| GO:0007010 | cytoskeleton organization | 972 | 109 | 72.82 | 8.10E-06 |
| GO:0019932 | second-messenger-mediated signaling | 145 | 27 | 10.86 | 8.40E-06 |
| GO:0009913 | epidermal cell differentiation | 98 | 21 | 7.34 | 9.00E-06 |
| GO:0120039 | plasma membrane bounded cell projection morphogenesis | 433 | 58 | 32.44 | 9.20E-06 |
| GO:0001707 | mesoderm formation | 37 | 12 | 2.77 | 9.30E-06 |
| GO:0048644 | muscle organ morphogenesis | 37 | 12 | 2.77 | 9.30E-06 |
| GO:0072078 | nephron tubule morphogenesis | 37 | 12 | 2.77 | 9.30E-06 |
| GO:0022414 | reproductive process | 757 | 89 | 56.72 | 9.80E-06 |
| GO:0033002 | muscle cell proliferation | 130 | 25 | 9.74 | 1.00E-05 |
| GO:0000003 | reproduction | 758 | 89 | 56.79 | 1.00E-05 |
| GO:0001649 | osteoblast differentiation | 155 | 28 | 11.61 | 1.00E-05 |
| GO:0030048 | actin filament-based movement | 70 | 17 | 5.24 | 1.10E-05 |

| | | | | | |
|---|---|---|---|---|---|
| GO:0043065 | positive regulation of apoptotic process | 406 | 55 | 30.42 | 1.10E-05 |
| GO:0048858 | cell projection morphogenesis | 436 | 58 | 32.67 | 1.10E-05 |
| GO:0006809 | nitric oxide biosynthetic process | 50 | 14 | 3.75 | 1.20E-05 |
| GO:0060993 | kidney morphogenesis | 50 | 14 | 3.75 | 1.20E-05 |
| GO:0034113 | heterotypic cell-cell adhesion | 32 | 11 | 2.4 | 1.20E-05 |
| GO:0071677 | positive regulation of mononuclear cell migration | 32 | 11 | 2.4 | 1.20E-05 |
| GO:0110053 | regulation of actin filament organization | 208 | 34 | 15.58 | 1.20E-05 |
| GO:0007389 | pattern specification process | 235 | 37 | 17.61 | 1.20E-05 |
| GO:0019371 | cyclooxygenase pathway | 9 | 6 | 0.67 | 1.20E-05 |
| GO:0099537 | trans-synaptic signaling | 329 | 47 | 24.65 | 1.20E-05 |
| GO:0097529 | myeloid leukocyte migration | 100 | 21 | 7.49 | 1.30E-05 |
| GO:0030193 | regulation of blood coagulation | 38 | 12 | 2.85 | 1.30E-05 |
| GO:1900046 | regulation of hemostasis | 38 | 12 | 2.85 | 1.30E-05 |
| GO:0110110 | positive regulation of animal organ morphogenesis | 17 | 8 | 1.27 | 1.30E-05 |
| GO:0002688 | regulation of leukocyte chemotaxis | 64 | 16 | 4.8 | 1.40E-05 |
| GO:0042445 | hormone metabolic process | 93 | 20 | 6.97 | 1.40E-05 |
| GO:0006875 | cellular metal ion homeostasis | 283 | 42 | 21.2 | 1.40E-05 |
| GO:0010562 | positive regulation of phosphorus metabolic process | 604 | 74 | 45.25 | 1.40E-05 |
| GO:0045937 | positive regulation of phosphate metabolic process | 604 | 74 | 45.25 | 1.40E-05 |
| GO:0010574 | regulation of vascular endothelial growth factor production | 22 | 9 | 1.65 | 1.50E-05 |
| GO:0019369 | arachidonic acid metabolic process | 22 | 9 | 1.65 | 1.50E-05 |
| GO:0072210 | metanephric nephron development | 22 | 9 | 1.65 | 1.50E-05 |
| GO:0001934 | positive regulation of protein phosphorylation | 511 | 65 | 38.29 | 1.50E-05 |
| GO:0030595 | leukocyte chemotaxis | 101 | 21 | 7.57 | 1.50E-05 |
| GO:0010954 | positive regulation of protein processing | 13 | 7 | 0.97 | 1.50E-05 |
| GO:0072109 | glomerular mesangium development | 13 | 7 | 0.97 | 1.50E-05 |
| GO:0071674 | mononuclear cell migration | 86 | 19 | 6.44 | 1.50E-05 |
| GO:0010720 | positive regulation of cell development | 202 | 33 | 15.13 | 1.60E-05 |
| GO:0071300 | cellular response to retinoic acid | 33 | 11 | 2.47 | 1.60E-05 |
| GO:0002042 | cell migration involved in sprouting angiogenesis | 45 | 13 | 3.37 | 1.60E-05 |
| GO:0071900 | regulation of protein serine/threonine kinase activity | 352 | 49 | 26.37 | 1.60E-05 |
| GO:0060021 | roof of mouth development | 65 | 16 | 4.87 | 1.70E-05 |
| GO:0048332 | mesoderm morphogenesis | 39 | 12 | 2.92 | 1.70E-05 |
| GO:0061005 | cell differentiation involved in kidney development | 39 | 12 | 2.92 | 1.70E-05 |
| GO:0140353 | lipid export from cell | 39 | 12 | 2.92 | 1.70E-05 |
| GO:0043068 | positive regulation of programmed cell death | 412 | 55 | 30.87 | 1.70E-05 |
| GO:0010634 | positive regulation of epithelial cell migration | 118 | 23 | 8.84 | 1.80E-05 |
| GO:0099536 | synaptic signaling | 344 | 48 | 25.77 | 1.90E-05 |
| GO:0010942 | positive regulation of cell death | 454 | 59 | 34.01 | 1.90E-05 |
| GO:0050790 | regulation of catalytic activity | 1684 | 169 | 126.17 | 2.00E-05 |
| GO:0050808 | synapse organization | 231 | 36 | 17.31 | 2.00E-05 |
| GO:0050804 | modulation of chemical synaptic transmission | 222 | 35 | 16.63 | 2.00E-05 |
| GO:0035150 | regulation of tube size | 66 | 16 | 4.94 | 2.10E-05 |
| GO:0035296 | regulation of tube diameter | 66 | 16 | 4.94 | 2.10E-05 |
| GO:0097746 | blood vessel diameter maintenance | 66 | 16 | 4.94 | 2.10E-05 |
| GO:0006796 | phosphate-containing compound metabolic process | 2118 | 205 | 158.69 | 2.10E-05 |
| GO:0030098 | lymphocyte differentiation | 187 | 31 | 14.01 | 2.20E-05 |
| GO:0099177 | regulation of trans-synaptic signaling | 223 | 35 | 16.71 | 2.20E-05 |
| GO:0060562 | epithelial tube morphogenesis | 205 | 33 | 15.36 | 2.20E-05 |
| GO:0072171 | mesonephric tubule morphogenesis | 34 | 11 | 2.55 | 2.30E-05 |
| GO:0010718 | positive regulation of epithelial to mesenchymal transition | 40 | 12 | 3 | 2.30E-05 |
| GO:0003205 | cardiac chamber development | 104 | 21 | 7.79 | 2.40E-05 |
| GO:0014909 | smooth muscle cell migration | 53 | 14 | 3.97 | 2.40E-05 |
| GO:1903131 | mononuclear cell differentiation | 215 | 34 | 16.11 | 2.40E-05 |
| GO:0033559 | unsaturated fatty acid metabolic process | 60 | 15 | 4.5 | 2.50E-05 |
| GO:0030282 | bone mineralization | 67 | 16 | 5.02 | 2.50E-05 |
| GO:0055080 | cation homeostasis | 378 | 51 | 28.32 | 2.60E-05 |
| GO:0032370 | positive regulation of lipid transport | 47 | 13 | 3.52 | 2.70E-05 |
| GO:1900024 | regulation of substrate adhesion-dependent cell spreading | 47 | 13 | 3.52 | 2.70E-05 |
| GO:1903319 | positive regulation of protein maturation | 14 | 7 | 1.05 | 2.80E-05 |
| GO:0001816 | cytokine production | 440 | 57 | 32.97 | 2.90E-05 |
| GO:0071396 | cellular response to lipid | 350 | 48 | 26.22 | 3.00E-05 |
| GO:0009991 | response to extracellular stimulus | 311 | 44 | 23.3 | 3.00E-05 |

| | | | | | |
|---|---|---|---|---|---|
| GO:0050818 | regulation of coagulation | 41 | 12 | 3.07 | 3.00E-05 |
| GO:0061035 | regulation of cartilage development | 41 | 12 | 3.07 | 3.00E-05 |
| GO:0080164 | regulation of nitric oxide metabolic process | 41 | 12 | 3.07 | 3.00E-05 |
| GO:0055002 | striated muscle cell development | 54 | 14 | 4.05 | 3.00E-05 |
| GO:0051896 | regulation of protein kinase B signaling | 130 | 24 | 9.74 | 3.10E-05 |
| GO:0060415 | muscle tissue morphogenesis | 35 | 11 | 2.62 | 3.10E-05 |
| GO:0085029 | extracellular matrix assembly | 35 | 11 | 2.62 | 3.10E-05 |
| GO:0032990 | cell part morphogenesis | 451 | 58 | 33.79 | 3.10E-05 |
| GO:0090288 | negative regulation of cellular response to growth factor stimulus | 61 | 15 | 4.57 | 3.10E-05 |
| GO:0015850 | organic hydroxy compound transport | 122 | 23 | 9.14 | 3.10E-05 |
| GO:0002576 | platelet degranulation | 83 | 18 | 6.22 | 3.30E-05 |
| GO:0008217 | regulation of blood pressure | 83 | 18 | 6.22 | 3.30E-05 |
| GO:0030501 | positive regulation of bone mineralization | 24 | 9 | 1.8 | 3.30E-05 |
| GO:1902533 | positive regulation of intracellular signal transduction | 641 | 76 | 48.03 | 3.40E-05 |
| GO:0010575 | positive regulation of vascular endothelial growth factor production | 19 | 8 | 1.42 | 3.40E-05 |
| GO:0060055 | angiogenesis involved in wound healing | 19 | 8 | 1.42 | 3.40E-05 |
| GO:0055082 | cellular chemical homeostasis | 433 | 56 | 32.44 | 3.60E-05 |
| GO:1905952 | regulation of lipid localization | 99 | 20 | 7.42 | 3.60E-05 |
| GO:0051179 | localization | 4321 | 378 | 323.74 | 3.70E-05 |
| GO:0098771 | inorganic ion homeostasis | 383 | 51 | 28.7 | 3.70E-05 |
| GO:0072593 | reactive oxygen species metabolic process | 175 | 29 | 13.11 | 4.00E-05 |
| GO:0001658 | branching involved in ureteric bud morphogenesis | 30 | 10 | 2.25 | 4.00E-05 |
| GO:0045778 | positive regulation of ossification | 30 | 10 | 2.25 | 4.00E-05 |
| GO:0055008 | cardiac muscle tissue morphogenesis | 30 | 10 | 2.25 | 4.00E-05 |
| GO:0051338 | regulation of transferase activity | 720 | 83 | 53.94 | 4.10E-05 |
| GO:0070663 | regulation of leukocyte proliferation | 116 | 22 | 8.69 | 4.20E-05 |
| GO:0051048 | negative regulation of secretion | 77 | 17 | 5.77 | 4.20E-05 |
| GO:0022409 | positive regulation of cell-cell adhesion | 141 | 25 | 10.56 | 4.30E-05 |
| GO:0050767 | regulation of neurogenesis | 230 | 35 | 17.23 | 4.30E-05 |
| GO:0014821 | phasic smooth muscle contraction | 7 | 5 | 0.52 | 4.30E-05 |
| GO:0042481 | regulation of odontogenesis | 7 | 5 | 0.52 | 4.30E-05 |
| GO:0072239 | metanephric glomerulus vasculature development | 7 | 5 | 0.52 | 4.30E-05 |
| GO:0009607 | response to biotic stimulus | 820 | 92 | 61.44 | 4.40E-05 |
| GO:0015711 | organic anion transport | 185 | 30 | 13.86 | 4.50E-05 |
| GO:0014812 | muscle cell migration | 63 | 15 | 4.72 | 4.70E-05 |
| GO:0050731 | positive regulation of peptidyl-tyrosine phosphorylation | 93 | 19 | 6.97 | 4.80E-05 |
| GO:0048286 | lung alveolus development | 25 | 9 | 1.87 | 4.90E-05 |
| GO:0003156 | regulation of animal organ formation | 15 | 7 | 1.12 | 4.90E-05 |
| GO:0007417 | central nervous system development | 605 | 72 | 45.33 | 4.90E-05 |
| GO:0042475 | odontogenesis of dentin-containing tooth | 43 | 12 | 3.22 | 5.00E-05 |
| GO:0045669 | positive regulation of osteoblast differentiation | 43 | 12 | 3.22 | 5.00E-05 |
| GO:0051336 | regulation of hydrolase activity | 857 | 95 | 64.21 | 5.20E-05 |
| GO:0043410 | positive regulation of MAPK cascade | 299 | 42 | 22.4 | 5.30E-05 |
| GO:1901654 | response to ketone | 126 | 23 | 9.44 | 5.30E-05 |
| GO:0001516 | prostaglandin biosynthetic process | 20 | 8 | 1.5 | 5.40E-05 |
| GO:0046457 | prostanoid biosynthetic process | 20 | 8 | 1.5 | 5.40E-05 |
| GO:0009893 | positive regulation of metabolic process | 2542 | 237 | 190.45 | 5.40E-05 |
| GO:0050730 | regulation of peptidyl-tyrosine phosphorylation | 143 | 25 | 10.71 | 5.40E-05 |
| GO:0002687 | positive regulation of leukocyte migration | 71 | 16 | 5.32 | 5.40E-05 |
| GO:0010717 | regulation of epithelial to mesenchymal transition | 71 | 16 | 5.32 | 5.40E-05 |
| GO:0042310 | vasoconstriction | 37 | 11 | 2.77 | 5.50E-05 |
| GO:0045123 | cellular extravasation | 37 | 11 | 2.77 | 5.50E-05 |
| GO:0006692 | prostanoid metabolic process | 31 | 10 | 2.32 | 5.50E-05 |
| GO:0006693 | prostaglandin metabolic process | 31 | 10 | 2.32 | 5.50E-05 |
| GO:0032890 | regulation of organic acid transport | 31 | 10 | 2.32 | 5.50E-05 |
| GO:0060420 | regulation of heart growth | 50 | 13 | 3.75 | 5.60E-05 |
| GO:0035270 | endocrine system development | 64 | 15 | 4.8 | 5.70E-05 |
| GO:0031639 | plasminogen activation | 11 | 6 | 0.82 | 5.80E-05 |
| GO:0045987 | positive regulation of smooth muscle contraction | 11 | 6 | 0.82 | 5.80E-05 |
| GO:0061042 | vascular wound healing | 11 | 6 | 0.82 | 5.80E-05 |
| GO:0048812 | neuron projection morphogenesis | 420 | 54 | 31.47 | 5.90E-05 |
| GO:0018212 | peptidyl-tyrosine modification | 215 | 33 | 16.11 | 5.90E-05 |
| GO:0042327 | positive regulation of phosphorylation | 566 | 68 | 42.41 | 5.90E-05 |

| | | | | | |
|---|---|---|---|---|---|
| GO:0032231 | regulation of actin filament bundle assembly | 79 | 17 | 5.92 | 6.00E-05 |
| GO:0043207 | response to external biotic stimulus | 794 | 89 | 59.49 | 6.10E-05 |
| GO:0051707 | response to other organism | 794 | 89 | 59.49 | 6.10E-05 |
| GO:0052547 | regulation of peptidase activity | 263 | 38 | 19.7 | 6.50E-05 |
| GO:0001837 | epithelial to mesenchymal transition | 111 | 21 | 8.32 | 6.50E-05 |
| GO:0046942 | carboxylic acid transport | 136 | 24 | 10.19 | 6.50E-05 |
| GO:0031667 | response to nutrient levels | 292 | 41 | 21.88 | 6.60E-05 |
| GO:0006986 | response to unfolded protein | 162 | 27 | 12.14 | 6.60E-05 |
| GO:0089718 | amino acid import across plasma membrane | 26 | 9 | 1.95 | 6.90E-05 |
| GO:0051384 | response to glucocorticoid | 80 | 17 | 5.99 | 7.10E-05 |
| GO:0043502 | regulation of muscle adaptation | 58 | 14 | 4.35 | 7.10E-05 |
| GO:0015837 | amine transport | 32 | 10 | 2.4 | 7.40E-05 |
| GO:0035850 | epithelial cell differentiation involved in kidney development | 32 | 10 | 2.4 | 7.40E-05 |
| GO:0051952 | regulation of amine transport | 32 | 10 | 2.4 | 7.40E-05 |
| GO:2000351 | regulation of endothelial cell apoptotic process | 32 | 10 | 2.4 | 7.40E-05 |
| GO:0051897 | positive regulation of protein kinase B signaling | 88 | 18 | 6.59 | 7.50E-05 |
| GO:0048639 | positive regulation of developmental growth | 104 | 20 | 7.79 | 7.60E-05 |
| GO:0030278 | regulation of ossification | 73 | 16 | 5.47 | 7.80E-05 |
| GO:1900120 | regulation of receptor binding | 16 | 7 | 1.2 | 8.10E-05 |
| GO:0006941 | striated muscle contraction | 81 | 17 | 6.07 | 8.40E-05 |
| GO:0042129 | regulation of T cell proliferation | 81 | 17 | 6.07 | 8.40E-05 |
| GO:0007492 | endoderm development | 52 | 13 | 3.9 | 8.70E-05 |
| GO:0070252 | actin-mediated cell contraction | 52 | 13 | 3.9 | 8.70E-05 |
| GO:0070373 | negative regulation of ERK1 and ERK2 cascade | 52 | 13 | 3.9 | 8.70E-05 |
| GO:0045165 | cell fate commitment | 105 | 20 | 7.87 | 8.70E-05 |
| GO:0050670 | regulation of lymphocyte proliferation | 105 | 20 | 7.87 | 8.70E-05 |
| GO:0051247 | positive regulation of protein metabolic process | 1073 | 113 | 80.39 | 8.80E-05 |
| GO:0050890 | cognition | 156 | 26 | 11.69 | 8.90E-05 |
| GO:0007409 | axonogenesis | 296 | 41 | 22.18 | 8.90E-05 |
| GO:2000241 | regulation of reproductive process | 74 | 16 | 5.54 | 9.30E-05 |
| GO:0030003 | cellular cation homeostasis | 336 | 45 | 25.17 | 9.30E-05 |
| GO:0051480 | regulation of cytosolic calcium ion concentration | 139 | 24 | 10.41 | 9.30E-05 |
| GO:0042698 | ovulation cycle | 39 | 11 | 2.92 | 9.30E-05 |
| GO:0045685 | regulation of glial cell differentiation | 39 | 11 | 2.92 | 9.30E-05 |
| GO:0050918 | positive chemotaxis | 39 | 11 | 2.92 | 9.30E-05 |
| GO:0043405 | regulation of MAP kinase activity | 211 | 32 | 15.81 | 9.60E-05 |
| GO:0003229 | ventricular cardiac muscle tissue development | 27 | 9 | 2.02 | 9.70E-05 |
| GO:0030239 | myofibril assembly | 27 | 9 | 2.02 | 9.70E-05 |
| GO:0045687 | positive regulation of glial cell differentiation | 27 | 9 | 2.02 | 9.70E-05 |
| GO:0070169 | positive regulation of biomineral tissue development | 27 | 9 | 2.02 | 9.70E-05 |
| GO:0071711 | basement membrane organization | 27 | 9 | 2.02 | 9.70E-05 |
| GO:0030968 | endoplasmic reticulum unfolded protein response | 114 | 21 | 8.54 | 9.70E-05 |
| GO:0019216 | regulation of lipid metabolic process | 268 | 38 | 20.08 | 9.80E-05 |
| GO:0034620 | cellular response to unfolded protein | 131 | 23 | 9.81 | 1.00E-04 |
| GO:0006636 | unsaturated fatty acid biosynthetic process | 33 | 10 | 2.47 | 1.00E-04 |
| GO:0060675 | ureteric bud morphogenesis | 33 | 10 | 2.47 | 1.00E-04 |
| GO:0090049 | regulation of cell migration involved in sprouting angiogenesis | 33 | 10 | 2.47 | 1.00E-04 |

**Table S4. KEGG pathway analysis of genes regulated by colchicine intervention compared to cholesterol loaded HASMCs.** HASMCs were cholesterol loaded (10µg/mL) for 72 hours, followed by treatment with colchicine (50nM) for 48 hours.

| ID | Description | GeneRatio | BgRatio | pvalue | p.adjust | qvalue | Count |
|---|---|---|---|---|---|---|---|
| hsa04060 | Cytokine-cytokine receptor interaction | 26/442 | 96/5193 | 5.45E-08 | 1.64E-05 | 1.39E-05 | 26 |
| hsa04350 | TGF-beta signaling pathway | 22/442 | 84/5193 | 1.10E-06 | 0.000166 | 0.000141 | 22 |
| hsa04510 | Focal adhesion | 31/442 | 151/5193 | 2.46E-06 | 0.000247 | 0.00021 | 31 |
| hsa04080 | Neuroactive ligand-receptor interaction | 17/442 | 63/5193 | 1.21E-05 | 0.000912 | 0.000775 | 17 |
| hsa04020 | Calcium signaling pathway | 24/442 | 112/5193 | 1.62E-05 | 0.000977 | 0.000831 | 24 |
| hsa04512 | ECM-receptor interaction | 14/442 | 47/5193 | 2.11E-05 | 0.000987 | 0.000838 | 14 |
| hsa04514 | Cell adhesion molecules | 17/442 | 66/5193 | 2.37E-05 | 0.000987 | 0.000838 | 17 |
| hsa05200 | Pathways in cancer | 53/442 | 356/5193 | 2.62E-05 | 0.000987 | 0.000838 | 53 |
| hsa05205 | Proteoglycans in cancer | 29/442 | 154/5193 | 3.01E-05 | 0.001006 | 0.000855 | 29 |
| hsa04530 | Tight junction | 23/442 | 115/5193 | 7.71E-05 | 0.002319 | 0.001971 | 23 |
| hsa04390 | Hippo signaling pathway | 22/442 | 109/5193 | 9.55E-05 | 0.002462 | 0.002092 | 22 |
| hsa05410 | Hypertrophic cardiomyopathy | 13/442 | 47/5193 | 9.82E-05 | 0.002462 | 0.002092 | 13 |
| hsa04670 | Leukocyte transendothelial migration | 16/442 | 69/5193 | 0.00016 | 0.003715 | 0.003157 | 16 |
| hsa04610 | Complement and coagulation cascades | 10/442 | 32/5193 | 0.000209 | 0.004292 | 0.003647 | 10 |
| hsa04933 | AGE-RAGE signaling pathway in diabetic complications | 18/442 | 85/5193 | 0.000218 | 0.004292 | 0.003647 | 18 |
| hsa04015 | Rap1 signaling pathway | 25/442 | 139/5193 | 0.000228 | 0.004292 | 0.003647 | 25 |
| hsa05217 | Basal cell carcinoma | 11/442 | 39/5193 | 0.000279 | 0.004949 | 0.004205 | 11 |
| hsa04151 | PI3K-Akt signaling pathway | 34/442 | 220/5193 | 0.00039 | 0.006527 | 0.005547 | 34 |
| hsa00140 | Steroid hormone biosynthesis | 7/442 | 18/5193 | 0.000428 | 0.006786 | 0.005767 | 7 |
| hsa04550 | Signaling pathways regulating pluripotency of stem cells | 18/442 | 91/5193 | 0.000529 | 0.007964 | 0.006768 | 18 |
| hsa04810 | Regulation of actin cytoskeleton | 26/442 | 156/5193 | 0.000594 | 0.008513 | 0.007234 | 26 |
| hsa05144 | Malaria | 8/442 | 25/5193 | 0.000762 | 0.010156 | 0.008631 | 8 |
| hsa05146 | Amoebiasis | 13/442 | 57/5193 | 0.000776 | 0.010156 | 0.008631 | 13 |
| hsa04010 | MAPK signaling pathway | 32/442 | 213/5193 | 0.000951 | 0.011933 | 0.010141 | 32 |
| hsa04270 | Vascular smooth muscle contraction | 15/442 | 75/5193 | 0.001349 | 0.016242 | 0.013802 | 15 |
| hsa05414 | Dilated cardiomyopathy | 11/442 | 47/5193 | 0.001557 | 0.018022 | 0.015315 | 11 |
| hsa04072 | Phospholipase D signaling pathway | 18/442 | 100/5193 | 0.001675 | 0.018671 | 0.015867 | 18 |
| hsa04061 | Viral protein interaction with cytokine and cytokine receptor | 8/442 | 28/5193 | 0.001745 | 0.018756 | 0.015939 | 8 |
| hsa04974 | Protein digestion and absorption | 10/442 | 42/5193 | 0.002201 | 0.02284 | 0.01941 | 10 |
| hsa04024 | cAMP signaling pathway | 19/442 | 116/5193 | 0.003866 | 0.038785 | 0.032959 | 19 |
| hsa05418 | Fluid shear stress and atherosclerosis | 18/442 | 109/5193 | 0.004451 | 0.043219 | 0.036728 | 18 |
| hsa04750 | Inflammatory mediator regulation of TRP channels | 12/442 | 61/5193 | 0.004599 | 0.043259 | 0.036761 | 12 |
| hsa04640 | Hematopoietic cell lineage | 8/442 | 33/5193 | 0.005342 | 0.048723 | 0.041405 | 8 |

**Table S5. Differentially expressed genes regulated by colchicine compared to control HASMCs.** HASMCs were treated with colchicine (50nM) for 24 hours.

| ENSEMBL | name | LogFC_Col | padj_Col |
|---|---|---|---|
| ENSG00000000971.16 | CFH | -0.72539 | 0.000335 |
| ENSG00000002834.18 | LASP1 | 0.770472 | 1.43E-07 |
| ENSG00000005102.14 | MEOX1 | 1.524317 | 0.000372 |
| ENSG00000005187.12 | ACSM3 | -0.98545 | 0.018102 |
| ENSG00000005469.12 | CROT | -0.7191 | 7.75E-08 |
| ENSG00000006327.14 | TNFRSF12A | 1.811959 | 8.97E-30 |
| ENSG00000006468.14 | ETV1 | -1.81244 | 4.51E-21 |
| ENSG00000007237.19 | GAS7 | -2.02027 | 9.95E-06 |
| ENSG00000007866.22 | TEAD3 | 0.61692 | 7.00E-06 |
| ENSG00000007944.15 | MYLIP | -0.77087 | 9.96E-07 |
| ENSG00000008118.10 | CAMK1G | -2.1047 | 4.71E-05 |
| ENSG00000008300.17 | CELSR3 | 0.913935 | 3.93E-09 |
| ENSG00000008517.19 | IL32 | 1.145476 | 3.69E-22 |
| ENSG00000011052.21 | NME1-NME2 | 0.714132 | 0.000292 |
| ENSG00000011347.10 | SYT7 | 1.021092 | 0.001813 |
| ENSG00000011422.12 | PLAUR | 1.133778 | 4.25E-14 |
| ENSG00000011465.18 | DCN | -0.81663 | 1.22E-23 |
| ENSG00000011523.14 | CEP68 | -0.63598 | 1.99E-07 |
| ENSG00000013375.16 | PGM3 | 0.691781 | 1.38E-12 |
| ENSG00000013588.9 | GPRC5A | 1.625655 | 1.13E-15 |
| ENSG00000013619.15 | MAMLD1 | 0.708169 | 5.98E-06 |
| ENSG00000017483.15 | SLC38A5 | 1.079958 | 4.67E-09 |
| ENSG00000018408.15 | WWTR1 | -1.05831 | 2.20E-21 |
| ENSG00000020577.14 | SAMD4A | 1.068499 | 1.13E-09 |
| ENSG00000023697.13 | DERA | 0.62386 | 2.22E-06 |
| ENSG00000023902.14 | PLEKHO1 | 0.813581 | 1.74E-09 |
| ENSG00000025434.19 | NR1H3 | -0.86127 | 6.34E-05 |
| ENSG00000026559.14 | KCNG1 | 1.067235 | 9.75E-10 |
| ENSG00000028137.19 | TNFRSF1B | -1.41193 | 1.63E-06 |
| ENSG00000029153.14 | ARNTL2 | 0.998229 | 1.03E-25 |
| ENSG00000033867.16 | SLC4A7 | 0.58505 | 8.55E-06 |
| ENSG00000034152.19 | MAP2K3 | 0.953897 | 8.79E-18 |
| ENSG00000035403.18 | VCL | 0.612251 | 0.000536 |
| ENSG00000037897.17 | METTL1 | 0.591059 | 0.001983 |
| ENSG00000038382.20 | TRIO | 0.786153 | 8.44E-21 |
| ENSG00000038945.15 | MSR1 | -1.05097 | 0.004408 |
| ENSG00000039560.14 | RAI14 | 1.051513 | 9.26E-14 |
| ENSG00000041982.16 | TNC | 1.696794 | 0.003426 |
| ENSG00000042062.13 | RIPOR3 | -1.93506 | 0.016604 |
| ENSG00000044574.9 | HSPA5 | 0.708958 | 9.78E-15 |
| ENSG00000046604.14 | DSG2 | 0.810775 | 9.78E-15 |
| ENSG00000047648.23 | ARHGAP6 | -0.92573 | 1.79E-06 |
| ENSG00000048052.23 | HDAC9 | 1.202875 | 3.46E-19 |
| ENSG00000048162.21 | NOP16 | 0.657127 | 0.008143 |
| ENSG00000049130.16 | KITLG | -0.68506 | 0.003389 |
| ENSG00000049192.15 | ADAMTS6 | 0.674853 | 1.76E-06 |
| ENSG00000049449.10 | RCN1 | 0.703526 | 1.91E-14 |
| ENSG00000050405.13 | LIMA1 | 1.05092 | 2.81E-13 |
| ENSG00000051108.15 | HERPUD1 | 0.930453 | 5.29E-09 |
| ENSG00000052749.14 | RRP12 | 0.689696 | 1.05E-05 |
| ENSG00000054392.13 | HHAT | -0.82709 | 0.001402 |
| ENSG00000058668.15 | ATP2B4 | 1.018609 | 9.48E-24 |
| ENSG00000058866.15 | DGKG | -1.09958 | 0.043455 |
| ENSG00000059728.11 | MXD1 | 0.822711 | 4.27E-05 |
| ENSG00000063180.9 | CA11 | -1.06371 | 4.66E-05 |
| ENSG00000064651.14 | SLC12A2 | -1.24739 | 5.78E-24 |

| | | | |
|---|---|---|---|
| ENSG00000064652.11 | SNX24 | 0.776409 | 3.50E-07 |
| ENSG00000064687.13 | ABCA7 | -1.22743 | 4.41E-16 |
| ENSG00000064763.11 | FAR2 | 1.043075 | 5.43E-11 |
| ENSG00000064886.14 | CHI3L2 | -1.71461 | 0.016704 |
| ENSG00000064989.13 | CALCRL | -1.35397 | 5.28E-10 |
| ENSG00000065060.17 | UHRF1BP1 | 0.595068 | 0.001402 |
| ENSG00000065320.9 | NTN1 | 0.88817 | 0.01668 |
| ENSG00000065802.12 | ASB1 | 1.022909 | 2.43E-20 |
| ENSG00000066279.19 | ASPM | -0.77732 | 0.013995 |
| ENSG00000067057.18 | PFKP | 1.195866 | 1.02E-23 |
| ENSG00000067082.15 | KLF6 | 0.652225 | 6.84E-08 |
| ENSG00000067715.14 | SYT1 | -1.2167 | 3.98E-28 |
| ENSG00000068366.21 | ACSL4 | 0.91022 | 5.07E-21 |
| ENSG00000068438.15 | FTSJ1 | 0.755977 | 6.60E-11 |
| ENSG00000068650.19 | ATP11A | 0.703987 | 2.27E-12 |
| ENSG00000068781.21 | STON1-GTF2A1L | -1.20581 | 0.000154 |
| ENSG00000068971.14 | PPP2R5B | 0.918392 | 1.94E-12 |
| ENSG00000069399.15 | BCL3 | -0.73435 | 8.28E-09 |
| ENSG00000069702.11 | TGFBR3 | -0.62151 | 0.009275 |
| ENSG00000070404.10 | FSTL3 | 1.611801 | 1.72E-33 |
| ENSG00000070495.15 | JMJD6 | 0.896434 | 2.31E-09 |
| ENSG00000070669.17 | ASNS | 1.320059 | 0.043822 |
| ENSG00000071054.16 | MAP4K4 | -0.59052 | 3.78E-06 |
| ENSG00000071127.17 | WDR1 | 0.895812 | 1.76E-11 |
| ENSG00000071282.12 | LMCD1 | 1.183352 | 2.67E-06 |
| ENSG00000071575.12 | TRIB2 | -0.81682 | 9.92E-05 |
| ENSG00000072071.16 | ADGRL1 | -0.63786 | 2.18E-06 |
| ENSG00000072110.16 | ACTN1 | 1.256238 | 2.35E-24 |
| ENSG00000072163.20 | LIMS2 | 1.55673 | 0.01922 |
| ENSG00000072195.15 | SPEG | 0.737645 | 1.37E-08 |
| ENSG00000072201.14 | LNX1 | -1.12525 | 0.00034 |
| ENSG00000072952.20 | IRAG1 | 0.953796 | 0.017887 |
| ENSG00000073008.15 | PVR | 1.366679 | 2.83E-13 |
| ENSG00000073712.15 | FERMT2 | 0.795312 | 2.95E-07 |
| ENSG00000073921.18 | PICALM | 0.594235 | 1.30E-09 |
| ENSG00000074416.15 | MGLL | 1.512633 | 2.77E-37 |
| ENSG00000074695.6 | LMAN1 | 0.703444 | 1.90E-17 |
| ENSG00000074800.16 | ENO1 | 0.714518 | 2.90E-19 |
| ENSG00000075223.14 | SEMA3C | -0.64268 | 3.18E-08 |
| ENSG00000075426.12 | FOSL2 | 0.923685 | 3.29E-07 |
| ENSG00000075624.17 | ACTB | 0.847765 | 5.24E-07 |
| ENSG00000075711.21 | DLG1 | 0.872054 | 4.66E-18 |
| ENSG00000075826.17 | SEC31B | -0.66841 | 0.007693 |
| ENSG00000076351.13 | SLC46A1 | 0.823062 | 8.41E-07 |
| ENSG00000076356.7 | PLXNA2 | -0.71833 | 4.95E-10 |
| ENSG00000076513.17 | ANKRD13A | 0.787223 | 5.80E-16 |
| ENSG00000076555.15 | ACACB | -0.63635 | 0.009335 |
| ENSG00000076706.17 | MCAM | 1.20822 | 4.50E-20 |
| ENSG00000077157.22 | PPP1R12B | 0.741117 | 0.008583 |
| ENSG00000077942.19 | FBLN1 | -0.99959 | 4.61E-12 |
| ENSG00000080031.10 | PTPRH | 1.474137 | 0.000711 |
| ENSG00000080573.7 | COL5A3 | 0.889916 | 9.66E-15 |
| ENSG00000080823.23 | MOK | 0.698058 | 0.000698 |
| ENSG00000081051.8 | AFP | 1.281458 | 0.021785 |
| ENSG00000081189.16 | MEF2C | -0.60334 | 0.000926 |
| ENSG00000081923.15 | ATP8B1 | 1.687556 | 5.96E-23 |
| ENSG00000082126.18 | MPP4 | 1.879369 | 8.31E-11 |
| ENSG00000082438.18 | COBLL1 | 0.60605 | 2.13E-05 |
| ENSG00000082497.12 | SERTAD4 | -1.26796 | 2.34E-06 |
| ENSG00000082512.15 | TRAF5 | 0.644771 | 5.45E-06 |
| ENSG00000086598.11 | TMED2 | 0.628155 | 9.62E-13 |
| ENSG00000087053.20 | MTMR2 | 0.640783 | 2.93E-12 |
| ENSG00000087076.9 | HSD17B14 | -0.60012 | 0.004459 |

| | | | |
|---|---|---|---|
| ENSG00000087116.16 | ADAMTS2 | 0.597517 | 4.50E-05 |
| ENSG00000087263.17 | OGFOD1 | 0.665442 | 1.00E-08 |
| ENSG00000087842.11 | PIR | -0.59438 | 0.000724 |
| ENSG00000088325.16 | TPX2 | -0.64756 | 0.035876 |
| ENSG00000088448.14 | ANKRD10 | -0.68458 | 4.47E-05 |
| ENSG00000088882.8 | CPXM1 | -0.64368 | 0.011154 |
| ENSG00000088899.16 | LZTS3 | 0.719784 | 2.30E-06 |
| ENSG00000089280.19 | FUS | 0.608607 | 1.85E-07 |
| ENSG00000090238.12 | YPEL3 | -0.64381 | 1.61E-05 |
| ENSG00000090447.12 | TFAP4 | -0.69414 | 4.54E-05 |
| ENSG00000090530.10 | P3H2 | 0.65471 | 0.000906 |
| ENSG00000092068.20 | SLC7A8 | -1.41259 | 1.42E-24 |
| ENSG00000092841.19 | MYL6 | 0.612757 | 0.001104 |
| ENSG00000092969.12 | TGFB2 | 1.073236 | 0.030526 |
| ENSG00000094804.12 | CDC6 | 0.752491 | 0.026641 |
| ENSG00000095380.11 | NANS | 0.597234 | 1.30E-06 |
| ENSG00000095383.20 | TBC1D2 | 0.595222 | 0.005513 |
| ENSG00000095752.7 | IL11 | 1.167029 | 3.85E-10 |
| ENSG00000099337.5 | KCNK6 | 0.98686 | 0.000296 |
| ENSG00000099849.15 | RASSF7 | 0.768471 | 0.036068 |
| ENSG00000099860.9 | GADD45B | 0.8498 | 0.001481 |
| ENSG00000099953.10 | MMP11 | -0.58666 | 0.00064 |
| ENSG00000100027.17 | YPEL1 | -0.94077 | 0.001216 |
| ENSG00000100092.24 | SH3BP1 | -0.63854 | 0.000307 |
| ENSG00000100100.13 | PIK3IP1 | -0.64489 | 1.47E-05 |
| ENSG00000100139.14 | MICALL1 | 0.957356 | 1.27E-12 |
| ENSG00000100196.11 | KDELR3 | 0.889957 | 2.26E-09 |
| ENSG00000100219.16 | XBP1 | 0.796909 | 8.25E-08 |
| ENSG00000100292.18 | HMOX1 | -0.90143 | 6.64E-05 |
| ENSG00000100342.21 | APOL1 | -0.94879 | 0.009948 |
| ENSG00000100345.22 | MYH9 | 1.359208 | 8.72E-20 |
| ENSG00000100350.15 | FOXRED2 | -0.64345 | 2.41E-06 |
| ENSG00000100522.10 | GNPNAT1 | 0.6754 | 4.31E-12 |
| ENSG00000100626.17 | GALNT16 | 0.617059 | 0.000166 |
| ENSG00000100784.12 | RPS6KA5 | -1.57795 | 3.24E-12 |
| ENSG00000100883.13 | SRP54 | 0.586251 | 1.38E-09 |
| ENSG00000100934.15 | SEC23A | 0.657265 | 1.18E-10 |
| ENSG00000100979.15 | PLTP | -0.7725 | 8.98E-11 |
| ENSG00000101224.18 | CDC25B | -1.00356 | 5.15E-06 |
| ENSG00000101310.17 | SEC23B | 0.671128 | 2.15E-11 |
| ENSG00000101333.18 | PLCB4 | 0.811573 | 0.03329 |
| ENSG00000101335.10 | MYL9 | 0.973314 | 1.46E-05 |
| ENSG00000101367.9 | MAPRE1 | 0.627699 | 1.29E-06 |
| ENSG00000101608.13 | MYL12A | 0.760063 | 1.75E-07 |
| ENSG00000101680.15 | LAMA1 | 0.977954 | 7.89E-09 |
| ENSG00000101928.13 | MOSPD1 | 0.674399 | 1.31E-08 |
| ENSG00000101955.15 | SRPX | -0.83018 | 1.12E-11 |
| ENSG00000102057.10 | KCND1 | -0.77833 | 0.001396 |
| ENSG00000102174.10 | PHEX | -1.45489 | 6.15E-07 |
| ENSG00000102271.14 | KLHL4 | -1.01063 | 7.65E-10 |
| ENSG00000102760.13 | RGCC | 0.824359 | 0.001248 |
| ENSG00000103064.15 | SLC7A6 | 0.679206 | 1.60E-10 |
| ENSG00000103145.11 | HCFC1R1 | -0.73014 | 0.019128 |
| ENSG00000103187.8 | COTL1 | 0.830441 | 1.73E-15 |
| ENSG00000103257.9 | SLC7A5 | 1.563708 | 0.001688 |
| ENSG00000103404.15 | USP31 | 0.617224 | 2.54E-08 |
| ENSG00000103485.19 | QPRT | -1.02624 | 5.18E-10 |
| ENSG00000103489.12 | XYLT1 | 0.73971 | 1.12E-07 |
| ENSG00000103490.14 | PYCARD | -0.80121 | 1.85E-05 |
| ENSG00000103742.12 | IGDCC4 | -0.709 | 0.00284 |
| ENSG00000103811.18 | CTSH | -0.68325 | 0.017969 |
| ENSG00000103888.17 | CEMIP | 0.59255 | 0.01352 |
| ENSG00000104081.14 | BMF | -0.9531 | 0.000645 |

| | | | |
|---|---|---|---|
| ENSG00000104356.11 | POP1 | 0.976929 | 3.14E-09 |
| ENSG00000104368.19 | PLAT | -1.41286 | 8.21E-12 |
| ENSG00000104447.13 | TRPS1 | -0.75538 | 3.51E-10 |
| ENSG00000104490.18 | NCALD | -0.94158 | 1.50E-07 |
| ENSG00000104812.15 | GYS1 | 0.812541 | 4.52E-12 |
| ENSG00000104881.16 | PPP1R13L | 0.896626 | 2.96E-07 |
| ENSG00000104884.17 | ERCC2 | 0.816143 | 2.15E-16 |
| ENSG00000105204.14 | DYRK1B | -0.58654 | 6.22E-05 |
| ENSG00000105499.14 | PLA2G4C | 0.862093 | 0.000379 |
| ENSG00000105516.11 | DBP | -1.62528 | 2.89E-13 |
| ENSG00000105559.12 | PLEKHA4 | -0.6673 | 7.13E-08 |
| ENSG00000105655.19 | ISYNA1 | -0.84784 | 4.84E-08 |
| ENSG00000105810.10 | CDK6 | 0.713277 | 6.56E-12 |
| ENSG00000105928.16 | GSDME | 0.779542 | 7.48E-10 |
| ENSG00000105996.7 | HOXA2 | -1.35463 | 7.45E-09 |
| ENSG00000106003.13 | LFNG | 0.640891 | 0.04794 |
| ENSG00000106034.18 | CPED1 | 0.695449 | 3.89E-10 |
| ENSG00000106080.11 | FKBP14 | 0.98862 | 2.57E-24 |
| ENSG00000106348.18 | IMPDH1 | 0.827662 | 3.03E-10 |
| ENSG00000106351.13 | AGFG2 | -0.63859 | 0.000176 |
| ENSG00000106366.9 | SERPINE1 | 2.458508 | ######## |
| ENSG00000106511.6 | MEOX2 | -2.54655 | 6.38E-12 |
| ENSG00000106537.8 | TSPAN13 | 0.765868 | 0.000166 |
| ENSG00000106546.14 | AHR | 0.701726 | 0.000189 |
| ENSG00000106617.15 | PRKAG2 | 0.946127 | 1.06E-07 |
| ENSG00000106688.12 | SLC1A1 | 0.763985 | 3.95E-05 |
| ENSG00000106799.13 | TGFBR1 | 2.043998 | 6.56E-58 |
| ENSG00000106804.8 | C5 | -0.78661 | 3.09E-05 |
| ENSG00000106809.11 | OGN | -1.75851 | 0.03223 |
| ENSG00000106829.20 | TLE4 | -0.66885 | 7.96E-12 |
| ENSG00000106976.21 | DNM1 | -0.59002 | 4.92E-08 |
| ENSG00000107562.16 | CXCL12 | -0.87615 | 0.000141 |
| ENSG00000107821.15 | KAZALD1 | -0.73816 | 0.032278 |
| ENSG00000107984.10 | DKK1 | 1.713798 | 2.70E-29 |
| ENSG00000108387.16 | SEPTIN4 | -0.99903 | 0.00076 |
| ENSG00000108576.10 | SLC6A4 | -2.83691 | 0.030063 |
| ENSG00000108639.8 | SYNGR2 | 0.744932 | 0.00011 |
| ENSG00000108771.13 | DHX58 | -0.74991 | 0.004327 |
| ENSG00000108798.9 | ABI3 | -1.22638 | 0.007664 |
| ENSG00000108829.10 | LRRC59 | 0.990899 | 2.66E-26 |
| ENSG00000108854.16 | SMURF2 | 1.449755 | 3.35E-25 |
| ENSG00000108861.9 | DUSP3 | 0.643394 | 4.72E-09 |
| ENSG00000108947.5 | EFNB3 | -0.83247 | 1.51E-09 |
| ENSG00000108950.12 | FAM20A | -3.78691 | 1.24E-09 |
| ENSG00000108984.15 | MAP2K6 | -1.90986 | 1.56E-31 |
| ENSG00000109089.7 | CDR2L | 0.776884 | 1.75E-15 |
| ENSG00000109265.15 | CRACD | -1.52549 | 9.24E-21 |
| ENSG00000109339.24 | MAPK10 | -0.8125 | 0.000722 |
| ENSG00000109670.16 | FBXW7 | -0.79047 | 2.82E-08 |
| ENSG00000109738.11 | GLRB | 0.599592 | 0.00032 |
| ENSG00000109743.11 | BST1 | 0.829122 | 3.14E-08 |
| ENSG00000109814.12 | UGDH | 0.754554 | 1.61E-12 |
| ENSG00000109846.9 | CRYAB | 0.893604 | 0.013386 |
| ENSG00000109881.17 | CCDC34 | -0.7194 | 0.012177 |
| ENSG00000110002.16 | VWA5A | -0.8495 | 1.29E-10 |
| ENSG00000110042.8 | DTX4 | -0.70272 | 0.005157 |
| ENSG00000110318.15 | CEP126 | -0.75356 | 0.003765 |
| ENSG00000110852.5 | CLEC2B | -1.63334 | 0.000415 |
| ENSG00000110876.10 | SELPLG | 0.742008 | 5.24E-07 |
| ENSG00000110880.11 | CORO1C | 0.841028 | 4.00E-20 |
| ENSG00000110881.12 | ASIC1 | -0.86881 | 1.71E-13 |
| ENSG00000111087.10 | GLI1 | -0.77901 | 0.019128 |
| ENSG00000111110.12 | PPM1H | 0.717894 | 0.00056 |

| | | | |
|---|---|---|---|
| ENSG00000111186.13 | WNT5B | 1.507478 | 2.82E-09 |
| ENSG00000111206.13 | FOXM1 | -0.71399 | 0.010264 |
| ENSG00000111266.9 | DUSP16 | 1.17746 | 1.81E-19 |
| ENSG00000111341.10 | MGP | -1.41596 | 0.001796 |
| ENSG00000111615.14 | KRR1 | -0.76394 | 5.38E-12 |
| ENSG00000111641.12 | NOP2 | 0.602369 | 1.62E-06 |
| ENSG00000111696.12 | NT5DC3 | 0.977672 | 3.48E-11 |
| ENSG00000111711.10 | GOLT1B | 0.848262 | 2.35E-16 |
| ENSG00000111799.22 | COL12A1 | 0.845865 | 2.79E-10 |
| ENSG00000111801.16 | BTN3A3 | -0.7181 | 0.000551 |
| ENSG00000111859.17 | NEDD9 | 2.472087 | 2.45E-26 |
| ENSG00000111885.7 | MAN1A1 | -0.58763 | 5.06E-08 |
| ENSG00000111981.5 | ULBP1 | 0.888651 | 0.049704 |
| ENSG00000112078.14 | KCTD20 | 0.61517 | 0.000139 |
| ENSG00000112183.15 | RBM24 | -0.67722 | 0.000272 |
| ENSG00000112208.11 | BAG2 | 1.181889 | 1.33E-22 |
| ENSG00000112210.12 | RAB23 | 1.478838 | 4.00E-20 |
| ENSG00000112320.12 | SOBP | -0.61612 | 6.34E-06 |
| ENSG00000112419.14 | PHACTR2 | 0.714689 | 6.16E-14 |
| ENSG00000112658.8 | SRF | 0.850493 | 1.59E-12 |
| ENSG00000112659.14 | CUL9 | -0.59006 | 2.45E-06 |
| ENSG00000112715.25 | VEGFA | 0.932192 | 1.20E-26 |
| ENSG00000112769.20 | LAMA4 | -0.71916 | 2.71E-13 |
| ENSG00000112773.16 | TENT5A | -1.13886 | 2.19E-16 |
| ENSG00000112893.10 | MAN2A1 | 0.654247 | 1.71E-05 |
| ENSG00000112936.19 | C7 | -1.73947 | 5.47E-06 |
| ENSG00000113070.8 | HBEGF | 3.267261 | 2.66E-59 |
| ENSG00000113083.15 | LOX | 1.23946 | 1.31E-20 |
| ENSG00000113356.13 | POLR3G | 0.688066 | 0.004366 |
| ENSG00000113532.13 | ST8SIA4 | -1.99504 | 2.18E-09 |
| ENSG00000113578.18 | FGF1 | 1.843488 | 1.98E-12 |
| ENSG00000113615.13 | SEC24A | 0.664717 | 8.65E-13 |
| ENSG00000113645.15 | WWC1 | 0.951064 | 1.20E-06 |
| ENSG00000113721.14 | PDGFRB | -0.67259 | 1.38E-12 |
| ENSG00000113739.11 | STC2 | 1.660764 | 7.15E-36 |
| ENSG00000113805.8 | CNTN3 | -0.62476 | 5.80E-07 |
| ENSG00000113811.11 | SELENOK | 0.591253 | 5.00E-06 |
| ENSG00000114115.10 | RBP1 | -0.58824 | 0.008583 |
| ENSG00000114126.18 | TFDP2 | -0.60983 | 1.55E-05 |
| ENSG00000114200.10 | BCHE | -0.90739 | 0.002312 |
| ENSG00000114251.15 | WNT5A | 1.284483 | 1.04E-25 |
| ENSG00000114529.13 | C3orf52 | 0.728716 | 0.00172 |
| ENSG00000114737.16 | CISH | -0.85288 | 0.002345 |
| ENSG00000114850.7 | SSR3 | 0.998494 | 5.35E-23 |
| ENSG00000115091.12 | ACTR3 | 0.785611 | 3.19E-10 |
| ENSG00000115257.15 | PCSK4 | -1.04924 | 0.005835 |
| ENSG00000115363.14 | EVA1A | 0.616628 | 7.52E-05 |
| ENSG00000115461.5 | IGFBP5 | 2.648193 | 7.25E-11 |
| ENSG00000115525.18 | ST3GAL5 | -0.93391 | 7.08E-10 |
| ENSG00000115556.14 | PLCD4 | -1.04234 | 2.02E-05 |
| ENSG00000115594.12 | IL1R1 | -0.72832 | 1.53E-06 |
| ENSG00000115604.12 | IL18R1 | -1.30532 | 0.000623 |
| ENSG00000115641.19 | FHL2 | 0.793567 | 4.72E-06 |
| ENSG00000115738.10 | ID2 | -0.99467 | 5.24E-07 |
| ENSG00000115758.13 | ODC1 | 1.063771 | 9.79E-28 |
| ENSG00000115841.21 | RMDN2 | -0.63778 | 0.004997 |
| ENSG00000115896.16 | PLCL1 | -0.86121 | 0.002509 |
| ENSG00000115902.11 | SLC1A4 | 1.337456 | 0.005413 |
| ENSG00000116132.12 | PRRX1 | -0.73724 | 5.71E-14 |
| ENSG00000116191.18 | RALGPS2 | 0.648533 | 0.001297 |
| ENSG00000116194.13 | ANGPTL1 | -0.69741 | 0.003455 |
| ENSG00000116584.20 | ARHGEF2 | 0.663876 | 0.002766 |
| ENSG00000116649.10 | SRM | 0.642247 | 6.16E-10 |

| | | | |
|---|---|---|---|
| ENSG00000116704.8 | SLC35D1 | 0.852183 | 2.63E-10 |
| ENSG00000116741.8 | RGS2 | -2.17071 | 0.001016 |
| ENSG00000116761.12 | CTH | 1.058214 | 8.11E-08 |
| ENSG00000116991.11 | SIPA1L2 | -2.3185 | 1.22E-12 |
| ENSG00000117143.13 | UAP1 | 1.471579 | 1.30E-17 |
| ENSG00000117152.14 | RGS4 | 1.456812 | 1.84E-28 |
| ENSG00000117228.11 | GBP1 | 0.882336 | 9.91E-10 |
| ENSG00000117308.15 | GALE | 0.641058 | 0.000171 |
| ENSG00000117318.9 | ID3 | -0.72051 | 4.65E-09 |
| ENSG00000117394.24 | SLC2A1 | 1.039733 | 2.23E-26 |
| ENSG00000117395.13 | EBNA1BP2 | 0.814441 | 1.48E-08 |
| ENSG00000117425.14 | PTCH2 | -1.09674 | 0.009693 |
| ENSG00000117461.15 | PIK3R3 | -0.80816 | 0.006908 |
| ENSG00000117479.15 | SLC19A2 | 0.757999 | 5.26E-07 |
| ENSG00000117525.14 | F3 | 1.697191 | 7.30E-15 |
| ENSG00000117632.23 | STMN1 | -0.7757 | 2.26E-08 |
| ENSG00000117724.13 | CENPF | -0.89509 | 0.024507 |
| ENSG00000117758.14 | STX12 | 0.940454 | 2.94E-17 |
| ENSG00000117877.11 | POLR1G | 0.698668 | 8.11E-06 |
| ENSG00000118263.15 | KLF7 | 0.759378 | 1.26E-07 |
| ENSG00000118508.5 | RAB32 | 1.280721 | 3.94E-23 |
| ENSG00000118596.12 | SLC16A7 | 0.735694 | 2.79E-08 |
| ENSG00000118804.9 | STBD1 | 1.002962 | 7.75E-07 |
| ENSG00000118898.16 | PPL | -1.5275 | 0.040074 |
| ENSG00000118985.16 | ELL2 | 0.693825 | 3.03E-09 |
| ENSG00000119699.8 | TGFB3 | -0.86912 | 1.31E-08 |
| ENSG00000119771.15 | KLHL29 | 0.585291 | 2.73E-05 |
| ENSG00000119812.19 | FAM98A | 0.66402 | 9.46E-12 |
| ENSG00000119917.15 | IFIT3 | -0.77149 | 0.011103 |
| ENSG00000119922.11 | IFIT2 | -0.90064 | 1.64E-05 |
| ENSG00000120129.6 | DUSP1 | 1.974393 | 1.32E-35 |
| ENSG00000120217.14 | CD274 | 3.248637 | 2.24E-68 |
| ENSG00000120526.12 | NUDCD1 | 0.600216 | 1.34E-05 |
| ENSG00000120594.17 | PLXDC2 | -0.74118 | 4.95E-07 |
| ENSG00000120693.14 | SMAD9 | -0.60592 | 1.42E-05 |
| ENSG00000120742.11 | SERP1 | 0.711247 | 2.62E-16 |
| ENSG00000120832.10 | MTERF2 | -0.83083 | 0.001792 |
| ENSG00000121005.9 | CRISPLD1 | -0.71743 | 5.31E-05 |
| ENSG00000121440.15 | PDZRN3 | -0.95539 | 4.21E-15 |
| ENSG00000121957.15 | GPSM2 | -0.8528 | 0.000307 |
| ENSG00000122420.10 | PTGFR | -1.31018 | 1.93E-07 |
| ENSG00000122641.11 | INHBA | 1.184618 | 5.04E-17 |
| ENSG00000122694.16 | GLIPR2 | 0.624672 | 0.00185 |
| ENSG00000122786.20 | CALD1 | 0.818274 | 7.41E-09 |
| ENSG00000122861.16 | PLAU | 0.674346 | 1.37E-06 |
| ENSG00000122877.17 | EGR2 | -2.26221 | 2.55E-06 |
| ENSG00000122966.17 | CIT | -0.9613 | 0.013139 |
| ENSG00000122986.14 | HVCN1 | -0.76408 | 0.027372 |
| ENSG00000123130.17 | ACOT9 | 0.733314 | 7.42E-10 |
| ENSG00000123213.23 | NLN | 0.725704 | 6.95E-07 |
| ENSG00000123219.13 | CENPK | 0.740964 | 0.032407 |
| ENSG00000123358.20 | NR4A1 | 1.023885 | 3.97E-05 |
| ENSG00000123562.18 | MORF4L2 | 0.637092 | 3.42E-07 |
| ENSG00000123689.6 | G0S2 | -2.4214 | 0.000396 |
| ENSG00000123977.10 | DAW1 | 1.412041 | 4.28E-06 |
| ENSG00000123983.15 | ACSL3 | 0.901949 | 1.40E-14 |
| ENSG00000124145.6 | SDC4 | 0.82343 | 1.33E-11 |
| ENSG00000124208.16 | PEDS1-UBE2V1 | 0.736341 | 0.002033 |
| ENSG00000124575.7 | H1-3 | -0.81346 | 0.000103 |
| ENSG00000124593.16 | RP11-298J23.10 | -0.86909 | 1.72E-07 |
| ENSG00000124766.7 | SOX4 | -0.63652 | 0.000222 |
| ENSG00000124831.19 | LRRFIP1 | 1.072954 | 4.08E-17 |
| ENSG00000124920.14 | MYRF | 0.799113 | 1.08E-05 |

| | | | |
|---|---|---|---|
| ENSG00000125510.18 | OPRL1 | -1.16739 | 3.49E-05 |
| ENSG00000125629.15 | INSIG2 | 0.640499 | 0.000364 |
| ENSG00000125637.16 | PSD4 | 0.681711 | 0.042342 |
| ENSG00000125730.17 | C3 | -1.17041 | 0.042767 |
| ENSG00000125753.14 | VASP | 1.034669 | 8.98E-11 |
| ENSG00000125827.9 | TMX4 | -0.82352 | 1.98E-05 |
| ENSG00000125864.14 | BFSP1 | -1.2069 | 2.04E-06 |
| ENSG00000125912.11 | NCLN | 0.647008 | 1.03E-10 |
| ENSG00000125945.15 | ZNF436 | -0.86149 | 2.47E-09 |
| ENSG00000125966.10 | MMP24 | 1.129284 | 8.48E-05 |
| ENSG00000125968.9 | ID1 | -1.65634 | 1.19E-08 |
| ENSG00000126016.17 | AMOT | -0.61089 | 9.43E-07 |
| ENSG00000126522.17 | ASL | 0.60575 | 2.20E-06 |
| ENSG00000126562.17 | WNK4 | 1.002074 | 7.29E-13 |
| ENSG00000126709.16 | IFI6 | -0.65947 | 2.32E-05 |
| ENSG00000126790.12 | L3HYPDH | 0.623186 | 5.39E-05 |
| ENSG00000126861.5 | OMG | -2.51851 | 0.011679 |
| ENSG00000126950.8 | TMEM35A | -0.77814 | 0.00441 |
| ENSG00000127561.15 | SYNGR3 | 0.782971 | 0.001497 |
| ENSG00000127585.12 | FBXL16 | 0.709448 | 0.018749 |
| ENSG00000127824.15 | TUBA4A | -0.61951 | 0.003109 |
| ENSG00000127920.6 | GNG11 | -0.8256 | 2.13E-05 |
| ENSG00000127951.8 | FGL2 | -2.00365 | 1.74E-15 |
| ENSG00000128045.7 | RASL11B | -1.80517 | 0.000199 |
| ENSG00000128052.10 | KDR | -1.60985 | 1.72E-13 |
| ENSG00000128228.5 | SDF2L1 | 0.852722 | 1.68E-06 |
| ENSG00000128283.7 | CDC42EP1 | 0.678253 | 0.000586 |
| ENSG00000128284.19 | APOL3 | -1.12253 | 0.001205 |
| ENSG00000128294.16 | TPST2 | 0.696861 | 3.63E-06 |
| ENSG00000128342.5 | LIF | 1.480218 | 0.01796 |
| ENSG00000128482.16 | RNF112 | -1.27319 | 1.08E-09 |
| ENSG00000128512.23 | DOCK4 | -0.83122 | 7.81E-08 |
| ENSG00000128591.16 | FLNC | 1.18013 | 1.01E-22 |
| ENSG00000128595.17 | CALU | 0.725865 | 4.36E-17 |
| ENSG00000128656.15 | CHN1 | -0.68813 | 0.000122 |
| ENSG00000128849.11 | CGNL1 | -0.87473 | 2.21E-12 |
| ENSG00000129048.7 | ACKR4 | -1.09777 | 0.000252 |
| ENSG00000129116.19 | PALLD | 1.068426 | 2.49E-12 |
| ENSG00000129128.13 | SPCS3 | 0.658206 | 4.83E-14 |
| ENSG00000129226.14 | CD68 | -0.69368 | 2.41E-05 |
| ENSG00000130052.14 | STARD8 | -0.74695 | 0.028719 |
| ENSG00000130066.17 | SAT1 | -0.6471 | 0.018927 |
| ENSG00000130176.8 | CNN1 | 2.195018 | 2.77E-05 |
| ENSG00000130203.10 | APOE | -1.0139 | 1.44E-05 |
| ENSG00000130204.13 | TOMM40 | 0.632084 | 1.05E-07 |
| ENSG00000130283.9 | GDF1 | -1.23615 | 0.00092 |
| ENSG00000130402.13 | ACTN4 | 0.948123 | 1.08E-13 |
| ENSG00000130513.6 | GDF15 | 0.740155 | 3.56E-12 |
| ENSG00000130592.17 | LSP1 | -1.02098 | 0.018855 |
| ENSG00000130600.19 | H19 | -1.0207 | 0.036947 |
| ENSG00000130653.16 | PNPLA7 | -0.90849 | 0.000264 |
| ENSG00000131015.5 | ULBP2 | 1.20149 | 4.17E-10 |
| ENSG00000131018.25 | SYNE1 | 0.768778 | 3.88E-18 |
| ENSG00000131019.11 | ULBP3 | 0.84672 | 7.19E-07 |
| ENSG00000131236.18 | CAP1 | 0.642006 | 6.20E-12 |
| ENSG00000131389.18 | SLC6A6 | -0.76969 | 0.001078 |
| ENSG00000131477.11 | RAMP2 | -0.64824 | 0.031125 |
| ENSG00000131634.14 | TMEM204 | -0.69483 | 0.001114 |
| ENSG00000131773.14 | KHDRBS3 | 0.91386 | 1.90E-08 |
| ENSG00000131778.19 | CHD1L | 0.655816 | 3.12E-07 |
| ENSG00000131871.15 | SELENOS | 0.753456 | 1.10E-08 |
| ENSG00000132122.12 | SPATA6 | -0.62927 | 9.49E-05 |
| ENSG00000132205.11 | EMILIN2 | 1.328047 | 3.07E-10 |

| | | | |
|---|---|---|---|
| ENSG00000132386.11 | SERPINF1 | -0.86817 | 5.62E-13 |
| ENSG00000132432.14 | SEC61G | 0.77641 | 1.27E-12 |
| ENSG00000132481.7 | TRIM47 | -0.63572 | 0.003244 |
| ENSG00000132768.14 | DPH2 | 0.678909 | 8.32E-05 |
| ENSG00000133107.15 | TRPC4 | 2.27671 | 1.87E-51 |
| ENSG00000133110.15 | POSTN | 0.632483 | 0.005571 |
| ENSG00000133321.11 | PLAAT4 | -1.40583 | 7.42E-05 |
| ENSG00000133466.14 | C1QTNF6 | -0.65651 | 1.55E-10 |
| ENSG00000133657.16 | ATP13A3 | 0.735456 | 1.34E-07 |
| ENSG00000133816.18 | MICAL2 | 1.481729 | 9.54E-19 |
| ENSG00000133818.14 | RRAS2 | 0.862749 | 9.40E-08 |
| ENSG00000133943.21 | DGLUCY | -0.6615 | 0.000149 |
| ENSG00000134245.18 | WNT2B | 1.012843 | 0.000342 |
| ENSG00000134259.6 | NGF | 1.203836 | 1.88E-05 |
| ENSG00000134285.11 | FKBP11 | 0.633546 | 6.37E-06 |
| ENSG00000134333.14 | LDHA | 0.718445 | 2.00E-10 |
| ENSG00000134343.14 | ANO3 | -1.17606 | 0.04619 |
| ENSG00000134352.20 | IL6ST | -0.65681 | 1.15E-12 |
| ENSG00000134375.11 | TIMM17A | 0.639495 | 2.84E-06 |
| ENSG00000134569.10 | LRP4 | -1.02607 | 2.67E-12 |
| ENSG00000134668.12 | SPOCD1 | 1.597356 | 1.33E-49 |
| ENSG00000134684.12 | YARS1 | 0.627563 | 0.000856 |
| ENSG00000134697.13 | GNL2 | 0.646097 | 6.83E-07 |
| ENSG00000134853.12 | PDGFRA | -1.11129 | 4.50E-29 |
| ENSG00000134871.19 | COL4A2 | 0.676979 | 6.27E-06 |
| ENSG00000134986.14 | NREP | 0.74363 | 3.04E-18 |
| ENSG00000135047.16 | CTSL | -0.71502 | 2.63E-07 |
| ENSG00000135074.16 | ADAM19 | 0.757631 | 2.84E-06 |
| ENSG00000135269.18 | TES | 0.83014 | 1.04E-12 |
| ENSG00000135272.12 | MDFIC | -0.67148 | 2.30E-09 |
| ENSG00000135312.7 | HTR1B | 2.562581 | 6.09E-10 |
| ENSG00000135318.12 | NT5E | -0.66408 | 3.64E-06 |
| ENSG00000135363.12 | LMO2 | -1.44158 | 0.000108 |
| ENSG00000135424.18 | ITGA7 | 0.920339 | 0.002447 |
| ENSG00000135472.9 | FAIM2 | -1.57588 | 7.96E-14 |
| ENSG00000135480.16 | KRT7 | 1.452769 | 0.02283 |
| ENSG00000135631.17 | RAB11FIP5 | 0.626709 | 2.87E-06 |
| ENSG00000135636.15 | DYSF | 0.844264 | 0.004948 |
| ENSG00000135698.10 | MPHOSPH6 | 0.646988 | 0.000189 |
| ENSG00000136026.14 | CKAP4 | 0.612832 | 1.06E-10 |
| ENSG00000136048.14 | DRAM1 | 0.601056 | 2.13E-05 |
| ENSG00000136068.16 | FLNB | 1.065802 | 5.88E-17 |
| ENSG00000136205.17 | TNS3 | -0.92316 | 4.91E-25 |
| ENSG00000136240.10 | KDELR2 | 0.714939 | 1.68E-09 |
| ENSG00000136404.16 | TM6SF1 | 0.691298 | 0.039517 |
| ENSG00000136492.10 | BRIP1 | 0.820163 | 0.000696 |
| ENSG00000136574.19 | GATA4 | -0.87342 | 2.17E-15 |
| ENSG00000136628.18 | EPRS1 | 0.587163 | 0.000498 |
| ENSG00000136826.15 | KLF4 | -0.85574 | 1.87E-08 |
| ENSG00000136928.7 | GABBR2 | 1.207365 | 5.04E-17 |
| ENSG00000136960.13 | ENPP2 | -1.09813 | 0.004215 |
| ENSG00000136997.21 | MYC | 1.003033 | 1.09E-12 |
| ENSG00000137054.16 | POLR1E | 0.649311 | 3.19E-05 |
| ENSG00000137076.21 | TLN1 | 0.657173 | 5.99E-07 |
| ENSG00000137124.8 | ALDH1B1 | 1.094227 | 2.43E-10 |
| ENSG00000137168.8 | PPIL1 | 0.752671 | 3.77E-07 |
| ENSG00000137198.10 | GMPR | -0.66062 | 0.002375 |
| ENSG00000137203.15 | TFAP2A | 0.682196 | 0.000416 |
| ENSG00000137312.15 | FLOT1 | -0.88131 | 2.70E-05 |
| ENSG00000137331.12 | IER3 | 1.50971 | 1.25E-32 |
| ENSG00000137393.10 | RNF144B | 1.009073 | 0.000813 |
| ENSG00000137449.17 | CPEB2 | 0.915622 | 5.94E-07 |
| ENSG00000137486.17 | ARRB1 | -0.71956 | 6.37E-09 |

| | | | |
|---|---|---|---|
| ENSG00000137501.17 | SYTL2 | 0.991665 | 1.54E-08 |
| ENSG00000137563.13 | GGH | 0.670903 | 5.85E-06 |
| ENSG00000137801.11 | THBS1 | 0.709625 | 0.000157 |
| ENSG00000137807.16 | KIF23 | 0.822973 | 0.001265 |
| ENSG00000137880.6 | GCHFR | -0.93697 | 0.042675 |
| ENSG00000137941.17 | TTLL7 | 0.719839 | 2.15E-05 |
| ENSG00000137955.16 | RABGGTB | 0.651727 | 1.08E-09 |
| ENSG00000137959.17 | IFI44L | -1.40751 | 0.030033 |
| ENSG00000137965.11 | IFI44 | -1.14777 | 0.000677 |
| ENSG00000137975.8 | CLCA2 | -0.99603 | 1.36E-05 |
| ENSG00000138018.18 | SELENOI | 0.812611 | 4.27E-16 |
| ENSG00000138061.12 | CYP1B1 | 1.051398 | 2.02E-05 |
| ENSG00000138071.14 | ACTR2 | 0.600786 | 1.04E-10 |
| ENSG00000138160.7 | KIF11 | -0.6876 | 0.010132 |
| ENSG00000138172.11 | CALHM2 | -0.63768 | 9.14E-05 |
| ENSG00000138316.11 | ADAMTS14 | 1.524299 | 2.73E-15 |
| ENSG00000138356.14 | AOX1 | 1.626257 | 2.64E-20 |
| ENSG00000138378.19 | STAT4 | 1.273304 | 4.24E-05 |
| ENSG00000138380.18 | CARF | -0.74648 | 2.50E-05 |
| ENSG00000138449.11 | SLC40A1 | -2.02735 | 9.89E-36 |
| ENSG00000138606.19 | SHF | -0.72924 | 0.007488 |
| ENSG00000138675.17 | FGF5 | 1.320642 | 1.59E-37 |
| ENSG00000138685.17 | FGF2 | 0.832583 | 7.67E-16 |
| ENSG00000138771.16 | SHROOM3 | 1.691215 | 7.56E-10 |
| ENSG00000138835.22 | RGS3 | 0.808175 | 6.39E-12 |
| ENSG00000139116.19 | KIF21A | 0.645494 | 0.000309 |
| ENSG00000139269.3 | INHBE | 2.186444 | 0.000109 |
| ENSG00000139278.10 | GLIPR1 | 0.976442 | 7.20E-27 |
| ENSG00000139289.14 | PHLDA1 | -1.43915 | 7.92E-37 |
| ENSG00000139329.5 | LUM | -0.60558 | 0.000108 |
| ENSG00000139514.13 | SLC7A1 | 1.265738 | 1.10E-10 |
| ENSG00000139531.13 | SUOX | -0.64071 | 5.02E-06 |
| ENSG00000139597.18 | N4BP2L1 | -1.20801 | 2.97E-06 |
| ENSG00000139625.13 | MAP3K12 | -0.59026 | 2.67E-05 |
| ENSG00000139629.16 | GALNT6 | 0.869695 | 1.53E-05 |
| ENSG00000139645.11 | ANKRD52 | 0.672438 | 4.72E-09 |
| ENSG00000139679.16 | LPAR6 | -0.86038 | 0.002471 |
| ENSG00000139734.19 | DIAPH3 | 1.750267 | 1.96E-37 |
| ENSG00000139874.6 | SSTR1 | 1.359226 | 2.72E-05 |
| ENSG00000139899.11 | CBLN3 | -0.73878 | 1.54E-05 |
| ENSG00000139926.16 | FRMD6 | 1.02631 | 8.80E-36 |
| ENSG00000140398.14 | NEIL1 | -0.95324 | 0.000247 |
| ENSG00000140416.23 | TPM1 | 1.233284 | 0.005204 |
| ENSG00000140538.16 | NTRK3 | -1.0035 | 0.000126 |
| ENSG00000140650.13 | PMM2 | 0.680873 | 2.30E-08 |
| ENSG00000140859.16 | KIFC3 | 0.615922 | 5.29E-09 |
| ENSG00000140873.16 | ADAMTS18 | -3.26832 | 1.70E-15 |
| ENSG00000140876.11 | NUDT7 | -0.8589 | 0.011207 |
| ENSG00000141338.14 | ABCA8 | -1.27654 | 1.24E-36 |
| ENSG00000141448.11 | GATA6 | 0.658014 | 0.041472 |
| ENSG00000141469.18 | SLC14A1 | -2.21367 | 2.34E-06 |
| ENSG00000141540.11 | TTYH2 | -0.83374 | 1.85E-05 |
| ENSG00000141682.12 | PMAIP1 | 1.522527 | 3.07E-11 |
| ENSG00000142197.12 | DOP1B | 1.027145 | 1.43E-16 |
| ENSG00000142235.13 | LMTK3 | -1.16076 | 0.000628 |
| ENSG00000142871.18 | CCN1 | 1.240062 | 7.46E-22 |
| ENSG00000142949.17 | PTPRF | 1.015716 | 3.60E-32 |
| ENSG00000142961.15 | MOB3C | -0.77649 | 1.59E-06 |
| ENSG00000143127.13 | ITGA10 | -0.79341 | 3.83E-08 |
| ENSG00000143179.16 | UCK2 | 1.354889 | 1.80E-25 |
| ENSG00000143322.21 | ABL2 | 0.645665 | 2.32E-09 |
| ENSG00000143367.16 | TUFT1 | 1.305406 | 1.37E-08 |
| ENSG00000143387.14 | CTSK | -0.91973 | 3.61E-08 |

| | | | |
|---|---|---|---|
| ENSG00000143409.16 | MINDY1 | -0.67456 | 1.02E-07 |
| ENSG00000143416.21 | SELENBP1 | -1.41301 | 1.19E-10 |
| ENSG00000143443.10 | C1orf56 | -0.70811 | 0.012523 |
| ENSG00000143494.16 | VASH2 | -1.42643 | 0.016774 |
| ENSG00000143578.16 | CREB3L4 | -0.62556 | 0.006882 |
| ENSG00000143603.19 | KCNN3 | -1.08165 | 0.013139 |
| ENSG00000143753.13 | DEGS1 | 0.640567 | 1.39E-12 |
| ENSG00000143867.7 | OSR1 | -1.26937 | 1.26E-14 |
| ENSG00000144283.22 | PKP4 | 0.639265 | 2.22E-07 |
| ENSG00000144339.12 | TMEFF2 | -1.61012 | 3.62E-09 |
| ENSG00000144583.5 | MARCHF4 | 0.913639 | 3.29E-08 |
| ENSG00000144591.20 | GMPPA | 0.720935 | 2.82E-12 |
| ENSG00000144655.15 | CSRNP1 | 0.597131 | 0.003723 |
| ENSG00000144681.11 | STAC | -0.78285 | 0.044656 |
| ENSG00000144824.21 | PHLDB2 | 1.494151 | 9.31E-20 |
| ENSG00000144867.13 | SRPRB | 0.915188 | 1.28E-16 |
| ENSG00000144959.11 | NCEH1 | 0.951469 | 0.000229 |
| ENSG00000145012.14 | LPP | 0.721067 | 1.25E-12 |
| ENSG00000145050.19 | MANF | 0.673381 | 2.69E-09 |
| ENSG00000145246.14 | ATP10D | 0.751916 | 3.49E-12 |
| ENSG00000145247.12 | OCIAD2 | 0.934974 | 8.64E-06 |
| ENSG00000145358.6 | DDIT4L | -1.95055 | 1.81E-09 |
| ENSG00000145390.11 | USP53 | 0.947916 | 2.02E-06 |
| ENSG00000145431.11 | PDGFC | 0.608805 | 6.52E-11 |
| ENSG00000145506.14 | NKD2 | -1.82868 | 0.000155 |
| ENSG00000145632.15 | PLK2 | -0.62621 | 3.99E-07 |
| ENSG00000145687.16 | SSBP2 | -0.82884 | 9.73E-10 |
| ENSG00000145779.8 | TNFAIP8 | -0.91767 | 2.75E-10 |
| ENSG00000145934.16 | TENM2 | 3.022896 | 8.88E-52 |
| ENSG00000145990.11 | GFOD1 | -0.73132 | 0.002553 |
| ENSG00000146021.15 | KLHL3 | -0.82512 | 0.008794 |
| ENSG00000146054.18 | TRIM7 | 0.933281 | 0.022426 |
| ENSG00000146072.6 | TNFRSF21 | -0.69006 | 8.38E-07 |
| ENSG00000146267.12 | FAXC | 0.720829 | 0.005725 |
| ENSG00000146278.11 | PNRC1 | -0.58871 | 2.45E-09 |
| ENSG00000146409.12 | SLC18B1 | -0.67764 | 0.029271 |
| ENSG00000146411.6 | SLC2A12 | -1.13105 | 8.42E-09 |
| ENSG00000146477.6 | SLC22A3 | -1.25503 | 0.000264 |
| ENSG00000146592.17 | CREB5 | 2.206294 | 1.00E-19 |
| ENSG00000147027.4 | TMEM47 | 0.706854 | 2.41E-05 |
| ENSG00000147065.17 | MSN | 0.6043 | 2.31E-09 |
| ENSG00000147224.13 | PRPS1 | 0.884692 | 2.56E-07 |
| ENSG00000147251.16 | DOCK11 | -0.67999 | 0.001933 |
| ENSG00000147408.14 | CSGALNACT1 | -1.67317 | 2.93E-31 |
| ENSG00000147576.17 | ADHFE1 | -0.77658 | 0.005482 |
| ENSG00000147650.12 | LRP12 | 0.587354 | 1.05E-09 |
| ENSG00000148154.10 | UGCG | 1.230094 | 5.11E-18 |
| ENSG00000148225.16 | WDR31 | -0.60036 | 0.015013 |
| ENSG00000148344.11 | PTGES | -1.02668 | 0.030033 |
| ENSG00000148516.23 | ZEB1 | -0.72836 | 2.61E-06 |
| ENSG00000148541.13 | FAM13C | -1.60564 | 1.17E-10 |
| ENSG00000148677.7 | ANKRD1 | 1.661131 | 7.99E-06 |
| ENSG00000148700.15 | ADD3 | -1.06603 | 1.72E-27 |
| ENSG00000148843.15 | PDCD11 | 0.629143 | 2.82E-09 |
| ENSG00000148848.15 | ADAM12 | 0.907428 | 2.10E-23 |
| ENSG00000148935.11 | GAS2 | -1.51622 | 2.35E-05 |
| ENSG00000149054.16 | ZNF215 | 0.707204 | 0.028645 |
| ENSG00000149212.12 | SESN3 | -1.03085 | 1.79E-12 |
| ENSG00000149256.16 | TENM4 | -0.91131 | 2.83E-13 |
| ENSG00000149428.19 | HYOU1 | 0.77394 | 7.08E-21 |
| ENSG00000149489.9 | ROM1 | -0.65292 | 0.015045 |
| ENSG00000149571.12 | KIRREL3 | 1.002581 | 2.71E-05 |
| ENSG00000149591.17 | TAGLN | 1.579784 | 0.019819 |

| | | | |
|---|---|---|---|
| ENSG00000149633.12 | KIAA1755 | -1.41912 | 2.41E-14 |
| ENSG00000149809.17 | TM7SF2 | -0.65161 | 0.000116 |
| ENSG00000150394.14 | CDH8 | 0.793808 | 0.021226 |
| ENSG00000150510.17 | FAM124A | 0.753478 | 0.002797 |
| ENSG00000150551.11 | LYPD1 | 0.717057 | 0.012105 |
| ENSG00000150593.18 | PDCD4 | -0.65984 | 1.49E-06 |
| ENSG00000150636.17 | CCDC102B | -1.59551 | 0.004818 |
| ENSG00000150753.12 | CCT5 | 0.586785 | 8.04E-10 |
| ENSG00000150787.8 | PTS | 0.593233 | 3.15E-05 |
| ENSG00000150907.10 | FOXO1 | -0.70496 | 2.57E-06 |
| ENSG00000150938.10 | CRIM1 | 1.082919 | 2.03E-12 |
| ENSG00000150961.15 | SEC24D | 0.732942 | 4.25E-12 |
| ENSG00000151136.15 | BTBD11 | -1.30496 | 2.38E-13 |
| ENSG00000151150.22 | ANK3 | -0.92874 | 1.51E-08 |
| ENSG00000151151.6 | IPMK | -0.77983 | 0.000187 |
| ENSG00000151632.17 | AKR1C2 | -1.88405 | 5.97E-18 |
| ENSG00000151692.15 | RNF144A | -1.46946 | 2.96E-10 |
| ENSG00000151729.11 | SLC25A4 | 0.598381 | 0.000595 |
| ENSG00000151835.17 | SACS | 1.330699 | 2.55E-26 |
| ENSG00000151892.16 | GFRA1 | 2.189782 | 1.28E-09 |
| ENSG00000152056.17 | AP1S3 | 1.608078 | 3.93E-18 |
| ENSG00000152137.8 | HSPB8 | 0.72551 | 9.20E-13 |
| ENSG00000152217.20 | SETBP1 | -0.83687 | 1.73E-05 |
| ENSG00000152413.15 | HOMER1 | 0.740329 | 4.93E-06 |
| ENSG00000152518.8 | ZFP36L2 | -0.75908 | 1.72E-10 |
| ENSG00000152689.18 | RASGRP3 | -0.81842 | 0.017213 |
| ENSG00000152749.8 | GPR180 | 0.64713 | 2.10E-08 |
| ENSG00000152763.17 | DNAI4 | -0.68548 | 0.009248 |
| ENSG00000152804.11 | HHEX | -1.28964 | 1.17E-09 |
| ENSG00000152952.12 | PLOD2 | 0.636596 | 2.46E-08 |
| ENSG00000153066.13 | TXNDC11 | 0.648835 | 8.66E-10 |
| ENSG00000153094.24 | BCL2L11 | -1.1388 | 7.78E-07 |
| ENSG00000153208.17 | MERTK | -0.76954 | 0.007606 |
| ENSG00000153832.12 | FBXO36 | -0.61621 | 0.041316 |
| ENSG00000153879.9 | CEBPG | 0.784273 | 0.002252 |
| ENSG00000153885.15 | KCTD15 | 0.614713 | 0.000595 |
| ENSG00000153904.21 | DDAH1 | 1.315882 | 9.73E-10 |
| ENSG00000153989.8 | NUS1 | 0.648243 | 4.44E-13 |
| ENSG00000154127.10 | UBASH3B | 0.900555 | 8.16E-07 |
| ENSG00000154146.13 | NRGN | -0.87611 | 0.045883 |
| ENSG00000154217.15 | PITPNC1 | -0.79049 | 0.000925 |
| ENSG00000154258.17 | ABCA9 | -0.96304 | 3.58E-27 |
| ENSG00000154262.13 | ABCA6 | -1.15862 | 9.21E-34 |
| ENSG00000154263.17 | ABCA10 | -0.8841 | 0.00151 |
| ENSG00000154380.17 | ENAH | 1.114522 | 6.84E-26 |
| ENSG00000154511.12 | DIPK1A | 0.612239 | 0.000393 |
| ENSG00000154545.17 | MAGED4 | 0.635412 | 8.87E-06 |
| ENSG00000154553.16 | PDLIM3 | 1.036582 | 1.14E-10 |
| ENSG00000154639.19 | CXADR | 1.254187 | 5.09E-08 |
| ENSG00000154640.15 | BTG3 | 0.628264 | 4.68E-06 |
| ENSG00000154721.15 | JAM2 | -1.49573 | 2.68E-16 |
| ENSG00000155096.15 | AZIN1 | 0.607095 | 9.66E-12 |
| ENSG00000155304.6 | HSPA13 | 0.587812 | 2.02E-08 |
| ENSG00000155755.19 | TMEM237 | 0.766997 | 5.80E-07 |
| ENSG00000155760.3 | FZD7 | -0.7602 | 0.003266 |
| ENSG00000155966.14 | AFF2 | -0.75937 | 2.91E-10 |
| ENSG00000156103.16 | MMP16 | -0.6321 | 2.78E-09 |
| ENSG00000156265.16 | MAP3K7CL | 2.334515 | 0.000156 |
| ENSG00000156273.16 | BACH1 | 0.797492 | 9.78E-08 |
| ENSG00000156398.13 | SFXN2 | 0.86938 | 0.001495 |
| ENSG00000156463.18 | SH3RF2 | -0.83378 | 0.004379 |
| ENSG00000157240.4 | FZD1 | -0.62265 | 0.000139 |
| ENSG00000157368.11 | IL34 | -0.66056 | 0.002101 |

| | | | |
|---|---|---|---|
| ENSG00000157404.16 | KIT | 3.233989 | 2.73E-46 |
| ENSG00000157514.17 | TSC22D3 | -0.76673 | 2.41E-12 |
| ENSG00000157680.16 | DGKI | 1.258691 | 7.38E-19 |
| ENSG00000158023.10 | CFAP251 | 0.655208 | 0.011114 |
| ENSG00000158270.12 | COLEC12 | -1.53902 | 3.19E-14 |
| ENSG00000158292.7 | GPR153 | -0.7618 | 6.14E-13 |
| ENSG00000158710.15 | TAGLN2 | 0.825715 | 6.56E-15 |
| ENSG00000158813.18 | EDA | -0.79511 | 0.011829 |
| ENSG00000158859.10 | ADAMTS4 | 0.840147 | 4.68E-06 |
| ENSG00000158966.16 | CACHD1 | -0.59671 | 0.001812 |
| ENSG00000159176.14 | CSRP1 | 1.247004 | 2.68E-16 |
| ENSG00000159199.14 | ATP5MC1 | 0.618748 | 3.06E-06 |
| ENSG00000159200.18 | RCAN1 | 1.642662 | 2.70E-29 |
| ENSG00000159261.12 | CLDN14 | 1.753381 | 1.29E-06 |
| ENSG00000159403.18 | C1R | -0.69112 | 1.98E-09 |
| ENSG00000159479.17 | MED8 | 0.597072 | 0.000295 |
| ENSG00000159761.15 | C16orf86 | -1.23936 | 0.017677 |
| ENSG00000159840.16 | ZYX | 1.087796 | 3.90E-10 |
| ENSG00000160072.20 | ATAD3B | 0.729202 | 1.24E-08 |
| ENSG00000160145.16 | KALRN | -0.60881 | 0.003162 |
| ENSG00000160193.12 | WDR4 | 0.618526 | 0.000984 |
| ENSG00000160870.15 | CYP3A7 | -1.30412 | 6.62E-05 |
| ENSG00000161638.11 | ITGA5 | 0.602054 | 5.77E-13 |
| ENSG00000161682.15 | FAM171A2 | -0.60634 | 3.89E-06 |
| ENSG00000161888.11 | SPC24 | -0.83054 | 0.033112 |
| ENSG00000162407.9 | PLPP3 | -1.12715 | 7.63E-28 |
| ENSG00000162433.15 | AK4 | 0.742886 | 6.27E-05 |
| ENSG00000162490.7 | DRAXIN | -1.47784 | 0.000593 |
| ENSG00000162493.17 | PDPN | 1.199022 | 0.000601 |
| ENSG00000162496.9 | DHRS3 | -0.9322 | 1.72E-06 |
| ENSG00000162520.15 | SYNC | 0.936898 | 2.25E-11 |
| ENSG00000162595.7 | DIRAS3 | -1.24403 | 5.98E-12 |
| ENSG00000162614.19 | NEXN | 0.852047 | 7.79E-05 |
| ENSG00000162616.9 | DNAJB4 | 1.05202 | 1.70E-31 |
| ENSG00000162631.20 | NTNG1 | -1.15045 | 3.49E-05 |
| ENSG00000162636.16 | FAM102B | -0.6112 | 0.000299 |
| ENSG00000162645.13 | GBP2 | -0.83058 | 6.15E-05 |
| ENSG00000162654.9 | GBP4 | -1.71372 | 2.46E-10 |
| ENSG00000162692.12 | VCAM1 | -2.15458 | 8.17E-24 |
| ENSG00000162702.8 | ZNF281 | 1.090182 | 2.48E-17 |
| ENSG00000162704.16 | ARPC5 | 0.728414 | 4.32E-09 |
| ENSG00000162745.11 | OLFML2B | -0.831 | 0.007095 |
| ENSG00000162772.17 | ATF3 | 1.761805 | 3.61E-08 |
| ENSG00000162804.14 | SNED1 | -0.80606 | 2.37E-05 |
| ENSG00000162944.11 | RFTN2 | -0.82893 | 2.45E-07 |
| ENSG00000163017.14 | ACTG2 | 3.179504 | 1.85E-08 |
| ENSG00000163050.18 | COQ8A | -1.03648 | 3.38E-11 |
| ENSG00000163110.15 | PDLIM5 | 0.953689 | 1.32E-09 |
| ENSG00000163170.12 | BOLA3 | 0.650722 | 0.002889 |
| ENSG00000163376.11 | KBTBD8 | 0.740199 | 0.000464 |
| ENSG00000163378.14 | EOGT | 0.765326 | 2.26E-12 |
| ENSG00000163513.19 | TGFBR2 | -0.6009 | 2.24E-05 |
| ENSG00000163545.11 | NUAK2 | 0.916779 | 3.46E-07 |
| ENSG00000163814.8 | CDCP1 | 0.925963 | 2.90E-07 |
| ENSG00000163827.14 | LRRC2 | 1.328253 | 1.23E-07 |
| ENSG00000163879.11 | DNALI1 | -0.74054 | 0.013913 |
| ENSG00000163884.4 | KLF15 | -1.15703 | 0.00795 |
| ENSG00000163947.12 | ARHGEF3 | -1.14901 | 1.23E-33 |
| ENSG00000163975.12 | MELTF | 0.69835 | 0.00023 |
| ENSG00000164035.10 | EMCN | -3.53536 | 6.76E-08 |
| ENSG00000164056.11 | SPRY1 | -0.72429 | 9.26E-06 |
| ENSG00000164066.13 | INTU | 0.743978 | 9.20E-09 |
| ENSG00000164093.17 | PITX2 | -0.73242 | 2.26E-05 |

| | | | |
|---|---|---|---|
| ENSG00000164161.10 | HHIP | -1.65894 | 6.91E-28 |
| ENSG00000164211.13 | STARD4 | 0.605074 | 1.54E-05 |
| ENSG00000164220.7 | F2RL2 | 0.640446 | 2.80E-07 |
| ENSG00000164440.15 | TXLNB | -2.29931 | 3.43E-29 |
| ENSG00000164442.10 | CITED2 | 0.861424 | 3.20E-06 |
| ENSG00000164483.17 | SAMD3 | -0.84296 | 0.002312 |
| ENSG00000164484.12 | TMEM200A | -0.59146 | 0.000224 |
| ENSG00000164535.15 | DAGLB | 0.64931 | 1.97E-07 |
| ENSG00000164620.9 | RELL2 | 0.730563 | 0.000108 |
| ENSG00000164663.15 | USP49 | -0.78081 | 0.000702 |
| ENSG00000164674.17 | SYTL3 | -0.85865 | 0.008713 |
| ENSG00000164736.6 | SOX17 | -0.58842 | 0.032838 |
| ENSG00000164741.15 | DLC1 | 0.65717 | 0.000698 |
| ENSG00000164761.9 | TNFRSF11B | 0.657546 | 3.05E-05 |
| ENSG00000164849.10 | GPR146 | -1.05038 | 0.033555 |
| ENSG00000164920.9 | OSR2 | -1.96103 | 8.50E-06 |
| ENSG00000164970.15 | FAM219A | 0.636902 | 5.60E-07 |
| ENSG00000165271.17 | NOL6 | 0.594057 | 5.46E-08 |
| ENSG00000165410.15 | CFL2 | 0.744319 | 1.11E-14 |
| ENSG00000165424.7 | ZCCHC24 | -0.93688 | 2.35E-26 |
| ENSG00000165527.7 | ARF6 | 0.734178 | 1.66E-11 |
| ENSG00000165617.15 | DACT1 | 1.182886 | 9.94E-09 |
| ENSG00000165661.17 | QSOX2 | 0.668735 | 4.65E-06 |
| ENSG00000165732.13 | DDX21 | 0.830724 | 9.78E-10 |
| ENSG00000165795.24 | NDRG2 | -1.28818 | 1.39E-09 |
| ENSG00000165996.14 | HACD1 | 1.164785 | 4.56E-07 |
| ENSG00000166002.7 | SMCO4 | 0.945715 | 0.004431 |
| ENSG00000166016.6 | ABTB2 | 0.596494 | 4.99E-05 |
| ENSG00000166046.11 | TCP11L2 | -0.65036 | 4.82E-10 |
| ENSG00000166106.4 | ADAMTS15 | -1.09863 | 1.13E-12 |
| ENSG00000166123.14 | GPT2 | 0.830409 | 0.000506 |
| ENSG00000166262.16 | FAM227B | -0.77472 | 0.045314 |
| ENSG00000166387.14 | PPFIBP2 | -0.58815 | 1.25E-05 |
| ENSG00000166483.12 | WEE1 | 0.723645 | 1.93E-09 |
| ENSG00000166562.9 | SEC11C | 1.148468 | 1.27E-12 |
| ENSG00000166579.16 | NDEL1 | 0.604211 | 8.27E-09 |
| ENSG00000166582.10 | CENPV | 1.321217 | 2.03E-12 |
| ENSG00000166670.10 | MMP10 | -1.79364 | 1.18E-08 |
| ENSG00000166741.8 | NNMT | 0.624573 | 0.011044 |
| ENSG00000166780.11 | BMERB1 | -0.64083 | 2.95E-07 |
| ENSG00000166793.13 | YPEL4 | -1.13169 | 0.002033 |
| ENSG00000166839.17 | ANKDD1A | -0.60148 | 0.047262 |
| ENSG00000166920.13 | C15orf48 | 2.558974 | 7.45E-08 |
| ENSG00000166922.8 | SCG5 | 1.57469 | 1.11E-12 |
| ENSG00000166923.12 | GREM1 | 1.146592 | 1.09E-39 |
| ENSG00000166949.17 | SMAD3 | -0.6424 | 3.98E-07 |
| ENSG00000166986.15 | MARS1 | 0.700073 | 0.00116 |
| ENSG00000167371.21 | PRRT2 | -1.25343 | 1.09E-15 |
| ENSG00000167460.17 | TPM4 | 0.860239 | 4.84E-13 |
| ENSG00000167549.19 | CORO6 | -0.68411 | 0.007595 |
| ENSG00000167645.17 | YIF1B | 0.685961 | 0.000373 |
| ENSG00000167657.14 | DAPK3 | 0.760709 | 9.62E-13 |
| ENSG00000167703.15 | SLC43A2 | -0.85596 | 5.78E-10 |
| ENSG00000167705.12 | RILP | -0.76546 | 0.00125 |
| ENSG00000167733.14 | HSD11B1L | -0.62182 | 0.009426 |
| ENSG00000167778.9 | SPRYD3 | 0.648831 | 4.22E-11 |
| ENSG00000167797.8 | CDK2AP2 | 0.7246 | 1.09E-10 |
| ENSG00000167964.13 | RAB26 | -1.534 | 0.000153 |
| ENSG00000167992.13 | VWCE | -0.92005 | 0.001203 |
| ENSG00000167994.13 | RAB3IL1 | -0.59904 | 1.77E-06 |
| ENSG00000168286.3 | THAP11 | -0.74165 | 2.39E-08 |
| ENSG00000168306.13 | ACOX2 | -0.64231 | 0.000156 |
| ENSG00000168374.11 | ARF4 | 0.736176 | 2.62E-09 |

| | | | |
|---|---|---|---|
| ENSG00000168386.19 | FILIP1L | 1.43541 | 8.92E-15 |
| ENSG00000168389.18 | MFSD2A | 1.501575 | 0.000635 |
| ENSG00000168517.11 | HEXIM2 | -1.05142 | 0.000174 |
| ENSG00000168575.10 | SLC20A2 | 1.582797 | 1.41E-24 |
| ENSG00000168621.15 | GDNF | 1.933524 | 1.48E-71 |
| ENSG00000168646.13 | AXIN2 | -0.75981 | 6.78E-05 |
| ENSG00000168765.17 | GSTM4 | -0.66361 | 5.24E-06 |
| ENSG00000168824.14 | NSG1 | -0.89825 | 0.000336 |
| ENSG00000168916.16 | ZNF608 | -1.01604 | 1.71E-05 |
| ENSG00000168917.9 | SLC35G2 | 0.586689 | 0.006303 |
| ENSG00000168961.17 | LGALS9 | -0.83133 | 0.006415 |
| ENSG00000168994.14 | PXDC1 | 0.998587 | 3.06E-06 |
| ENSG00000169122.11 | FAM110B | -0.80906 | 0.002793 |
| ENSG00000169359.16 | SLC33A1 | 0.737628 | 1.04E-15 |
| ENSG00000169439.12 | SDC2 | 0.728577 | 4.77E-08 |
| ENSG00000169715.15 | MT1E | -0.77701 | 0.045516 |
| ENSG00000169744.13 | LDB2 | -1.07046 | 1.80E-18 |
| ENSG00000169756.16 | LIMS1 | 0.604969 | 1.02E-12 |
| ENSG00000169857.9 | AVEN | 0.874626 | 5.25E-05 |
| ENSG00000169908.12 | TM4SF1 | -0.64224 | 1.59E-09 |
| ENSG00000169991.11 | IFFO2 | 0.731619 | 4.32E-10 |
| ENSG00000170006.12 | TMEM154 | 0.694645 | 3.96E-05 |
| ENSG00000170017.12 | ALCAM | 0.721352 | 3.18E-10 |
| ENSG00000170271.11 | FAXDC2 | -0.76132 | 2.75E-10 |
| ENSG00000170345.10 | FOS | -1.16126 | 0.005842 |
| ENSG00000170485.17 | NPAS2 | 0.62375 | 2.66E-07 |
| ENSG00000170624.14 | SGCD | -0.6142 | 4.28E-06 |
| ENSG00000170801.10 | HTRA3 | -1.20708 | 9.71E-09 |
| ENSG00000170873.19 | MTSS1 | -0.65304 | 4.92E-10 |
| ENSG00000170961.7 | HAS2 | 1.226891 | 0.001306 |
| ENSG00000170962.13 | PDGFD | -1.17276 | 5.70E-17 |
| ENSG00000171067.11 | C11orf24 | 0.93154 | 1.06E-16 |
| ENSG00000171150.9 | SOCS5 | -0.66246 | 9.27E-12 |
| ENSG00000171310.11 | CHST11 | -0.66656 | 0.004174 |
| ENSG00000171522.6 | PTGER4 | -0.8958 | 9.93E-05 |
| ENSG00000171617.15 | ENC1 | 1.178955 | 6.56E-22 |
| ENSG00000171793.16 | CTPS1 | 1.189208 | 3.83E-14 |
| ENSG00000171877.21 | FRMD5 | 0.615602 | 1.34E-05 |
| ENSG00000172020.13 | GAP43 | -0.84482 | 0.028887 |
| ENSG00000172057.10 | ORMDL3 | 0.680733 | 5.19E-09 |
| ENSG00000172071.15 | EIF2AK3 | 0.744119 | 1.32E-10 |
| ENSG00000172115.9 | CYCS | 0.975513 | 7.35E-14 |
| ENSG00000172137.19 | CALB2 | 0.932957 | 0.0133 |
| ENSG00000172159.16 | FRMD3 | -2.10026 | 9.70E-21 |
| ENSG00000172260.15 | NEGR1 | 1.423461 | 4.43E-33 |
| ENSG00000172296.13 | SPTLC3 | -0.82041 | 0.00141 |
| ENSG00000172399.6 | MYOZ2 | 1.685714 | 4.58E-07 |
| ENSG00000172432.19 | GTPBP2 | 0.625505 | 2.02E-05 |
| ENSG00000172594.13 | SMPDL3A | -0.66474 | 0.00062 |
| ENSG00000172733.12 | PURG | -0.76442 | 0.005089 |
| ENSG00000172738.12 | TMEM217 | 1.312338 | 9.78E-08 |
| ENSG00000172965.17 | MIR4435-2HG | 0.629587 | 8.02E-10 |
| ENSG00000173114.13 | LRRN3 | -0.94394 | 0.033555 |
| ENSG00000173207.13 | CKS1B | -0.61817 | 0.004694 |
| ENSG00000173210.20 | ABLIM3 | -0.75286 | 0.001663 |
| ENSG00000173276.14 | ZBTB21 | 0.805661 | 4.82E-08 |
| ENSG00000173334.4 | TRIB1 | 0.87518 | 6.20E-06 |
| ENSG00000173531.15 | MST1 | -0.84252 | 3.18E-07 |
| ENSG00000173559.15 | NABP1 | 0.600094 | 1.66E-05 |
| ENSG00000173641.18 | HSPB7 | 1.06129 | 0.035656 |
| ENSG00000173848.19 | NET1 | -0.76093 | 9.86E-05 |
| ENSG00000174099.12 | MSRB3 | 0.72204 | 8.17E-06 |
| ENSG00000174136.13 | RGMB | 1.006742 | 3.85E-06 |

| | | | |
|---|---|---|---|
| ENSG00000174437.18 | ATP2A2 | 0.692058 | 1.95E-17 |
| ENSG00000174718.12 | RESF1 | -0.83407 | 2.75E-10 |
| ENSG00000174804.4 | FZD4 | -1.04165 | 8.46E-12 |
| ENSG00000174851.16 | YIF1A | 0.619619 | 3.94E-05 |
| ENSG00000174939.11 | ASPHD1 | 0.823748 | 0.000204 |
| ENSG00000175066.16 | GK5 | 0.714707 | 3.80E-10 |
| ENSG00000175287.19 | PHYHD1 | -0.82939 | 0.000102 |
| ENSG00000175471.19 | MCTP1 | -0.84409 | 0.010617 |
| ENSG00000175592.9 | FOSL1 | 0.905559 | 2.40E-05 |
| ENSG00000175745.14 | NR2F1 | -1.06436 | 3.06E-12 |
| ENSG00000175874.10 | CREG2 | -3.00788 | 7.99E-06 |
| ENSG00000176170.14 | SPHK1 | 1.325956 | 7.75E-28 |
| ENSG00000176438.13 | SYNE3 | -1.26842 | 8.11E-16 |
| ENSG00000176692.8 | FOXC2 | 1.146588 | 1.72E-08 |
| ENSG00000176697.20 | BDNF | 1.6663 | 1.03E-20 |
| ENSG00000176720.6 | BOK | 0.646184 | 0.000357 |
| ENSG00000176723.10 | ZNF843 | -0.6316 | 0.049945 |
| ENSG00000176771.17 | NCKAP5 | 1.018734 | 4.82E-10 |
| ENSG00000176871.9 | WSB2 | 0.639922 | 1.63E-12 |
| ENSG00000176907.5 | TCIM | -0.71149 | 0.032654 |
| ENSG00000176912.5 | TYMSOS | -1.45504 | 0.00892 |
| ENSG00000177042.16 | TMEM80 | -0.60001 | 0.001032 |
| ENSG00000177335.11 | C8orf31 | -1.24728 | 0.028565 |
| ENSG00000177374.13 | HIC1 | -0.75683 | 6.11E-07 |
| ENSG00000177425.11 | PAWR | 0.896201 | 2.66E-07 |
| ENSG00000178662.16 | CSRNP3 | -0.97383 | 3.49E-07 |
| ENSG00000178726.7 | THBD | -2.08958 | 9.75E-18 |
| ENSG00000178773.15 | CPNE7 | 0.886058 | 0.004361 |
| ENSG00000178776.5 | C5orf46 | 2.031038 | 0.012664 |
| ENSG00000178878.13 | APOLD1 | 0.696456 | 0.005519 |
| ENSG00000178922.18 | HYI | 0.786336 | 4.31E-11 |
| ENSG00000179071.5 | CCDC89 | -0.7694 | 0.030985 |
| ENSG00000179104.9 | TMTC2 | -0.74202 | 5.98E-12 |
| ENSG00000179242.16 | CDH4 | 0.955746 | 0.000501 |
| ENSG00000179604.10 | CDC42EP4 | -0.66813 | 3.40E-05 |
| ENSG00000179776.19 | CDH5 | -0.84733 | 3.46E-11 |
| ENSG00000179820.16 | MYADM | 0.872449 | 6.41E-16 |
| ENSG00000179833.4 | SERTAD2 | 0.605549 | 1.35E-09 |
| ENSG00000179965.12 | ZNF771 | -0.59495 | 0.007228 |
| ENSG00000179981.11 | TSHZ1 | -0.73678 | 5.29E-09 |
| ENSG00000180263.14 | FGD6 | -0.5936 | 2.44E-09 |
| ENSG00000180425.11 | C11orf71 | -0.72049 | 0.027915 |
| ENSG00000180440.4 | SERTM1 | -3.06079 | 6.46E-13 |
| ENSG00000180447.7 | GAS1 | -1.36166 | 4.31E-11 |
| ENSG00000180537.13 | RNF182 | 0.673766 | 0.005499 |
| ENSG00000180611.7 | MB21D2 | 0.757421 | 0.000109 |
| ENSG00000180769.10 | WDFY3-AS2 | -0.77075 | 0.002716 |
| ENSG00000180801.14 | ARSJ | 2.041118 | 4.51E-36 |
| ENSG00000180875.5 | GREM2 | -1.14441 | 0.005269 |
| ENSG00000180881.19 | CAPS2 | -1.12082 | 0.004389 |
| ENSG00000180884.10 | ZNF792 | -0.89752 | 4.48E-05 |
| ENSG00000180914.11 | OXTR | 1.822995 | 0.021856 |
| ENSG00000181218.5 | H2AW | -0.63702 | 0.016536 |
| ENSG00000181444.13 | ZNF467 | -0.92198 | 7.19E-06 |
| ENSG00000181649.8 | PHLDA2 | 0.986078 | 2.09E-12 |
| ENSG00000181804.15 | SLC9A9 | -1.48433 | 6.61E-15 |
| ENSG00000182054.10 | IDH2 | 0.616299 | 4.32E-08 |
| ENSG00000182165.18 | TP53TG1 | -0.75571 | 0.000147 |
| ENSG00000182175.14 | RGMA | -1.26694 | 1.35E-08 |
| ENSG00000182179.13 | UBA7 | -0.6623 | 7.25E-11 |
| ENSG00000182185.18 | RAD51B | -0.66595 | 0.048985 |
| ENSG00000182326.15 | C1S | -0.59603 | 7.02E-10 |
| ENSG00000182534.14 | MXRA7 | 0.618485 | 1.29E-06 |

| | | | |
|---|---|---|---|
| ENSG00000182568.17 | SATB1 | -0.62368 | 2.70E-05 |
| ENSG00000182585.10 | EPGN | 2.113038 | 1.14E-07 |
| ENSG00000182621.18 | PLCB1 | -0.76598 | 9.90E-08 |
| ENSG00000182752.10 | PAPPA | 1.178666 | 3.23E-14 |
| ENSG00000182796.14 | TMEM198B | -0.69332 | 2.43E-06 |
| ENSG00000183010.17 | PYCR1 | 0.955219 | 6.49E-07 |
| ENSG00000183087.15 | GAS6 | 0.772105 | 4.61E-11 |
| ENSG00000183098.11 | GPC6 | -0.7366 | 1.55E-16 |
| ENSG00000183111.12 | ARHGEF37 | -0.66315 | 0.002913 |
| ENSG00000183605.17 | SFXN4 | 0.61426 | 0.000909 |
| ENSG00000183696.14 | UPP1 | 0.681689 | 0.037752 |
| ENSG00000183715.14 | OPCML | 1.233898 | 0.001402 |
| ENSG00000183722.9 | LHFPL6 | -0.59043 | 0.000525 |
| ENSG00000183876.9 | ARSI | 1.011121 | 3.55E-11 |
| ENSG00000183963.19 | SMTN | 0.726466 | 6.74E-11 |
| ENSG00000184260.6 | H2AC20 | -0.59452 | 1.65E-06 |
| ENSG00000184270.6 | H2AC21 | -0.8528 | 0.001881 |
| ENSG00000184357.5 | H1-5 | -0.79529 | 0.022075 |
| ENSG00000184545.11 | DUSP8 | 2.499197 | 6.07E-17 |
| ENSG00000184575.12 | XPOT | 0.640657 | 0.00151 |
| ENSG00000184916.9 | JAG2 | -0.78007 | 0.001265 |
| ENSG00000184985.16 | SORCS2 | -0.77052 | 0.00898 |
| ENSG00000184988.8 | TMEM106A | 0.868019 | 1.98E-05 |
| ENSG00000184992.13 | BRI3BP | 0.812358 | 4.89E-06 |
| ENSG00000185010.15 | F8 | -0.69142 | 0.000977 |
| ENSG00000185015.8 | CA13 | 0.778443 | 0.00487 |
| ENSG00000185052.13 | SLC24A3 | 0.761495 | 1.86E-05 |
| ENSG00000185070.12 | FLRT2 | -0.65298 | 3.40E-08 |
| ENSG00000185261.15 | KIAA0825 | -0.78664 | 8.43E-05 |
| ENSG00000185338.7 | SOCS1 | -0.8174 | 0.016004 |
| ENSG00000185339.9 | TCN2 | -0.68907 | 0.003109 |
| ENSG00000185432.12 | METTL7A | -2.51335 | 2.26E-34 |
| ENSG00000185561.10 | TLCD2 | -0.7182 | 0.01285 |
| ENSG00000185565.12 | LSAMP | -1.16401 | 0.003389 |
| ENSG00000185614.7 | INKA1 | -0.84831 | 7.71E-05 |
| ENSG00000185634.12 | SHC4 | -1.42687 | 4.02E-07 |
| ENSG00000185697.17 | MYBL1 | 1.579363 | 1.16E-08 |
| ENSG00000185745.10 | IFIT1 | -1.61354 | 7.27E-19 |
| ENSG00000185760.17 | KCNQ5 | 0.638098 | 0.014205 |
| ENSG00000185803.12 | SLC52A2 | 0.624268 | 6.64E-06 |
| ENSG00000185885.17 | IFITM1 | -1.18545 | 4.34E-14 |
| ENSG00000185909.15 | KLHDC8B | -0.63894 | 1.18E-10 |
| ENSG00000185920.17 | PTCH1 | -0.77003 | 2.64E-07 |
| ENSG00000185972.6 | CCIN | 1.183229 | 0.00497 |
| ENSG00000186153.18 | WWOX | -0.75722 | 0.001154 |
| ENSG00000186469.8 | GNG2 | -0.63866 | 0.002871 |
| ENSG00000186470.14 | BTN3A2 | -0.7088 | 0.000211 |
| ENSG00000186567.13 | CEACAM19 | 0.823658 | 0.000947 |
| ENSG00000186575.19 | NF2 | 1.159767 | 1.42E-23 |
| ENSG00000186594.14 | MIR22HG | 0.684068 | 3.96E-05 |
| ENSG00000186615.11 | KTN1-AS1 | -0.64953 | 0.038833 |
| ENSG00000186907.8 | RTN4RL2 | -0.70633 | 0.031569 |
| ENSG00000187134.14 | AKR1C1 | -1.68413 | 4.91E-15 |
| ENSG00000187164.20 | SHTN1 | 0.832587 | 1.68E-09 |
| ENSG00000187266.14 | EPOR | -0.69072 | 0.000371 |
| ENSG00000187498.16 | COL4A1 | 1.043364 | 3.68E-10 |
| ENSG00000187720.14 | THSD4 | 1.598977 | 9.75E-25 |
| ENSG00000187840.5 | EIF4EBP1 | 0.73555 | 0.000731 |
| ENSG00000187955.12 | COL14A1 | -2.00539 | 0.000758 |
| ENSG00000188112.9 | C6orf132 | 1.027257 | 1.71E-05 |
| ENSG00000188211.9 | NCR3LG1 | 0.596453 | 9.81E-07 |
| ENSG00000188290.11 | HES4 | 1.335077 | 0.000585 |
| ENSG00000188312.14 | CENPP | -1.10367 | 2.37E-10 |

| | | | |
|---|---|---|---|
| ENSG00000188313.13 | PLSCR1 | -0.87126 | 3.89E-06 |
| ENSG00000188522.15 | FAM83G | 0.700222 | 6.63E-13 |
| ENSG00000188641.14 | DPYD | -0.62896 | 1.22E-07 |
| ENSG00000189159.16 | JPT1 | 0.610318 | 1.87E-08 |
| ENSG00000189184.12 | PCDH18 | -0.89622 | 9.75E-25 |
| ENSG00000189320.9 | FAM180A | -1.39458 | 0.000321 |
| ENSG00000189410.12 | SH2D5 | 1.827262 | 8.59E-36 |
| ENSG00000196104.11 | SPOCK3 | -0.80913 | 0.0028 |
| ENSG00000196123.13 | KIAA0895L | -0.58856 | 0.000103 |
| ENSG00000196139.14 | AKR1C3 | -1.91435 | 2.69E-19 |
| ENSG00000196154.12 | S100A4 | -0.90964 | 0.04857 |
| ENSG00000196155.13 | PLEKHG4 | 0.660241 | 0.014233 |
| ENSG00000196305.19 | IARS1 | 0.729641 | 4.51E-05 |
| ENSG00000196352.16 | CD55 | 0.753058 | 1.75E-10 |
| ENSG00000196358.11 | NTNG2 | -0.75756 | 0.001651 |
| ENSG00000196449.4 | YRDC | 0.860522 | 2.37E-05 |
| ENSG00000196460.14 | RFX8 | -1.10174 | 0.030063 |
| ENSG00000196502.12 | SULT1A1 | -0.66177 | 0.024301 |
| ENSG00000196628.20 | TCF4 | -0.8134 | 2.64E-10 |
| ENSG00000196747.4 | H2AC13 | -0.82052 | 0.002999 |
| ENSG00000196782.12 | MAML3 | -1.28773 | 4.38E-34 |
| ENSG00000196787.3 | H2AC11 | -0.78904 | 2.25E-06 |
| ENSG00000196843.17 | ARID5A | 0.873296 | 0.000219 |
| ENSG00000196923.14 | PDLIM7 | 0.890678 | 7.19E-06 |
| ENSG00000196924.19 | FLNA | 0.807321 | 1.01E-05 |
| ENSG00000197256.11 | KANK2 | -0.58701 | 6.96E-10 |
| ENSG00000197321.15 | SVIL | -0.66181 | 1.49E-06 |
| ENSG00000197381.17 | ADARB1 | 0.647647 | 0.000138 |
| ENSG00000197461.13 | PDGFA | 0.597857 | 0.037665 |
| ENSG00000197467.17 | COL13A1 | -0.81406 | 7.02E-08 |
| ENSG00000197558.13 | SSPOP | -1.2053 | 2.37E-05 |
| ENSG00000197582.5 | GPX1P1 | -1.08829 | 0.037886 |
| ENSG00000197586.13 | ENTPD6 | 0.805418 | 4.27E-16 |
| ENSG00000197632.9 | SERPINB2 | 1.798446 | 0.023134 |
| ENSG00000197646.8 | PDCD1LG2 | 2.717688 | 2.01E-62 |
| ENSG00000197780.10 | TAF13 | 0.732805 | 2.84E-12 |
| ENSG00000197785.14 | ATAD3A | 0.733101 | 1.00E-06 |
| ENSG00000197860.10 | SGTB | 0.945178 | 8.98E-14 |
| ENSG00000197977.4 | ELOVL2 | -0.63092 | 0.016059 |
| ENSG00000198018.7 | ENTPD7 | 0.834855 | 2.92E-12 |
| ENSG00000198053.12 | SIRPA | 0.586354 | 0.000143 |
| ENSG00000198074.10 | AKR1B10 | -1.97272 | 4.56E-08 |
| ENSG00000198121.14 | LPAR1 | -1.1667 | 4.64E-16 |
| ENSG00000198208.12 | RPS6KL1 | 1.070802 | 0.043199 |
| ENSG00000198286.10 | CARD11 | 0.643099 | 0.007165 |
| ENSG00000198380.13 | GFPT1 | 0.641009 | 4.90E-07 |
| ENSG00000198467.16 | TPM2 | 0.644581 | 0.003622 |
| ENSG00000198624.13 | CCDC69 | -0.67868 | 0.00029 |
| ENSG00000198682.13 | PAPSS2 | 0.800475 | 4.67E-14 |
| ENSG00000198795.11 | ZNF521 | -1.12978 | 7.78E-08 |
| ENSG00000198796.7 | ALPK2 | 1.112879 | 1.00E-05 |
| ENSG00000198805.12 | PNP | 0.697694 | 0.003423 |
| ENSG00000198843.13 | SELENOT | 0.624422 | 0.001254 |
| ENSG00000198853.12 | RUSC2 | 0.655572 | 0.000663 |
| ENSG00000198855.7 | FICD | 0.879075 | 2.96E-10 |
| ENSG00000198937.9 | CCDC167 | 0.634462 | 0.000854 |
| ENSG00000198948.12 | MFAP3L | 0.777507 | 4.07E-07 |
| ENSG00000203497.2 | PDCD4-AS1 | -1.46172 | 0.004647 |
| ENSG00000203880.12 | PCMTD2 | -0.62231 | 3.87E-08 |
| ENSG00000204131.9 | NHSL2 | -0.85018 | 0.001026 |
| ENSG00000204209.13 | DAXX | 0.61628 | 7.19E-06 |
| ENSG00000204311.15 | PJVK | -0.90567 | 0.026772 |
| ENSG00000204314.12 | PRRT1 | -0.94406 | 0.010944 |

| | | | |
|---|---|---|---|
| ENSG00000204381.12 | LAYN | 0.770984 | 3.01E-08 |
| ENSG00000204394.13 | VARS1 | 0.629634 | 8.58E-11 |
| ENSG00000204634.13 | TBC1D8 | -0.61624 | 0.013828 |
| ENSG00000204642.14 | HLA-F | -0.71276 | 0.024229 |
| ENSG00000204941.14 | PSG5 | 2.305885 | 1.30E-05 |
| ENSG00000205464.13 | ATP6AP1L | -0.65893 | 0.044251 |
| ENSG00000205795.4 | CYS1 | -0.90518 | 0.007704 |
| ENSG00000205978.6 | NYNRIN | -0.81539 | 4.99E-11 |
| ENSG00000206190.12 | ATP10A | 1.269723 | 0.000488 |
| ENSG00000206417.8 | H1-10-AS1 | -1.29489 | 0.020281 |
| ENSG00000206538.9 | VGLL3 | 1.00681 | 2.82E-07 |
| ENSG00000206588.1 | RNU1-28P | 0.92468 | 0.035179 |
| ENSG00000206596.1 | RNU1-27P | 0.92468 | 0.035179 |
| ENSG00000206652.1 | RNU1-1 | 0.92468 | 0.035179 |
| ENSG00000206737.1 | RNVU1-18 | 0.92468 | 0.035179 |
| ENSG00000207005.1 | RNU1-2 | 0.92468 | 0.035179 |
| ENSG00000207389.1 | RNU1-4 | 0.92468 | 0.035179 |
| ENSG00000207513.1 | RNU1-3 | 0.92468 | 0.035179 |
| ENSG00000211455.8 | STK38L | 1.040683 | 4.70E-16 |
| ENSG00000212232.1 | SNORD17 | 0.718864 | 0.031387 |
| ENSG00000212694.8 | LINC01089 | -0.7421 | 2.84E-05 |
| ENSG00000212864.3 | RNF208 | -1.01498 | 0.001869 |
| ENSG00000213523.12 | SRA1 | 0.650219 | 3.02E-06 |
| ENSG00000213760.11 | ATP6V1G2 | -1.12739 | 0.000472 |
| ENSG00000214212.9 | C19orf38 | -1.19227 | 0.020863 |
| ENSG00000214706.12 | IFRD2 | 0.762966 | 5.12E-07 |
| ENSG00000215068.8 | AC025171.1 | -0.7456 | 4.05E-05 |
| ENSG00000215861.6 | WI2-1896O14.1 | 1.103724 | 7.38E-10 |
| ENSG00000215883.11 | CYB5RL | -0.67097 | 5.00E-06 |
| ENSG00000217801.10 | RP11-465B22.3 | 2.489672 | 4.06E-05 |
| ENSG00000221818.9 | EBF2 | -1.62488 | 3.61E-15 |
| ENSG00000221821.4 | C6orf226 | -1.04333 | 0.005819 |
| ENSG00000221968.9 | FADS3 | 0.698431 | 7.94E-13 |
| ENSG00000222041.11 | CYTOR | 0.832026 | 3.41E-11 |
| ENSG00000223403.6 | MEG9 | 0.744121 | 0.001121 |
| ENSG00000223485.4 | LINC01615 | 0.985066 | 0.01416 |
| ENSG00000223745.8 | CCDC18-AS1 | -0.64563 | 0.004266 |
| ENSG00000223802.9 | CERS1 | -0.73865 | 4.58E-05 |
| ENSG00000224715.1 | CITF22-49D8.1 | -1.9361 | 0.011659 |
| ENSG00000225614.4 | ZNF469 | 0.758289 | 9.94E-12 |
| ENSG00000225783.8 | MIAT | -0.81675 | 7.84E-06 |
| ENSG00000225855.7 | RUSC1-AS1 | -0.99525 | 0.040912 |
| ENSG00000226380.9 | AC058791.1 | 0.791864 | 0.002256 |
| ENSG00000227051.7 | C14orf132 | -0.92079 | 5.77E-15 |
| ENSG00000227268.5 | KLLN | -0.80562 | 0.001939 |
| ENSG00000227496.3 | RP11-145A3.1 | 2.238592 | 5.54E-05 |
| ENSG00000228451.4 | SDAD1P1 | -0.61806 | 0.031606 |
| ENSG00000229124.7 | VIM-AS1 | -0.65254 | 0.012811 |
| ENSG00000229619.4 | MBNL1-AS1 | 0.815961 | 0.00113 |
| ENSG00000229809.9 | ZNF688 | -0.67975 | 0.006939 |
| ENSG00000232533.1 | AC093673.5 | 0.606145 | 0.040244 |
| ENSG00000232859.10 | LYRM9 | -0.65508 | 0.006382 |
| ENSG00000233098.9 | CCDC144NL-AS1 | 1.501013 | 9.96E-09 |
| ENSG00000233117.4 | LINC00702 | 0.808053 | 0.008405 |
| ENSG00000233178.7 | RP11-88I18.2 | -0.63833 | 0.013639 |
| ENSG00000233223.3 | AC113189.5 | -1.01989 | 0.017923 |
| ENSG00000233695.2 | GAS6-AS1 | -1.48651 | 1.89E-20 |
| ENSG00000234857.2 | HNRNPUL2-BSCL2 | 0.662794 | 0.002451 |
| ENSG00000235162.9 | C12orf75 | 0.624051 | 7.32E-05 |
| ENSG00000235217.6 | TSPY26P | -0.81757 | 1.62E-05 |
| ENSG00000235863.4 | B3GALT4 | -1.15278 | 0.001829 |
| ENSG00000237037.9 | NDUFA6-DT | -0.81046 | 0.000373 |
| ENSG00000237352.4 | LINC01358 | -0.88977 | 0.029271 |

| | | | |
|---|---|---|---|
| ENSG00000237765.8 | FAM200B | 0.590742 | 4.31E-07 |
| ENSG00000237807.4 | RP11-400K9.4 | -0.67974 | 1.80E-06 |
| ENSG00000238917.1 | SNORD10 | 1.016755 | 0.010688 |
| ENSG00000239332.6 | LINC01119 | -0.97327 | 0.016264 |
| ENSG00000239521.9 | CASTOR3 | -0.62568 | 4.40E-07 |
| ENSG00000239672.8 | NME1 | 0.823904 | 7.48E-14 |
| ENSG00000239704.11 | CDRT4 | 0.901345 | 0.044119 |
| ENSG00000240038.7 | AMY2B | -0.72026 | 0.034622 |
| ENSG00000240694.9 | PNMA2 | 0.713909 | 0.018986 |
| ENSG00000241399.7 | CD302 | -0.70377 | 6.93E-09 |
| ENSG00000242242.6 | NECTIN3-AS1 | 2.932318 | 1.44E-08 |
| ENSG00000242265.6 | PEG10 | -0.94102 | 1.86E-14 |
| ENSG00000243137.8 | PSG4 | 2.274487 | 2.22E-14 |
| ENSG00000243156.9 | MICAL3 | 0.646049 | 3.55E-11 |
| ENSG00000243244.7 | STON1 | -0.87776 | 7.77E-12 |
| ENSG00000243649.9 | CFB | -0.86296 | 0.001046 |
| ENSG00000243811.12 | APOBEC3D | -0.75555 | 0.043106 |
| ENSG00000244026.6 | FAM86DP | 0.740558 | 0.000162 |
| ENSG00000244270.1 | RPL32P29 | -0.74933 | 0.043533 |
| ENSG00000244694.8 | PTCHD4 | -0.68828 | 2.57E-05 |
| ENSG00000245573.9 | BDNF-AS | -1.07936 | 0.003396 |
| ENSG00000246430.7 | LINC00968 | 1.755591 | 0.00202 |
| ENSG00000246898.1 | LINC00920 | -1.33389 | 0.000493 |
| ENSG00000247271.7 | ZBED5-AS1 | -0.74568 | 0.003242 |
| ENSG00000247746.5 | USP51 | -0.95426 | 6.22E-05 |
| ENSG00000247950.7 | SEC24B-AS1 | -0.94139 | 0.04925 |
| ENSG00000248441.7 | LINC01197 | -1.22448 | 0.007109 |
| ENSG00000248890.2 | HHIP-AS1 | -1.20059 | 2.47E-06 |
| ENSG00000249087.7 | ZNF436-AS1 | -0.75449 | 0.039827 |
| ENSG00000249279.6 | LINC02057 | 1.41645 | 2.94E-06 |
| ENSG00000249669.10 | CARMN | 0.722555 | 2.30E-11 |
| ENSG00000250320.6 | EDIL3-DT | 0.805637 | 0.002554 |
| ENSG00000250510.8 | GPR162 | -0.92082 | 2.07E-06 |
| ENSG00000250644.3 | RP11-295K3.1 | -0.82092 | 0.000712 |
| ENSG00000251493.5 | FOXD1 | 0.829202 | 7.33E-11 |
| ENSG00000251669.6 | FAM86EP | 0.648153 | 0.002101 |
| ENSG00000252010.1 | SCARNA5 | 0.988033 | 0.017463 |
| ENSG00000253661.2 | ZFHX4-AS1 | -0.77175 | 0.006494 |
| ENSG00000253982.2 | CTD-2336O2.1 | -0.82383 | 5.90E-06 |
| ENSG00000254602.2 | AP000662.4 | -0.73168 | 0.044659 |
| ENSG00000254682.1 | RP11-660L16.2 | -0.74429 | 0.042006 |
| ENSG00000254726.3 | MEX3A | -0.76671 | 9.72E-07 |
| ENSG00000255439.6 | RP11-196G11.1 | -2.82076 | 0.017922 |
| ENSG00000255471.1 | RP11-736K20.5 | -1.80963 | 4.65E-07 |
| ENSG00000255690.3 | TRIL | -2.20184 | 7.60E-09 |
| ENSG00000257108.2 | NHLRC4 | -1.02501 | 0.00252 |
| ENSG00000257219.6 | LNCOG | 1.835724 | 5.46E-05 |
| ENSG00000257453.1 | RP11-290L1.3 | -1.34827 | 0.028353 |
| ENSG00000257702.3 | LBX2-AS1 | -0.78343 | 0.018611 |
| ENSG00000258388.7 | PPT2-EGFL8 | -1.50641 | 0.006008 |
| ENSG00000258947.8 | TUBB3 | -0.7404 | 5.40E-05 |
| ENSG00000259132.1 | RP11-298I3.5 | -3.19938 | 0.000799 |
| ENSG00000259207.9 | ITGB3 | 2.309164 | 1.62E-21 |
| ENSG00000259275.4 | RP11-522B15.3 | -0.96792 | 0.002788 |
| ENSG00000259426.6 | RP11-253M7.1 | 1.49915 | 7.59E-08 |
| ENSG00000259877.2 | RP11-46C24.7 | -0.58541 | 0.028768 |
| ENSG00000260077.1 | RP11-254F7.2 | -0.77093 | 0.002152 |
| ENSG00000260196.1 | RP1-239B22.5 | 0.620902 | 0.002817 |
| ENSG00000260293.2 | RP11-715J22.6 | -0.82687 | 0.018992 |
| ENSG00000260604.2 | RP1-140K8.5 | 2.293211 | 1.98E-09 |
| ENSG00000261040.7 | WFDC21P | 0.973095 | 0.002787 |
| ENSG00000261087.1 | ZNNT1 | -0.91447 | 0.038605 |
| ENSG00000261371.6 | PECAM1 | -0.98765 | 0.000241 |

| | | | |
|---|---|---|---|
| ENSG00000261468.1 | RP11-1024P17.1 | -1.16961 | 2.93E-05 |
| ENSG00000261487.1 | AC135048.13 | -0.92722 | 0.010613 |
| ENSG00000261490.1 | RP11-448G15.3 | 0.635111 | 0.024601 |
| ENSG00000261801.6 | LOXL1-AS1 | 0.639981 | 0.001541 |
| ENSG00000263426.2 | RN7SL471P | 1.059665 | 8.06E-05 |
| ENSG00000266074.10 | BAHCC1 | -0.77763 | 3.30E-15 |
| ENSG00000266094.8 | RASSF5 | -1.09283 | 2.76E-05 |
| ENSG00000266714.9 | MYO15B | -1.08565 | 0.026641 |
| ENSG00000267002.4 | RP11-242D8.1 | -0.61425 | 0.035239 |
| ENSG00000267303.1 | CTD-2369P2.12 | 3.920773 | 1.04E-11 |
| ENSG00000267361.1 | SEC24AP1 | -1.92257 | 0.006125 |
| ENSG00000267645.5 | RP11-577H5.5 | -0.61563 | 0.003314 |
| ENSG00000269378.1 | ITGB1P1 | 0.774127 | 2.82E-05 |
| ENSG00000270149.5 | RP11-544M22.13 | -0.61219 | 0.013886 |
| ENSG00000271605.6 | MILR1 | -0.71904 | 0.001869 |
| ENSG00000272031.3 | ANKRD34A | 0.645341 | 0.012012 |
| ENSG00000272686.1 | WASL-DT | -0.69057 | 0.043926 |
| ENSG00000272695.2 | GAS6-DT | 0.823336 | 0.00128 |
| ENSG00000273213.4 | H3-2 | 0.640269 | 2.21E-06 |
| ENSG00000273542.2 | H4C12 | -0.85145 | 2.22E-14 |
| ENSG00000273768.1 | RNVU1-29 | 0.92468 | 0.035179 |
| ENSG00000274618.2 | H4C6 | -0.58526 | 1.15E-06 |
| ENSG00000275342.5 | PRAG1 | -0.94637 | 3.55E-11 |
| ENSG00000275379.2 | H3C11 | -0.76115 | 3.09E-05 |
| ENSG00000275405.1 | U1 | 0.92468 | 0.035179 |
| ENSG00000275713.2 | H2BC9 | -0.66797 | 0.0051 |
| ENSG00000275714.2 | H3C1 | -0.64096 | 0.001413 |
| ENSG00000275894.1 | RP3-453C12.14 | 1.48972 | 0.033354 |
| ENSG00000276043.5 | UHRF1 | 0.799324 | 1.77E-06 |
| ENSG00000276903.2 | H2AC16 | -1.15027 | 0.001891 |
| ENSG00000277075.2 | H2AC8 | -0.61236 | 0.000331 |
| ENSG00000277224.2 | H2BC7 | -0.62912 | 0.014605 |
| ENSG00000277775.2 | H3C7 | -0.99228 | 0.000551 |
| ENSG00000277778.2 | PGM5P2 | 1.223627 | 0.001115 |
| ENSG00000278249.1 | SCARNA2 | -0.62205 | 2.70E-07 |
| ENSG00000278463.2 | H2AC4 | -0.88749 | 0.000286 |
| ENSG00000278637.2 | H4C1 | -0.68877 | 0.00589 |
| ENSG00000278677.2 | H2AC17 | -0.64849 | 6.14E-05 |
| ENSG00000278828.1 | H3C10 | -0.72654 | 0.000116 |
| ENSG00000278948.1 | RP5-1039K5.12 | 0.835788 | 1.02E-08 |
| ENSG00000279041.1 | CTD-2373N4.3 | 0.976593 | 0.005629 |
| ENSG00000279821.1 | RP11-1334A24.5 | 1.3827 | 3.79E-08 |
| ENSG00000280143.1 | AP000892.6 | 2.257829 | 2.17E-14 |
| ENSG00000280339.1 | RP11-736K20.4 | -1.73154 | 2.78E-09 |
| ENSG00000280441.3 | CH507-528H12.1 | -0.59017 | 0.002224 |
| ENSG00000280893.1 | AC009133.23 | -1.33483 | 0.000403 |
| ENSG00000282057.1 | RP4-621F18.2 | 0.974629 | 0.00012 |
| ENSG00000282849.1 | RP11-121P12.1 | 3.611728 | 5.85E-06 |
| ENSG00000283154.2 | IQCJ-SCHIP1 | 1.336809 | 2.79E-08 |
| ENSG00000284693.1 | LINC02606 | -0.81763 | 0.038979 |
| ENSG00000284879.1 | RP13-511M20.1 | 0.602779 | 0.001414 |
| ENSG00000285106.2 | RP11-36B6.2 | 0.631796 | 0.00505 |
| ENSG00000285283.1 | RP1-65P5.6 | 0.704775 | 0.046156 |
| ENSG00000285901.2 | RP11-388F6.5 | 1.279307 | 1.89E-05 |
| ENSG00000286190.2 | RP11-807C20.3 | 0.786834 | 4.42E-12 |
| ENSG00000287080.2 | H3C3 | -0.84366 | 0.033992 |
| ENSG00000287104.1 | RP11-326F6.1 | -0.80778 | 0.033543 |
| ENSG00000288031.1 | RP11-1L5.2 | 0.791092 | 0.03877 |
| ENSG00000288686.2 | XGY2 | 1.167652 | 0.024513 |

**Table S6. Gene Ontology analysis of genes regulated by colchicine compared to control HASMCs.** HASMCs were treated with colchicine (50nM) for 24 hours.

| GO.ID | Term | Annotated | Significant | Expected | classicFisher |
|---|---|---|---|---|---|
| GO:0008360 | regulation of cell shape | 120 | 25 | 11.23 | 0.0001 |
| GO:0032526 | response to retinoic acid | 67 | 17 | 6.27 | 0.00011 |
| GO:0045137 | development of primary sexual characteristics | 149 | 29 | 13.95 | 0.00011 |
| GO:0035025 | positive regulation of Rho protein signal transduction | 18 | 8 | 1.68 | 0.00011 |
| GO:0007389 | pattern specification process | 264 | 44 | 24.71 | 0.00011 |
| GO:1904888 | cranial skeletal system development | 49 | 14 | 4.59 | 0.00011 |
| GO:0001823 | mesonephros development | 61 | 16 | 5.71 | 0.00011 |
| GO:0055017 | cardiac muscle tissue growth | 61 | 16 | 5.71 | 0.00011 |
| GO:0015849 | organic acid transport | 164 | 31 | 15.35 | 0.00011 |
| GO:0070372 | regulation of ERK1 and ERK2 cascade | 164 | 31 | 15.35 | 0.00011 |
| GO:0050890 | cognition | 179 | 33 | 16.76 | 0.00011 |
| GO:0050807 | regulation of synapse organization | 121 | 25 | 11.33 | 0.00012 |
| GO:0043583 | ear development | 114 | 24 | 10.67 | 0.00012 |
| GO:0001759 | organ induction | 14 | 7 | 1.31 | 0.00012 |
| GO:0008584 | male gonad development | 94 | 21 | 8.8 | 0.00013 |
| GO:0051918 | negative regulation of fibrinolysis | 7 | 5 | 0.66 | 0.00013 |
| GO:0048701 | embryonic cranial skeleton morphogenesis | 33 | 11 | 3.09 | 0.00013 |
| GO:0055002 | striated muscle cell development | 33 | 11 | 3.09 | 0.00013 |
| GO:0072078 | nephron tubule morphogenesis | 44 | 13 | 4.12 | 0.00013 |
| GO:0035023 | regulation of Rho protein signal transduction | 62 | 16 | 5.8 | 0.00013 |
| GO:0032101 | regulation of response to external stimulus | 571 | 80 | 53.45 | 0.00014 |
| GO:0065009 | regulation of molecular function | 2053 | 237 | 192.17 | 0.00014 |
| GO:0010232 | vascular transport | 56 | 15 | 5.24 | 0.00014 |
| GO:0048704 | embryonic skeletal system morphogenesis | 56 | 15 | 5.24 | 0.00014 |
| GO:0150104 | transport across blood-brain barrier | 56 | 15 | 5.24 | 0.00014 |
| GO:0060485 | mesenchyme development | 212 | 37 | 19.84 | 0.00014 |
| GO:0030030 | cell projection organization | 1035 | 131 | 96.88 | 0.00014 |
| GO:0001818 | negative regulation of cytokine production | 159 | 30 | 14.88 | 0.00015 |
| GO:0060284 | regulation of cell development | 341 | 53 | 31.92 | 0.00015 |
| GO:0046546 | development of primary male sexual characteristics | 95 | 21 | 8.89 | 0.00015 |
| GO:0045216 | cell-cell junction organization | 130 | 26 | 12.17 | 0.00015 |
| GO:0007266 | Rho protein signal transduction | 102 | 22 | 9.55 | 0.00015 |
| GO:0051403 | stress-activated MAPK cascade | 182 | 33 | 17.04 | 0.00015 |
| GO:0003206 | cardiac chamber morphogenesis | 82 | 19 | 7.68 | 0.00016 |
| GO:0042110 | T cell activation | 301 | 48 | 28.18 | 0.00016 |
| GO:0030323 | respiratory tube development | 138 | 27 | 12.92 | 0.00017 |
| GO:0090257 | regulation of muscle system process | 138 | 27 | 12.92 | 0.00017 |
| GO:0006869 | lipid transport | 245 | 41 | 22.93 | 0.00017 |
| GO:0034446 | substrate adhesion-dependent cell spreading | 89 | 20 | 8.33 | 0.00017 |
| GO:0030199 | collagen fibril organization | 51 | 14 | 4.77 | 0.00017 |
| GO:1900024 | regulation of substrate adhesion-dependent cell spreading | 51 | 14 | 4.77 | 0.00017 |
| GO:0007611 | learning or memory | 153 | 29 | 14.32 | 0.00017 |
| GO:0007267 | cell-cell signaling | 891 | 115 | 83.4 | 0.00017 |
| GO:0071674 | mononuclear cell migration | 96 | 21 | 8.99 | 0.00017 |
| GO:0090049 | regulation of cell migration involved in sprouting angiogenesis | 34 | 11 | 3.18 | 0.00017 |
| GO:0008544 | epidermis development | 168 | 31 | 15.73 | 0.00017 |
| GO:0016310 | phosphorylation | 1275 | 156 | 119.35 | 0.00017 |
| GO:0050803 | regulation of synapse structure or activity | 124 | 25 | 11.61 | 0.00017 |
| GO:0120039 | plasma membrane bounded cell projection morphogenesis | 420 | 62 | 39.31 | 0.00019 |
| GO:0035633 | maintenance of blood-brain barrier | 24 | 9 | 2.25 | 0.00019 |
| GO:0071677 | positive regulation of mononuclear cell migration | 40 | 12 | 3.74 | 0.0002 |
| GO:0050731 | positive regulation of peptidyl-tyrosine phosphorylation | 97 | 21 | 9.08 | 0.0002 |
| GO:0031639 | plasminogen activation | 15 | 7 | 1.4 | 0.0002 |
| GO:0043535 | regulation of blood vessel endothelial cell migration | 77 | 18 | 7.21 | 0.00021 |
| GO:0035924 | cellular response to vascular endothelial growth factor stimulus | 46 | 13 | 4.31 | 0.00021 |
| GO:1903036 | positive regulation of response to wounding | 46 | 13 | 4.31 | 0.00021 |

| | | | | | |
|---|---|---|---|---|---|
| GO:0002521 | leukocyte differentiation | 346 | 53 | 32.39 | 0.00021 |
| GO:0007369 | gastrulation | 133 | 26 | 12.45 | 0.00022 |
| GO:0022409 | positive regulation of cell-cell adhesion | 178 | 32 | 16.66 | 0.00023 |
| GO:0048858 | cell projection morphogenesis | 423 | 62 | 39.6 | 0.00023 |
| GO:0010876 | lipid localization | 281 | 45 | 26.3 | 0.00023 |
| GO:0045596 | negative regulation of cell differentiation | 432 | 63 | 40.44 | 0.00023 |
| GO:0080134 | regulation of response to stress | 972 | 123 | 90.99 | 0.00024 |
| GO:0046661 | male sex differentiation | 105 | 22 | 9.83 | 0.00024 |
| GO:0032102 | negative regulation of response to external stimulus | 241 | 40 | 22.56 | 0.00024 |
| GO:0007548 | sex differentiation | 171 | 31 | 16.01 | 0.00024 |
| GO:0019932 | second-messenger-mediated signaling | 149 | 28 | 13.95 | 0.00026 |
| GO:0060038 | cardiac muscle cell proliferation | 41 | 12 | 3.84 | 0.00026 |
| GO:0001657 | ureteric bud development | 59 | 15 | 5.52 | 0.00026 |
| GO:0032231 | regulation of actin filament bundle assembly | 85 | 19 | 7.96 | 0.00026 |
| GO:0010543 | regulation of platelet activation | 30 | 10 | 2.81 | 0.00026 |
| GO:0015844 | monoamine transport | 30 | 10 | 2.81 | 0.00026 |
| GO:0031098 | stress-activated protein kinase signaling cascade | 187 | 33 | 17.5 | 0.00026 |
| GO:0014910 | regulation of smooth muscle cell migration | 53 | 14 | 4.96 | 0.00027 |
| GO:0048638 | regulation of developmental growth | 211 | 36 | 19.75 | 0.00028 |
| GO:0061448 | connective tissue development | 188 | 33 | 17.6 | 0.00029 |
| GO:0007409 | axonogenesis | 284 | 45 | 26.58 | 0.00029 |
| GO:0071560 | cellular response to transforming growth factor beta stimulus | 204 | 35 | 19.1 | 0.0003 |
| GO:0090303 | positive regulation of wound healing | 36 | 11 | 3.37 | 0.00031 |
| GO:0120035 | regulation of plasma membrane bounded cell projection organization | 454 | 65 | 42.5 | 0.00032 |
| GO:0072163 | mesonephric epithelium development | 60 | 15 | 5.62 | 0.00032 |
| GO:0072164 | mesonephric tubule development | 60 | 15 | 5.62 | 0.00032 |
| GO:0014902 | myotube differentiation | 73 | 17 | 6.83 | 0.00032 |
| GO:0030282 | bone mineralization | 73 | 17 | 6.83 | 0.00032 |
| GO:0051057 | positive regulation of small GTPase mediated signal transduction | 42 | 12 | 3.93 | 0.00033 |
| GO:0002455 | humoral immune response mediated by circulating immunoglobulin | 16 | 7 | 1.5 | 0.00033 |
| GO:0060395 | SMAD protein signal transduction | 54 | 14 | 5.05 | 0.00033 |
| GO:0002685 | regulation of leukocyte migration | 129 | 25 | 12.08 | 0.00033 |
| GO:0050678 | regulation of epithelial cell proliferation | 237 | 39 | 22.18 | 0.00034 |
| GO:0031344 | regulation of cell projection organization | 464 | 66 | 43.43 | 0.00034 |
| GO:0030278 | regulation of ossification | 80 | 18 | 7.49 | 0.00034 |
| GO:0003308 | negative regulation of Wnt signaling pathway involved in heart development | 5 | 4 | 0.47 | 0.00035 |
| GO:1903131 | mononuclear cell differentiation | 270 | 43 | 25.27 | 0.00036 |
| GO:0010975 | regulation of neuron projection development | 312 | 48 | 29.21 | 0.00038 |
| GO:0010518 | positive regulation of phospholipase activity | 26 | 9 | 2.43 | 0.00038 |
| GO:0045214 | sarcomere organization | 21 | 8 | 1.97 | 0.00038 |
| GO:0001756 | somitogenesis | 37 | 11 | 3.46 | 0.0004 |
| GO:0050804 | modulation of chemical synaptic transmission | 239 | 39 | 22.37 | 0.0004 |
| GO:0060541 | respiratory system development | 153 | 28 | 14.32 | 0.0004 |
| GO:0002040 | sprouting angiogenesis | 88 | 19 | 8.24 | 0.00041 |
| GO:1902903 | regulation of supramolecular fiber organization | 297 | 46 | 27.8 | 0.00043 |
| GO:0046578 | regulation of Ras protein signal transduction | 146 | 27 | 13.67 | 0.00043 |
| GO:0071559 | response to transforming growth factor beta | 208 | 35 | 19.47 | 0.00044 |
| GO:0099177 | regulation of trans-synaptic signaling | 240 | 39 | 22.47 | 0.00044 |
| GO:0016485 | protein processing | 169 | 30 | 15.82 | 0.00044 |
| GO:0051247 | positive regulation of protein metabolic process | 1072 | 132 | 100.35 | 0.00045 |
| GO:0015850 | organic hydroxy compound transport | 139 | 26 | 13.01 | 0.00045 |
| GO:0051338 | regulation of transferase activity | 647 | 86 | 60.56 | 0.00046 |
| GO:0006940 | regulation of smooth muscle contraction | 32 | 10 | 3 | 0.00047 |
| GO:0015804 | neutral amino acid transport | 32 | 10 | 3 | 0.00047 |
| GO:0060193 | positive regulation of lipase activity | 32 | 10 | 3 | 0.00047 |
| GO:0022612 | gland morphogenesis | 82 | 18 | 7.68 | 0.00047 |
| GO:0050920 | regulation of chemotaxis | 132 | 25 | 12.36 | 0.00048 |
| GO:0030038 | contractile actin filament bundle assembly | 89 | 19 | 8.33 | 0.00048 |
| GO:0043149 | stress fiber assembly | 89 | 19 | 8.33 | 0.00048 |
| GO:0003151 | outflow tract morphogenesis | 56 | 14 | 5.24 | 0.0005 |
| GO:0001819 | positive regulation of cytokine production | 266 | 42 | 24.9 | 0.0005 |
| GO:0030194 | positive regulation of blood coagulation | 17 | 7 | 1.59 | 0.00052 |
| GO:0060561 | apoptotic process involved in morphogenesis | 17 | 7 | 1.59 | 0.00052 |

| | | | | | |
|---|---|---|---|---|---|
| GO:0061298 | retina vasculature development in camera-type eye | 17 | 7 | 1.59 | 0.00052 |
| GO:1900048 | positive regulation of hemostasis | 17 | 7 | 1.59 | 0.00052 |
| GO:0061053 | somite development | 50 | 13 | 4.68 | 0.00052 |
| GO:0150117 | positive regulation of cell-substrate junction organization | 27 | 9 | 2.53 | 0.00053 |
| GO:0048706 | embryonic skeletal system development | 76 | 17 | 7.11 | 0.00053 |
| GO:0055007 | cardiac muscle cell differentiation | 76 | 17 | 7.11 | 0.00053 |
| GO:0048660 | regulation of smooth muscle cell proliferation | 104 | 21 | 9.74 | 0.00055 |
| GO:0010927 | cellular component assembly involved in morphogenesis | 63 | 15 | 5.9 | 0.00056 |
| GO:0006937 | regulation of muscle contraction | 90 | 19 | 8.42 | 0.00056 |
| GO:0070663 | regulation of leukocyte proliferation | 141 | 26 | 13.2 | 0.00057 |
| GO:0044093 | positive regulation of molecular function | 1043 | 128 | 97.63 | 0.00063 |
| GO:0006958 | complement activation, classical pathway | 13 | 6 | 1.22 | 0.00064 |
| GO:0002088 | lens development in camera-type eye | 51 | 13 | 4.77 | 0.00064 |
| GO:0007416 | synapse assembly | 91 | 19 | 8.52 | 0.00065 |
| GO:0048812 | neuron projection morphogenesis | 405 | 58 | 37.91 | 0.00065 |
| GO:0072102 | glomerulus morphogenesis | 9 | 5 | 0.84 | 0.00065 |
| GO:0072160 | nephron tubule epithelial cell differentiation | 9 | 5 | 0.84 | 0.00065 |
| GO:0015837 | amine transport | 39 | 11 | 3.65 | 0.00066 |
| GO:0051952 | regulation of amine transport | 39 | 11 | 3.65 | 0.00066 |
| GO:0051302 | regulation of cell division | 120 | 23 | 11.23 | 0.00067 |
| GO:0071900 | regulation of protein serine/threonine kinase activity | 278 | 43 | 26.02 | 0.00067 |
| GO:0032990 | cell part morphogenesis | 441 | 62 | 41.28 | 0.0007 |
| GO:0051347 | positive regulation of transferase activity | 389 | 56 | 36.41 | 0.0007 |
| GO:0048562 | embryonic organ morphogenesis | 166 | 29 | 15.54 | 0.00071 |
| GO:0089718 | amino acid import across plasma membrane | 28 | 9 | 2.62 | 0.00071 |
| GO:0007009 | plasma membrane organization | 106 | 21 | 9.92 | 0.00072 |
| GO:0070374 | positive regulation of ERK1 and ERK2 cascade | 106 | 21 | 9.92 | 0.00072 |
| GO:0048534 | hematopoietic or lymphoid organ development | 629 | 83 | 58.88 | 0.00073 |
| GO:0035282 | segmentation | 58 | 14 | 5.43 | 0.00073 |
| GO:0042592 | homeostatic process | 1038 | 127 | 97.16 | 0.00074 |
| GO:0006955 | immune response | 869 | 109 | 81.34 | 0.00075 |
| GO:0010720 | positive regulation of cell development | 214 | 35 | 20.03 | 0.00075 |
| GO:0010863 | positive regulation of phospholipase C activity | 18 | 7 | 1.68 | 0.00078 |
| GO:0110110 | positive regulation of animal organ morphogenesis | 18 | 7 | 1.68 | 0.00078 |
| GO:0010517 | regulation of phospholipase activity | 34 | 10 | 3.18 | 0.0008 |
| GO:0034113 | heterotypic cell-cell adhesion | 34 | 10 | 3.18 | 0.0008 |
| GO:1901654 | response to ketone | 129 | 24 | 12.08 | 0.0008 |
| GO:0007159 | leukocyte cell-cell adhesion | 223 | 36 | 20.87 | 0.00081 |
| GO:0048659 | smooth muscle cell proliferation | 107 | 21 | 10.02 | 0.00081 |
| GO:0060420 | regulation of heart growth | 46 | 12 | 4.31 | 0.00082 |
| GO:0071902 | positive regulation of protein serine/threonine kinase activity | 152 | 27 | 14.23 | 0.00082 |
| GO:0032873 | negative regulation of stress-activated MAPK cascade | 40 | 11 | 3.74 | 0.00083 |
| GO:0045123 | cellular extravasation | 40 | 11 | 3.74 | 0.00083 |
| GO:0048644 | muscle organ morphogenesis | 40 | 11 | 3.74 | 0.00083 |
| GO:0070303 | negative regulation of stress-activated protein kinase signaling cascade | 40 | 11 | 3.74 | 0.00083 |
| GO:0072171 | mesonephric tubule morphogenesis | 40 | 11 | 3.74 | 0.00083 |
| GO:0010634 | positive regulation of epithelial cell migration | 122 | 23 | 11.42 | 0.00084 |
| GO:0061138 | morphogenesis of a branching epithelium | 122 | 23 | 11.42 | 0.00084 |
| GO:0009607 | response to biotic stimulus | 807 | 102 | 75.54 | 0.00085 |
| GO:0048839 | inner ear development | 93 | 19 | 8.71 | 0.00085 |
| GO:0002252 | immune effector process | 315 | 47 | 29.49 | 0.00086 |
| GO:0034620 | cellular response to unfolded protein | 86 | 18 | 8.05 | 0.00086 |
| GO:2000106 | regulation of leukocyte apoptotic process | 59 | 14 | 5.52 | 0.00087 |
| GO:0035265 | organ growth | 115 | 22 | 10.76 | 0.00089 |
| GO:0050877 | nervous system process | 481 | 66 | 45.02 | 0.00092 |
| GO:0007610 | behavior | 342 | 50 | 32.01 | 0.00095 |
| GO:0060043 | regulation of cardiac muscle cell proliferation | 29 | 9 | 2.71 | 0.00095 |
| GO:0006952 | defense response | 894 | 111 | 83.68 | 0.00097 |
| GO:0048762 | mesenchymal cell differentiation | 177 | 30 | 16.57 | 0.00097 |
| GO:0043588 | skin development | 146 | 26 | 13.67 | 0.00098 |
| GO:0003307 | regulation of Wnt signaling pathway involved in heart development | 6 | 4 | 0.56 | 0.00098 |
| GO:0006837 | serotonin transport | 6 | 4 | 0.56 | 0.00098 |
| GO:0010757 | negative regulation of plasminogen activation | 6 | 4 | 0.56 | 0.00098 |

| | | | | | |
|---|---|---|---|---|---|
| GO:0035696 | monocyte extravasation | 6 | 4 | 0.56 | 0.00098 |
| GO:0061043 | regulation of vascular wound healing | 6 | 4 | 0.56 | 0.00098 |
| GO:0061304 | retinal blood vessel morphogenesis | 6 | 4 | 0.56 | 0.00098 |
| GO:0072378 | blood coagulation, fibrin clot formation | 6 | 4 | 0.56 | 0.00098 |
| GO:2000563 | positive regulation of CD4-positive, alpha-beta T cell proliferation | 6 | 4 | 0.56 | 0.00098 |
| GO:0001704 | formation of primary germ layer | 80 | 17 | 7.49 | 0.00099 |
| GO:0048144 | fibroblast proliferation | 80 | 17 | 7.49 | 0.00099 |
| GO:0030856 | regulation of epithelial cell differentiation | 87 | 18 | 8.14 | 0.00099 |
| GO:0001763 | morphogenesis of a branching structure | 131 | 24 | 12.26 | 0.001 |
| GO:0032835 | glomerulus development | 47 | 12 | 4.4 | 0.00101 |
| GO:0060191 | regulation of lipase activity | 47 | 12 | 4.4 | 0.00101 |
| GO:1903317 | regulation of protein maturation | 47 | 12 | 4.4 | 0.00101 |
| GO:0010544 | negative regulation of platelet activation | 14 | 6 | 1.31 | 0.00103 |
| GO:0042474 | middle ear morphogenesis | 14 | 6 | 1.31 | 0.00103 |
| GO:0003208 | cardiac ventricle morphogenesis | 41 | 11 | 3.84 | 0.00104 |
| GO:0048662 | negative regulation of smooth muscle cell proliferation | 41 | 11 | 3.84 | 0.00104 |
| GO:0060389 | pathway-restricted SMAD protein phosphorylation | 41 | 11 | 3.84 | 0.00104 |
| GO:0006939 | smooth muscle contraction | 60 | 14 | 5.62 | 0.00104 |
| GO:0014897 | striated muscle hypertrophy | 60 | 14 | 5.62 | 0.00104 |
| GO:0007520 | myoblast fusion | 24 | 8 | 2.25 | 0.00108 |
| GO:0007519 | skeletal muscle tissue development | 102 | 20 | 9.55 | 0.00109 |
| GO:0110148 | biomineralization | 102 | 20 | 9.55 | 0.00109 |
| GO:0048145 | regulation of fibroblast proliferation | 67 | 15 | 6.27 | 0.0011 |
| GO:0090288 | negative regulation of cellular response to growth factor stimulus | 67 | 15 | 6.27 | 0.0011 |
| GO:0051493 | regulation of cytoskeleton organization | 414 | 58 | 38.75 | 0.00112 |
| GO:0009226 | nucleotide-sugar biosynthetic process | 19 | 7 | 1.78 | 0.00113 |
| GO:0044331 | cell-cell adhesion mediated by cadherin | 19 | 7 | 1.78 | 0.00113 |
| GO:0050820 | positive regulation of coagulation | 19 | 7 | 1.78 | 0.00113 |
| GO:0070528 | protein kinase C signaling | 19 | 7 | 1.78 | 0.00113 |
| GO:0090183 | regulation of kidney development | 19 | 7 | 1.78 | 0.00113 |
| GO:0030510 | regulation of BMP signaling pathway | 74 | 16 | 6.93 | 0.00114 |
| GO:2000345 | regulation of hepatocyte proliferation | 10 | 5 | 0.94 | 0.0012 |
| GO:0055013 | cardiac muscle cell development | 48 | 12 | 4.49 | 0.00123 |
| GO:0014896 | muscle hypertrophy | 61 | 14 | 5.71 | 0.00124 |
| GO:0060348 | bone development | 164 | 28 | 15.35 | 0.00125 |
| GO:0048589 | developmental growth | 434 | 60 | 40.63 | 0.00128 |
| GO:0045685 | regulation of glial cell differentiation | 42 | 11 | 3.93 | 0.00129 |
| GO:0055021 | regulation of cardiac muscle tissue growth | 42 | 11 | 3.93 | 0.00129 |
| GO:0001658 | branching involved in ureteric bud morphogenesis | 36 | 10 | 3.37 | 0.00131 |
| GO:0045778 | positive regulation of ossification | 36 | 10 | 3.37 | 0.00131 |
| GO:0071300 | cellular response to retinoic acid | 36 | 10 | 3.37 | 0.00131 |
| GO:0085029 | extracellular matrix assembly | 36 | 10 | 3.37 | 0.00131 |
| GO:0071466 | cellular response to xenobiotic stimulus | 89 | 18 | 8.33 | 0.00131 |
| GO:0014855 | striated muscle cell proliferation | 55 | 13 | 5.15 | 0.00137 |
| GO:0060411 | cardiac septum morphogenesis | 55 | 13 | 5.15 | 0.00137 |
| GO:0051960 | regulation of nervous system development | 288 | 43 | 26.96 | 0.00138 |
| GO:0010594 | regulation of endothelial cell migration | 134 | 24 | 12.54 | 0.00139 |
| GO:0048667 | cell morphogenesis involved in neuron differentiation | 374 | 53 | 35.01 | 0.00139 |
| GO:0001649 | osteoblast differentiation | 173 | 29 | 16.19 | 0.00139 |
| GO:0048754 | branching morphogenesis of an epithelial tube | 104 | 20 | 9.74 | 0.00139 |
| GO:0015711 | organic anion transport | 189 | 31 | 17.69 | 0.0014 |
| GO:0070482 | response to oxygen levels | 255 | 39 | 23.87 | 0.00144 |
| GO:0031214 | biomineral tissue development | 97 | 19 | 9.08 | 0.00145 |
| GO:0045667 | regulation of osteoblast differentiation | 97 | 19 | 9.08 | 0.00145 |
| GO:0051147 | regulation of muscle cell differentiation | 97 | 19 | 9.08 | 0.00145 |
| GO:0014888 | striated muscle adaptation | 25 | 8 | 2.34 | 0.00146 |
| GO:0032570 | response to progesterone | 25 | 8 | 2.34 | 0.00146 |
| GO:2000403 | positive regulation of lymphocyte migration | 25 | 8 | 2.34 | 0.00146 |
| GO:0048008 | platelet-derived growth factor receptor signaling pathway | 49 | 12 | 4.59 | 0.00149 |
| GO:0030968 | endoplasmic reticulum unfolded protein response | 69 | 15 | 6.46 | 0.00151 |
| GO:0050921 | positive regulation of chemotaxis | 83 | 17 | 7.77 | 0.00152 |
| GO:0051492 | regulation of stress fiber assembly | 76 | 16 | 7.11 | 0.00153 |
| GO:0002684 | positive regulation of immune system process | 482 | 65 | 45.12 | 0.00156 |

| | | | | | |
|---|---|---|---|---|---|
| GO:0008015 | blood circulation | 273 | 41 | 25.55 | 0.00158 |
| GO:0048732 | gland development | 273 | 41 | 25.55 | 0.00158 |
| GO:0072111 | cell proliferation involved in kidney development | 15 | 6 | 1.4 | 0.00158 |
| GO:0031099 | regeneration | 120 | 22 | 11.23 | 0.00158 |
| GO:0010718 | positive regulation of epithelial to mesenchymal transition | 43 | 11 | 4.03 | 0.00159 |
| GO:0031102 | neuron projection regeneration | 43 | 11 | 4.03 | 0.00159 |
| GO:0030097 | hemopoiesis | 600 | 78 | 56.16 | 0.00159 |
| GO:0031128 | developmental induction | 20 | 7 | 1.87 | 0.0016 |
| GO:0090050 | positive regulation of cell migration involved in sprouting angiogenesis | 20 | 7 | 1.87 | 0.0016 |
| GO:1900274 | regulation of phospholipase C activity | 20 | 7 | 1.87 | 0.0016 |
| GO:0071711 | basement membrane organization | 31 | 9 | 2.9 | 0.00161 |
| GO:0042102 | positive regulation of T cell proliferation | 56 | 13 | 5.24 | 0.00164 |
| GO:0050673 | epithelial cell proliferation | 282 | 42 | 26.4 | 0.00164 |
| GO:0007613 | memory | 63 | 14 | 5.9 | 0.00172 |
| GO:0002687 | positive regulation of leukocyte migration | 84 | 17 | 7.86 | 0.00175 |
| GO:0007268 | chemical synaptic transmission | 343 | 49 | 32.11 | 0.00175 |
| GO:0098916 | anterograde trans-synaptic signaling | 343 | 49 | 32.11 | 0.00175 |
| GO:0002367 | cytokine production involved in immune response | 70 | 15 | 6.55 | 0.00176 |
| GO:0002718 | regulation of cytokine production involved in immune response | 70 | 15 | 6.55 | 0.00176 |
| GO:0031401 | positive regulation of protein modification process | 685 | 87 | 64.12 | 0.00177 |
| GO:0097529 | myeloid leukocyte migration | 121 | 22 | 11.33 | 0.00177 |
| GO:0051056 | regulation of small GTPase mediated signal transduction | 225 | 35 | 21.06 | 0.00184 |
| GO:0022414 | reproductive process | 779 | 97 | 72.92 | 0.00184 |
| GO:0002695 | negative regulation of leukocyte activation | 99 | 19 | 9.27 | 0.00186 |
| GO:0019722 | calcium-mediated signaling | 99 | 19 | 9.27 | 0.00186 |
| GO:0009725 | response to hormone | 567 | 74 | 53.07 | 0.00188 |
| GO:0071345 | cellular response to cytokine stimulus | 513 | 68 | 48.02 | 0.00191 |
| GO:0070613 | regulation of protein processing | 44 | 11 | 4.12 | 0.00195 |
| GO:0051128 | regulation of cellular component organization | 1748 | 197 | 163.62 | 0.00197 |
| GO:0006986 | response to unfolded protein | 122 | 22 | 11.42 | 0.00197 |
| GO:0043409 | negative regulation of MAPK cascade | 122 | 22 | 11.42 | 0.00197 |
| GO:0090100 | positive regulation of transmembrane receptor protein serine/threonine kinase signaling pathway | 85 | 17 | 7.96 | 0.002 |
| GO:0008356 | asymmetric cell division | 11 | 5 | 1.03 | 0.00203 |
| GO:0023019 | signal transduction involved in regulation of gene expression | 11 | 5 | 1.03 | 0.00203 |
| GO:0034112 | positive regulation of homotypic cell-cell adhesion | 11 | 5 | 1.03 | 0.00203 |
| GO:0032729 | positive regulation of interferon-gamma production | 38 | 10 | 3.56 | 0.00205 |
| GO:1905330 | regulation of morphogenesis of an epithelium | 38 | 10 | 3.56 | 0.00205 |
| GO:0000768 | syncytium formation by plasma membrane fusion | 32 | 9 | 3 | 0.00206 |
| GO:0022602 | ovulation cycle process | 32 | 9 | 3 | 0.00206 |
| GO:0031641 | regulation of myelination | 32 | 9 | 3 | 0.00206 |
| GO:0035850 | epithelial cell differentiation involved in kidney development | 32 | 9 | 3 | 0.00206 |
| GO:0140253 | cell-cell fusion | 32 | 9 | 3 | 0.00206 |
| GO:0043086 | negative regulation of catalytic activity | 515 | 68 | 48.21 | 0.00211 |
| GO:0003306 | Wnt signaling pathway involved in heart development | 7 | 4 | 0.66 | 0.00212 |
| GO:0032693 | negative regulation of interleukin-10 production | 7 | 4 | 0.66 | 0.00212 |
| GO:0033605 | positive regulation of catecholamine secretion | 7 | 4 | 0.66 | 0.00212 |
| GO:0072203 | cell proliferation involved in metanephros development | 7 | 4 | 0.66 | 0.00212 |
| GO:0120180 | cell-substrate junction disassembly | 7 | 4 | 0.66 | 0.00212 |
| GO:0120181 | focal adhesion disassembly | 7 | 4 | 0.66 | 0.00212 |
| GO:0006820 | anion transport | 252 | 38 | 23.59 | 0.00213 |
| GO:0050670 | regulation of lymphocyte proliferation | 123 | 22 | 11.51 | 0.0022 |
| GO:0044087 | regulation of cellular component biogenesis | 718 | 90 | 67.21 | 0.0022 |
| GO:0050790 | regulation of catalytic activity | 1604 | 182 | 150.14 | 0.00221 |
| GO:0099173 | postsynapse organization | 108 | 20 | 10.11 | 0.00224 |
| GO:0050789 | regulation of biological process | 7214 | 717 | 675.28 | 0.00226 |
| GO:0051216 | cartilage development | 131 | 23 | 12.26 | 0.00227 |
| GO:0003299 | muscle hypertrophy in response to stress | 16 | 6 | 1.5 | 0.00233 |
| GO:0014887 | cardiac muscle adaptation | 16 | 6 | 1.5 | 0.00233 |
| GO:0014898 | cardiac muscle hypertrophy in response to stress | 16 | 6 | 1.5 | 0.00233 |
| GO:0060795 | cell fate commitment involved in formation of primary germ layer | 16 | 6 | 1.5 | 0.00233 |
| GO:0060977 | coronary vasculature morphogenesis | 16 | 6 | 1.5 | 0.00233 |
| GO:1903319 | positive regulation of protein maturation | 16 | 6 | 1.5 | 0.00233 |
| GO:0000003 | reproduction | 785 | 97 | 73.48 | 0.00234 |

| | | | | | |
|---|---|---|---|---|---|
| GO:0043954 | cellular component maintenance | 45 | 11 | 4.21 | 0.00236 |
| GO:0071346 | cellular response to interferon-gamma | 72 | 15 | 6.74 | 0.00237 |
| GO:0099537 | trans-synaptic signaling | 348 | 49 | 32.57 | 0.00237 |
| GO:0071396 | cellular response to lipid | 384 | 53 | 35.94 | 0.00249 |
| GO:0071248 | cellular response to metal ion | 109 | 20 | 10.2 | 0.00251 |
| GO:0065007 | biological regulation | 7650 | 756 | 716.09 | 0.00252 |
| GO:0060393 | regulation of pathway-restricted SMAD protein phosphorylation | 39 | 10 | 3.65 | 0.00252 |
| GO:0060675 | ureteric bud morphogenesis | 39 | 10 | 3.65 | 0.00252 |
| GO:0061005 | cell differentiation involved in kidney development | 39 | 10 | 3.65 | 0.00252 |
| GO:0001953 | negative regulation of cell-matrix adhesion | 27 | 8 | 2.53 | 0.00252 |
| GO:0014911 | positive regulation of smooth muscle cell migration | 27 | 8 | 2.53 | 0.00252 |
| GO:0045687 | positive regulation of glial cell differentiation | 27 | 8 | 2.53 | 0.00252 |
| GO:0051937 | catecholamine transport | 27 | 8 | 2.53 | 0.00252 |
| GO:0099054 | presynapse assembly | 27 | 8 | 2.53 | 0.00252 |
| GO:0010001 | glial cell differentiation | 140 | 24 | 13.1 | 0.00256 |
| GO:0040008 | regulation of growth | 420 | 57 | 39.31 | 0.00256 |
| GO:0055123 | digestive system development | 87 | 17 | 8.14 | 0.00259 |
| GO:0070665 | positive regulation of leukocyte proliferation | 87 | 17 | 8.14 | 0.00259 |
| GO:0009952 | anterior/posterior pattern specification | 117 | 21 | 10.95 | 0.00262 |
| GO:0043085 | positive regulation of catalytic activity | 779 | 96 | 72.92 | 0.00267 |
| GO:0071887 | leukocyte apoptotic process | 80 | 16 | 7.49 | 0.00267 |
| GO:0110020 | regulation of actomyosin structure organization | 80 | 16 | 7.49 | 0.00267 |
| GO:0003300 | cardiac muscle hypertrophy | 59 | 13 | 5.52 | 0.00269 |
| GO:0002520 | immune system development | 677 | 85 | 63.37 | 0.00276 |
| GO:0030098 | lymphocyte differentiation | 239 | 36 | 22.37 | 0.00281 |
| GO:0046942 | carboxylic acid transport | 141 | 24 | 13.2 | 0.00282 |
| GO:0006984 | ER-nucleus signaling pathway | 46 | 11 | 4.31 | 0.00285 |
| GO:0014068 | positive regulation of phosphatidylinositol 3-kinase signaling | 46 | 11 | 4.31 | 0.00285 |
| GO:0002694 | regulation of leukocyte activation | 334 | 47 | 31.26 | 0.00291 |
| GO:0010469 | regulation of signaling receptor activity | 88 | 17 | 8.24 | 0.00294 |
| GO:0048639 | positive regulation of developmental growth | 103 | 19 | 9.64 | 0.00298 |
| GO:0051894 | positive regulation of focal adhesion assembly | 22 | 7 | 2.06 | 0.00299 |
| GO:0072210 | metanephric nephron development | 22 | 7 | 2.06 | 0.00299 |
| GO:1901889 | negative regulation of cell junction assembly | 22 | 7 | 2.06 | 0.00299 |
| GO:1905207 | regulation of cardiocyte differentiation | 22 | 7 | 2.06 | 0.00299 |
| GO:0032944 | regulation of mononuclear cell proliferation | 126 | 22 | 11.79 | 0.003 |
| GO:0051051 | negative regulation of transport | 274 | 40 | 25.65 | 0.00301 |
| GO:0006865 | amino acid transport | 81 | 16 | 7.58 | 0.00305 |
| GO:0048565 | digestive tract development | 81 | 16 | 7.58 | 0.00305 |
| GO:0010596 | negative regulation of endothelial cell migration | 40 | 10 | 3.74 | 0.00309 |
| GO:2000243 | positive regulation of reproductive process | 40 | 10 | 3.74 | 0.00309 |
| GO:0060538 | skeletal muscle organ development | 111 | 20 | 10.39 | 0.00312 |
| GO:0006959 | humoral immune response | 74 | 15 | 6.93 | 0.00313 |
| GO:1901655 | cellular response to ketone | 74 | 15 | 6.93 | 0.00313 |
| GO:0007263 | nitric oxide mediated signal transduction | 12 | 5 | 1.12 | 0.00322 |
| GO:0010771 | negative regulation of cell morphogenesis involved in differentiation | 12 | 5 | 1.12 | 0.00322 |
| GO:0090136 | epithelial cell-cell adhesion | 12 | 5 | 1.12 | 0.00322 |
| GO:1900025 | negative regulation of substrate adhesion-dependent cell spreading | 12 | 5 | 1.12 | 0.00322 |
| GO:0006949 | syncytium formation | 34 | 9 | 3.18 | 0.00325 |
| GO:0043537 | negative regulation of blood vessel endothelial cell migration | 28 | 8 | 2.62 | 0.00325 |
| GO:1902235 | regulation of endoplasmic reticulum stress-induced intrinsic apoptotic signaling pathway | 28 | 8 | 2.62 | 0.00325 |
| GO:0009792 | embryo development ending in birth or egg hatching | 488 | 64 | 45.68 | 0.00331 |
| GO:0001504 | neurotransmitter uptake | 17 | 6 | 1.59 | 0.00332 |
| GO:0001975 | response to amphetamine | 17 | 6 | 1.59 | 0.00332 |
| GO:0002053 | positive regulation of mesenchymal cell proliferation | 17 | 6 | 1.59 | 0.00332 |
| GO:0050927 | positive regulation of positive chemotaxis | 17 | 6 | 1.59 | 0.00332 |
| GO:2000738 | positive regulation of stem cell differentiation | 17 | 6 | 1.59 | 0.00332 |
| GO:0003018 | vascular process in circulatory system | 159 | 26 | 14.88 | 0.0034 |
| GO:0048878 | chemical homeostasis | 608 | 77 | 56.91 | 0.00344 |
| GO:0042063 | gliogenesis | 192 | 30 | 17.97 | 0.00352 |
| GO:0043207 | response to external biotic stimulus | 777 | 95 | 72.73 | 0.00355 |
| GO:0051707 | response to other organism | 777 | 95 | 72.73 | 0.00355 |
| GO:0001738 | morphogenesis of a polarized epithelium | 75 | 15 | 7.02 | 0.00358 |

| | | | | | |
|---|---|---|---|---|---|
| GO:0050671 | positive regulation of lymphocyte proliferation | 75 | 15 | 7.02 | 0.00358 |
| GO:0048863 | stem cell differentiation | 176 | 28 | 16.47 | 0.00364 |
| GO:0014015 | positive regulation of gliogenesis | 41 | 10 | 3.84 | 0.00375 |
| GO:0010811 | positive regulation of cell-substrate adhesion | 90 | 17 | 8.42 | 0.00375 |
| GO:0009893 | positive regulation of metabolic process | 2683 | 287 | 251.15 | 0.0039 |
| GO:0002577 | regulation of antigen processing and presentation | 8 | 4 | 0.75 | 0.00393 |
| GO:0007379 | segment specification | 8 | 4 | 0.75 | 0.00393 |
| GO:0015816 | glycine transport | 8 | 4 | 0.75 | 0.00393 |
| GO:0030638 | polyketide metabolic process | 8 | 4 | 0.75 | 0.00393 |
| GO:0030647 | aminoglycoside antibiotic metabolic process | 8 | 4 | 0.75 | 0.00393 |
| GO:0044597 | daunorubicin metabolic process | 8 | 4 | 0.75 | 0.00393 |
| GO:0044598 | doxorubicin metabolic process | 8 | 4 | 0.75 | 0.00393 |
| GO:0072376 | protein activation cascade | 8 | 4 | 0.75 | 0.00393 |
| GO:1903587 | regulation of blood vessel endothelial cell proliferation involved in sprouting angiogenesis | 8 | 4 | 0.75 | 0.00393 |
| GO:0034341 | response to interferon-gamma | 83 | 16 | 7.77 | 0.00393 |
| GO:0010464 | regulation of mesenchymal cell proliferation | 23 | 7 | 2.15 | 0.00396 |
| GO:0010574 | regulation of vascular endothelial growth factor production | 23 | 7 | 2.15 | 0.00396 |
| GO:0017145 | stem cell division | 23 | 7 | 2.15 | 0.00396 |
| GO:0035886 | vascular associated smooth muscle cell differentiation | 23 | 7 | 2.15 | 0.00396 |
| GO:1904706 | negative regulation of vascular associated smooth muscle cell proliferation | 23 | 7 | 2.15 | 0.00396 |
| GO:1905332 | positive regulation of morphogenesis of an epithelium | 23 | 7 | 2.15 | 0.00396 |
| GO:0045668 | negative regulation of osteoblast differentiation | 35 | 9 | 3.28 | 0.00401 |
| GO:2000351 | regulation of endothelial cell apoptotic process | 35 | 9 | 3.28 | 0.00401 |
| GO:0007586 | digestion | 48 | 11 | 4.49 | 0.00406 |
| GO:0032233 | positive regulation of actin filament bundle assembly | 48 | 11 | 4.49 | 0.00406 |
| GO:1904705 | regulation of vascular associated smooth muscle cell proliferation | 48 | 11 | 4.49 | 0.00406 |
| GO:0010604 | positive regulation of macromolecule metabolic process | 2471 | 266 | 231.3 | 0.00408 |
| GO:0034097 | response to cytokine | 575 | 73 | 53.82 | 0.00408 |
| GO:0003229 | ventricular cardiac muscle tissue development | 29 | 8 | 2.71 | 0.00413 |
| GO:0010862 | positive regulation of pathway-restricted SMAD protein phosphorylation | 29 | 8 | 2.71 | 0.00413 |
| GO:0030574 | collagen catabolic process | 29 | 8 | 2.71 | 0.00413 |
| GO:0035272 | exocrine system development | 29 | 8 | 2.71 | 0.00413 |
| GO:0035967 | cellular response to topologically incorrect protein | 106 | 19 | 9.92 | 0.00415 |
| GO:0002761 | regulation of myeloid leukocyte differentiation | 69 | 14 | 6.46 | 0.0042 |
| GO:0030301 | cholesterol transport | 69 | 14 | 6.46 | 0.0042 |
| GO:0072091 | regulation of stem cell proliferation | 55 | 12 | 5.15 | 0.00421 |
| GO:0050794 | regulation of cellular process | 6853 | 681 | 641.48 | 0.00425 |
| GO:0003205 | cardiac chamber development | 114 | 20 | 10.67 | 0.00428 |
| GO:0042129 | regulation of T cell proliferation | 99 | 18 | 9.27 | 0.00445 |
| GO:0043281 | regulation of cysteine-type endopeptidase activity involved in apoptotic process | 146 | 24 | 13.67 | 0.00446 |
| GO:0007417 | central nervous system development | 661 | 82 | 61.87 | 0.0045 |
| GO:0032890 | regulation of organic acid transport | 42 | 10 | 3.93 | 0.00452 |
| GO:0044092 | negative regulation of molecular function | 746 | 91 | 69.83 | 0.00457 |
| GO:0034694 | response to prostaglandin | 18 | 6 | 1.68 | 0.00459 |
| GO:0034698 | response to gonadotropin | 18 | 6 | 1.68 | 0.00459 |
| GO:0050926 | regulation of positive chemotaxis | 18 | 6 | 1.68 | 0.00459 |
| GO:0043009 | chordate embryonic development | 477 | 62 | 44.65 | 0.00466 |
| GO:0051604 | protein maturation | 238 | 35 | 22.28 | 0.00472 |
| GO:0003158 | endothelium development | 92 | 17 | 8.61 | 0.00474 |
| GO:0050680 | negative regulation of epithelial cell proliferation | 92 | 17 | 8.61 | 0.00474 |
| GO:0001837 | epithelial to mesenchymal transition | 123 | 21 | 11.51 | 0.00483 |
| GO:0006596 | polyamine biosynthetic process | 13 | 5 | 1.22 | 0.00483 |
| GO:0032354 | response to follicle-stimulating hormone | 13 | 5 | 1.22 | 0.00483 |
| GO:0035739 | CD4-positive, alpha-beta T cell proliferation | 13 | 5 | 1.22 | 0.00483 |
| GO:0150011 | regulation of neuron projection arborization | 13 | 5 | 1.22 | 0.00483 |
| GO:1903672 | positive regulation of sprouting angiogenesis | 13 | 5 | 1.22 | 0.00483 |
| GO:2000047 | regulation of cell-cell adhesion mediated by cadherin | 13 | 5 | 1.22 | 0.00483 |
| GO:2000561 | regulation of CD4-positive, alpha-beta T cell proliferation | 13 | 5 | 1.22 | 0.00483 |
| GO:0016525 | negative regulation of angiogenesis | 63 | 13 | 5.9 | 0.00491 |
| GO:1901343 | negative regulation of vasculature development | 63 | 13 | 5.9 | 0.00491 |
| GO:2000181 | negative regulation of blood vessel morphogenesis | 63 | 13 | 5.9 | 0.00491 |
| GO:0003333 | amino acid transmembrane transport | 56 | 12 | 5.24 | 0.00492 |
| GO:0033627 | cell adhesion mediated by integrin | 56 | 12 | 5.24 | 0.00492 |

| | | | | | |
|---|---|---|---|---|---|
| GO:0051153 | regulation of striated muscle cell differentiation | 56 | 12 | 5.24 | 0.00492 |
| GO:0098773 | skin epidermis development | 56 | 12 | 5.24 | 0.00492 |
| GO:0040007 | growth | 626 | 78 | 58.6 | 0.00495 |
| GO:0051962 | positive regulation of nervous system development | 180 | 28 | 16.85 | 0.00501 |
| GO:0030183 | B cell differentiation | 85 | 16 | 7.96 | 0.00502 |
| GO:0014741 | negative regulation of muscle hypertrophy | 24 | 7 | 2.25 | 0.00514 |
| GO:0032613 | interleukin-10 production | 24 | 7 | 2.25 | 0.00514 |
| GO:0032653 | regulation of interleukin-10 production | 24 | 7 | 2.25 | 0.00514 |
| GO:0072012 | glomerulus vasculature development | 24 | 7 | 2.25 | 0.00514 |
| GO:1902742 | apoptotic process involved in development | 24 | 7 | 2.25 | 0.00514 |
| GO:0010463 | mesenchymal cell proliferation | 30 | 8 | 2.81 | 0.00517 |
| GO:0030501 | positive regulation of bone mineralization | 30 | 8 | 2.81 | 0.00517 |
| GO:0031670 | cellular response to nutrient | 30 | 8 | 2.81 | 0.00517 |
| GO:0045332 | phospholipid translocation | 30 | 8 | 2.81 | 0.00517 |
| GO:0099172 | presynapse organization | 30 | 8 | 2.81 | 0.00517 |
| GO:0032946 | positive regulation of mononuclear cell proliferation | 78 | 15 | 7.3 | 0.00527 |
| GO:0072089 | stem cell proliferation | 78 | 15 | 7.3 | 0.00527 |
| GO:0002573 | myeloid leukocyte differentiation | 132 | 22 | 12.36 | 0.00535 |
| GO:0001947 | heart looping | 43 | 10 | 4.03 | 0.0054 |
| GO:1902475 | L-alpha-amino acid transmembrane transport | 43 | 10 | 4.03 | 0.0054 |
| GO:0070661 | leukocyte proliferation | 181 | 28 | 16.94 | 0.00541 |
| GO:0008037 | cell recognition | 71 | 14 | 6.65 | 0.00549 |
| GO:0051250 | negative regulation of lymphocyte activation | 86 | 16 | 8.05 | 0.00564 |
| GO:0060041 | retina development in camera-type eye | 86 | 16 | 8.05 | 0.00564 |
| GO:0042303 | molting cycle | 64 | 13 | 5.99 | 0.00565 |
| GO:0042633 | hair cycle | 64 | 13 | 5.99 | 0.00565 |
| GO:0010633 | negative regulation of epithelial cell migration | 50 | 11 | 4.68 | 0.00565 |
| GO:0042149 | cellular response to glucose starvation | 50 | 11 | 4.68 | 0.00565 |
| GO:1990874 | vascular associated smooth muscle cell proliferation | 50 | 11 | 4.68 | 0.00565 |
| GO:0006793 | phosphorus metabolic process | 2000 | 218 | 187.21 | 0.0057 |
| GO:0048678 | response to axon injury | 57 | 12 | 5.34 | 0.00571 |
| GO:1901890 | positive regulation of cell junction assembly | 57 | 12 | 5.34 | 0.00571 |
| GO:0099536 | synaptic signaling | 364 | 49 | 34.07 | 0.0058 |
| GO:0007229 | integrin-mediated signaling pathway | 79 | 15 | 7.39 | 0.00596 |
| GO:0031103 | axon regeneration | 37 | 9 | 3.46 | 0.00597 |
| GO:0060415 | muscle tissue morphogenesis | 37 | 9 | 3.46 | 0.00597 |
| GO:0036293 | response to decreased oxygen levels | 233 | 34 | 21.81 | 0.00598 |
| GO:0051130 | positive regulation of cellular component organization | 782 | 94 | 73.2 | 0.00603 |
| GO:1902532 | negative regulation of intracellular signal transduction | 401 | 53 | 37.54 | 0.00612 |
| GO:0001702 | gastrulation with mouth forming second | 19 | 6 | 1.78 | 0.00619 |
| GO:0099068 | postsynapse assembly | 19 | 6 | 1.78 | 0.00619 |
| GO:2000516 | positive regulation of CD4-positive, alpha-beta T cell activation | 19 | 6 | 1.78 | 0.00619 |
| GO:0051336 | regulation of hydrolase activity | 641 | 79 | 60 | 0.00619 |
| GO:0043500 | muscle adaptation | 72 | 14 | 6.74 | 0.00625 |
| GO:0051249 | regulation of lymphocyte activation | 286 | 40 | 26.77 | 0.00637 |
| GO:1902905 | positive regulation of supramolecular fiber organization | 134 | 22 | 12.54 | 0.00641 |
| GO:0002762 | negative regulation of myeloid leukocyte differentiation | 31 | 8 | 2.9 | 0.00641 |
| GO:0006335 | DNA replication-dependent chromatin assembly | 31 | 8 | 2.9 | 0.00641 |
| GO:0034405 | response to fluid shear stress | 31 | 8 | 2.9 | 0.00641 |
| GO:0044319 | wound healing, spreading of cells | 31 | 8 | 2.9 | 0.00641 |
| GO:0048546 | digestive tract morphogenesis | 31 | 8 | 2.9 | 0.00641 |
| GO:0090505 | epiboly involved in wound healing | 31 | 8 | 2.9 | 0.00641 |
| GO:0140467 | integrated stress response signaling | 31 | 8 | 2.9 | 0.00641 |
| GO:0061371 | determination of heart left/right asymmetry | 44 | 10 | 4.12 | 0.00642 |
| GO:0006048 | UDP-N-acetylglucosamine biosynthetic process | 9 | 4 | 0.84 | 0.00655 |
| GO:0009158 | ribonucleoside monophosphate catabolic process | 9 | 4 | 0.84 | 0.00655 |
| GO:0014745 | negative regulation of muscle adaptation | 9 | 4 | 0.84 | 0.00655 |
| GO:0015801 | aromatic amino acid transport | 9 | 4 | 0.84 | 0.00655 |
| GO:0031642 | negative regulation of myelination | 9 | 4 | 0.84 | 0.00655 |
| GO:0031645 | negative regulation of nervous system process | 9 | 4 | 0.84 | 0.00655 |
| GO:0032026 | response to magnesium ion | 9 | 4 | 0.84 | 0.00655 |
| GO:0048103 | somatic stem cell division | 9 | 4 | 0.84 | 0.00655 |
| GO:0051549 | positive regulation of keratinocyte migration | 9 | 4 | 0.84 | 0.00655 |

| | | | | | |
|---|---|---|---|---|---|
| GO:0051956 | negative regulation of amino acid transport | 9 | 4 | 0.84 | 0.00655 |
| GO:0061299 | retina vasculature morphogenesis in camera-type eye | 9 | 4 | 0.84 | 0.00655 |
| GO:0070234 | positive regulation of T cell apoptotic process | 9 | 4 | 0.84 | 0.00655 |
| GO:2000696 | regulation of epithelial cell differentiation involved in kidney development | 9 | 4 | 0.84 | 0.00655 |
| GO:0007193 | adenylate cyclase-inhibiting G protein-coupled receptor signaling pathway | 25 | 7 | 2.34 | 0.00657 |
| GO:0034110 | regulation of homotypic cell-cell adhesion | 25 | 7 | 2.34 | 0.00657 |
| GO:0060317 | cardiac epithelial to mesenchymal transition | 25 | 7 | 2.34 | 0.00657 |
| GO:0035050 | embryonic heart tube development | 58 | 12 | 5.43 | 0.00661 |
| GO:0002011 | morphogenesis of an epithelial sheet | 51 | 11 | 4.77 | 0.00661 |
| GO:0098739 | import across plasma membrane | 95 | 17 | 8.89 | 0.00661 |
| GO:0048871 | multicellular organismal homeostasis | 305 | 42 | 28.55 | 0.00691 |
| GO:0035148 | tube formation | 111 | 19 | 10.39 | 0.00693 |
| GO:0035966 | response to topologically incorrect protein | 143 | 23 | 13.39 | 0.00694 |
| GO:0071379 | cellular response to prostaglandin stimulus | 14 | 5 | 1.31 | 0.00694 |
| GO:0071636 | positive regulation of transforming growth factor beta production | 14 | 5 | 1.31 | 0.00694 |
| GO:0072202 | cell differentiation involved in metanephros development | 14 | 5 | 1.31 | 0.00694 |
| GO:0072574 | hepatocyte proliferation | 14 | 5 | 1.31 | 0.00694 |
| GO:0072575 | epithelial cell proliferation involved in liver morphogenesis | 14 | 5 | 1.31 | 0.00694 |
| GO:0042098 | T cell proliferation | 119 | 20 | 11.14 | 0.007 |
| GO:0002578 | negative regulation of antigen processing and presentation | 5 | 3 | 0.47 | 0.00708 |
| GO:0002692 | negative regulation of cellular extravasation | 5 | 3 | 0.47 | 0.00708 |
| GO:0003278 | apoptotic process involved in heart morphogenesis | 5 | 3 | 0.47 | 0.00708 |
| GO:0003284 | septum primum development | 5 | 3 | 0.47 | 0.00708 |
| GO:0003289 | atrial septum primum morphogenesis | 5 | 3 | 0.47 | 0.00708 |
| GO:0003334 | keratinocyte development | 5 | 3 | 0.47 | 0.00708 |
| GO:0008354 | germ cell migration | 5 | 3 | 0.47 | 0.00708 |
| GO:0021936 | regulation of cerebellar granule cell precursor proliferation | 5 | 3 | 0.47 | 0.00708 |
| GO:0035701 | hematopoietic stem cell migration | 5 | 3 | 0.47 | 0.00708 |
| GO:0035905 | ascending aorta development | 5 | 3 | 0.47 | 0.00708 |
| GO:0045741 | positive regulation of epidermal growth factor-activated receptor activity | 5 | 3 | 0.47 | 0.00708 |
| GO:0046035 | CMP metabolic process | 5 | 3 | 0.47 | 0.00708 |
| GO:0051610 | serotonin uptake | 5 | 3 | 0.47 | 0.00708 |
| GO:0051891 | positive regulation of cardioblast differentiation | 5 | 3 | 0.47 | 0.00708 |
| GO:0060978 | angiogenesis involved in coronary vascular morphogenesis | 5 | 3 | 0.47 | 0.00708 |
| GO:0061343 | cell adhesion involved in heart morphogenesis | 5 | 3 | 0.47 | 0.00708 |
| GO:0071374 | cellular response to parathyroid hormone stimulus | 5 | 3 | 0.47 | 0.00708 |
| GO:0098713 | leucine import across plasma membrane | 5 | 3 | 0.47 | 0.00708 |
| GO:1900019 | regulation of protein kinase C activity | 5 | 3 | 0.47 | 0.00708 |
| GO:1900020 | positive regulation of protein kinase C activity | 5 | 3 | 0.47 | 0.00708 |
| GO:1903801 | L-leucine import across plasma membrane | 5 | 3 | 0.47 | 0.00708 |
| GO:1905005 | regulation of epithelial to mesenchymal transition involved in endocardial cushion formation | 5 | 3 | 0.47 | 0.00708 |
| GO:1905007 | positive regulation of epithelial to mesenchymal transition involved in endocardial cushion formation | 5 | 3 | 0.47 | 0.00708 |
| GO:2000822 | regulation of behavioral fear response | 5 | 3 | 0.47 | 0.00708 |
| GO:0035270 | endocrine system development | 73 | 14 | 6.83 | 0.00708 |
| GO:0072507 | divalent inorganic cation homeostasis | 193 | 29 | 18.07 | 0.0071 |
| GO:0030514 | negative regulation of BMP signaling pathway | 38 | 9 | 3.56 | 0.00719 |
| GO:0043551 | regulation of phosphatidylinositol 3-kinase activity | 38 | 9 | 3.56 | 0.00719 |
| GO:0045047 | protein targeting to ER | 38 | 9 | 3.56 | 0.00719 |
| GO:0061008 | hepaticobiliary system development | 96 | 17 | 8.99 | 0.00735 |
| GO:1903037 | regulation of leukocyte cell-cell adhesion | 202 | 30 | 18.91 | 0.00738 |
| GO:0007265 | Ras protein signal transduction | 271 | 38 | 25.37 | 0.00739 |
| GO:0051145 | smooth muscle cell differentiation | 45 | 10 | 4.21 | 0.00758 |
| GO:0002449 | lymphocyte mediated immunity | 136 | 22 | 12.73 | 0.00763 |
| GO:0022408 | negative regulation of cell-cell adhesion | 112 | 19 | 10.48 | 0.00764 |
| GO:0035567 | non-canonical Wnt signaling pathway | 52 | 11 | 4.87 | 0.00769 |
| GO:0055074 | calcium ion homeostasis | 169 | 26 | 15.82 | 0.00772 |
| GO:0090504 | epiboly | 32 | 8 | 3 | 0.00786 |
| GO:0051048 | negative regulation of secretion | 89 | 16 | 8.33 | 0.00791 |
| GO:1905039 | carboxylic acid transmembrane transport | 89 | 16 | 8.33 | 0.00791 |
| GO:0010717 | regulation of epithelial to mesenchymal transition | 74 | 14 | 6.93 | 0.008 |
| GO:0045944 | positive regulation of transcription by RNA polymerase II | 867 | 102 | 81.16 | 0.00804 |
| GO:0046651 | lymphocyte proliferation | 153 | 24 | 14.32 | 0.00806 |
| GO:0001662 | behavioral fear response | 20 | 6 | 1.87 | 0.00815 |

| | | | | | |
|---|---|---|---|---|---|
| GO:0002369 | T cell cytokine production | 20 | 6 | 1.87 | 0.00815 |
| GO:0002724 | regulation of T cell cytokine production | 20 | 6 | 1.87 | 0.00815 |
| GO:0010875 | positive regulation of cholesterol efflux | 20 | 6 | 1.87 | 0.00815 |
| GO:0015872 | dopamine transport | 20 | 6 | 1.87 | 0.00815 |
| GO:0034656 | nucleobase-containing small molecule catabolic process | 20 | 6 | 1.87 | 0.00815 |
| GO:0035767 | endothelial cell chemotaxis | 20 | 6 | 1.87 | 0.00815 |
| GO:0050433 | regulation of catecholamine secretion | 20 | 6 | 1.87 | 0.00815 |
| GO:0051954 | positive regulation of amine transport | 20 | 6 | 1.87 | 0.00815 |
| GO:0099174 | regulation of presynapse organization | 20 | 6 | 1.87 | 0.00815 |
| GO:1905606 | regulation of presynapse assembly | 20 | 6 | 1.87 | 0.00815 |
| GO:0032872 | regulation of stress-activated MAPK cascade | 145 | 23 | 13.57 | 0.0082 |
| GO:0010573 | vascular endothelial growth factor production | 26 | 7 | 2.43 | 0.00828 |
| GO:0033687 | osteoblast proliferation | 26 | 7 | 2.43 | 0.00828 |
| GO:0035459 | vesicle cargo loading | 26 | 7 | 2.43 | 0.00828 |
| GO:0061437 | renal system vasculature development | 26 | 7 | 2.43 | 0.00828 |
| GO:0061440 | kidney vasculature development | 26 | 7 | 2.43 | 0.00828 |
| GO:0097242 | amyloid-beta clearance | 26 | 7 | 2.43 | 0.00828 |
| GO:0001666 | response to hypoxia | 221 | 32 | 20.69 | 0.0084 |
| GO:0060349 | bone morphogenesis | 67 | 13 | 6.27 | 0.0084 |
| GO:0050769 | positive regulation of neurogenesis | 162 | 25 | 15.16 | 0.00854 |
| GO:0060976 | coronary vasculature development | 39 | 9 | 3.65 | 0.00859 |
| GO:0072577 | endothelial cell apoptotic process | 39 | 9 | 3.65 | 0.00859 |
| GO:0097006 | regulation of plasma lipoprotein particle levels | 39 | 9 | 3.65 | 0.00859 |
| GO:0097035 | regulation of membrane lipid distribution | 39 | 9 | 3.65 | 0.00859 |
| GO:2000401 | regulation of lymphocyte migration | 39 | 9 | 3.65 | 0.00859 |
| GO:0051049 | regulation of transport | 1034 | 119 | 96.79 | 0.00867 |
| GO:1902105 | regulation of leukocyte differentiation | 179 | 27 | 16.76 | 0.00872 |
| GO:0014013 | regulation of gliogenesis | 60 | 12 | 5.62 | 0.00872 |
| GO:0035710 | CD4-positive, alpha-beta T cell activation | 60 | 12 | 5.62 | 0.00872 |
| GO:0048661 | positive regulation of smooth muscle cell proliferation | 60 | 12 | 5.62 | 0.00872 |
| GO:1902106 | negative regulation of leukocyte differentiation | 60 | 12 | 5.62 | 0.00872 |
| GO:1903825 | organic acid transmembrane transport | 90 | 16 | 8.42 | 0.00881 |
| GO:0006821 | chloride transport | 46 | 10 | 4.31 | 0.00889 |
| GO:0070252 | actin-mediated cell contraction | 53 | 11 | 4.96 | 0.0089 |
| GO:0051495 | positive regulation of cytoskeleton organization | 146 | 23 | 13.67 | 0.0089 |
| GO:0008154 | actin polymerization or depolymerization | 155 | 24 | 14.51 | 0.00945 |
| GO:0010769 | regulation of cell morphogenesis involved in differentiation | 83 | 15 | 7.77 | 0.00947 |
| GO:0002688 | regulation of leukocyte chemotaxis | 68 | 13 | 6.37 | 0.00952 |
| GO:0043277 | apoptotic cell clearance | 33 | 8 | 3.09 | 0.00955 |
| GO:2000107 | negative regulation of leukocyte apoptotic process | 33 | 8 | 3.09 | 0.00955 |
| GO:0003215 | cardiac right ventricle morphogenesis | 15 | 5 | 1.4 | 0.00962 |
| GO:0009164 | nucleoside catabolic process | 15 | 5 | 1.4 | 0.00962 |
| GO:0010002 | cardioblast differentiation | 15 | 5 | 1.4 | 0.00962 |
| GO:0010954 | positive regulation of protein processing | 15 | 5 | 1.4 | 0.00962 |
| GO:0014046 | dopamine secretion | 15 | 5 | 1.4 | 0.00962 |
| GO:0014059 | regulation of dopamine secretion | 15 | 5 | 1.4 | 0.00962 |
| GO:0030220 | platelet formation | 15 | 5 | 1.4 | 0.00962 |
| GO:0032891 | negative regulation of organic acid transport | 15 | 5 | 1.4 | 0.00962 |
| GO:0045649 | regulation of macrophage differentiation | 15 | 5 | 1.4 | 0.00962 |
| GO:0051895 | negative regulation of focal adhesion assembly | 15 | 5 | 1.4 | 0.00962 |
| GO:0070293 | renal absorption | 15 | 5 | 1.4 | 0.00962 |
| GO:0072576 | liver morphogenesis | 15 | 5 | 1.4 | 0.00962 |
| GO:0090110 | COPII-coated vesicle cargo loading | 15 | 5 | 1.4 | 0.00962 |
| GO:0150118 | negative regulation of cell-substrate junction organization | 15 | 5 | 1.4 | 0.00962 |
| GO:2001026 | regulation of endothelial cell chemotaxis | 15 | 5 | 1.4 | 0.00962 |
| GO:0001570 | vasculogenesis | 61 | 12 | 5.71 | 0.00995 |
| GO:0009653 | anatomical structure morphogenesis | 1729 | 273 | 161.85 | 6.00E-21 |
| GO:0007155 | cell adhesion | 880 | 168 | 82.37 | 1.00E-20 |
| GO:0030154 | cell differentiation | 2518 | 358 | 235.7 | 5.70E-20 |
| GO:0048856 | anatomical structure development | 3564 | 467 | 333.61 | 6.70E-20 |
| GO:0032501 | multicellular organismal process | 4067 | 517 | 380.7 | 7.80E-20 |
| GO:0040011 | locomotion | 855 | 162 | 80.03 | 1.30E-19 |
| GO:0048869 | cellular developmental process | 2542 | 359 | 237.95 | 1.60E-19 |

| | | | | | |
|---|---|---|---|---|---|
| GO:0048513 | animal organ development | 2154 | 314 | 201.63 | 9.20E-19 |
| GO:0032502 | developmental process | 3904 | 495 | 365.44 | 2.50E-18 |
| GO:0048731 | system development | 2717 | 371 | 254.33 | 1.10E-17 |
| GO:0009888 | tissue development | 1187 | 198 | 111.11 | 1.90E-17 |
| GO:0007275 | multicellular organism development | 3022 | 399 | 282.88 | 1.10E-16 |
| GO:0030334 | regulation of cell migration | 645 | 124 | 60.38 | 1.40E-15 |
| GO:0051239 | regulation of multicellular organismal process | 1676 | 249 | 156.88 | 1.70E-15 |
| GO:0016477 | cell migration | 975 | 166 | 91.27 | 2.00E-15 |
| GO:2000145 | regulation of cell motility | 679 | 128 | 63.56 | 2.30E-15 |
| GO:0048870 | cell motility | 1053 | 174 | 98.57 | 6.80E-15 |
| GO:0009887 | animal organ morphogenesis | 622 | 119 | 58.22 | 8.10E-15 |
| GO:0061061 | muscle structure development | 442 | 94 | 41.37 | 9.60E-15 |
| GO:0009611 | response to wounding | 362 | 82 | 33.89 | 1.40E-14 |
| GO:0040012 | regulation of locomotion | 701 | 128 | 65.62 | 2.80E-14 |
| GO:0042060 | wound healing | 281 | 69 | 26.3 | 2.90E-14 |
| GO:0072359 | circulatory system development | 779 | 138 | 72.92 | 2.90E-14 |
| GO:0098609 | cell-cell adhesion | 531 | 105 | 49.7 | 3.90E-14 |
| GO:0007154 | cell communication | 3662 | 452 | 342.79 | 7.10E-14 |
| GO:0050793 | regulation of developmental process | 1627 | 237 | 152.3 | 1.10E-13 |
| GO:0050896 | response to stimulus | 5137 | 593 | 480.86 | 1.50E-13 |
| GO:0022603 | regulation of anatomical structure morphogenesis | 654 | 119 | 61.22 | 3.30E-13 |
| GO:0023052 | signaling | 3614 | 444 | 338.29 | 3.50E-13 |
| GO:0007165 | signal transduction | 3357 | 418 | 314.24 | 3.90E-13 |
| GO:0070887 | cellular response to chemical stimulus | 1928 | 268 | 180.47 | 4.50E-13 |
| GO:0009605 | response to external stimulus | 1552 | 226 | 145.28 | 5.10E-13 |
| GO:0048468 | cell development | 1305 | 197 | 122.16 | 8.30E-13 |
| GO:0048646 | anatomical structure formation involved in morphogenesis | 751 | 130 | 70.3 | 1.10E-12 |
| GO:0007166 | cell surface receptor signaling pathway | 1664 | 236 | 155.76 | 2.40E-12 |
| GO:0051241 | negative regulation of multicellular organismal process | 631 | 113 | 59.07 | 4.10E-12 |
| GO:0030036 | actin cytoskeleton organization | 520 | 98 | 48.68 | 6.00E-12 |
| GO:0030029 | actin filament-based process | 567 | 104 | 53.07 | 7.20E-12 |
| GO:0003008 | system process | 922 | 148 | 86.3 | 1.00E-11 |
| GO:0042221 | response to chemical | 2479 | 321 | 232.05 | 1.10E-11 |
| GO:0042127 | regulation of cell population proliferation | 1049 | 163 | 98.19 | 1.20E-11 |
| GO:0051716 | cellular response to stimulus | 4428 | 516 | 414.49 | 1.30E-11 |
| GO:0071310 | cellular response to organic substance | 1544 | 219 | 144.53 | 1.90E-11 |
| GO:2000026 | regulation of multicellular organismal development | 893 | 143 | 83.59 | 3.00E-11 |
| GO:0001944 | vasculature development | 519 | 96 | 48.58 | 3.00E-11 |
| GO:0060429 | epithelium development | 716 | 121 | 67.02 | 3.70E-11 |
| GO:0042692 | muscle cell differentiation | 256 | 59 | 23.96 | 3.80E-11 |
| GO:0006935 | chemotaxis | 338 | 71 | 31.64 | 3.80E-11 |
| GO:0042330 | taxis | 338 | 71 | 31.64 | 3.80E-11 |
| GO:0035239 | tube morphogenesis | 600 | 106 | 56.16 | 4.70E-11 |
| GO:0030155 | regulation of cell adhesion | 488 | 91 | 45.68 | 6.60E-11 |
| GO:0030198 | extracellular matrix organization | 208 | 51 | 19.47 | 7.70E-11 |
| GO:0043062 | extracellular structure organization | 208 | 51 | 19.47 | 7.70E-11 |
| GO:0035295 | tube development | 741 | 123 | 69.36 | 8.30E-11 |
| GO:0045229 | external encapsulating structure organization | 209 | 51 | 19.56 | 9.30E-11 |
| GO:0034330 | cell junction organization | 434 | 83 | 40.63 | 1.30E-10 |
| GO:0007167 | enzyme-linked receptor protein signaling pathway | 650 | 111 | 60.84 | 1.40E-10 |
| GO:0060537 | muscle tissue development | 258 | 58 | 24.15 | 1.60E-10 |
| GO:0008283 | cell population proliferation | 1249 | 181 | 116.91 | 3.00E-10 |
| GO:0010646 | regulation of cell communication | 2255 | 290 | 211.08 | 4.20E-10 |
| GO:0030335 | positive regulation of cell migration | 378 | 74 | 35.38 | 4.60E-10 |
| GO:0022008 | neurogenesis | 1051 | 157 | 98.38 | 6.40E-10 |
| GO:2000147 | positive regulation of cell motility | 388 | 75 | 36.32 | 6.40E-10 |
| GO:0048729 | tissue morphogenesis | 418 | 79 | 39.13 | 6.70E-10 |
| GO:0001568 | blood vessel development | 494 | 89 | 46.24 | 6.80E-10 |
| GO:0031589 | cell-substrate adhesion | 254 | 56 | 23.78 | 7.20E-10 |
| GO:0048699 | generation of neurons | 903 | 139 | 84.53 | 9.60E-10 |
| GO:0002009 | morphogenesis of an epithelium | 357 | 70 | 33.42 | 1.30E-09 |
| GO:0040017 | positive regulation of locomotion | 396 | 75 | 37.07 | 1.70E-09 |
| GO:0023051 | regulation of signaling | 2266 | 288 | 212.11 | 1.90E-09 |

| | | | | | |
|---|---|---|---|---|---|
| GO:0051146 | striated muscle cell differentiation | 175 | 43 | 16.38 | 2.20E-09 |
| GO:0051094 | positive regulation of developmental process | 866 | 133 | 81.06 | 2.70E-09 |
| GO:0010033 | response to organic substance | 1952 | 254 | 182.72 | 2.70E-09 |
| GO:0065008 | regulation of biological quality | 2257 | 286 | 211.27 | 3.00E-09 |
| GO:0071363 | cellular response to growth factor stimulus | 478 | 85 | 44.74 | 3.30E-09 |
| GO:0007596 | blood coagulation | 134 | 36 | 12.54 | 3.50E-09 |
| GO:0030182 | neuron differentiation | 855 | 131 | 80.03 | 4.20E-09 |
| GO:0050817 | coagulation | 135 | 36 | 12.64 | 4.40E-09 |
| GO:0072001 | renal system development | 220 | 49 | 20.59 | 5.90E-09 |
| GO:0001775 | cell activation | 619 | 102 | 57.94 | 6.00E-09 |
| GO:0051240 | positive regulation of multicellular organismal process | 912 | 137 | 85.37 | 6.60E-09 |
| GO:0001822 | kidney development | 214 | 48 | 20.03 | 6.70E-09 |
| GO:0007599 | hemostasis | 137 | 36 | 12.82 | 6.80E-09 |
| GO:0070848 | response to growth factor | 496 | 86 | 46.43 | 9.10E-09 |
| GO:0007399 | nervous system development | 1559 | 209 | 145.93 | 1.00E-08 |
| GO:1901888 | regulation of cell junction assembly | 128 | 34 | 11.98 | 1.30E-08 |
| GO:0050878 | regulation of body fluid levels | 213 | 47 | 19.94 | 1.70E-08 |
| GO:0097435 | supramolecular fiber organization | 566 | 94 | 52.98 | 1.70E-08 |
| GO:0048514 | blood vessel morphogenesis | 432 | 77 | 40.44 | 1.70E-08 |
| GO:0000902 | cell morphogenesis | 691 | 109 | 64.68 | 2.00E-08 |
| GO:0001655 | urogenital system development | 244 | 51 | 22.84 | 2.80E-08 |
| GO:0048583 | regulation of response to stimulus | 2617 | 318 | 244.97 | 2.90E-08 |
| GO:0010941 | regulation of cell death | 1142 | 160 | 106.9 | 4.40E-08 |
| GO:0034329 | cell junction assembly | 255 | 52 | 23.87 | 4.80E-08 |
| GO:0010810 | regulation of cell-substrate adhesion | 160 | 38 | 14.98 | 5.10E-08 |
| GO:0007507 | heart development | 420 | 74 | 39.31 | 5.30E-08 |
| GO:0009966 | regulation of signal transduction | 2046 | 257 | 191.52 | 6.30E-08 |
| GO:0008285 | negative regulation of cell population proliferation | 469 | 80 | 43.9 | 6.40E-08 |
| GO:0000165 | MAPK cascade | 493 | 83 | 46.15 | 6.50E-08 |
| GO:0007423 | sensory organ development | 316 | 60 | 29.58 | 6.80E-08 |
| GO:1903034 | regulation of response to wounding | 100 | 28 | 9.36 | 7.40E-08 |
| GO:0030855 | epithelial cell differentiation | 373 | 67 | 34.92 | 1.10E-07 |
| GO:0035051 | cardiocyte differentiation | 102 | 28 | 9.55 | 1.20E-07 |
| GO:0008284 | positive regulation of cell population proliferation | 573 | 92 | 53.64 | 1.20E-07 |
| GO:0043067 | regulation of programmed cell death | 1035 | 146 | 96.88 | 1.30E-07 |
| GO:0001525 | angiogenesis | 367 | 66 | 34.35 | 1.30E-07 |
| GO:0061045 | negative regulation of wound healing | 39 | 16 | 3.65 | 1.50E-07 |
| GO:0043534 | blood vessel endothelial cell migration | 98 | 27 | 9.17 | 1.80E-07 |
| GO:0042981 | regulation of apoptotic process | 1015 | 143 | 95.01 | 1.90E-07 |
| GO:0001932 | regulation of protein phosphorylation | 730 | 110 | 68.33 | 2.10E-07 |
| GO:0014706 | striated muscle tissue development | 155 | 36 | 14.51 | 2.10E-07 |
| GO:0045595 | regulation of cell differentiation | 1018 | 143 | 95.29 | 2.20E-07 |
| GO:0043408 | regulation of MAPK cascade | 419 | 72 | 39.22 | 2.20E-07 |
| GO:0048598 | embryonic morphogenesis | 396 | 69 | 37.07 | 2.30E-07 |
| GO:0031032 | actomyosin structure organization | 143 | 34 | 13.39 | 2.50E-07 |
| GO:0001934 | positive regulation of protein phosphorylation | 460 | 77 | 43.06 | 2.50E-07 |
| GO:0030168 | platelet activation | 82 | 24 | 7.68 | 2.50E-07 |
| GO:0061041 | regulation of wound healing | 77 | 23 | 7.21 | 3.00E-07 |
| GO:0090130 | tissue migration | 241 | 48 | 22.56 | 3.30E-07 |
| GO:0007015 | actin filament organization | 330 | 60 | 30.89 | 3.30E-07 |
| GO:0055001 | muscle cell development | 113 | 29 | 10.58 | 3.40E-07 |
| GO:0008219 | cell death | 1485 | 193 | 139.01 | 4.40E-07 |
| GO:0007186 | G protein-coupled receptor signaling pathway | 295 | 55 | 27.61 | 4.50E-07 |
| GO:0045785 | positive regulation of cell adhesion | 295 | 55 | 27.61 | 4.50E-07 |
| GO:0034109 | homotypic cell-cell adhesion | 62 | 20 | 5.8 | 4.50E-07 |
| GO:0042476 | odontogenesis | 73 | 22 | 6.83 | 4.60E-07 |
| GO:0007160 | cell-matrix adhesion | 160 | 36 | 14.98 | 4.80E-07 |
| GO:0048666 | neuron development | 709 | 106 | 66.37 | 5.20E-07 |
| GO:0001503 | ossification | 289 | 54 | 27.05 | 5.30E-07 |
| GO:0007517 | muscle organ development | 209 | 43 | 19.56 | 5.40E-07 |
| GO:0010631 | epithelial cell migration | 238 | 47 | 22.28 | 5.60E-07 |
| GO:0090132 | epithelium migration | 238 | 47 | 22.28 | 5.60E-07 |
| GO:0048584 | positive regulation of response to stimulus | 1368 | 179 | 128.05 | 8.10E-07 |

| | | | | | |
|---|---|---|---|---|---|
| GO:0060021 | roof of mouth development | 70 | 21 | 6.55 | 9.00E-07 |
| GO:0098742 | cell-cell adhesion via plasma-membrane adhesion molecules | 131 | 31 | 12.26 | 9.30E-07 |
| GO:1901342 | regulation of vasculature development | 199 | 41 | 18.63 | 9.40E-07 |
| GO:0071495 | cellular response to endogenous stimulus | 892 | 126 | 83.5 | 9.60E-07 |
| GO:1903035 | negative regulation of response to wounding | 49 | 17 | 4.59 | 1.00E-06 |
| GO:0072073 | kidney epithelium development | 88 | 24 | 8.24 | 1.10E-06 |
| GO:0032970 | regulation of actin filament-based process | 296 | 54 | 27.71 | 1.20E-06 |
| GO:0051093 | negative regulation of developmental process | 585 | 90 | 54.76 | 1.20E-06 |
| GO:0042327 | positive regulation of phosphorylation | 511 | 81 | 47.83 | 1.20E-06 |
| GO:0045766 | positive regulation of angiogenesis | 120 | 29 | 11.23 | 1.30E-06 |
| GO:1904018 | positive regulation of vasculature development | 120 | 29 | 11.23 | 1.30E-06 |
| GO:0072006 | nephron development | 95 | 25 | 8.89 | 1.30E-06 |
| GO:0045165 | cell fate commitment | 133 | 31 | 12.45 | 1.30E-06 |
| GO:0048585 | negative regulation of response to stimulus | 1131 | 152 | 105.87 | 1.40E-06 |
| GO:0009719 | response to endogenous stimulus | 1041 | 142 | 97.44 | 1.40E-06 |
| GO:0010647 | positive regulation of cell communication | 1105 | 149 | 103.43 | 1.50E-06 |
| GO:0000904 | cell morphogenesis involved in differentiation | 489 | 78 | 45.77 | 1.50E-06 |
| GO:0002376 | immune system process | 1522 | 194 | 142.47 | 1.60E-06 |
| GO:0010562 | positive regulation of phosphorus metabolic process | 557 | 86 | 52.14 | 1.80E-06 |
| GO:0045937 | positive regulation of phosphate metabolic process | 557 | 86 | 52.14 | 1.80E-06 |
| GO:0045765 | regulation of angiogenesis | 197 | 40 | 18.44 | 1.90E-06 |
| GO:0001656 | metanephros development | 51 | 17 | 4.77 | 1.90E-06 |
| GO:0003012 | muscle system process | 241 | 46 | 22.56 | 2.00E-06 |
| GO:0009967 | positive regulation of signal transduction | 1013 | 138 | 94.82 | 2.20E-06 |
| GO:0012501 | programmed cell death | 1371 | 177 | 128.33 | 2.30E-06 |
| GO:0003007 | heart morphogenesis | 171 | 36 | 16.01 | 2.50E-06 |
| GO:0006915 | apoptotic process | 1337 | 173 | 125.15 | 2.70E-06 |
| GO:0043010 | camera-type eye development | 200 | 40 | 18.72 | 2.80E-06 |
| GO:0061333 | renal tubule morphogenesis | 47 | 16 | 4.4 | 2.80E-06 |
| GO:0044703 | multi-organism reproductive process | 118 | 28 | 11.05 | 2.90E-06 |
| GO:0072009 | nephron epithelium development | 69 | 20 | 6.46 | 3.00E-06 |
| GO:0045860 | positive regulation of protein kinase activity | 267 | 49 | 24.99 | 3.00E-06 |
| GO:0030195 | negative regulation of blood coagulation | 28 | 12 | 2.62 | 3.10E-06 |
| GO:1900047 | negative regulation of hemostasis | 28 | 12 | 2.62 | 3.10E-06 |
| GO:0061326 | renal tubule development | 58 | 18 | 5.43 | 3.20E-06 |
| GO:0048738 | cardiac muscle tissue development | 145 | 32 | 13.57 | 3.20E-06 |
| GO:0023056 | positive regulation of signaling | 1111 | 148 | 104 | 3.20E-06 |
| GO:0062009 | secondary palate development | 16 | 9 | 1.5 | 3.30E-06 |
| GO:0010648 | negative regulation of cell communication | 967 | 132 | 90.52 | 3.30E-06 |
| GO:0007157 | heterophilic cell-cell adhesion via plasma membrane cell adhesion molecules | 24 | 11 | 2.25 | 3.60E-06 |
| GO:0023057 | negative regulation of signaling | 970 | 132 | 90.8 | 3.90E-06 |
| GO:1901700 | response to oxygen-containing compound | 1043 | 140 | 97.63 | 4.10E-06 |
| GO:0042325 | regulation of phosphorylation | 837 | 117 | 78.35 | 4.10E-06 |
| GO:0022604 | regulation of cell morphogenesis | 241 | 45 | 22.56 | 4.80E-06 |
| GO:0007169 | transmembrane receptor protein tyrosine kinase signaling pathway | 405 | 66 | 37.91 | 4.80E-06 |
| GO:0030509 | BMP signaling pathway | 108 | 26 | 10.11 | 5.00E-06 |
| GO:0070371 | ERK1 and ERK2 cascade | 183 | 37 | 17.13 | 5.10E-06 |
| GO:0150116 | regulation of cell-substrate junction organization | 60 | 18 | 5.62 | 5.40E-06 |
| GO:0045859 | regulation of protein kinase activity | 464 | 73 | 43.43 | 5.50E-06 |
| GO:0007178 | transmembrane receptor protein serine/threonine kinase signaling pathway | 265 | 48 | 24.81 | 5.50E-06 |
| GO:0043410 | positive regulation of MAPK cascade | 288 | 51 | 26.96 | 5.60E-06 |
| GO:0006950 | response to stress | 2599 | 302 | 243.28 | 5.80E-06 |
| GO:0048146 | positive regulation of fibroblast proliferation | 39 | 14 | 3.65 | 5.80E-06 |
| GO:0044706 | multi-multicellular organism process | 122 | 28 | 11.42 | 5.90E-06 |
| GO:0001952 | regulation of cell-matrix adhesion | 90 | 23 | 8.42 | 5.90E-06 |
| GO:0032989 | cellular component morphogenesis | 490 | 76 | 45.87 | 6.00E-06 |
| GO:0031175 | neuron projection development | 634 | 93 | 59.35 | 6.10E-06 |
| GO:0051893 | regulation of focal adhesion assembly | 55 | 17 | 5.15 | 6.30E-06 |
| GO:0090109 | regulation of cell-substrate junction assembly | 55 | 17 | 5.15 | 6.30E-06 |
| GO:0006957 | complement activation, alternative pathway | 5 | 5 | 0.47 | 7.10E-06 |
| GO:0009790 | embryo development | 785 | 110 | 73.48 | 7.30E-06 |
| GO:0045597 | positive regulation of cell differentiation | 577 | 86 | 54.01 | 7.40E-06 |
| GO:0050819 | negative regulation of coagulation | 30 | 12 | 2.81 | 7.40E-06 |

| | | | | | |
|---|---|---|---|---|---|
| GO:0007411 | axon guidance | 137 | 30 | 12.82 | 7.60E-06 |
| GO:0002042 | cell migration involved in sprouting angiogenesis | 45 | 15 | 4.21 | 7.80E-06 |
| GO:0018212 | peptidyl-tyrosine modification | 223 | 42 | 20.87 | 7.90E-06 |
| GO:0001667 | ameboidal-type cell migration | 323 | 55 | 30.23 | 8.10E-06 |
| GO:0072080 | nephron tubule development | 56 | 17 | 5.24 | 8.20E-06 |
| GO:0048518 | positive regulation of biological process | 4146 | 453 | 388.09 | 8.40E-06 |
| GO:0097485 | neuron projection guidance | 138 | 30 | 12.92 | 8.80E-06 |
| GO:0035556 | intracellular signal transduction | 1829 | 222 | 171.21 | 9.00E-06 |
| GO:0150115 | cell-substrate junction organization | 86 | 22 | 8.05 | 9.30E-06 |
| GO:0032956 | regulation of actin cytoskeleton organization | 270 | 48 | 25.27 | 9.40E-06 |
| GO:0007044 | cell-substrate junction assembly | 80 | 21 | 7.49 | 9.60E-06 |
| GO:0071772 | response to BMP | 112 | 26 | 10.48 | 1.00E-05 |
| GO:0071773 | cellular response to BMP stimulus | 112 | 26 | 10.48 | 1.00E-05 |
| GO:0050808 | synapse organization | 248 | 45 | 23.21 | 1.00E-05 |
| GO:0061564 | axon development | 318 | 54 | 29.77 | 1.10E-05 |
| GO:0048522 | positive regulation of cellular process | 3793 | 418 | 355.05 | 1.10E-05 |
| GO:0061572 | actin filament bundle organization | 126 | 28 | 11.79 | 1.10E-05 |
| GO:0010628 | positive regulation of gene expression | 714 | 101 | 66.83 | 1.20E-05 |
| GO:0003002 | regionalization | 197 | 38 | 18.44 | 1.20E-05 |
| GO:0048608 | reproductive structure development | 190 | 37 | 17.79 | 1.20E-05 |
| GO:0001501 | skeletal system development | 344 | 57 | 32.2 | 1.30E-05 |
| GO:0048568 | embryonic organ development | 281 | 49 | 26.3 | 1.30E-05 |
| GO:1901701 | cellular response to oxygen-containing compound | 769 | 107 | 71.98 | 1.30E-05 |
| GO:0060993 | kidney morphogenesis | 58 | 17 | 5.43 | 1.40E-05 |
| GO:0072028 | nephron morphogenesis | 47 | 15 | 4.4 | 1.40E-05 |
| GO:0061042 | vascular wound healing | 11 | 7 | 1.03 | 1.50E-05 |
| GO:0009410 | response to xenobiotic stimulus | 236 | 43 | 22.09 | 1.50E-05 |
| GO:0018108 | peptidyl-tyrosine phosphorylation | 221 | 41 | 20.69 | 1.50E-05 |
| GO:0019220 | regulation of phosphate metabolic process | 941 | 126 | 88.08 | 1.50E-05 |
| GO:0051174 | regulation of phosphorus metabolic process | 942 | 126 | 88.18 | 1.60E-05 |
| GO:0030193 | regulation of blood coagulation | 42 | 14 | 3.93 | 1.60E-05 |
| GO:1900046 | regulation of hemostasis | 42 | 14 | 3.93 | 1.60E-05 |
| GO:0007162 | negative regulation of cell adhesion | 178 | 35 | 16.66 | 1.70E-05 |
| GO:0061458 | reproductive system development | 193 | 37 | 18.07 | 1.80E-05 |
| GO:0001817 | regulation of cytokine production | 430 | 67 | 40.25 | 1.90E-05 |
| GO:0070527 | platelet aggregation | 48 | 15 | 4.49 | 1.90E-05 |
| GO:0150063 | visual system development | 231 | 42 | 21.62 | 1.90E-05 |
| GO:0150146 | cell junction disassembly | 15 | 8 | 1.4 | 2.00E-05 |
| GO:1900120 | regulation of receptor binding | 15 | 8 | 1.4 | 2.00E-05 |
| GO:0060055 | angiogenesis involved in wound healing | 19 | 9 | 1.78 | 2.10E-05 |
| GO:0043549 | regulation of kinase activity | 541 | 80 | 50.64 | 2.10E-05 |
| GO:0002683 | negative regulation of immune system process | 247 | 44 | 23.12 | 2.10E-05 |
| GO:0051781 | positive regulation of cell division | 54 | 16 | 5.05 | 2.10E-05 |
| GO:0051963 | regulation of synapse assembly | 54 | 16 | 5.05 | 2.10E-05 |
| GO:0055006 | cardiac cell development | 54 | 16 | 5.05 | 2.10E-05 |
| GO:0048880 | sensory system development | 232 | 42 | 21.72 | 2.10E-05 |
| GO:2000146 | negative regulation of cell motility | 217 | 40 | 20.31 | 2.20E-05 |
| GO:0007565 | female pregnancy | 110 | 25 | 10.3 | 2.20E-05 |
| GO:0006468 | protein phosphorylation | 1059 | 138 | 99.13 | 2.30E-05 |
| GO:0051017 | actin filament bundle assembly | 124 | 27 | 11.61 | 2.40E-05 |
| GO:0045321 | leukocyte activation | 528 | 78 | 49.42 | 2.70E-05 |
| GO:0060548 | negative regulation of cell death | 702 | 98 | 65.71 | 2.80E-05 |
| GO:0033002 | muscle cell proliferation | 153 | 31 | 14.32 | 2.80E-05 |
| GO:0033674 | positive regulation of kinase activity | 313 | 52 | 29.3 | 2.90E-05 |
| GO:0048041 | focal adhesion assembly | 73 | 19 | 6.83 | 2.90E-05 |
| GO:0006936 | muscle contraction | 175 | 34 | 16.38 | 2.90E-05 |
| GO:0043542 | endothelial cell migration | 175 | 34 | 16.38 | 2.90E-05 |
| GO:0010942 | positive regulation of cell death | 428 | 66 | 40.06 | 3.00E-05 |
| GO:0009968 | negative regulation of signal transduction | 910 | 121 | 85.18 | 3.10E-05 |
| GO:0098657 | import into cell | 119 | 26 | 11.14 | 3.10E-05 |
| GO:1902531 | regulation of intracellular signal transduction | 1215 | 154 | 113.73 | 3.20E-05 |
| GO:0006956 | complement activation | 20 | 9 | 1.87 | 3.50E-05 |
| GO:0010092 | specification of animal organ identity | 20 | 9 | 1.87 | 3.50E-05 |

| | | | | | |
|---|---|---|---|---|---|
| GO:0001816 | cytokine production | 439 | 67 | 41.09 | 3.60E-05 |
| GO:0003013 | circulatory system process | 332 | 54 | 31.08 | 3.70E-05 |
| GO:0010812 | negative regulation of cell-substrate adhesion | 45 | 14 | 4.21 | 3.80E-05 |
| GO:0050818 | regulation of coagulation | 45 | 14 | 4.21 | 3.80E-05 |
| GO:0001654 | eye development | 230 | 41 | 21.53 | 3.90E-05 |
| GO:0050900 | leukocyte migration | 200 | 37 | 18.72 | 4.00E-05 |
| GO:0060326 | cell chemotaxis | 156 | 31 | 14.6 | 4.10E-05 |
| GO:0048519 | negative regulation of biological process | 3667 | 401 | 343.25 | 4.30E-05 |
| GO:0051917 | regulation of fibrinolysis | 9 | 6 | 0.84 | 4.30E-05 |
| GO:0010632 | regulation of epithelial cell migration | 179 | 34 | 16.76 | 4.70E-05 |
| GO:0050730 | regulation of peptidyl-tyrosine phosphorylation | 150 | 30 | 14.04 | 4.80E-05 |
| GO:0046649 | lymphocyte activation | 443 | 67 | 41.47 | 4.90E-05 |
| GO:0040013 | negative regulation of locomotion | 240 | 42 | 22.47 | 4.90E-05 |
| GO:0042698 | ovulation cycle | 46 | 14 | 4.31 | 5.00E-05 |
| GO:0072088 | nephron epithelium morphogenesis | 46 | 14 | 4.31 | 5.00E-05 |
| GO:0014812 | muscle cell migration | 70 | 18 | 6.55 | 5.50E-05 |
| GO:0071675 | regulation of mononuclear cell migration | 70 | 18 | 6.55 | 5.50E-05 |
| GO:1902533 | positive regulation of intracellular signal transduction | 661 | 92 | 61.87 | 5.50E-05 |
| GO:0001709 | cell fate determination | 21 | 9 | 1.97 | 5.50E-05 |
| GO:0043065 | positive regulation of apoptotic process | 378 | 59 | 35.38 | 5.60E-05 |
| GO:0050866 | negative regulation of cell activation | 116 | 25 | 10.86 | 5.60E-05 |
| GO:0002682 | regulation of immune system process | 842 | 112 | 78.82 | 6.10E-05 |
| GO:0033993 | response to lipid | 558 | 80 | 52.23 | 6.30E-05 |
| GO:0010755 | regulation of plasminogen activation | 13 | 7 | 1.22 | 6.40E-05 |
| GO:0042730 | fibrinolysis | 13 | 7 | 1.22 | 6.40E-05 |
| GO:0003156 | regulation of animal organ formation | 17 | 8 | 1.59 | 6.50E-05 |
| GO:0044057 | regulation of system process | 291 | 48 | 27.24 | 6.90E-05 |
| GO:0032609 | interferon-gamma production | 53 | 15 | 4.96 | 6.90E-05 |
| GO:0032649 | regulation of interferon-gamma production | 53 | 15 | 4.96 | 6.90E-05 |
| GO:0032147 | activation of protein kinase activity | 97 | 22 | 9.08 | 6.90E-05 |
| GO:0090287 | regulation of cellular response to growth factor stimulus | 228 | 40 | 21.34 | 6.90E-05 |
| GO:0046620 | regulation of organ growth | 65 | 17 | 6.08 | 7.00E-05 |
| GO:0060419 | heart growth | 65 | 17 | 6.08 | 7.00E-05 |
| GO:0007010 | cytoskeleton organization | 1047 | 134 | 98.01 | 7.20E-05 |
| GO:0120036 | plasma membrane bounded cell projection organization | 1010 | 130 | 94.54 | 7.20E-05 |
| GO:0022407 | regulation of cell-cell adhesion | 276 | 46 | 25.84 | 7.50E-05 |
| GO:0043405 | regulation of MAP kinase activity | 132 | 27 | 12.36 | 7.60E-05 |
| GO:0048645 | animal organ formation | 42 | 13 | 3.93 | 7.60E-05 |
| GO:0009913 | epidermal cell differentiation | 98 | 22 | 9.17 | 8.20E-05 |
| GO:0008406 | gonad development | 147 | 29 | 13.76 | 8.30E-05 |
| GO:0090092 | regulation of transmembrane receptor protein serine/threonine kinase signaling pathway | 199 | 36 | 18.63 | 8.30E-05 |
| GO:0060562 | epithelial tube morphogenesis | 230 | 40 | 21.53 | 8.50E-05 |
| GO:0046579 | positive regulation of Ras protein signal transduction | 37 | 12 | 3.46 | 8.60E-05 |
| GO:0043068 | positive regulation of programmed cell death | 384 | 59 | 35.94 | 8.80E-05 |
| GO:0043069 | negative regulation of programmed cell death | 625 | 87 | 58.5 | 8.80E-05 |
| GO:0014909 | smooth muscle cell migration | 60 | 16 | 5.62 | 8.80E-05 |
| GO:0048523 | negative regulation of cellular process | 3315 | 364 | 310.3 | 9.10E-05 |
| GO:0050865 | regulation of cell activation | 368 | 57 | 34.45 | 9.20E-05 |
| GO:0043406 | positive regulation of MAP kinase activity | 79 | 19 | 7.39 | 9.30E-05 |
| GO:2000027 | regulation of animal organ morphogenesis | 79 | 19 | 7.39 | 9.30E-05 |
| GO:0030239 | myofibril assembly | 32 | 11 | 3 | 9.30E-05 |
| GO:0030336 | negative regulation of cell migration | 208 | 37 | 19.47 | 9.50E-05 |
| GO:0043066 | negative regulation of apoptotic process | 609 | 85 | 57.01 | 9.70E-05 |
| GO:0048705 | skeletal system morphogenesis | 141 | 28 | 13.2 | 9.70E-05 |
| GO:0110053 | regulation of actin filament organization | 216 | 38 | 20.22 | 9.80E-05 |
| GO:0030324 | lung development | 134 | 27 | 12.54 | 9.90E-05 |
| GO:0031638 | zymogen activation | 43 | 13 | 4.03 | 1.00E-04 |
